# Coupling of oceanographic state to the dark proteome: a foundation for genome-informed marine productivity modeling

**DOI:** 10.64898/2026.09.23.753693

**Authors:** David Roy Nelson, Maxence Plouviez, Sarah Daakour, Ashish Jaiswal, Weiqi Fu, Shady A. Amin, Kourosh Salehi-Ashtiani

## Abstract

Phytoplankton drive *∼*46% of global primary production, yet whether their protein repertoires encode ocean conditions—and whether reference databases capture the environmentally responsive fraction—is unknown. We used a protein language model to classify 447.7 million proteins from 2,357 samples and analyzed domain profiles for 231.7 million algal proteins across 2,044 samples. Under spatial block cross-validation, environment predicted individual domain abundances at *R*^2^ up to 0.59, while domain profiles predicted sea surface temperature at *R*^2^ = 0.38. The strongest coupling lay beyond annotated sequence space: 33,950 families clustered from 201 million Pfam-dark proteins coupled to environment 2.29-fold more strongly than Pfam domains. Cross-taxonomic selection, protein language modeling, and AlphaFold 3 predictions of 138 well-folded, InterPro-unannotated representatives support the Pfam-dark proteome as a biologically structured evolutionary compartment. Thus, the proteins most tightly coupled to ocean state are those least represented in reference databases.

**GRAPHICAL ABSTRACT:** Language modeling with AI for algal amino acid Sequence Representation (LA^4^SR) classified 447.7 million proteins from a 2,357-sample input collection; the 2,044-sample domain-analysis set contained 231.7 million proteins classified as algal. Protein domain composition and ocean state predict each other, and 33,950 Pfam-dark protein families show 2.29-fold stronger environmental coupling than Pfam domains.

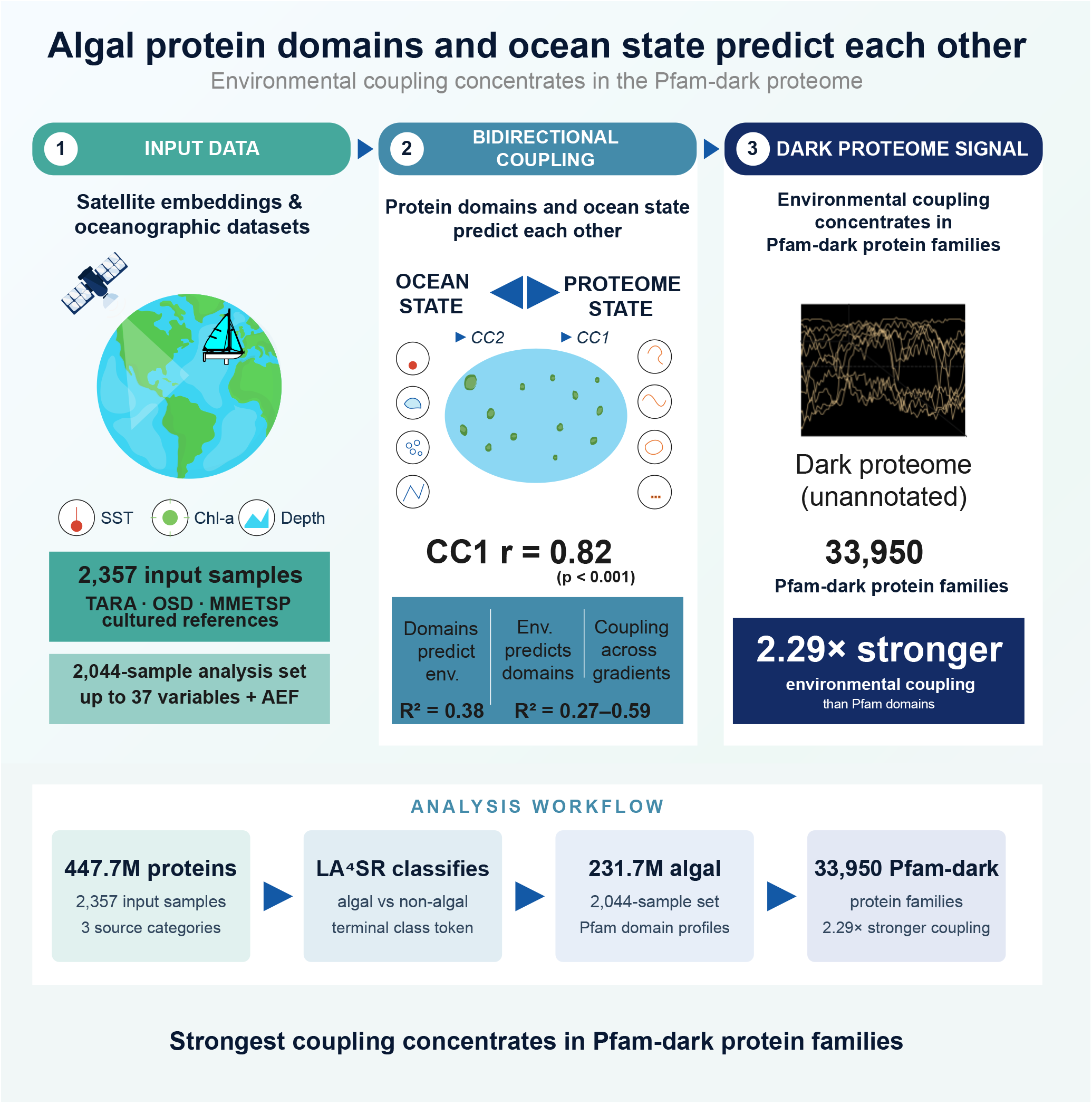

## INTRODUCTION

Marine phytoplankton account for roughly half of global net primary productivity,^1^ yet their genomes and environments are usually studied separately. Satellite remote sensing measures ocean state globally,^2^ while ocean metagenomics has generated billions of predicted proteins across major basins.^3–6^ Protein language models can now classify these sequences from learned amino acid representations without sequence homology.^7,8^ Whether environmental state and protein domain composition predict each other has not been tested at this scale. Here, “coupling” denotes out-of-sample predictability, not direct causation.

Protein repertoire helps shape community productivity, yet satellite-based productivity models do not directly represent the functional capacity of the organisms responsible for carbon fixation. Genome-scale metabolic models offer one route,^9^ but require organism-specific curation that does not scale to thousands of uncultured taxa. A domain-level link between protein repertoire and environment could provide a more scalable genomic layer.

Two barriers have prevented a direct test. Functional annotation is incomplete even in model microalgae and poorer in non-model species. Extracting eukaryotic algal signals from mixed metagenomes is also difficult: marker-gene and co-abundance approaches^10^ face scaling and fragmentation limits, and homology-independent extraction has only recently become feasible.

Natural selection can filter functional space:^11^ environmental conditions constrain viable protein repertoires, potentially creating learnable structure between genomes and co-located environments. Neutral processes, dispersal limitation, and history also shape communities. We therefore ask whether detectable coupling exists and whether it is concentrated in accessory and niche-specific rather than universally conserved domains.

Protein family (Pfam) domains provide a cross-lineage vocabulary of modular functional units.^12^ Environmental measurements can be complemented by AlphaEarth Foundations (AEF) satellite embeddings,^13^ which compress multispectral imagery into 64-dimensional descriptors. Together, these representations enable a direct test of domain–environment coupling.

TARA Oceans studies linked community composition to carbon export and environmental gradients to phytoplankton physiology,^9^ diversity,^6^ and gene expression.^14^ Global prokaryotic gene– environment associations are also established.^15^ Bidirectional predictability between protein-domain composition and environmental state, however, remains untested.

Here we test this directly. We apply Language modeling with AI for algal amino acid Sequence Representation (LA^4^SR),^16^ a transformer-based protein language model, to extract algal sequences from ocean metagenomes (TARA Oceans and Ocean Sampling Day [OSD]^17^), Marine Microbial Eukaryote Transcriptome Sequencing Project (MMETSP) transcriptomes,^18^ and cultured reference proteomes. Each sample is paired with 37 interpretable oceanographic variables (Google Earth Engine and World Ocean Atlas 2023 [WOA23] nutrients) and 64-dimensional AEF satellite embeddings.^13^ We then ask: do protein domain composition and environmental state predict each other?

Environment and domain composition predicted each other. Canonical correlation analysis (CCA) identified a dominant component (CC1; canonical correlation *r* = 0.82) associated with sea surface temperature (SST), while CC2–CC3 captured chlorophyll and ocean color. The strongest associations concentrated in 33,950 Pfam-dark families, whose biological sequence organization we assessed by cross-taxonomic selection, protein language modeling, and structure prediction. We focus on eukaryotic microalgae because LA^4^SR enables their homology-independent extraction from mixed metagenomes; matched prokaryotic domain-level coupling remains outside this study.

## RESULTS

### A protein language model recovers *∼*44,000–86,000 algal taxa from a global marine and reference collection

Testing whether protein domain composition and ocean state predict each other requires first extracting eukaryotic algal sequences from mixed microbial metagenomes, a task for which homology-based methods discard the *∼*75% of algal sequences lacking database matches (Table S1D). We applied LA^4^SR,^16^ a transformer-based protein language model, to a 2,357-sample collection of ocean metagenomes, marine eukaryotic transcriptomes, and cultured reference proteomes.

LA^4^SR classified 447.7 million predicted proteins. Of the 2,357 input samples, 2,044 had non-empty Pfam-A results and entered the domain-analysis set, which contained 231.7 million proteins classified as algal (51.8% of classifier inputs; Table S1). The input collection comprised 1,203 ocean metagenome assemblies (1,063 TARA Oceans^3^ plus 140 Ocean Sampling Day;^17^ all size fractions, no pre-filtering), 676 MMETSP transcriptomes, and 478 cultured reference proteomes.

We annotated the retained proteins by profile hidden Markov model (HMM) search (hmm-search)^19^ against the curated Pfam-A domain database.^12^ We paired georeferenced samples with 37 interpretable oceanographic variables^2,20,21^ and 64-dimensional AEF satellite embeddings. Spearman correlation screened domain–environment associations; XGBoost^22^ tested out-of-sample predictability, and SHAP (SHapley Additive exPlanations)^23^ quantified each feature’s contribution to the predictions (Methods).

The geographic coverage and basin-level source composition of the 1,810 Global Positioning System (GPS)-mapped samples are shown in Figure 1A–B.

**Figure 1:**
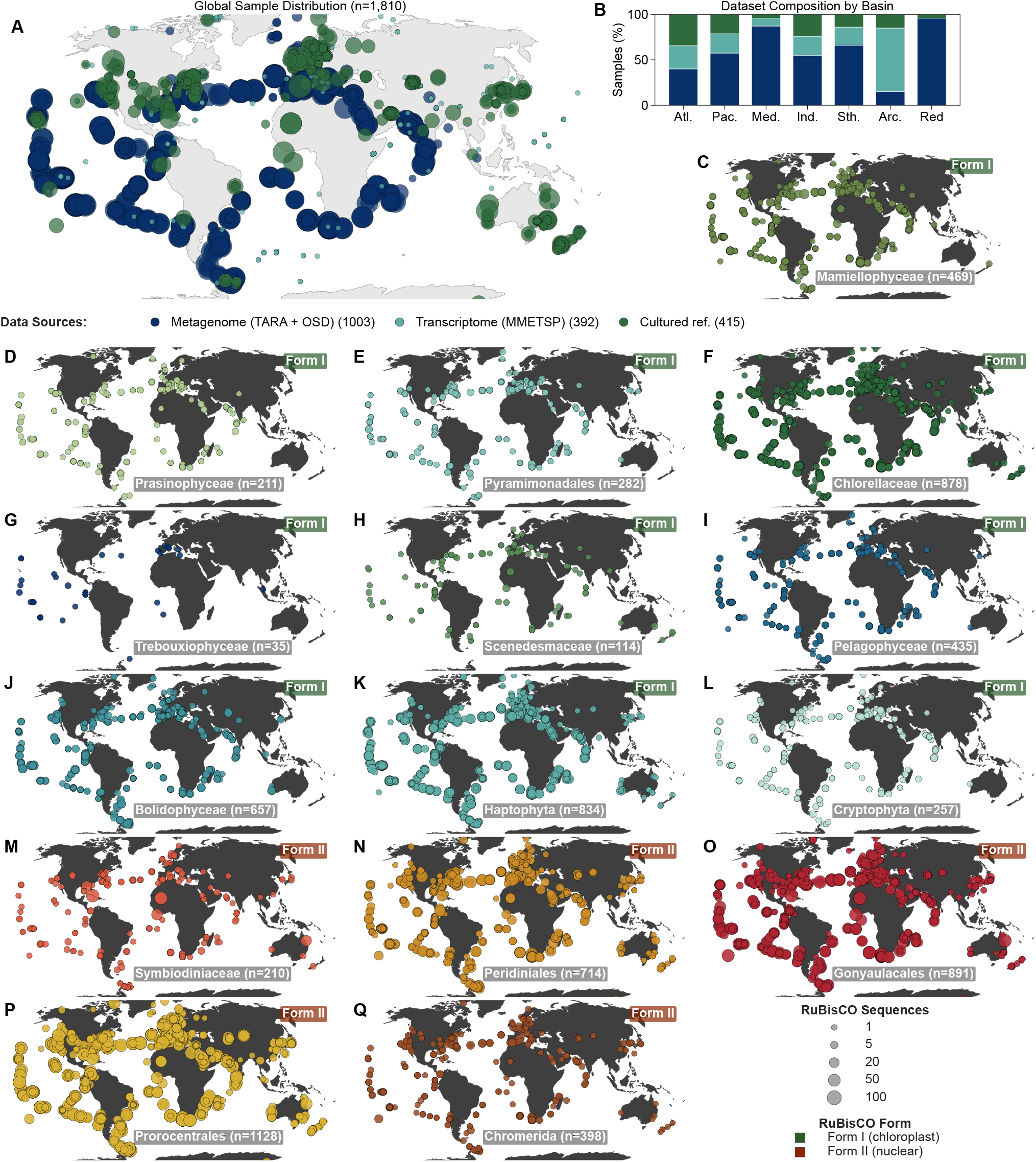
Global sample coverage, dataset composition, and algal lineage distributions. **(A)** Global distribution of 1,810 Global Positioning System (GPS)-mapped samples across the three source categories (ocean metagenomes, Marine Microbial Eukaryote Transcriptome Sequencing Project [MMETSP] single-organism transcriptomes from cultured isolates, and cultured reference proteomes), colored by category (Table S1). Marker area scales with the percentile rank of total protein count per sample. MMETSP transcriptomes are not environmental metatranscriptomes; they are mapped to the geographic origin of each culture strain. **(B)** Source-category composition per ocean basin (stacked percentage bars). **(C–L)** Geographic distribution of ten algal lineages identified via Form I ribulose-1,5-bisphosphate carboxylase/oxygenase (RuBisCO) hidden Markov models (HMMs; chloroplast-encoded); marker size proportional to sequence count. Lineage names and sample counts shown per panel. **(M–Q)** Geographic distribution of five myzozoan lineages via Form II RuBisCO HMMs (nuclear-encoded).^24^ Marker size proportional to sequence count. Robinson projection; Form I in cool tones, Form II in warm tones.

Within the 2,044-sample analysis set, 1,005 ocean metagenomes contributed 178.4 million algal sequences, 606 MMETSP transcriptomes contributed 18.3 million, and 433 reference proteomes contributed 35.0 million (Table S1). The resulting 231.7-million-sequence catalog increases sample coverage 5.3-fold over the Marine Atlas of Tara Oceans Unigenes (MATOU).^5^

Classification used LA^4^SR’s algaGPT decoding mode, in which top-*k* sampling generates the terminal token that encodes the algal or non-algal class (Methods). Retention scales with source purity (cultured references 93.0%, MMETSP 73.9%, metagenomes 61.8%; Table S1). In the 352.6-million-protein Data S8 comparison subset, non-algal-classified sequences were 2.9-fold more likely to have homologs in the National Center for Biotechnology Information (NCBI) non-redundant (NR) protein database than algal-classified ones (73.2% vs. 25.1%), supporting preferential retention of database-underrepresented algal sequences.^16^ Restricting subsequent analyses to metagenomes alone or to the algal-dominated 20–180 µm size fraction retained or strengthened coupling (see below; Supplemental Text).

Of the 2,044 analysis-set samples, 1,810 (88.6%) had both valid Global Positioning System (GPS) coordinates and named basin assignments in the frozen basin table, spanning seven ocean basins dominated by the Atlantic (644) and Pacific (617; Figure 1A). A subsequent GPS-recovery pass added coordinates for 68 samples that lacked a basin assignment in that table, yielding the 1,878-sample satellite-only validation set used below; basin-resolved analyses retained the frozen 1,810-sample set. Depth distributions are surface-biased (median 5 m), consistent with euphotic-zone-focused sampling; environmental variable coverage ranges from 49–94% depending on the variable (Table S1).

To independently assess whether the retained proteins recover plausible algal community structure, we searched for RuBisCO large-subunit markers from 15 algal lineages (10 Form I and 5 Form II myzozoan; Figure 1C–Q; Figure S1A–C). The lineage-specific HMMs detected 149,733 unique sequences (Table S3; Data S4) and RuBisCO in 92.9% (1,118) of the 1,203 ocean metagenome assemblies (TARA Oceans and OSD), with Chlorellaceae and Haptophyta the most widely detected Form I lineages among GPS-mapped samples (Figure 1C–L). These rank-order prevalences agree with 18S ribosomal DNA (rDNA) metabarcoding of the same samples,^6^ supporting recovery of ecologically realistic lineage composition (Supplemental Text).

Because Form I rbcL is typically single-copy^25^ while dinoflagellate Form II is multi-copy (2–6 per genome^24,26^), copy-number correction brackets the taxon estimate at *∼*44,000–86,000 photosynthetic algal taxa (Supplemental Text), providing the statistical power required to test domain–environment coupling across 7 ocean basins while retaining the *∼*75% of algal sequences that BLAST-based pipelines discard (Table S1D). These results establish that existing marine metagenome collections contain a large, previously inaccessible algal proteome; protein language models now make it tractable for domain-level analysis across ocean basins.

### Satellite imagery and ocean climatologies define the environmental axis

Of the 1,810 GPS-mapped analysis-set samples, 1,809 satisfied the completeness criteria for the raw-environment matrix (Table S1C; Methods). We described each sampling site with 37 interpretable oceanographic variables (SST, chlorophyll-a, bathymetry, ocean color, nutrients, dissolved oxygen, mixed layer depth; compiled from Google Earth Engine^2^ [GEE], WOA23,^20^ and mixed-layer-depth [MLD] climatology^21^) plus 64-dimensional AEF satellite embeddings.^13^ AEF embeddings derive from annual satellite embeddings for 2017 onward (mosaicked across the available annual layers at extraction), whereas samples were collected in 2009–2014; associations therefore reflect long-term spatial environmental state rather than contemporaneous conditions. Temporal mismatch may attenuate relationships for variable features and precludes contemporaneous interpretation; correlations are 2.4-fold stronger for stable variables (median *|ρ|* = 0.14) than variable ones (0.06; *p <* 10^−16^; Data S1; Supplemental Text).

Of these, 995 (55.0%) had AEF records: 969 contained no zero-filled dimensions and 26 were partially zero-filled during feature preparation (coastal coverage *>*90% vs. open-ocean *∼*30%; Table S1). AEF–GEE correlation showed that the embeddings capture oceanographically meaningful variation (44.1% of dimension–variable pairs significant after false-discovery-rate [FDR] correction; max *|ρ|* = 0.68; Data S1; Data S5).

The 1,809-sample dataset with raw environmental variables was used for the two-view CCA and bidirectional modeling (Figure 2B–C, E–F; Figure S2), while the 995-sample AEF subset was used for three-view generalized CCA (GCCA; Figure 2A, D; Figure S2H) and embedding-dimension correlation analysis (Figure S3F–G; Table S1C). This coastal and shelf bias limits generalizability to deep-ocean and pelagic environments. With the genomic and environmental axes now defined at the same locations, we next tested whether they are coupled.

**Figure 2:**
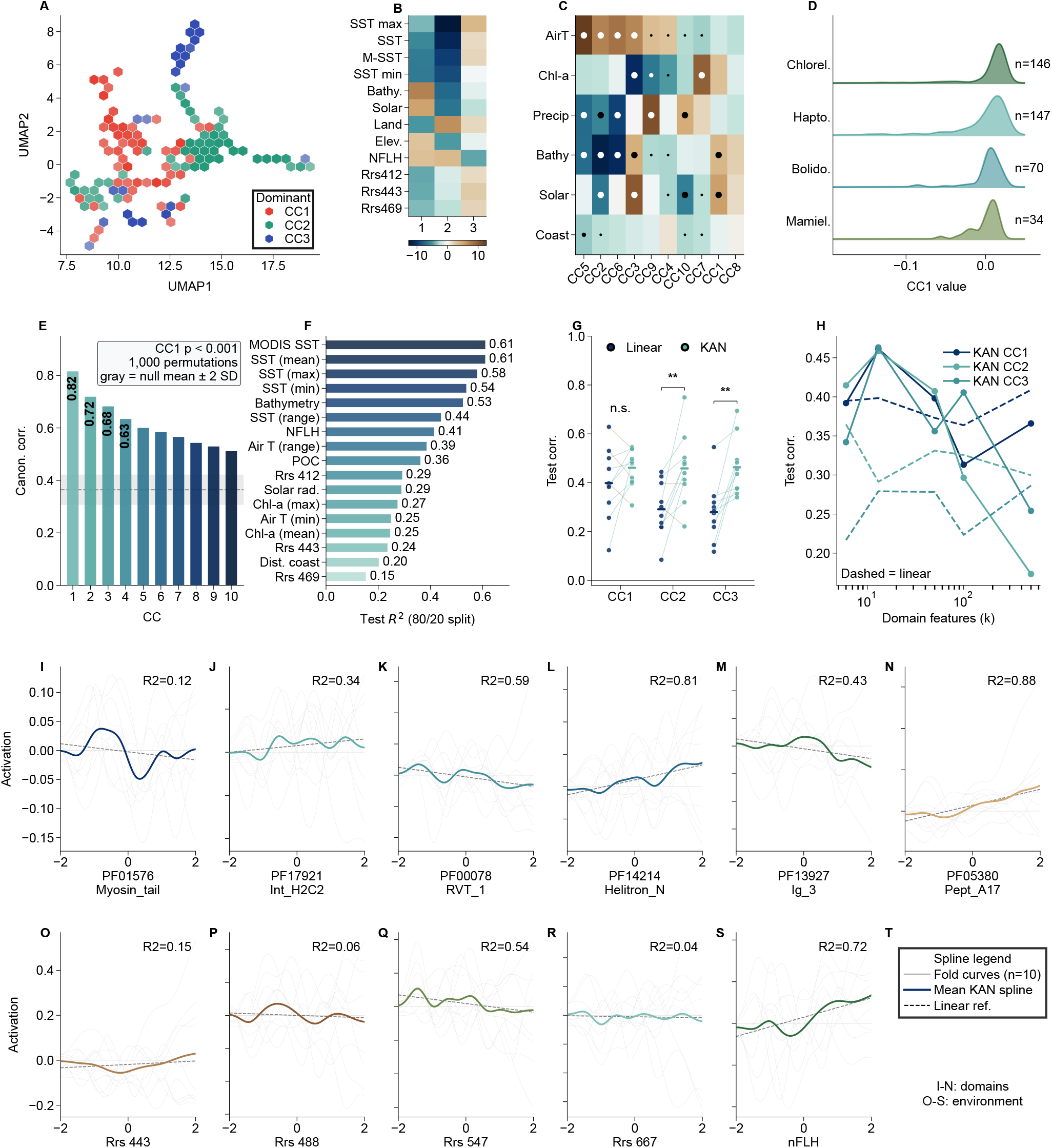
Canonical correlation manifold, environment–genome associations, and canonical correlation analysis (CCA) nonlinearity. **(A)** Uniform Manifold Approximation and Projection (UMAP) of ten three-view generalized CCA (GCCA) components derived from AlphaEarth Foundations embeddings, raw environmental variables, and Pfam profiles (*n* = 995). After removal of 68 interquartile-range (IQR) outliers, 927 samples were retained for visualization. Hexagons are colored by the majority dominant component (red, CC1; teal, CC2; blue, CC3); saturation encodes the mean gap between the two largest component magnitudes. **(B)** Environmental loadings on CC1–CC3 from the two-view environment–Pfam CCA (*n* = 1,809; top 12 variables by absolute loading). Sea surface temperature (SST) variants are collinear (variance inflation factor [VIF] *>* 10; Table S8; Supplemental Text); loadings within collinear groups should be interpreted collectively. **(C)** Pearson correlations between six environmental variables and ten components from the two-view CCA (*n* = 1,809), hierarchically clustered. Dot size encodes significance (*p <* 0.05, *p <* 0.01, or *p <* 0.001); color shows *r* on a −0.6 to 0.6 scale. **(D)** GCCA CC1 density by dominant RuBisCO lineage (*n* shown for each lineage). **(E)** Two-view observed canonical correlations versus a standard row-permutation null (*n* = 1,809; 1,000 shuffles; gray band, null mean 2 standard deviations [SD]); CC1 *p <* 0.001. The prespecified spatial-block permutation result is reported in the Results, Data S1, and Supplemental Text. **(F)** Held-out 20% test-set *R*^2^ for reverse models (Pfam environment) with *R*^2^ *>* 0.1 (17 of 29 targets), using target-specific complete cases (*n* = 146–1,279 per target) and the same fixed 80/20 split as Figure 3E; 95% bootstrap CIs are shown in Figure 3E. **(G)** Kolmogorov–Arnold network CCA (KAN-CCA) versus linear CCA test correlations across ten spatial CV folds in the satellite-only analysis (*n* = 969; 19 environmental variables; *k* = 13 sparse domain features). Points are fold values, horizontal ticks are means, and lines pair methods within fold. Two-sided paired *t* -tests: \*\**p <* 0.01; not significant (n.s.), *p ≥*0.05. **(H)** Mean test correlation across the same ten folds as a function of domain-feature sparsity: KAN (solid) versus linear (dashed) CCA for CC1–CC3. **(I–N)** Learned B-spline activation functions for the six top-weighted domain features (KAN-CCA, CC1). Gray: individual folds; colored: mean. *R*^2^ = linearity of spline (lower = more nonlinear). **(O–S)** Learned B-spline activations for the five top-weighted environmental features. **(T)** Line-style key for panels I–S (gray, individual folds; colored, mean KAN spline; dashed, linear reference).

### Domain–environment associations are pervasive and environmentally structured

If domain composition and environmental state share statistical structure, the first signature should be pervasive, albeit individually weak, covariation across hundreds of thousands of domain–dimension pairs. Spearman correlation analysis tested 761,472 associations between 11,898 Pfam domains and 64 AEF dimensions (*n* = 995; algaGPT extraction). At Benjamini-Hochberg FDR *<* 0.05,^27^ 279,575 associations (36.7%) passed the FDR threshold, indicating widespread statistical covariation (Figure S3F). XGBoost regressors confirm that these correlations encode learnable structure: models predicting embedding dimensions from Pfam composition reached *R*^2^ up to 0.17 in five-fold cross-validation (*n* = 995; Figure S3G), with SHAP analysis identifying consistent top features across dimensions. A centered log-ratio (CLR) sensitivity analysis confirmed that broader significance counts are comparable between normalizations, though the top-ranked domains differ (Jaccard = 0.21 at top-50; Supplemental Text); all downstream modeling uses CLR-transformed inputs (Methods; Data S1).

### Temperature dominates the leading axis of domain–environment coupling

The 11,898 *×* 64 domain–environment association matrix (Spearman *ρ* for each Pfam domain against each AEF dimension) reveals that associations cluster by environmental axis rather than distributing uniformly across dimensions. Uniform Manifold Approximation and Projection (UMAP)^28^ of the permissive Pfam-A v37.2 abundance matrix (*E <* 10^−5^; 20,318 raw feature columns, 1,810 GPS-mapped samples; Figure S1D–K) confirms this: samples separate along temperature gradients in protein composition space, a pattern robust across UMAP neighbor counts and t-distributed Stochastic Neighbor Embedding (t-SNE) perplexity values (Figure S1D– K). The domain–environment correlation structure underlying this separation is quantified in Figure S3F–G; underlying correlation tables and embedding coordinates are in Data S1, and the AEF embedding matrix is in Data S5.

Canonical correlation analysis (CCA)^29^ across 1,809 samples (Figure 2) yielded CC1 (*r* = 0.82), CC2 (*r* = 0.72), and CC3 (*r* = 0.68). The prespecified spatial-block permutation test supported CC1 (*p <* 0.001; *z* = 7.3), which showed split-half reproducibility *r* = 0.99 (9.7% shrinkage; Data S1; Supplemental Text); CC2 and CC3 were not tested with the spatial-block permutation procedure.

SST variables dominate CC1 loadings while ocean color and chlorophyll variables load on CC2– CC3 (Figure 2B–C). Figure 2E shows the observed canonical correlations against the standard row-permutation null; the spatial-block test was limited to CC1. Ridge density plots of CC1 by dominant RuBisCO lineage revealed lineage-specific positioning: Haptophyta-dominated samples shifted toward negative CC1 values relative to Chlorellaceae (Figure 2D), linking the canonical structure to taxonomic turnover. Three of the top 10 SHAP-ranked domains appeared among the top 7 CC1 loadings (Figure S2C–D), indicating convergent identification by supervised and unsupervised frameworks.

Given the nonlinear nature of ecological processes, is the dominant domain–environment coupling itself nonlinear? A satellite-only sparse CCA and Kolmogorov–Arnold network CCA (KAN-CCA)^30^ analysis (*n* = 969; 19 environmental variables) tested this: counter to expectation, KAN-CCA did not improve CC1 at the prespecified 13-domain sparsity level (*p* = 0.201), while secondary variates contained nonlinear structure (CC2 +57.4%, *q* = 0.012; CC3 +65.9%, *q* = 0.006; exploratory; Supplemental Text; Figure 2G–H; Data S6). A separate nutrient-expanded sparse CCA (*n* = 786; 24 variables) recovered a leading temperature–oxygen axis (Supplemental Text; Data S6). Together, these analyses indicate that the dominant coupling is largely linear at the scale of global ocean gradients, with nonlinear structure concentrated in secondary axes.

Of the 13 highest-weighted sparse CCA domains on CC1 in the satellite-only analysis (Data S6), 12 encode transposable element (TE) machinery, converging with the supervised finding that 11 of 18 most predictable domains are mobile genetic elements. TE dominance likely reflects dinoflagellate genome architecture,^31^ but lineage-resolved partial correlations indicate that TE– environment coupling persists beyond pure taxonomic sorting (Supplemental Text).

### Environment predicts which domains are present, and mobile elements dominate

Domain–environment covariation predicts held-out samples in both directions under 10-fold spatial block cross-validation (CV; 2° grid cells; Data S1; Figures 3–5): genome-to-environment (*R*^2^ = 0.38 for SST) and environment-to-genome (*R*^2^ up to 0.59). The environment-to-genome direction is the stringent test, since predicting individual domain abundances from a few dozen environmental variables is dimensionally harder than predicting one target from *∼*10,000 domains. Pfam composition contributes unique variance beyond a nonlinear latitude-only baseline for all four flagship targets (partial *R*^2^: SST 0.27, bathymetry 0.55, nitrate 0.21, dissolved oxygen 0.21; Supplemental Text).

**Figure 3:**
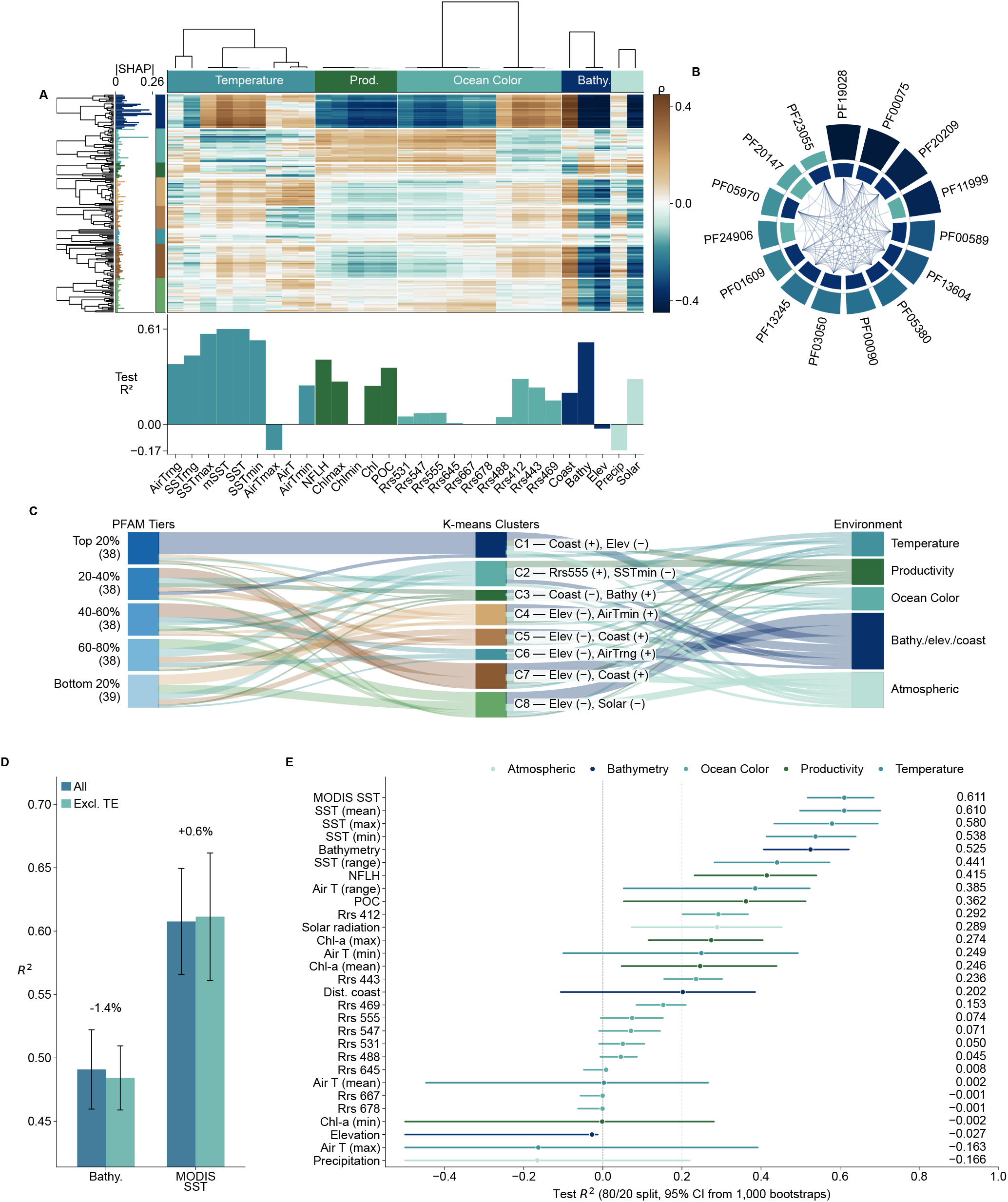
Modular organization of domain–environment coupling. **(A)** K-means biclustered heatmap of Spearman *ρ*: 384 Pfam domains (mean normalized SHapley Additive exPlanations [SHAP] importance *≥*5% of maximum) clustered at *k* = 8 across 29 environmental variables; up to 30 domains per cluster are displayed (191 rows). Left marginal, mean SHAP importance; bottom, reverse-model test-set *R*^2^ per variable (same estimates as panel E; negative values below the zero line). **(B)** Top 15 domains by mean SHAP importance. Inner ring, cluster membership; outer ring, SHAP magnitude. **(C)** Sankey diagram from SHAP-importance quintiles through clusters to environmental categories. Tier-to-cluster widths count domains; cluster-to-environment widths are rescaled mean absolute Spearman associations, with outgoing flow normalized to each cluster’s incoming count. Cluster labels give the two variables with the largest absolute z-scored centroid (sign from mean *ρ*). The Bathy./elev./coast category comprises bathymetry, elevation, and distance to coast; the Atmospheric category comprises solar radiation and precipitation (air-temperature variables are grouped with Temperature; the same categories are used in panels A, C, and E). **(D)** Five-fold CV sensitivity of reverse-model *R*^2^ to exclusion of 23 transposable-element (TE) domains (*n* = 1,279). Bars show fold mean SD for the full and TE-excluded feature sets; labels show relative percent change. The y-axis is truncated to resolve the small changes. **(E)** Reverse-model test-set *R*^2^ for target-specific complete cases under a fixed 80/20 split. Points are test-set estimates and horizontal lines are 95% CIs from 1,000 bootstrap resamples of held-out predictions (CI lower bounds below 0.5 are truncated at 0.5: Chl-a min, elevation, air temperature max, precipitation); colors indicate environmental category.

**Figure 4:**
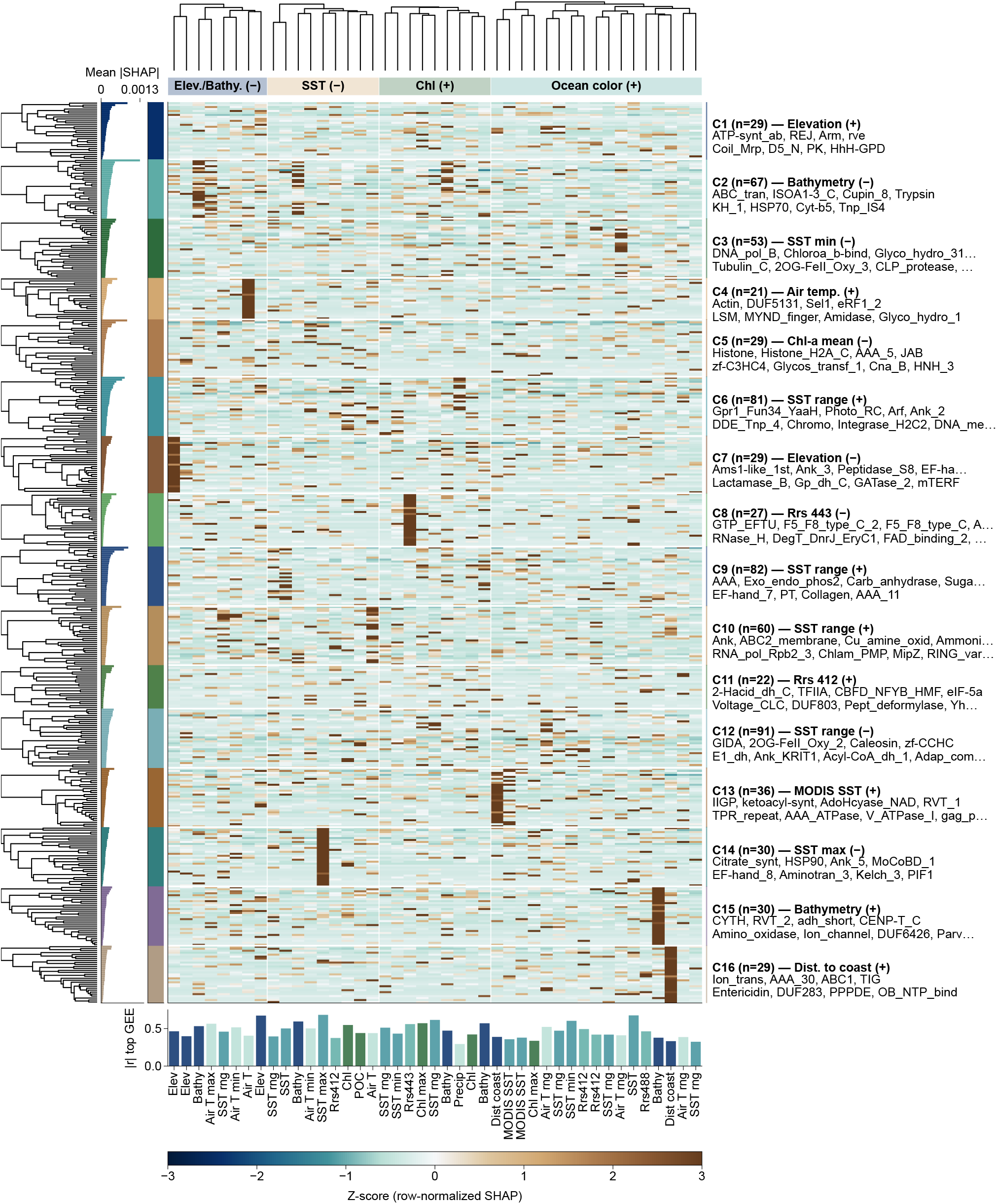
Pfam–environment SHapley Additive exPlanations (SHAP) landscape across AlphaEarth Foundations (AEF) embedding dimensions. K-means was fitted to 716 Pfam domains across all 64 AEF dimensions (*n* = 995 samples; *k* = 16). The heatmap displays up to 30 domains with the largest mean SHAP per cluster (456 total) and 43 dimensions assigned to four interpretable groups (elevation/bathymetry, sea surface temperature [SST], chlorophyll, and ocean color). Cluster *n* labels report membership in the full 716-domain clustering. Values are z-scored SHAP importance. Left marginal, mean SHAP and cluster assignment; each cluster is labeled by its dominant environmental association and top domains. Clusters span SST (7 clusters), bathymetry/elevation (4), chlorophyll (1), ocean color (2), air temperature (1), and coastal distance (1). Bottom, absolute Pearson *r* between each displayed AEF dimension and its most-correlated Google Earth Engine (GEE) variable, colored by environmental category.

**Figure 5:**
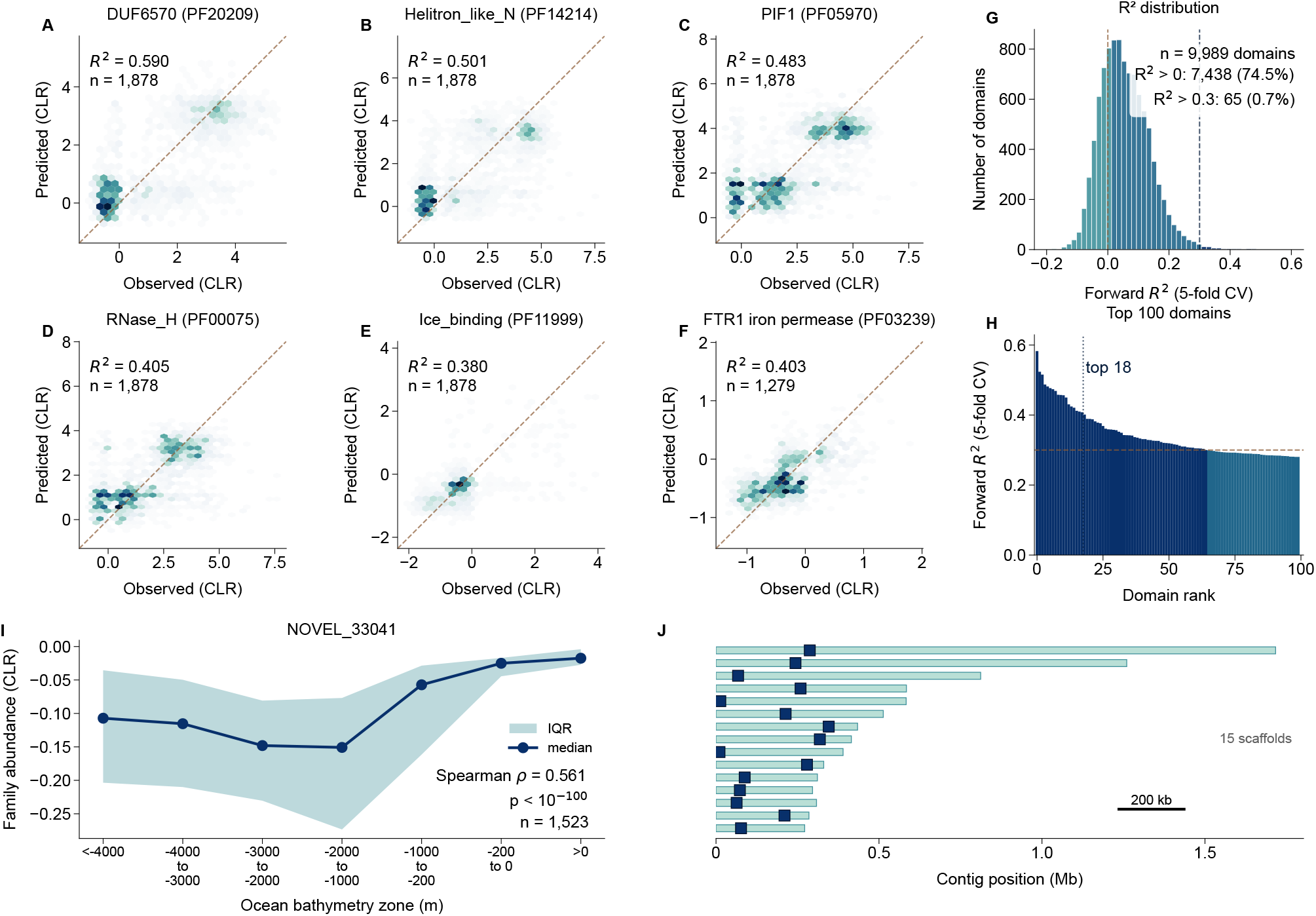
Forward model calibration, prediction accuracy, and biological proof-of-concept. **(A– E)** Observed versus predicted centered log-ratio (CLR)-transformed Pfam abundance (10-fold spatial block cross-validation [CV], *n* = 1,878 Global Positioning System [GPS]-recovered samples; satellite-only environmental features) for the five most predictable non-phage domains: DUF6570 (*R*^2^ = 0.59), Helitron_like_N (0.501), PIF1 (0.483), RNase_H (0.405), Ice_binding (0.380). Hexbin density; dashed line = identity. This later GPS-recovered set includes 68 samples that lacked a named basin assignment in the frozen 1,810-sample basin table; Table S2 reports the World Ocean Atlas 2023 (WOA23)-merged, basin-resolved analysis and therefore differs slightly in sample composition and *R*^2^. **(F)** FTR1 iron permease (PF03239): observed versus predicted CLR-transformed abundance (5-fold CV, *n* = 1,279; *R*^2^ = 0.40). FTR1 encodes a high-affinity iron uptake permease induced under iron limitation. ^32^ **(G)** Forward *R*^2^ distribution across 9,989 domains in the five-fold CV screen (*n* = 1,279): 65 (0.7%; fold-bootstrap 95% confidence interval [CI]: 51–118) exceed *R*^2^ *>* 0.3; 74.5% exceed *R*^2^ *>* 0 (versus 5% expected under permutation null). **(H)** Top 100 domains from the same screen; dotted line, top-18 cutoff; dashed line, *R*^2^ = 0.3. **(I)** NOVEL_33041 Pfam-dark protein-family abundance (CLR) across ocean bathymetry zones (median with interquartile range [IQR]; *n* = 1,523). The archived panel-reproduction vector gives Spearman *ρ* = 0.561 (*p <* 10^−100^); the frozen primary correlation table gives *ρ* = 0.566 for the same sample count (Data S7). NOVEL_33041 has no Pfam or integrated InterPro functional annotation yet is the tightest single family–environment coupling in the dataset; a PDB homology result is not available for this family. **(J)** Genomic context of NOVEL_33041 in the *Dunaliella* ROIL 10x Genomics assembly. Fifteen of the 20 loci on chromosome-scale scaffolds (*≥*100 predicted genes each) are shown; the family occurs at 66 loci on 66 distinct contigs overall, supporting dispersed multi-copy expansion. Within-genome pairwise *d*_N_*/d*_S_ = 0.33 (97.4% of 2,145 pairs *<* 1; Table S13), supporting purifying selection.

In the genome-to-environment direction, CLR-transformed Pfam abundances predicted SST at *R*^2^ = 0.38 and bathymetry at *R*^2^ = 0.42 under spatial block CV, with 10 of 37 targets exceeding *R*^2^ *>* 0.2 (Data S1). Fixed 80/20 split estimates are shown separately in Figure 3E. Restricting to TARA metagenomes alone *increased* SST *R*^2^ to 0.44 (metagenome-only CCA CC1 = 0.89 vs. 0.82 full dataset; Supplemental Text). Dissolved nutrients were also predictable (nitrate *R*^2^ = 0.42, dissolved oxygen 0.40, phosphate 0.37). Leave-one-basin-out (LOBO) CV yielded pooled held-out *R*^2^ = 0.31 for bathymetry, with positive basin-level performance in 3 of 7 basins that together contained 84.9% of evaluated samples. SST retained pooled global-scale signal (*R*^2^ = 0.24), but basin-level performance was negative in 6 of 7 basins (Table S6), indicating that SST– domain mappings do not transfer consistently within individual basins.

Temporal holdouts further reduced SST performance to *R*^2^ = 0.16–0.25, indicating that cross-sectional estimates partly capture stable site effects (Figure S5A–D; Supplemental Text).

In the environment-to-genome direction, a two-tier design (5-fold CV screen of 9,989 domains, then 10-fold spatial block CV of the 18 top-ranked; Table S2; Methods) revealed that mobile genetic element domains constituted 11 of 18 selected domains (*R*^2^ = 0.27–0.59; median 0.40). Because the targets were selected on the overlapping 1,279-sample discovery cohort before spatial validation, these spatial-CV scores condition on screen selection and are not selection-independent estimates; the maximum may therefore be optimistic. DUF6570, a domain of unknown function in dinoflagellate transposons, achieved the largest observed spatial-CV value (*R*^2^ = 0.59 from satellite imagery alone; Figure 5A; Table S2).

Within the Table S2 spatial-validation panel, the most predictable non-TE domains include PIF1 helicase (*R*^2^ = 0.46), Ice_binding (PF11999; *R*^2^ = 0.42; the antifreeze protein polar diatoms secrete to survive subzero waters), and AAA ATPases (0.38–0.40). Additional non-TE domains from the separate five-fold discovery screen included iron permease FTR1 (*R*^2^ = 0.40; Figure 5F) and dUTPase (*R*^2^ = 0.40; Table S9); neither belongs to the Table S2 spatial panel.

These non-TE domains recapitulate known ecophysiology without supervision: FTR1 iron permeases are induced under iron limitation,^32^ the primary constraint on Southern Ocean productivity, and eukaryotic carbonic anhydrase (PF00194) co-segregates with the SST-range cluster C9 (Figure 4). Recovery of these relationships from domain abundances supports a biological basis for the coupling; domain-by-domain interpretation is in Table S9 and the Supplemental Text. Calibration plots show that predictions track observed abundances (Figure 5A–F).

Predictability was concentrated in an elite subset: 65 domains (0.7% of 9,989 tested; fold-bootstrap 95% confidence interval [CI]: 51–118) exceeded *R*^2^ *>* 0.3, enriched for mobile elements and DNA processing machinery (Figure 5G–H; Data S1). More broadly, 27.3% of domains exceeded *R*^2^ *>* 0.1 and 4.4% exceeded *R*^2^ *>* 0.2 (5-fold CV; versus *∼*5% expected at *R*^2^ *>* 0 under permutation null), indicating that meaningful environmental signal extends well beyond the elite subset, albeit at modest effect sizes for most domains.

SHAP decomposition of the XGBoost models and biclustering across 64 AEF embedding dimensions revealed 16 functionally coherent domain groups (Figure 4). Two domains lead the SHAP rankings of the reverse models: PF11999 (Ice_binding) carries the highest SHAP importance for SST, while Transposase IS66 (PF03050), a member of the DUF6570-anchored TE ensemble, ranks first for bathymetry. Together with PF00692 (dUTPase), they define cold adaptation, mobile element activity, and DNA metabolism as the three most environment-responsive functional axes.

K-means clustering of the Spearman correlation matrix (384 Pfams *×* 29 environmental variables; *k* = 8, silhouette = 0.286) revealed coherent modules of domains sharing similar environmental association profiles (Figure 3A).

Unsupervised Hierarchical Density-Based Spatial Clustering of Applications with Noise (HDB-SCAN)^33^ of domain abundances alone, without geographic input, recovered functional biomes moderately concordant with Longhurst biogeochemical provinces (adjusted Rand index [ARI] = 0.503; Figure S4B). Gene Ontology (GO) enrichment supported functional coherence among environment-predictable domains (*q <* 0.05; exploratory; Figure S4A; Table S5; Supplemental Text). Excluding 23 transposable-element (TE) domains left reverse-model performance nearly unchanged (bathymetry *R*^2^: 0.491*→*0.484; SST: 0.608*→*0.611), showing that non-TE features retained predictive signal; this ablation does not estimate the importance of the excluded TE features (Figure 3D; Supplemental Text).

The 16 SHAP-biclustered domain groups span temperature, bathymetry, chlorophyll, and ocean color axes (Figure 4; Supplemental Text). Partial Spearman correlations controlling for RuBisCO lineage identity confirmed that 31 of 32 tested associations retained significance (*n* = 1,005; Supplemental Text), and all 44 TE–environment associations retained significance after controlling for the five myzozoan Form II lineages (four dinoflagellate lineages plus Chromerida; Bonferroni-corrected; Supplemental Text).

Distance-based redundancy analysis (db-RDA; *n* = 786) using Aitchison distance partitioned domain composition into pure environmental (adj. *R*^2^ = 0.029), pure geographic (0.034), and shared (0.017) fractions (conditional permutation *p* = 0.001 for both pure fractions; total adj. *R*^2^ = 0.080, comparable to published marine plankton values^34^; Supplemental Text). The combined environmental signal (pure plus shared) accounts for 57.6% of explained variance versus 42.4% for pure geography.

### The proteins most coupled to ocean state are sparsely functionally annotated

Pfam-A v37.2 annotation at the strict threshold of *E <* 10^−9^ leaves 86.9% of predicted proteins without any domain assignment, the “Pfam-dark proteome,” disproportionately concentrated in algal sequences (Data S8). Yet this dark fraction contains recurrent protein clusters with 2.29-fold stronger environmental coupling than Pfam domains (matched design; bootstrap 95% CI [2.28, 2.30]; Data S7).

Physicochemical profiling (500,000 open reading frames [ORFs] per class; Figure 6E–K) reveals that dark proteins are shorter (median 115 vs. 205 amino acids [aa]), more basic (isoelectric point [pI] 9.22 vs. 6.41), more disordered (0.556 vs. 0.500), of lower sequence complexity (Shannon entropy 3.87 vs. 4.05 bits), and enriched in amino acids with GC-rich codons (GC-proxy median 0.213 vs. 0.064)—hallmarks of rapidly evolving lineage-specific proteins whose divergence erodes BLAST and profile-HMM sensitivity.^35^ This compositional divergence extends to dipeptide usage (1.54-fold wider observed/expected [O/E] distribution; Figure S6C–J; Table S10; Supplemental Text).

**Figure 6:**
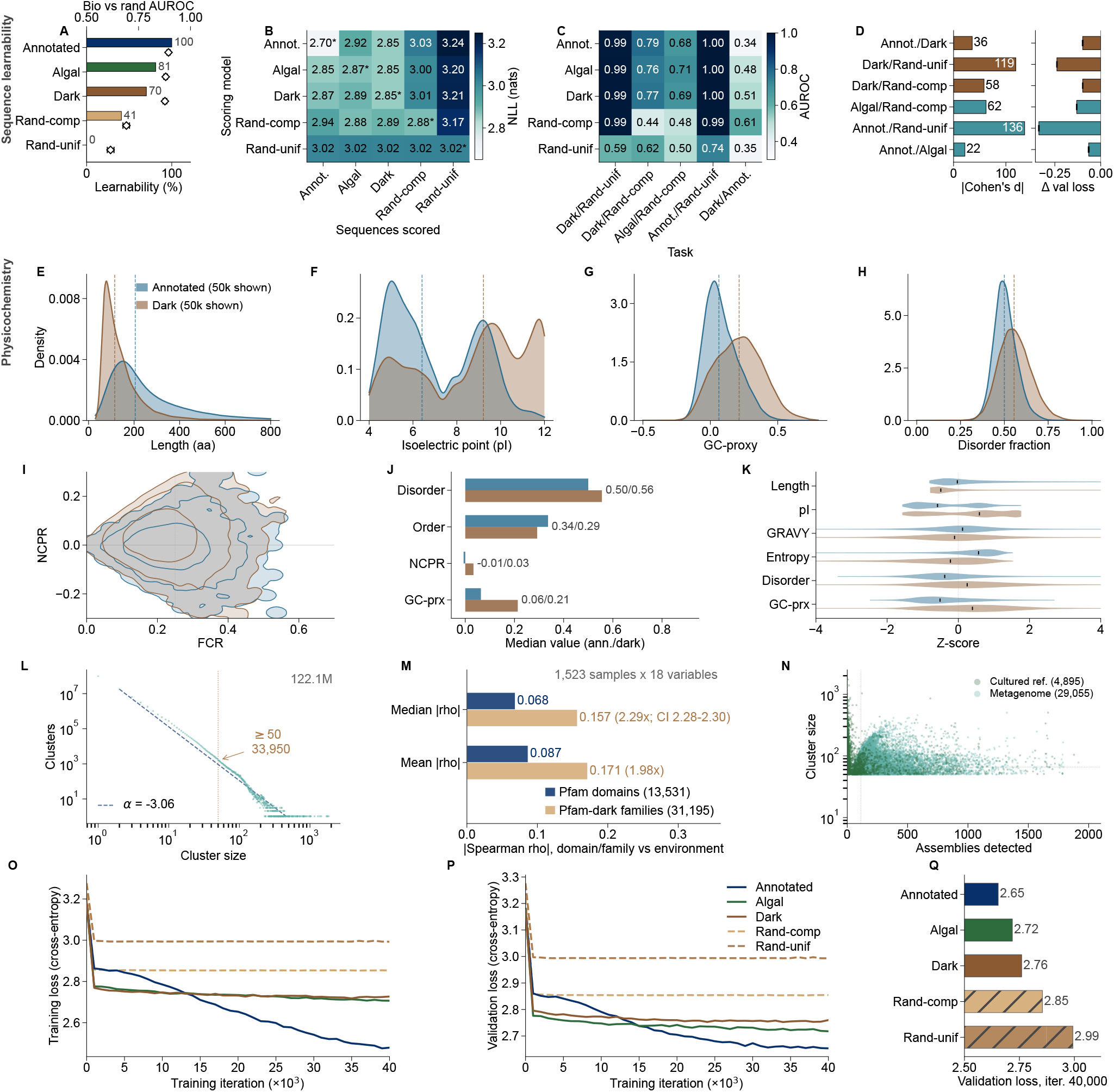
Dark proteins carry biological sequence structure, are physicochemically distinct, and couple tightly to ocean state. The figure is organized into four blocks: sequence learnability (A– D), physicochemistry (E–K), discovery and environmental coupling (L–N), and training dynamics (O–Q). Structure predictions for well-folded Pfam-dark family representatives are shown in Figure 7. *Sequence learnability (A–D).* Five generative pre-trained transformer (GPT)-2 Small models (85 million parameters, character-level; nanoGPT) were trained from scratch under identical hyperparameters on five corpora: annotated proteins (Annotated), algal proteome (Algal), dark proteins (Dark), composition-matched random sequences (Rand-comp), and uniform random sequences (Rand-unif). Per-sequence discrimination used 1,000 held-out sequences per corpus. **(A)** Relative learnability *L*(*D*) = 100 (*L*_floor_ *L_D_*)*/*(*L*_floor_ *L*_best_) with aggregate biological-vs-random area under the receiver operating characteristic curve (AUROC) per scoring model (diamonds, 95% bootstrap confidence interval [CI]). **(B)** Cross-dataset per-sequence mean negative log-likelihood (NLL; 1,000 held-out sequences per corpus; rows, scoring model; columns, corpus scored; asterisks, model’s own corpus). **(C)** Cross-model AUROC heatmap for five binary classification tasks. **(D)** Cohen’s *d* effect sizes computed over the last ten validation-loss evaluations of each run (left) and final validation-loss differences with 95% bootstrap CI (right; 9,999 resamples). Reported *p*-values are from one-tailed Mann–Whitney *U* tests on the final ten evaluation steps (all *p <* 0.001). *Physico-chemistry (E–K).* Annotated (blue) versus dark (brown) proteins from fixed 500,000-sequence-per-class source samples drawn from the 2,044-sample mixed-source analysis set (seed = 42). Panels E–H display 50,000 sequences per class and dashed lines mark medians. **(E)** Sequence length. **(F)** Isoelectric point. **(G)** Guanine–cytosine (GC)-content proxy. **(H)** Disorder fraction. **(I)** Charge landscape for 15,000 sequences per class: fraction of charged residues (FCR) versus net charge per residue (NCPR). **(J)** Medians computed from all 500,000 sequences per class across four metrics. **(K)** Z-score summary violins for 10,000 sequences per class across six metrics. *Discovery and environmental coupling (L–N).* **(L)** Cluster size distribution (122.1 million clusters; 33,950 hidden Markov models [HMMs] at ≥50-member threshold). **(M)** Environmental coupling in the matched design (same 1,523 samples and 18 variables): median and mean *ρ* for 13,531 Pfam domains and 31,195 Pfam-dark protein families (median ratio 2.29; bootstrap 95% CI [2.28, 2.30]). **(N)** Cluster size versus prevalence for 33,950 Pfam-dark protein families, colored by source type. *Training dynamics (O–Q).* GPT-2 training curves for five protein corpora: Annotated, Algal, Dark, Rand-comp, and Rand-unif. Dark proteins converge to lower loss than random-sequence controls, supporting learnable biological structure beyond amino-acid composition alone. **(O)** Training loss (cross-entropy) versus iteration. **(P)** Validation loss versus iteration. **(Q)** Validation loss at iteration 40,000 (horizontal bars); biological corpora achieve lower loss than random controls.

Two independent lines of evidence support biological sequence constraint rather than simple accumulation of spurious ORFs.

#### Cross-taxonomic selection analysis

Population-genomic data from four lineages (three eukaryotic algae and the cyanobacterium *Synechococcus*) show altered selective constraint in Pfam-dark genes, using metrics appropriate to each dataset (Table S11; Table S12; Data S9). Dark genes had higher *π*_N_*/π*_S_ in *Chlamydomonas reinhardtii* ^36^ (2.30-fold; median 0.277 vs. 0.121; *p <* 10^−300^; *n* = 11,123) and *Seminavis robusta* ^37^ (1.46-fold), higher *d*_N_*/d*_S_ in *Synechococcus* ^38^ (1.84-fold), and a greater incidence of site-model evidence for positive selection in *Thalassiosira pseudonana* ^39^ (9.3% vs. 4.9%; rate ratio 1.90; Supplemental Text). These metrics operate at different evolutionary scales and were not pooled. Together they support altered constraint as one mechanism by which sequence similarity can fall below homology-detection thresholds. This evidence is assembly-independent, deriving from population-genomic data in cultured organisms rather than from the metagenomic samples.

#### Protein language model learnability

We defined learnability as the reduction in negative log-likelihood (NLL; mean next-residue prediction loss, where lower values indicate more predictable sequence) relative to random-sequence baselines. Five generative pre-trained transformer (GPT)-2 Small models^40^ (85M parameters; nanoGPT^41^) trained on annotated, algal, dark, composition-matched random, and uniform-random corpora placed dark proteins at 70.2% of the range between the random floor and annotated proteins (Methods; Data S11; Figure 6A). The area under the receiver operating characteristic curve (AUROC; 0.5 = chance, 1 = perfect separation) was 0.990 (95% CI [0.987, 0.993]) for dark versus uniform-random sequences and 0.766 ([0.745, 0.787]) for dark versus composition-matched random sequences. Dark-versus-annotated discrimination was at chance (AUROC = 0.506; Figure 6B–D). All biological models assigned similar NLL to the biological corpora (2.70–2.92 nats) and higher NLL to random sequences (3.00–3.24 nats), supporting comparable learnable sequence structure in dark and annotated proteins.

Together, altered selective constraint (selection analysis) and full biological sequence structure (language model) frame the dark proteome as a distinct evolutionary compartment rather than an annotation gap. To determine whether these proteins respond to the environment, MMseqs2 linclust^42^ clustering of 201 million Pfam-dark proteins at 30% identity and 80% bidirectional coverage yielded 33,950 whole-protein families (*≥*50 members each) from which family-profile HMMs were built and searched against all samples (Methods; Data S7). These are operational full-protein units, not independently delimited or validated protein domains.

Environmental correlation analysis under a matched design (same 1,523 samples, same 18 GEE+WOA23 variables) yielded median *|ρ|* = 0.157 for Pfam-dark protein families versus 0.068 for Pfam domains, a 2.29-fold enrichment (bootstrap 95% CI [2.28, 2.30]; Mann-Whitney *p <* 10^−300^; Figure 6M; Data S7). This enrichment is robust across E-value thresholds (2.32-fold at *E <* 10^−5^), persists after prevalence matching (3.0-fold; Supplemental Text), and holds when restricted to high-prevalence families largely free of zero-inflation (1.81-fold at *>*75% prevalence; *p <* 10^−53^).

The strongest individual association belongs to the Pfam-dark protein family NOVEL_33041, conserved across five algal phyla (Chlorophyta, Stramenopiles, Haptophyta, Euglenozoa, Charophyta), which tracks ocean bathymetry at *ρ* = 0.566 (*n* = 1,523; Figure 5I), the tightest single family–environment coupling in the dataset. NOVEL_33041 had no Pfam or integrated InterPro functional annotation. DIAMOND returned five weak or partial matches—four hypothetical proteins and one alignment to the non-reverse-transcriptase region of a reverse-transcriptase-labelled multidomain protein—but none assigned a credible function; a PDB homology result is not available for this family (Supplemental Text). Across all 30,994 prevalent Pfam-dark protein families, 99.3% had at least one significant nutrient correlation, indicating broad coupling to the dissolved nutrient environment.

Geographic breadth analysis supports a lineage-restriction mechanism: Pfam-dark protein families with narrower geographic ranges couple more tightly to local conditions (*ρ* = *−*0.349, *n* = 7,935), the opposite of the Pfam-domain pattern (*ρ* = +0.145; Supplemental Text). Controlling for 15 RuBisCO lineage counts attenuated the enrichment from 2.08-fold to 1.83-fold in the 1,133 samples with Pfam, Pfam-dark-family, environmental, and RuBisCO data (a 22.5% reduction of the enrichment above 1; *p <* 10^−300^), confirming that Pfam-dark families retain stronger environmental coupling than Pfam domains after lineage correction (Data S4; Data S7). InterProScan cross-checking of the 33,950-family HMM catalogue found that 86 of its 100 top-ranked families lacked an integrated InterPro entry. A separate photosynthetic-anchor-neighbor screen began with 58,570 unannotated proteins and produced 35,667 MMseqs2 cluster representatives; across this independent set, 96.8% lacked a functional annotation from InterPro member databases, and AntiFam flagged only 10 sequences (0.03%) as potential spurious ORFs (Table S4; Data S7). The 35,667 representatives are therefore not an unfiltered stage of the 33,950-family HMM catalogue.

#### Structure prediction

Boltz-2^43^ and AlphaFold 3^44^ predictions across 3,600 rank-binned Pfam-dark family representatives showed that the dark proteome is predominantly disordered (65.0% with disorder fraction *≥* 0.9). AlphaFold 3 nevertheless identified a well-folded tail of 138 representatives (pTM *≥* 0.5; maximum 0.89), none with an InterPro annotation; 12 high-confidence examples (pTM = 0.78–0.89) are shown in Figure 7. These families span all environmental-coupling deciles (Figure S7; Data S7; Data S10). ESMFold^8^/Foldseek^45^ analysis of the ten highest-ranked families found five with no structural homolog (*E <* 10^−3^; Data S7; Supplemental Text). Thus, even within a predominantly disordered dark proteome, a database-unannotated subset is predicted to adopt stable folds.

**Figure 7:**
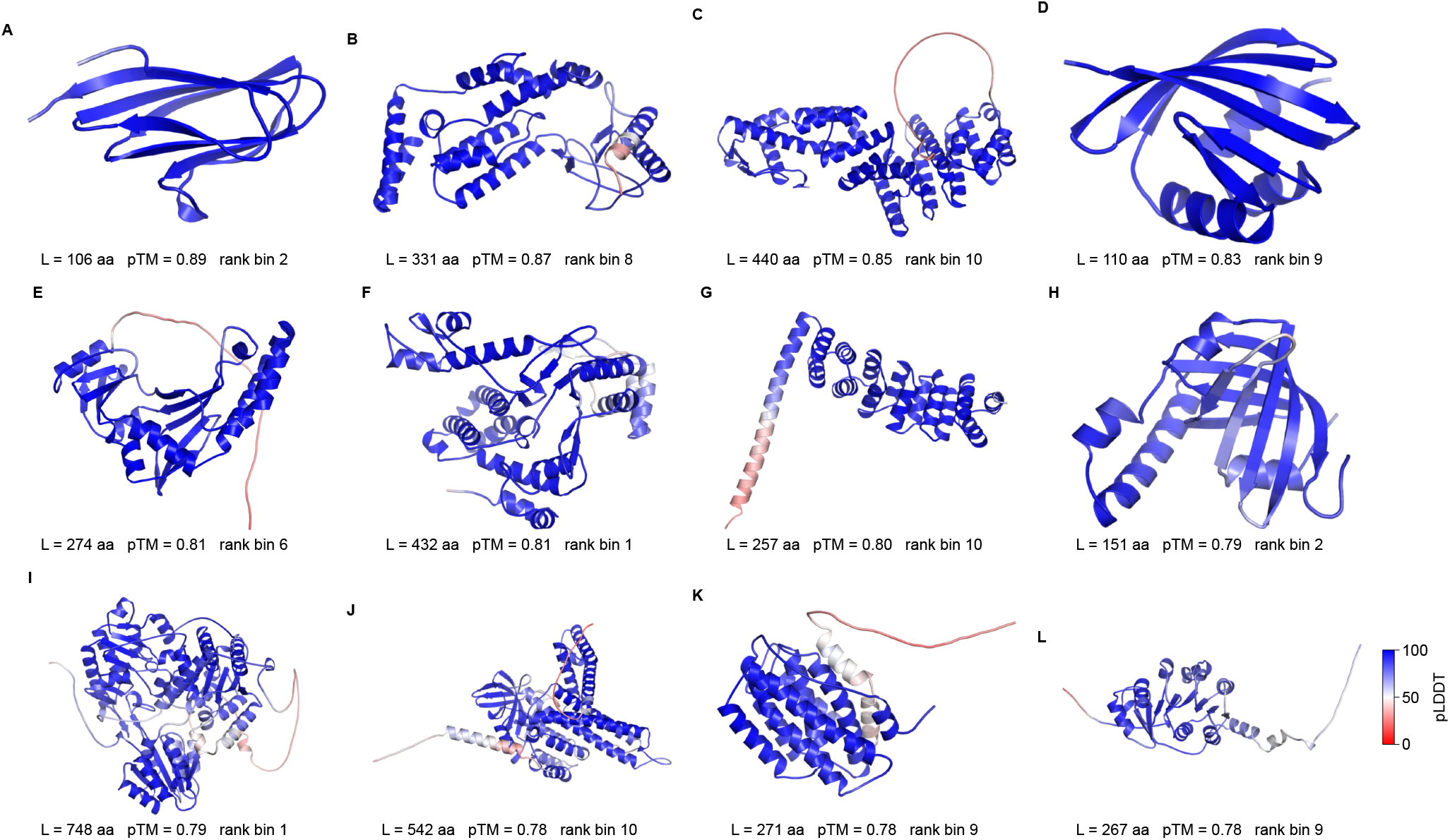
AlphaFold 3 structure predictions for twelve well-folded Pfam-dark protein-family representatives. All twelve representatives (predicted template modeling score [pTM] = 0.78–0.89) lack an InterProScan annotation; none of the 138 well-folded representatives carried a functional annotation. Structures are colored by predicted local distance difference test (pLDDT; continuous red–white–blue scale from 0 [red] through 50 [white] to 100 [blue]). Metadata below each render give sequence length, pTM, and rank bin (decile of maximum *ρ* with any environmental variable; bin 1 = strongest, bin 10 = weakest). The twelve examples fall in rank bins 1, 2, 6, 8, 9, and 10. These predictions provide high-confidence structural models despite absent InterPro annotation; structural homology was assessed separately for the ten highest-ranked families (Results; Data S7).

### Genomic composition adds modest information to satellite-based productivity estimates

To test whether the bidirectional framework extends to direct productivity prediction, we trained XGBoost models on CLR-transformed domain profiles (principal component analysis [PCA]-reduced to 100 components) to predict three satellite-derived productivity proxies collocated at each sampling coordinate: chlorophyll-*a* concentration, particulate organic carbon (POC), and normalized fluorescence line height (NFLH). Under 10-fold spatial block CV (*n* = 1,152 for chl-a and POC; *n* = 1,151 for NFLH; 277 spatial blocks at 2-degree resolution), domain composition alone yielded median *R*^2^ = 0.07 (chl-a), *−*0.12 (POC), and 0.12 (NFLH), indicating that Pfam profiles carry minimal direct productivity signal when evaluated across spatially independent folds. Environment-only models (26 non-productivity GEE variables) performed well: median *R*^2^ = 0.80 (chl-a), 0.59 (POC), 0.83 (NFLH). Combined models (domains + environment, 126 features) improved over environment-only for chl-a (median *R*^2^ = 0.83, a +0.04 increment) and POC (*R*^2^ = 0.66, +0.07) but not NFLH (*R*^2^ = 0.80, *−*0.03; Figure S5; Data S1). SHAP analysis of the combined chl-a model identified remote-sensing reflectance (RRS) bands at 443, 412, and 645 nm as dominant predictors (mean *|*SHAP*|* = 0.87, 0.36, 0.23), with four Pfam domains contributing modestly at ranks 11–15 (PF07727, PF09815, PF00589, PF07707; mean *|*SHAP*|* = 0.020–0.034). Thus, adding domain features modestly improved two of three targets; all productivity-derived variables were excluded from the predictors (Methods).

A self-supervised joint embedding using Variance-Invariance-Covariance Regularization (VICReg)^46^ captures biogeographic structure (Longhurst ARI = 0.52; Supplemental Text) but fails to generalize under spatial block CV for productivity prediction (negative *R*^2^ on held-out basins; Supplemental Text), identifying an architectural bottleneck in the multilayer perceptron (MLP) prediction head rather than a data limitation. Full results and model checkpoints are released as an open problem.^47^

Restricting to the algal-dominated 20–180 µm TARA fraction alone (*n* = 150) confirmed retained coupling: XGBoost SST *R*^2^ = 0.29 under spatial block CV and CCA CC1 = 0.917 (permutation *p* = 0.001, *Z* = 3.62; Table S7; Supplemental Text), with the reduced *R*^2^ (vs. 0.39 for the full dataset) consistent with a 7.8-fold smaller sample size. These results demonstrate that environment–genome coupling is detectable from existing metagenomic and remote-sensing data, providing a quantitative basis for incorporating functional genomic information into marine productivity models.

## DISCUSSION

Satellite-based productivity models estimate marine productivity without directly representing which organisms or molecular machinery are present. Our results show that metagenomes contain information relevant to this gap: protein domain composition and ocean state predict each other from co-located observations across *∼*44,000–86,000 algal taxa and seven ocean basins. The coupling is asymmetric: the environment-to-genome direction concentrates in a small, functionally enriched tail, whereas the genome-to-environment direction (*∼*10,000 domains predicting one target) is dimensionally easier.

### Predictability differs among environmental variables

Microalgal domain composition organizes along coherent environmental gradients, consistent with temperature as a primary filter on functional diversity.^9^ Predictability was variable-specific: nitrate (*R*^2^ = 0.42) and dissolved oxygen (0.40) matched or exceeded bathymetry (0.42) and SST (0.38), whereas silicate was lower (0.23). These cross-sectional associations do not establish directionality. Leave-one-basin-out CV (Table S6) further shows that bathymetry transfers across most represented open-ocean samples, whereas SST does not transfer consistently within individual basins.

The framework cannot fully distinguish environmental filtering from dispersal limitation, taxonomic turnover, or unmeasured top-down controls. Pure geographic effects slightly exceed pure environmental effects, although the combined environmental fraction accounts for 57.6% of explained variance. Lineage-controlled partial correlations retain domain–environment coupling, but generalizability beyond TARA Oceans and OSD remains untested; CCA components CC6– CC10 are exploratory.

### Environment-predictive domains recapitulate known ecophysiology without supervision

The top environment-predictive domains show functional coherence consistent with known marine biology. Ice_binding PF11999, an antifreeze protein in polar diatoms, is the top SST predictor in both directions of coupling. PF00692 encodes dUTPase, which sanitizes nucleotide pools,^48^ and its abundance associates here with bathymetry; a role in depth-related genome maintenance remains a hypothesis. PEPCK PF17297 contributes to light-independent carbon fixation in diatoms^49^ and associates here near-exclusively with normalized fluorescence line height. Extended per-domain interpretations are provided in Table S9 and the Supplemental Text.

At a broader scale, temperature-associated clusters split photosynthetic and genome-maintenance machinery between two thermal axes (Figure 4): chlorophyll *a*/*b*-binding and family-B DNA polymerase domains cluster with low minimum SST (C3), whereas photosynthetic reaction center (C6), carbonic anhydrase, and AAA ATPase (C9) domains cluster with SST range. The co-occurrence of AAA ATPase chaperones with carbonic anhydrase in C9 links protein quality control to thermal variability rather than to mean temperature. That unsupervised analysis recovers this partitioning from domain abundances alone confirms that functional composition encodes the dominant axis of ecological variation (CC1 captures 17% of total shared variance).

#### Transposable elements as environmental sensors

The prominence of TE domains among environment-predictive features (11 of 18 top forward-model domains, 12 of 13 top sparse CCA loadings on CC1, with DUF6570 alone achieving *R*^2^ = 0.59) requires explanation. Three non-exclusive hypotheses can explain it: (i) taxonomic covariation, in which TE-rich dinoflagellate genomes^31^ shift community fraction along environmental gradients, dragging TE counts with them; (ii) stress-induced TE activation, in which environmental conditions directly modulate transposon expression and mobilization; and (iii) TE-mediated adaptive divergence, in which transposon insertions rewire regulatory networks under selection.

Hypothesis (i) alone is insufficient at the tested taxonomic resolution: partial correlations controlling for the five myzozoan Form II lineages (four dinoflagellate lineages plus Chromerida) retained 100% of TE–environment associations (Supplemental Text). Prior experiments in diatoms^50–52^ and dinoflagellates^53^ support stress-induced TE activation as a plausible mechanism, and an independent analysis of the same TARA metagenomes found TE genes reaching 13.5% of total gene abundance.^54^ Our abundance data do not measure transposon expression or mobilization and therefore cannot identify a primary mechanism. Longitudinal metatranscriptomic data would be needed to separate stress-induced activation from finer taxonomic sorting and TE-mediated regulatory divergence.

Domain-based predictions (*R*^2^ = 0.38–0.42) match or exceed reported trait-based models (*R*^2^ = 0.1–0.3^55^). In a proof of concept, adding domain composition modestly improved two of three remote-sensing productivity targets (Figure S5). Pfam-dark protein families couple more than twice as strongly as Pfam domains (Results).

### The most environment-coupled proteins are sparsely functionally annotated

The dark proteome analysis (Results) reveals that the Pfam-unannotated fraction contains recurrent protein families with stronger environmental coupling than curated domains and sparse informative functional annotation. Some top families nevertheless returned integrated InterPro or structural homolog matches, so database absence is not universal. This pattern supports enrichment for recently diverged, lineage-specific proteins that Pfam’s conservation-based curation systematically underrepresents.

Selection, language-model, and structure analyses support the dark proteome as a genuine evolutionary compartment. Within-study nonsynonymous/synonymous contrasts were elevated in three lineages, and positive-selection incidence was elevated in a fourth; these distinct metrics were interpreted separately. GPT-2 models place dark proteins 70.2% of the way from random controls to annotated proteins in learnability, with dark-versus-annotated discrimination at chance (AUROC = 0.506). AlphaFold 3 identifies 138 well-folded but InterPro-unannotated family representatives within the predominantly disordered fraction. Thus, database absence does not imply random sequence or universal disorder.

This inverse relationship between coupling strength and database representation is most acute for NOVEL_33041. The tightest family–environment coupling in 231 million proteins (*ρ* = 0.566, bathymetry) belongs to this family, conserved across five algal phyla yet lacking a Pfam or integrated InterPro functional annotation. DIAMOND returned five weak or partial matches, none with a credible functional assignment; a PDB homology result is not available for NOVEL_33041. In the *Dunaliella* ROIL 10x Genomics assembly, where it was first characterized, it is dispersed across 66 loci on 66 distinct contigs, including 20 chromosome-scale scaffolds (*≥*100 predicted genes each; 15 shown in Figure 5J), a distribution consistent with a functional gene family rather than a transposon artifact.

Four independent observations support biological function: (i) within-genome *d*_N_*/d*_S_ = 0.33 across 66 copies (97.4% of pairs *<* 1; Table S13), consistent with purifying selection; (ii) transcription in four MMETSP transcriptomes and detection in 118 metagenome samples; (iii) conservation across five algal phyla spanning *∼*1.5 Gyr (Chlorophyta, Stramenopiles, Haptophyta, Euglenozoa, Charophyta); and (iv) only five weak or partial DIAMOND BLASTp hits against NCBI nr, four labelled hypothetical and one aligning outside the annotated reverse-transcriptase region of a multidomain protein (Supplemental Text).

The 198-aa NOVEL_33041 representative contains cysteine-rich motifs (CVC, CPC; 4.5% of residues, 3.3*×* the Swiss-Prot average), tryptophan enrichment (7.1%, 6.5*×*), and an estimated net charge of +9.7 at pH 7 (BioPython `ProteinAnalysis`); its genomic co-localization with an antifreeze type I protein (IPR000104) is consistent with a possible extracellular or surface-associated role. Heuristic analysis identified a possible but atypical signal peptide and no transmembrane domain; localization remains untested (Supplemental Text). That a protein shared across five algal phyla lacks informative functional annotation illustrates how database architecture constrains what we can interpret.

We hypothesize that the dark proteome is enriched in environmental interface proteins: the top-ranked non-TE annotated domains discussed above (ice-binding, iron permease) mediate a direct interaction between organism and environment, the functional class expected to diverge fastest under habitat-specific selection. NOVEL_33041’s compositional signature and purifying selection (*ω* = 0.33) are consistent with this prediction; whether it extends to the full unannotated fraction requires experimental testing.

The enrichment is robust across E-value thresholds, and catalogue absence was cross-checked with InterProScan (Results; Table S4). Because protein-family databases favor conservation across broad phylogenetic ranges, they underrepresent lineage-specific, rapidly diverging proteins. Data-driven family-profile HMMs can recover part of this missing signal; the 33,950 Pfam-dark protein-family HMMs released in Data S7 provide a starting point for experimental characterization.

### Limitations of the study

This study has several limitations. First, the cross-sectional design cannot distinguish environmental filtering from historical biogeography or unmeasured top-down controls; pure geographic effects (adj. *R*^2^ = 0.034) slightly exceed pure environmental effects (adj. *R*^2^ = 0.029). Second, AEF records cover only 55.0% of GPS-mapped samples and are biased toward coastal and shelf sites. Third, leave-one-basin-out SST performance is negative in 6 of 7 basins; the positive pooled score (*R*^2^ = 0.24) reflects the global SST range rather than consistent within-basin transfer. Fourth, temporal holdout attenuated SST *R*^2^ to 0.16–0.25, indicating that cross-sectional estimates partly capture stable site effects. Fifth, experimental characterization is required to distinguish new functions from lineage-restricted markers. Controlling for 15 RuBisCO lineages attenuated Pfam-dark-family enrichment from 2.08-fold to 1.83-fold, showing that lineage composition explains part of the signal; independent selection and language-model analyses support biological sequence constraint without resolving function.

## DATA AND CODE AVAILABILITY

### Data

All processed data supporting this study are publicly available as Data S1–S11. Stable repository identifiers and the contents of each deposit are listed in the Supplemental Information index and the Resources table (Methods). Source metagenome assemblies are available from EBI MGnify^56,57^ (MGYA accessions), and MMETSP transcriptomes^18,58^ are available from NCBI (BioProject PRJNA231566).

### Code

All original analysis code (371 scripts organized into 15 modules; Code S1) has been deposited at Zenodo^59^ and is publicly available (DOI: https://doi.org/10.5281/zenodo.22910378). The trained algaGPT classifier,^60^ bidirectional XGBoost models,^61^ VICReg joint-embedding checkpoints,^47^ and dark-whiteGPLM protein-language-model checkpoints^62^ are available through Hugging Face. The archived protein-language-model analysis code and results are provided as Data S11.^63^

### Additional information

Any additional information required to reanalyze the data reported in this paper is available from the corresponding authors upon request.

## ACKNOWLEDGMENTS

We thank the TARA Oceans Consortium for making metagenomic data publicly available, the Ocean Sampling Day (OSD) initiative for coordinated coastal metagenome sampling, the Marine Microbial Eukaryote Transcriptome Sequencing Project (MMETSP) for transcriptome resources, and NCBI GenBank/RefSeq and the authors of the reference genome assemblies listed in Table S14 for cultured reference genome data. Computational resources were provided by New York University Abu Dhabi High Performance Computing facilities. The Cawthron Institute is also kindly acknowledged for supporting this research. This work was supported by NYUAD Faculty Research Funds (grant AD060).

## AUTHOR CONTRIBUTIONS

Conceptualization, D.R.N. and K.S.-A.; Methodology, D.R.N. and M.P.; Software, D.R.N.; Investigation, D.R.N., M.P., S.D., A.J., and W.F.; Formal Analysis, D.R.N. and S.D.; Writing–Original Draft, D.R.N.; Writing–Review & Editing, all authors; Supervision, S.A.A. and K.S.-A.; Funding Acquisition, K.S.-A.

## USE OF AI-ASSISTED TECHNOLOGIES

During the preparation of this work, the authors used Claude (Anthropic) and Codex (OpenAI) to assist with code development, data analysis, and manuscript preparation. The authors reviewed and edited all AI-assisted outputs and take full responsibility for the content of the publication.

## METHODS

### Resources

| RESOURCE | SOURCE | IDENTIFIER |
| --- | --- | --- |
| **Deposited data** |  |  |
| TARA Oceans metagenome assemblies (1,063) | EBI MGnify<sup>56,57</sup> | MGYA accessions; https://www.ebi.ac.uk/metagenomics/ |
| Ocean Sampling Day metagenome assemblies (140) | EBI MGnify<sup>17</sup> | MGnify study MGYS00001977 |
| MMETSP transcriptomes (676) | iMicrobe/NCBI<sup>18,58</sup> | PRJNA231566 |
| Cultured microalgal reference genome assemblies (478: 284 NCBI GenBank/RefSeq, 194 other published assemblies) | NCBI GenBank/RefSeq;<sup>65</sup> Nelson et al.<sup>66,67</sup> and original genome publications (Table S14) | GCA/GCF accessions and the publishing study per assembly in Table S14 |
| Pfam-A database (v37.2, November 2024; 24,076 families) | EMBL-EBI<sup>12,68</sup> | https://ftp.ebi.ac.uk/pub/databases/Pfam/releases/Pfam37.2/ |

| **RESOURCE** (*continued*) | **SOURCE** | **IDENTIFIER** |
| --- | --- | --- |
| AlphaEarth satellite embeddings | Google Earth Engine<sup>2,13,69</sup> | GOOGLE/<br>SATELLITE_EMBEDDING/<br>V1/ANNUAL |
| World Ocean Atlas 2023 (WOA23) annual climatology (nitrate, phosphate, silicate, dissolved oxygen) | NCEI/NOAA<sup>20</sup> | https://www.ncei.noaa.gov/access/world-ocean-atlas-2023/ |
| Mixed layer depth climatology | de Boyer Montégut et al.<sup>21</sup> | https://doi.org/10.17882/91774 |
| TARA Oceans station registry | PANGAEA<sup>3</sup> | https://doi.org/10.1594/PANGAEA.842237 |
| Pfam2GO mappings | Gene Ontology Consortium<sup>70,71</sup> | http://current.geneontology.org/ontology/external2go/pfam2go |
| Domain–environment association tables and modeling results (Data S1) | This study; Zenodo<sup>72</sup> | DOI: https://doi.org/10.5281/zenodo.18538438 |
| LA<sup>4</sup>SR-classified algal protein sequences (Data S2) | This study; Zenodo<sup>73</sup> | DOI: https://doi.org/10.5281/zenodo.18728836 |
| Pfam-A hmmsearch results archive (Data S3) | This study; Zenodo<sup>74</sup> | DOI: https://doi.org/10.5281/zenodo.18786751 |
| RuBisCO lineage analysis package (Data S4) | This study; Zenodo<sup>75</sup> | DOI: https://doi.org/10.5281/zenodo.18786775 |
| AlphaEarth satellite embedding matrix (Data S5) | This study; Zenodo<sup>76</sup> | DOI: https://doi.org/10.5281/zenodo.18786761 |

| **RESOURCE** *(continued)* | **SOURCE** | **IDENTIFIER** |
| --- | --- | --- |
| KAN-CCA and sparse CCA results (Data S6) | This study; Zenodo<sup>77</sup> | DOI: https://doi.org/10.5281/zenodo.18786766 |
| Pfam-dark protein-family discovery results incl. environmental correlations, ESMFold structures, and Foldseek homology searches (Data S7) | This study; Zenodo<sup>78</sup> | DOI: https://doi.org/10.5281/zenodo.18786771 |
| DIAMOND BLASTp comparison results (Data S8) | This study; Zenodo<sup>64,79</sup> | DOI: https://doi.org/10.5281/zenodo.19441355 |
| Dark-proteome selection analysis and 15M dark/annotated GPLM training sequences (Data S9) | This study; Zenodo<sup>80</sup> | DOI: https://doi.org/10.5281/zenodo.20486973 |
| Boltz-2 and AlphaFold 3 structure predictions for 3,600 Pfam-dark protein-family representatives (Data S10) | This study; Zenodo<sup>43,44,81</sup> | DOI: https://doi.org/10.5281/zenodo.20508680 |

| RESOURCE (*continued*) | SOURCE | IDENTIFIER |
| --- | --- | --- |
| Protein language model learnability analysis: code and results, checkpoints, and training data (Data S11) | This study; Zenodo,<sup>63</sup><br>Hugging Face<sup>62,82</sup> | Code and results: https://doi.org/10.5281/zenodo.20933332; checkpoints: https://huggingface.co/SarahDaakour/dark-whiteGPLM; training data: https://huggingface.co/datasets/SarahDaakour/dark-whiteGPLM-data |
| **Software and algorithms** |  |  |
| LA<sup>4</sup>SR/algaGPT classifier | Nelson et al.<sup>16,60</sup> | https://huggingface.co/GreenGenomicsLab/algaGPT |
| TARA-XGBoost-Bidirectional (bidirectional models) | This study<sup>61</sup> | https://huggingface.co/GreenGenomicsLab/TARA-XGBoost-Bidirectional |
| TARA-WorldModel-VICReg (joint embedding) | This study<sup>47</sup> | https://huggingface.co/GreenGenomicsLab/TARA-WorldModel-VICReg |
| HMMER (v3.3; hmmbuild v3.3.2) | Eddy Lab<sup>19,83</sup> | http://hmmer.org/ |
| SNAP gene predictor | Korf Lab<sup>84</sup> | https://github.com/KorfLab/SNAP |
| seqtk | Heng Li | https://github.com/lh3/seqtk |
| MAFFT (v7) | Katoh and Standley<sup>85</sup> | https://mafft.cbrc.jp/alignment/software/; DOI: 10.1093/molbev/mst010 |

| **RESOURCE** *(continued)* | **SOURCE** | **IDENTIFIER** |
| --- | --- | --- |
| MMseqs2 | Steinegger and Söding<sup>42</sup> | https://github.com/soedinglab/MMseqs2; DOI: 10.1038/nbt.3988 |
| InterProScan (v5.74-105.0) | EMBL-EBI<sup>86</sup> | https://interproscan-docs.readthedocs.io/ |
| DIAMOND | Buchfink et al.<sup>64</sup> | https://github.com/bbuchfink/diamond; DOI: 10.1038/nmeth.3176 |
| Foldseek | van Kempen et al.<sup>45</sup> | https://github.com/steineggerlab/foldseek; DOI: 10.1038/s41587-023-01773-0 |
| Boltz-2 (v2.2.1) | Passaro et al.<sup>43</sup> | https://github.com/jwohlwend/boltz |
| ESMFold API (v1) | Lin et al.<sup>8</sup> | https://api.esmatlas.com/foldSequence/v1/pdb/ |
| CD-HIT | Fu et al.<sup>87</sup> | https://github.com/weizhongli/cdhit; DOI: 10.1093/bioinformatics/bts565 |
| IQ-TREE 2 (v2.1.4-beta) | Minh et al.<sup>88</sup> | https://github.com/iqtree/iqtree2 |
| FastTree 2 | Price et al.<sup>89</sup> | http://www.microbesonline.org/fasttree/; DOI: 10.1371/journal.pone.0009490 |
| Biopython | Cock et al.<sup>90</sup> | https://biopython.org/;<br>DOI: 10.1093/bioinformatics/btp163 |
| PyMOL Open-Source | Schrödinger, LLC | https://github.com/schrodinger/pymol-open-source |
| Python (v3.10.12) | Python Software Foundation<sup>91</sup> | https://www.python.org/;<br>RRID:SCR_008394 |
| PyTorch (v2.1) | Meta AI | https://pytorch.org/;<sup>92</sup>;<br>RRID:SCR_018536 |
| nanoGPT | Karpathy<sup>41</sup> | Code snapshot archived with Data S11; DOI: 10.5281/zenodo.20933332 |
| HuggingFace Transformers | Hugging Face<sup>93</sup> | https://huggingface.co/docs/transformers; DOI: 10.18653/v1/2020.emnlp-demos.6 |
| XGBoost (v1.7.4) | XGBoost Developers | https://xgboost.readthedocs.io/;<sup>22</sup> |
| SHAP (v0.41.0) | Lundberg & Lee | https://shap.readthedocs.io/;<sup>23</sup> |
| UMAP (v0.5.3) | McInnes et al. | https://umap-learn.readthedocs.io/;<sup>28</sup> |
| scikit-learn (v1.2.2) | scikit-learn Developers | https://scikit-learn.org/;<sup>94</sup>; RRID:SCR_002577 |
| hdbscan | McInnes et al.<sup>33</sup> | https://github.com/scikit-learn-contrib/hdbscan |

| RESOURCE (continued) | SOURCE | IDENTIFIER |
| --- | --- | --- |
| SciPy (v1.9.3) | SciPy Developers | https://scipy.org/; <sup>95</sup>;<br>RRID:SCR_008058 |
| NumPy (v1.24.3) | NumPy Developers | https://numpy.org/; <sup>96</sup>;<br>RRID:SCR_008633 |
| pandas (v1.5.3) | pandas Development Team | https://pandas.pydata.org/; <sup>97</sup>; RRID:SCR_018214 |
| statsmodels (v0.13.5) | statsmodels Developers | https://www.statsmodels.org/; <sup>98</sup>; RRID:SCR_016074 |
| Google Earth Engine Python API | Google<sup>2</sup> | https://developers.google.com/earth-engine/; DOI: 10.1016/j.rse.2017.06.031 |
| xarray | Hoyer and Hamman<sup>99</sup> | https://xarray.dev/; DOI: 10.5334/jors.148 |
| t-SNE implementation | scikit-learn | 100 |
| AlphaFold 3 (v3.0.1) | Google DeepMind<sup>44</sup> | https://github.com/google-deepmind/alphafold3 |
| Analysis code (Code S1; 371 scripts, 15 modules) | This study; Zenodo<sup>59</sup> | DOI: https://doi.org/10.5281/zenodo.22910378 |
| Slurm Workload Manager | SchedMD | https://slurm.schedmd.com/ |
| **Other** |  |  |
| NVIDIA A100 graphics processing unit (GPU; 40 GB) | NVIDIA | LA<sup>4</sup>SR inference, neural network training, statistical analyses |
| NVIDIA GB10 GPU | NVIDIA | Boltz-2 structure prediction |

| RESOURCE *(continued)* | SOURCE | IDENTIFIER |
| --- | --- | --- |
| NVIDIA RTX A6000 GPU<br>(48 GB) | NVIDIA | AlphaFold 3 structure prediction |

### Study design

This study is entirely computational and does not involve experimental models, organisms, or study participants. All analyses use publicly available metagenomic sequence data from prior ocean sampling campaigns (TARA Oceans, Ocean Sampling Day, MMETSP) and satellite-derived environmental data products. No new biological samples were collected or processed.

### Method details

#### Reference dataset assembly

The broad input collection comprised 2,357 proteomes: 1,203 ocean metagenome assemblies, 676 MMETSP transcriptomes, and 478 cultured reference proteomes. After LA^4^SR classification and the requirement for non-empty Pfam-A results, 2,044 samples entered the domain-analysis set: 1,005 ocean metagenomes, 606 MMETSP transcriptomes, 236 NCBI reference genomes, and 197 other cultured references (Table S1; Figure S6). This analysis set contained 231,673,564 proteins assigned to the algal class. Counts for the broader input collection and the domain-analysis set therefore describe different stages of the pipeline.

Validation datasets comprised 1,154 reference proteomes organized into two categories: 478 cultured reference proteomes from genome assemblies (284 downloaded from NCBI GenBank and RefSeq^65^ and 194 published in our previous genome sequencing studies^66,67^ or in the original genome publications; accessions and the publishing study for every assembly are listed in Table S14) plus 676 transcriptomes from the Marine Microbial Eukaryote Transcriptome Sequencing Project (MMETSP).^18^ Reference proteomes span major algal lineages (Chlorophyta, Stramenopiles, Haptophyta, Cryptophyta) with verified taxonomic assignments.

#### algaGPT classification

Protein sequences were classified using algaGPT, a 70-million-parameter GPT-2-based transformer classifier implemented in the LA^4^SR (Language modeling with AI for algal amino acid Sequence Representation) framework and previously described and validated in Nelson et al.^16^ The character-level tokenizer maps each amino acid to one token. For each query sequence, the model autoregressively generates a terminal marker token encoding an algal or non-algal (bacterial, viral, or fungal) class; no separate classification head is used.

We used the algaGPT decoding mode, which restricts each decoding step to the ten highest-probability tokens before multinomial sampling (do_sample=True, temperature=0.1, top_k=10). Top-*k* sampling controls how LA^4^SR generates the terminal class marker; it is not a separate classifier. Across 222 algal reference genomes, this mode retained 83.3% of proteins (16.7% unretained), while precision for the bacterial class exceeded 95% in the matched benchmark.^16^ Published whole-proteome tests generated predictions for *>*99% of sequences across 166 microalgal genomes; error rates were lineage-dependent, with up to 10.56% *±* 0.36% of bona fide algal proteins called contaminant-like in the most challenging tested lineage.^16^ These benchmark rates describe reference proteomes and need not transfer unchanged to mixed environmental assemblies.

Inference parameters: max_new_tokens=14, batch size 512, maximum sequence length 512 tokens. Sequences exceeding this length were truncated, meaning classification was based on the N-terminal portion. Inference used mixed-precision (float16) computation via Slurm Workload Manager job arrays, processing 447.7 million sequences at *∼*280 sequences/second (hardware and runtime summary in Statistical analysis).

#### DIAMOND homology comparison

To compare database representation between LA^4^SR classes, we searched 352,615,196 proteins from the 2,044-sample domain-analysis set against NCBI NR using DIAMOND BLASTp^64^ in very-sensitive mode (*E <* 10^−9^). This comparison subset excludes proteins from the 313 input samples that did not enter the Pfam/domain analysis and is therefore smaller than the 447.7-million-protein classifier input. We compared the proportions of algal- and non-algal-classified sequences with at least one NR homolog; complete per-sample results and execution logs are provided in Data S8.

##### Operational scope of the “algal” classification

LA^4^SR is applied to marine metagenomic assemblies, where the non-target population consists of marine heterotrophic microeukaryotes rather than terrestrial model organisms. Within this operational scope, non-photosynthetic marine protists (including apicomplexans, plastid-loss dinoflagellates, choanoflagellates, kinetoplastids, and kleptoplastic ciliates) share phylogenetic proximity, evolutionary history, and compositional features with algae. Many such lineages have photosynthetic ancestry and retain genomic signatures of algal heritage. When LA^4^SR includes such organisms in the “algal” fraction, this represents a slight scope widening from strict photosynthetic eukaryotes to photosynthetic eukaryotes plus their close marine protistan relatives, a biologically coherent unit representing the algal-derived functional repertoire in the marine plankton. Benchmarks against phylogenetically distant eukaryotic references (fungi, metazoans, non-algal protists from terrestrial culture collections) are included as generalization probes but lie beyond the realistic operational scope.

#### Gene prediction and Pfam domain annotation

Protein-coding genes were predicted from metagenomic assemblies using SNAP (Semi-HMM-based Nucleic Acid Parser)^84^ trained on *Arabidopsis thaliana* gene models, validated for cross-clade microalgal annotation across 22 phylogenetically diverse reference genomes in Nelson et al.,^66^ achieving *>*90% sensitivity and *>*85% specificity across Chlorophyta, Streptophyta, Stramenopiles, Haptophyta, Rhodophyta, and Cryptophyta, including both intron-rich (*Chlamy-domonas*, 7.4 introns/gene) and intron-poor (*Ostreococcus*, 0.3 introns/gene) architectures. SNAP is an *ab initio* predictor that models eukaryotic gene structure (splice sites, exon–intron transitions, start/stop codons) from sequence alone, without reference database dependence. Because our study targets the dark proteome, the 86.9% of proteins lacking any Pfam hit, *ab initio* prediction is the appropriate design: reference-based eukaryotic gene callers (e.g., MetaEuk^101^) retrieve only proteins with detectable homology to existing databases and would systematically exclude novel protein clusters from the pipeline by construction, biasing the analysis toward the annotated fraction we are trying to extend. SNAP operates upstream of LA^4^SR, so sequences assigned to the non-algal class do not enter downstream Pfam analysis; the benchmark rates above delimit the expected classification error. The 1,039 MMETSP transcriptome and reference genome samples (606 + 433, respectively) bypass SNAP entirely, using gene calls from transcript assembly or genome annotation and providing an internal control: coupling patterns are consistent across SNAP-predicted and independently called gene sets (TARA-only vs. full-dataset comparison; Results).

On fragmented metagenomic contigs (*<*1 kb), SNAP’s sensitivity is reduced because intact intron–exon boundary signals may be truncated. Because assembly contiguity correlates with sequencing depth and organism abundance, the resulting Pfam domain matrix is biased toward well-assembled, high-abundance taxa. This bias is partially mitigated by the domain-level unit of analysis: a domain need only be detected once per sample to contribute to the count matrix, so missed gene copies reduce abundance estimates but are unlikely to produce false absences for domains carried by abundant organisms.

Domain annotation was performed using hmmsearch (HMMER v3.3)^19^ against Pfam-A v37.2 (November 2024; 24,076 families).^12,68^ Two thresholds served distinct analytical purposes. The strict screen (*E <* 10^−9^) defined the Pfam-*dark* proteome and detected 17,245 Pfam families across 2,318 assemblies. The permissive screen (*E <* 10^−5^) maximized sensitivity for domain-abundance and domain–environment analyses across the 2,044-sample non-empty analysis set. Its merged matrix contained 20,317 valid versioned Pfam accessions plus one additional unversioned duplicate identifier (`PF00128`), yielding 20,318 raw feature columns. Domain counts per genome or assembly retained integer hit counts per feature and sample. Subset-specific prevalence filters then retained 11,898 features present in at least 10 samples in the AEF correlation subset (*n* = 995), 9,611 features present in at least 5% of the 1,809-sample complete-coordinate matrix (used for CCA, GO enrichment, and projection modeling), and 9,989 features in the 1,810-sample GEE subset used for forward modeling. The stricter threshold was used for the dark-proteome definition to restrict the annotated fraction to high-confidence matches; sensitivity analysis at *E <* 10^−5^ confirmed that the Pfam-dark-family effect size was stable (2.29-fold enrichment at the primary family-profile threshold of *E <* 10^−9^ vs. 2.32-fold at *E <* 10^−5^; Results). An output-integrity audit identified 45 of the 1,809 samples (2.49%) whose HMMER tabular outputs lacked the terminal `# [ok]` completion marker and ended mid-record. We therefore repeated the prevalence filter and CLR–PCA–CCA workflow after excluding these samples. Among the remaining 1,764 samples, 9,616 domains passed the recalculated *≥*5% threshold; 9,600 of the original 9,611 domains were retained (99.89%; feature-set Jaccard index = 0.997), and the first canonical correlation changed from 0.816 to 0.820 (Δ*r* = 0.005).

#### Lineage-specific RuBisCO HMM construction

To enable taxonomic assignment of photosynthetic lineages within extracted proteomes, we constructed lineage-specific profile HMMs for ribulose-1,5-bisphosphate carboxylase/oxygenase (RuBisCO) large subunit (rbcL) sequences. Reference rbcL protein sequences were retrieved from NCBI for ten Form I algal lineages: Mamiellophyceae (28 sequences), Prasinophyceae *sensu lato* (50), Pyramimonadales (30), Chlorellaceae (50), Trebouxiophyceae (100), Scenedes-maceae (50), Pelagophyceae (50), Bolidophyceae (20), Haptophyta (50), and Cryptophyta (50), totaling 478 reference sequences (Data S1). Sequences shorter than 300 amino acids were removed. Multiple sequence alignments were generated for each lineage using Multiple Alignment using Fast Fourier Transform (MAFFT) L-INS-i^85^ `(–localpair –maxiterate 1000`), and profile HMMs were built using `hmmbuild` (HMMER v3.3.2). Combined “green lineage” (Form IB RuBisCO: Chlorophyta) and “red lineage” (Form ID: Stramenopiles, Haptophyta, Cryptophyta) HMMs were constructed from pooled alignments for broad screening. Lineage assignment used hierarchical searching: sequences were first screened against broad HMMs (*E <* 10^−7^), then scored against all lineage-specific HMMs (*E <* 10^−10^) and assigned to the lineage with highest bit score.

To capture dinoflagellate and chromerid phototrophs, which encode Form II RuBisCO in the nuclear genome rather than the plastid genome,^24^ we constructed five additional lineage-specific HMMs: Symbiodiniaceae (284 filtered sequences after polyprotein splitting), Peridiniales *sensu lato* (296), Gonyaulacales (235), Prorocentrales (243), and Chromerida (125), plus a broad Form II HMM pooling all eukaryotic Form II sequences with proteobacterial outgroup representatives (clustered at 85% identity). Dinoflagellate Form II rbcL is frequently encoded as polyproteins containing 2–4 tandem copies of the *∼*470-amino-acid monomer;^26^ sequences exceeding 600 aa were split into individual monomer units before alignment. Sequences were clustered at 90% identity using CD-HIT,^87^ aligned with MAFFT L-INS-i, and HMMs built with `hmmbuild`. Because Form II rbcL is nuclear-encoded, LA^4^SR may not classify dinoflagellate nuclear contigs as “algal” (the classifier targets chloroplast gene structure signatures); therefore, the extended search queries *all* predicted proteins per assembly rather than only algal-classified proteins, ensuring global detection of nuclear-encoded photosynthetic markers.

#### AEF embedding extraction

Environmental embeddings were extracted via Google Earth Engine’s Python application programming interface (API) using the AEF annual satellite embedding collection.^2^ This collection provides 64-dimensional embeddings (A00–A63) derived from annual satellite imagery syntheses (Earth Engine collection `GOOGLE/SATELLITE_EMBEDDING/V1/ANNUAL`, 2017 onward; all available annual layers were combined with `ImageCollection.mosaic()` before extraction). A fundamental caveat applies to all satellite-derived environmental variables in this study: the 27 GEE variables are multi-year composites (typically 2013–2021 averaging windows), the 5 WOA23/MLD variables are 1991–2020 decadal climatologies, and the AEF embeddings derive from annual satellite syntheses for 2017 onward. None represents conditions at the time of biological sampling (TARA Oceans 2009–2013, OSD 2014, MMETSP various years). Domain– environment associations therefore reflect coupling to long-term mean environmental state, not contemporaneous conditions. Temporal mismatch may attenuate relationships for variable features and can also preserve or confound stable spatial gradients; it does not guarantee a lower bound on true coupling. Correlations are 2.4-fold stronger for temporally stable variables (median *|ρ|* = 0.14, 75.9% FDR-significant) than for temporally varying ones (median *|ρ|* = 0.06, 39.2% significant; *p <* 10^−16^), and temporal stability under year-based holdout is assessed in the Supplemental Text. Extraction used 10-meter spatial resolution with mean value aggregation at each GPS coordinate. Batch processing (100 samples per request with 2-second rate limiting) yielded AEF records for 995 of 1,810 GPS-mapped samples (55.0%); 969 had no zero-filled dimensions and 26 were partially zero-filled during downstream feature preparation. Missing records were concentrated in open-ocean locations (*>*50 km offshore, *>*200 m depth), where the AEF model’s terrestrial/coastal training data provides insufficient coverage. Coastal and shallow-water samples had record coverage *>*90%.

#### Nutrient and mixed-layer-depth extraction

Dissolved inorganic nutrient concentrations (nitrate, phosphate, silicate) and dissolved oxygen were obtained from the World Ocean Atlas 2023 (WOA23) objectively analyzed annual climatology at 1° spatial resolution.^20^ For each variable, the surface-layer (0 m depth) annual mean field was extracted from the 1991–2020 decadal average climatology (“decav91C0” product). Mixed layer depth (MLD) was obtained from the de Boyer Montégut climatology,^21^ defined using a density threshold criterion (0.03 kg/m^3^ from a 10 m reference depth); monthly MLD fields were averaged to obtain annual mean values. For each of the 1,810 GPS-mapped samples, nutrient and MLD values were extracted by nearest-neighbor lookup on the respective climatological grids using `xarray`.^99^ Surface values were used throughout, consistent with the median TARA sampling depth of 5 m. These five variables (nitrate, phosphate, silicate, dissolved oxygen, MLD) supplement the 30 GEE-derived environmental variables and 2 sample metadata variables (sampling depth, estimated salinity), yielding 37 interpretable oceanographic variables total.

#### Domain-environment correlation analysis

Spearman rank correlations were computed between each Pfam domain count vector and each AEF embedding dimension. Because Pfam abundances are compositional data (counts per assembly subject to a sample-dependent total), correlations on raw counts may be affected by spurious associations induced by the constant-sum constraint.^102^ The initial discovery screen uses raw domain counts with Spearman correlation; a sensitivity analysis using centered log-ratio (CLR)-transformed abundances (Data S1) showed that the top-ranked domains differ between normalizations (Jaccard = 0.21 at the top-50 level; mean *|*Δ*ρ|* = 0.066), while broader significance counts are comparable (65.6% agreement rate across 279,834 raw vs. 268,909 CLR FDR-significant associations in this sensitivity run, which tested 13,335 domains rather than the 11,898 of the primary screen). This analysis quantifies the effect of normalization rather than assuming invariance: CLR is computed within samples and is not a monotonic transformation of each domain vector across samples. All major quantitative findings (XGBoost bidirectional *R*^2^ values, CCA canonical correlations, and SHAP importance rankings) use CLR-transformed Pfam inputs (pseudocount = 0.5 or 1; see respective subsections). CLR normalization removes between-sample differences in total domain yield but does not correct domain-specific copy-number amplification within a genome (e.g. multi-copy Form II rbcL in dinoflagellates or tandem transposase arrays), which would require per-cell normalization data. Lineage-resolved partial correlations (controlling for five dinoflagellate orders; see below) address the measured taxonomic-sorting component. For paired observations (*X_i_*, *Y_i_*), the Spearman correlation coefficient is:

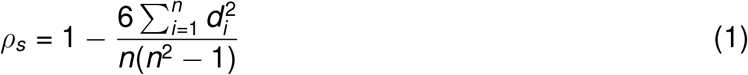

where *d_i_* = rank(*X_i_*) *−* rank(*Y_i_*) is the difference in ranks and *n* is the sample size. Computations used `scipy.stats.spearmanr`^95^ with `nan_policy=’omit’`. Multiple testing correction employed Benjamini-Hochberg FDR^27^ via `statsmodels.stats.multitest.fdrcorrection`. Associations with FDR *<* 0.05 were considered significant.

To quantify the correlation structure among tests, we computed the effective number of independent tests (M_eff_) using eigenvalue decomposition of both the Pfam correlation matrix (9,466 domains at *≥*5% prevalence) and the AEF embedding correlation matrix (64 dimensions). M_eff_ was calculated using the Nyholt formula:^103^

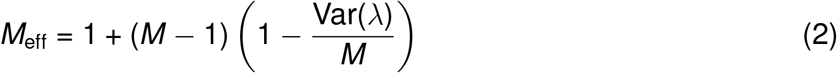

where *M* is the number of variables and *λ*_1_, *…*, *λ_M_* are the eigenvalues of the correlation matrix. Total effective tests were computed as M_eff,PFAM_ *×* M_eff,AEF_. The M_eff_ correction assumes that the correlation structure among Pfam domains is stable across samples; systematic differences between source categories (ocean metagenomes, MMETSP transcriptomes, cultured reference proteomes), ocean basins, and environmental regimes may violate this assumption. To assess sensitivity, we repeated the correlation analysis stratified by data source (Data S1). For family-wise error rate (FWER) control, we performed 1,000 permutation tests shuffling sample labels while preserving correlation structure, recording the maximum *|ρ|* per permutation to establish the 95th percentile threshold.

#### XGBoost validation

XGBoost gradient boosting regressors validated correlation-based associations by predicting AEF embedding dimensions from Pfam domain composition (AEF subset, *n* = 995).^22^ Note: this section describes the embedding dimension validation models; the bidirectional environmental prediction models (see Bidirectional predictive modeling, below) use a separate 10-fold spatial block CV design on the larger GEE dataset. XGBoost minimizes a regularized objective function:

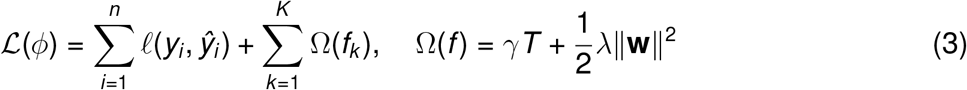

where *ℓ* is a differentiable loss function, 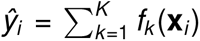 is the prediction from *K* additive trees, and Ω(*f*) penalizes model complexity via the number of leaves *T* and leaf weights **w**. Model parameters: max depth 6, learning rate 0.1, 200 estimators, subsample 0.8, column sample by tree 0.8, min child weight 3, L1 regularization 0.1, L2 regularization 1.0, random seed 42. Five-fold cross-validation assessed predictive performance (*R*^2^). Hyperparameters were fixed across all models based on preliminary tuning on a held-out development set (Atlantic samples); the Atlantic was chosen because it contains the most samples (644, 35.6% of total), providing the most statistically stable tuning estimates. This introduces potential overfitting to Atlantic-specific characteristics; however, the 10-fold spatial block cross-validation design, which evaluates generalization to geographically separated grid cells, serves as the primary test of geographic robustness. Sensitivity analysis varying max_depth (4–8), learning rate (0.05–0.2), and n_estimators (100–500) showed stable *R*^2^ estimates with maximum range *<*0.09 across all six evaluated targets (Data S1), indicating results are robust to hyperparameter choice within reasonable ranges. Because conclusions depend on relative effect sizes across validation designs rather than absolute *R*^2^, this insensitivity to hyperparameter choice ensures that the scientific interpretation is not contingent on the Atlantic-derived tuning.

Feature importance was determined via SHAP (SHapley Additive exPlanations) values,^23^ which decompose predictions into additive feature contributions based on cooperative game theory. For a prediction *f* (**x**), the SHAP value for feature *j* is:

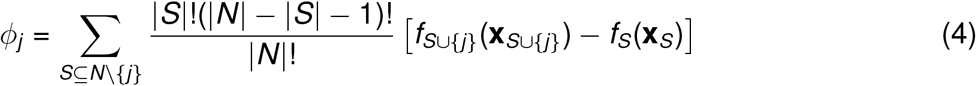

where *N* is the set of all features, *S* is a subset excluding feature *j*, and *f_S_* is the model prediction using only features in *S*. SHAP values satisfy local accuracy 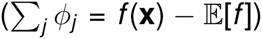, consistency, and missingness properties.

#### Dimensionality reduction

UMAP and t-SNE embeddings (Figure S1D–K) were computed on the permissive Pfam-A count matrix (20,318 raw feature columns *×* 1,810 GPS-mapped samples; per-sample integer counts with missing values set to zero, without standardization) to visualize structure in protein composition space.^28,100^ Principal Component Analysis (PCA) was computed using `sklearn.decomposition.PCA`^94^ with 50 components retained. UMAP (Uniform Manifold Approximation and Projection) constructs a fuzzy topological representation of the high-dimensional data and optimizes a low-dimensional embedding by minimizing the cross-entropy:

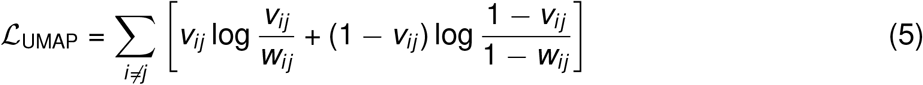

where *v_ij_* and *w_ij_* are fuzzy set membership strengths in high- and low-dimensional spaces, respectively.^28^ UMAP used `umap-learn` with n_neighbors = 15, 30, and 50, min_dist=0.1, metric=‘euclidean’, and random_state=42. t-SNE (t-distributed Stochastic Neighbor Embedding) minimizes the Kullback-Leibler divergence between probability distributions over pairwise similarities:

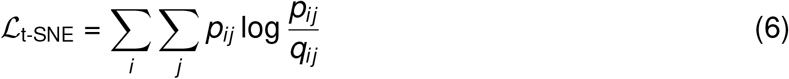

where *p_ij_*are Gaussian-based similarities in high-dimensional space and *q_ij_*are Student-t-based similarities in the embedding.^100^ t-SNE was computed using sklearn.manifold.TSNE at perplexities 5, 30, and 50 (1,000 iterations, random_state=42; Figure S1H–K).

#### Canonical correlation analysis

Canonical Correlation Analysis (CCA)^29^ jointly embedded Pfam and environmental matrices to identify maximally correlated linear combinations. CCA seeks weight vectors **a** and **b** that maximize:

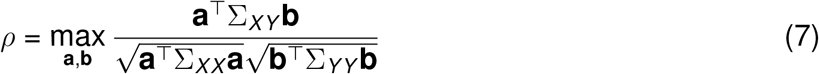

where Σ*_XY_* is the cross-covariance matrix between Pfam (*X*) and environment (*Y*), and Σ*_XX_*, Σ*_Y Y_* are the respective within-set covariance matrices. CCA significance was evaluated with two distribution-free permutation tests; the parametric Wilks’ Λ approximation was not used because multivariate normality was not expected for compositional Pfam data. The tests were (1) standard row shuffling (1,000 iterations, shuffling sample–environment pairings while preserving within-matrix correlation structure) and (2) spatial-block shuffling (1,000 iterations, permuting entire 2° grid cells to preserve within-block spatial autocorrelation under the null). Both tests report the full null distribution of CC1 (mean, SD, maximum) and compute permutation *p*-values as the fraction of permuted CC1 values *≥* the observed CC1. The spatial-block test is the conservative estimate, matching the 2° spatial block CV design used for XGBoost validation. CCA was implemented via `sklearn.cross_decomposition.CCA` with 10 components and default internal scaling (`scale=True`, which centers and unit-variance-normalizes both views prior to fitting). The environmental matrix (*n* = 1,809 samples *×* 31 variables; 30 GEE features plus sampling depth, with estimated salinity dropped for complete missingness) was z-score standardized; the Pfam matrix was CLR-transformed (pseudocount = 1) and reduced to 100 principal components via PCA (capturing 76.0% of total variance in the 9,611-domain CLR matrix) before input to CCA. The 100-component threshold balances variance retention against the *n ≫ p* + *q* requirement for stable CCA estimation (*n* = 1,809 versus *p* + *q* = 131); sensitivity analysis at 50, 100, 150, and 200 components confirmed stable CC1 (0.79–0.85; Supplemental Text). Environmental loadings on canonical components were extracted to identify variables most strongly coupled to domain composition. The nutrient-expanded L1-penalized sparse CCA used *n* = 786 samples and 24 environmental variables, including dissolved nitrate, phosphate, silicate, oxygen, and mixed layer depth from WOA23, with the same CLR + PCA(100) domain preprocessing. The KANCCA analysis and domain-sparsity ablation plotted in Figure 2G–T used the earlier satellite-only complete-case matrix (*n* = 969; 19 environmental variables) and ten 2° spatial-block folds. These two analysis variants are reported separately in Data S6 and Supplemental Text.

#### Bidirectional predictive modeling

To validate domain-environment associations beyond correlation, we trained XGBoost regressors in two directions. Reverse modeling predicted each of 37 environmental variables from Pfam domain composition (*n* = 1,064–1,600 depending on target variable completeness) using 10-fold spatial block cross-validation (2° grid cells assigned to folds, ensuring geographically proximate samples are in the same fold). Forward modeling (environment *→* individual domain abundance) used a two-tier design because fitting spatial block CV models for all 9,989 domains independently would require *∼*100,000 model fits. First, all 9,989 domains were screened under standard 5-fold CV (*n* = 1,279; 25 environmental features) to rank domains by out-of-sample *R*^2^. Second, the 18 top-ranked domains were re-evaluated under 10-fold spatial block CV in two analysis-specific datasets: the WOA23-merged, basin-resolved set (*n* = 1,810; Table S2) and the later GPS-recovered, satellite-only set (*n* = 1,878; Figure 5A–E). The latter includes 68 samples for which coordinates were recovered after the frozen basin table had been generated; those samples therefore enter the satellite-only analysis but not analyses requiring a named basin assignment. The initial screen used the initial algaGPT extraction (*n* = 1,279 after requiring complete satellite-derived environmental variables). *R*^2^ values for individually named domains are linked to their stated dataset (Figure 5 or Table S2), whereas aggregate statistics (e.g., 65 domains exceeding *R*^2^ *>* 0.3; fold-bootstrap 95% CI: 51–118, *B* = 10,000) derive from the broader 5-fold screen. The spatial-validation values (0.27–0.59) condition on target selection in an overlapping cohort and therefore characterize the selected top tail rather than providing selection-independent estimates; the maximum may be optimistic. Sensitivity analyses at 5° and 10° grid sizes, a latitude-only baseline, metagenome-only subsetting, permutational multivariate analysis of variance (PERMANOVA) batch-effect testing, and cross-basin holdout are reported in Supplemental Text. As a secondary evaluation, data were also split 80/20 for training/testing with stratification by latitude quartile to ensure both training and test sets span comparable thermal and light regimes, since latitude is the dominant proxy for the primary environmental gradients (temperature, photoperiod) and prevents the test set from being drawn entirely from a single climatic zone. This stratification approach complements the 10-fold spatial block cross-validation (primary design), which ensures geographically proximate samples are evaluated together. Model parameters were identical to the embedding dimension validation above. SHAP values were computed using TreeExplainer to identify the most predictive features across all models.

#### Joint embedding approach (VICReg)

As an alternative to the XGBoost bidirectional framework, we explored a self-supervised joint embedding approach using VICReg (Variance-Invariance-Covariance Regularization)^46^ to align environmental and Pfam representations in a shared latent space. A two-layer MLP encoder (*∼*53K–64K parameters; latent dimension = 32; dropout = 0.3; VICReg loss weights: variance = 25.0, invariance = 25.0, covariance = 1.0) was trained on 1,151 samples with complete productivity data (chlorophyll-*a*, POC, NFLH). Under 6-fold leave-one-basin-out (LOBO) cross-validation, the joint model achieved POC *R*^2^ = 0.532 versus an environment-only XGBoost baseline of *R*^2^ = 0.422, but the improvement was not statistically significant (Cohen’s *d* = 0.026, *p* = 0.38). Under 9-fold spatial block cross-validation (the primary XGBoost models used 10 folds), the MLP architecture produced negative *R*^2^ on held-out basins (POC *R*^2^ = *−*2.05 to *−*4.22 across Pfam dimensionalities of 20, 32, and 64), driven by catastrophic generalization failures on spatially distinctive basins (Mediterranean, mid-Pacific). The architecture confound (XGBoost’s tree-based partitioning handles distribution shift in small tabular datasets [*n ≈* 1,100] more effectively than shallow neural networks) was the dominant factor, rather than absence of the Pfam alignment signal per se. We retained the XGBoost bidirectional framework as the primary modeling approach; VICReg checkpoints and hyperparameter sweep results are deposited for reproducibility.^47^

#### Lineage-resolved transposase analysis

To assess whether TE–environment coupling reflects finer-grained dinoflagellate taxonomic sorting rather than within-lineage TE variation, we performed three analyses using per-sample RuBisCO lineage counts from the 15-lineage HMM search (Data S4) matched to Pfam domain abundances (*n* = 1,438 samples with both Pfam profiles and RuBisCO assignments). Sample IDs were matched by stripping file-extension suffixes (`.fa.aa`, `.aa`) from the RuBisCO sample identifiers. The 11 TE domains were defined as the mobile genetic element domains among the 18 most environment-predictable (Table S2). Because dinoflagellate genomes are characterized by extensive polyploidy, large genome sizes (1–250 Gbp), and high transposon copy numbers,^31^ TE domain counts in dinoflagellate-dominated samples may be inflated by genome architecture independently of environmental selection; the partial correlation analyses below are designed to control for this taxonomic component.

##### TE–RuBisCO covariation

Spearman rank correlations were computed between each of the 11 TE domain abundances (CLR-transformed) and each of the 18 RuBisCO lineage counts across matched samples, yielding 198 tests. Bonferroni correction was applied for multiple testing.

##### Within-dinoflagellate variance

Among samples with detected Form II RuBisCO (*n* = 1,227), the standard deviation of each TE domain’s CLR-transformed abundance was computed and compared to the genome-wide distribution of all 20,318 domain SDs using a one-sided Mann– Whitney U test (alternative: TE domains show greater variance). Standard deviation rather than coefficient of variation was used because CLR-transformed values center near zero, rendering CV unreliable.

##### Lineage-controlled partial correlations

Partial Spearman correlations between each TE domain and four environmental variables (SST, bathymetry, air temperature, chlorophyll-a) were computed controlling for five Form II dinoflagellate lineage counts (Symbiodiniaceae, Peridiniales, Gonyaulacales, Prorocentrales, Chromerida) as continuous covariates. The rank-then-regress approach was used: all variables were rank-transformed, ordinary least squares residuals were computed for both the TE domain ranks and environmental variable ranks after regressing on the ranked covariates, and the partial correlation was taken as the Pearson correlation of the residuals. *P*-values were computed from the *t* -distribution with *n − k −* 2 degrees of freedom, where *k* = 5 covariates. Bonferroni correction was applied across all 44 tests.

#### Functional biomes

Functional biomes were defined by clustering Pfam composition profiles with HDBSCAN^33^ (min_cluster_size=20, min_samples=15) and comparing clusters to Longhurst biogeochemical provinces using Adjusted Rand Index (ARI) and Normalized Mutual Information (NMI).

#### GO enrichment analysis

Gene Ontology (GO) enrichment^70^ was performed using hypergeometric tests to identify overrepresented biological processes among environment-associated Pfam domains. The hypergeometric test calculates the probability of observing *k* or more successes when drawing *n* items from a population of *N* items containing *K* successes:

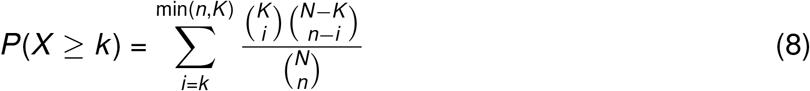

where *N* is the total number of Pfams in the background, *K* is the number of Pfams annotated with a given GO term, *n* is the size of the test set (environment-associated Pfams), and *k* is the observed overlap. Pfam2GO mappings were obtained from the Gene Ontology Consortium (http://current.geneontology.org/ontology/external2go/pfam2go), providing 9,842 Pfam-to-GO associations across 5,182 Pfam families. Pfam descriptions were retrieved from the InterPro API^86,104^ (https://www.ebi.ac.uk/interpro/api).

For each environmental group (temperature, depth, light, optical), the top 100 Pfams ranked by mean SHAP importance across relevant environmental variables were tested against the background of all 9,611 Pfams in the dataset. Environment-predictable Pfams (the 22 domains with forward test *R*^2^ *>* 0.3 in an earlier 100-Pfam single-split forward screen; distinct from the 9,989-domain five-fold screen) were analyzed as a separate set. P-values were computed using scipy.stats.hypergeom^95^ and corrected for multiple testing using Benjamini-Hochberg FDR via statsmodels.stats.multitest.fdrcorrection. GO terms present in fewer than 3 background Pfams were excluded from testing.

#### Pfam-to-AEF projection model

The Pfam-to-AEF neural projection model architecture and training procedure are described in the Supplemental Text. This model is not used for primary analyses; it serves as a methodological complement to the XGBoost and CCA frameworks, confirming that domain–environment coupling supports cross-modal prediction.

#### Environment-to-Pfam prediction model

As a complementary environment-to-Pfam model, we trained deep multilayer perceptrons mapping 94 environmental input features (30 Google Earth Engine variables and 64 AEF embedding dimensions) to CLR-normalized Pfam domain profiles. The permissive Pfam-A v37.2 matrix (*E <* 10^−5^; 20,318 raw feature columns across 2,044 samples) was used. Missing GEE values and absent or partially missing AEF dimensions were encoded as zero after feature preparation, without a separate missingness mask; among the 995 samples with AEF records, 969 had no zero-filled AEF dimensions and 26 were partially zero-filled, while 1,049 samples without AEF records had all 64 AEF inputs set to zero. Data were randomly split 70/15/15 for training, validation, and testing (*n*=1,430/306/308), and continuous inputs were z-score standardized using training-set statistics.

Two model architectures were evaluated: a light variant (94*→*256*→*512*→*1,024*→*2,048*→*output) and a full variant (94*→*512*→*1,024*→*2,048*→*4,096*→*output), each with batch normalization, rectified linear unit (ReLU) activations, and dropout (*p*=0.2) at every hidden layer, using Kaiming He initialization.^105^ Models were trained with AdamW optimizer^106^ (weight decay 10^−4^), mean squared error (MSE) loss, and learning rates of 10^−3^ and 10^−4^, for up to 200 epochs with early stopping (patience=20, min delta=10^−6^) on validation loss. All training used GPU-accelerated mixed-precision computation with batch size 32 and fixed random seed 42.

Model performance was evaluated on held-out test sets using global *R*^2^, mean cosine similarity, MSE, and mean absolute error (MAE). To assess domain-level predictability, predictions were binarized using the CLR zero-crossing threshold (CLR *>* 0 indicates domain abundance above the sample geometric mean), and per-domain area under the receiver operating characteristic curve (AUC) was computed for all domains with non-trivial prevalence. The best configuration (full architecture, lr=10^−3^) achieved test *R*^2^=0.475, cosine similarity=0.666, and mean per-domain AUC=0.598 across 15,665 evaluable domains, with 76.4% of domains exceeding random (AUC *>* 0.5), 31.2% achieving AUC *>* 0.7, and 175 domains (1.1%) reaching AUC *>* 0.9. Because the split was random and zero filling encodes data availability, this secondary exploratory model may learn coverage patterns and does not establish spatial transfer.

#### Pfam-dark protein-family discovery

To characterize the fraction of the algal proteome inaccessible to homology-based annotation, we developed a pipeline to identify recurrent whole-protein clusters in the Pfam-dark proteome, defined as proteins with no Pfam-A v37.2 hit at the strict threshold of *E <* 10^−9^. Because clustering used 80% bidirectional whole-sequence coverage and did not identify independent domain boundaries, we refer to these operational units as Pfam-dark protein families rather than novel domains. The pipeline operates in eight stages across the 2,044 mixed-source sample proteomes with non-empty Pfam-A results. The upstream HMMER processing archive contains outputs for 2,318 of the 2,357 input samples; 2,044 of those outputs met the non-empty Pfam-A inclusion criterion for the domain-analysis set.

##### Dark proteome extraction (Steps 1–2)

For each sample, protein IDs present in the predicted proteome FASTA but absent from the corresponding hmmsearch tblout (sorted set difference via `comm -23`) were classified as Pfam-dark. A protein with any Pfam-A hit, even a single domain, was excluded from the dark set. Per-sample arithmetic cross-checks verified that dark + annotated = total. Dark protein sequences were then extracted using `seqtk subseq` (with a streaming Python fallback for compatibility).

##### Sequence filtering (Step 3)

Dark sequences were filtered to remove artifacts (applied in order, each sequence counted under the first failing criterion):

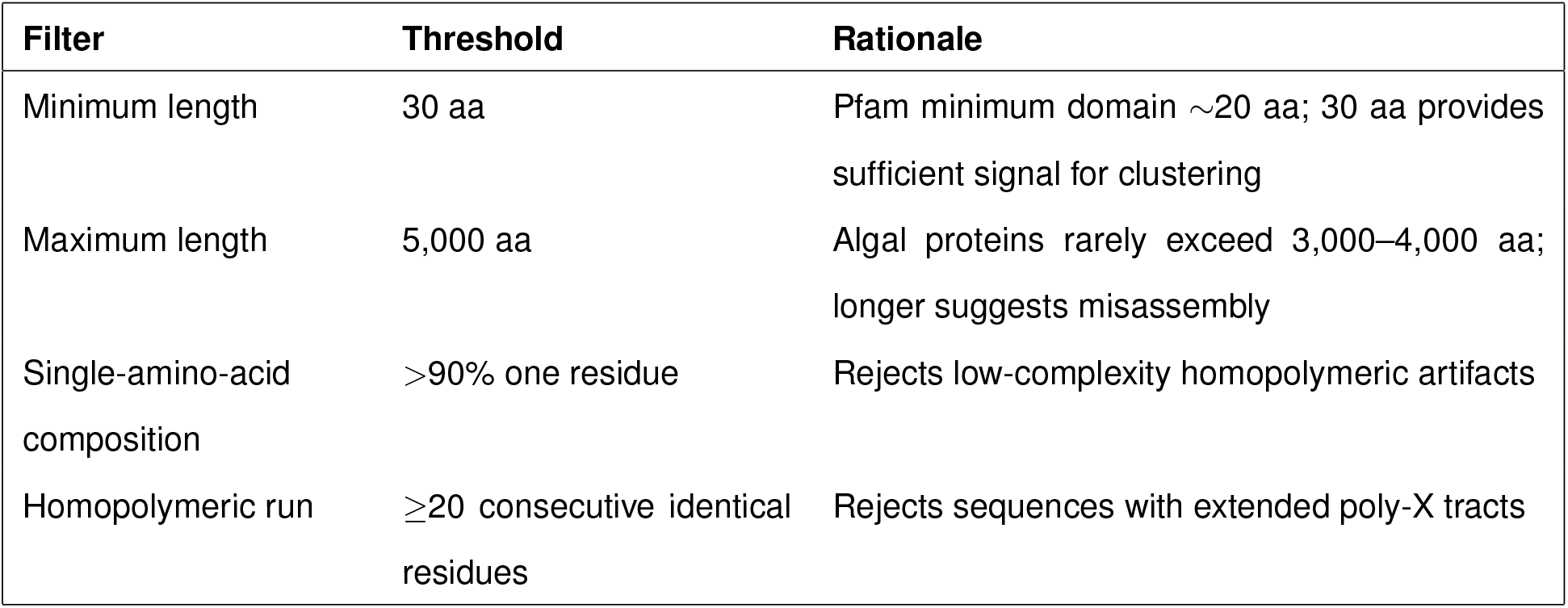

| Filter | Threshold | Rationale |
| --- | --- | --- |
| Minimum length | 30 aa | Pfam minimum domain ~20 aa; 30 aa provides sufficient signal for clustering |
| Maximum length | 5,000 aa | Algal proteins rarely exceed 3,000–4,000 aa; longer suggests misassembly |
| Single-amino-acid composition | >90% one residue | Rejects low-complexity homopolymeric artifacts |
| Homopolymeric run | $\geq 20$ consecutive identical residues | Rejects sequences with extended poly-X tracts |

Passing sequences from all 2,044 samples were concatenated into a single filtered dark proteome. Of the 201,336,126 input dark proteins, 201,179,616 passed all filters (99.9% pass rate); rejected sequences comprised 519 below the minimum length, 375 above the maximum length, 539 with low-complexity composition, and 155,077 with homopolymeric runs.

##### Clustering (Step 4)

The filtered dark proteome was clustered using MMseqs2 easy-linclust^42^ (a linear-time algorithm required for datasets exceeding 100 million sequences) at two identity thresholds:

| Parameter | Value | Rationale |
| --- | --- | --- |
| Identity threshold (Tier A) | 30% (`-min-seq-id 0.3`) | Twilight-zone boundary; captures broad whole-protein families |
| Identity threshold (Tier B) | 50% (`-min-seq-id 0.5`) | Subfamilies with higher functional coherence |
| Coverage | 80% (`-c 0.8`) | Ensures substantial alignment overlap |
| Coverage mode | Bidirectional (`-cov-mode 0`) | Both query and target must reach 80% coverage |
| Large cluster threshold | $\geq 50$ members | Minimum for reliable HMM profile construction |

At 30% identity, clustering produced 122,052,662 clusters from 201,179,616 filtered sequences, of which 1,324,206 contained *≥*10 members and 33,950 contained *≥*50 members.

##### HMM profile construction (Steps 5–5b)

For each cluster with *≥*50 members at 30% identity, member sequences were extracted and aligned with MAFFT (`–auto`, which adaptively selects L-INS-i for small clusters and FFT-NS-2 for large ones).^85^ Family-profile HMMs were built with `hmmbuild` (`–amino`), named `NOVEL_{representative_ID}` for traceability. Individual HMMs were concatenated into a single database and indexed with `hmmpress`. This yielded 33,950 Pfam-dark protein-family HMMs. Quality assessment of the resulting profiles showed median model length 110 match states (IQR: 78–189) and median effective number of sequences (Neff) 0.47 (IQR: 0.42–0.57); 100% of profiles have Neff *<* 2, as expected for *de novo* profiles built from tight 30% identity clusters within a single metagenomic dataset (Pfam achieves higher Neff through iterative curation aggregating sequences across evolutionary time). Zero profiles fall below 30 aa length. Model length describes the aligned family profile and should not be interpreted as evidence of independently delimited domain boundaries.

##### Pfam-dark family search (Step 6)

The concatenated family-profile HMM database was searched against the full algal proteome (all 2,044 samples, including both dark and annotated proteins) using `hmmsearch`. The primary family threshold was *E <* 10^−9^, yielding 30,994 prevalent Pfam-dark families and 557,892 family–environment correlations. A matched-design comparison (same 1,523 samples, same 18 GEE+WOA variables, same prevalence threshold) against the permissive Pfam-A v37.2 matrix (*E <* 10^−5^) yielded a Pfam-dark-family median *|ρ|* = 0.157 versus Pfam-domain median 0.068 (2.29-fold enrichment; bootstrap 95% CI [2.28, 2.30]; Mann-Whitney *p <* 10^−300^). Sensitivity analysis at two additional family-profile thresholds, *E <* 10^−7^ and *E <* 10^−5^, was performed by post-filtering the hmmsearch results and recomputing all environmental correlations; at *E <* 10^−5^, 31,257 prevalent Pfam-dark families yielded median *|ρ|* = 0.162 and 2.32-fold enrichment, confirming that the enrichment is robust across family-profile search thresholds.

##### Count matrix and characterization (Steps 6b–7)

Per-sample hmmsearch tblout files were parsed into a 2,044 *×* 33,950 sample *×* Pfam-dark-family count matrix matching the format of the existing Pfam count matrix. Families were characterized by cluster size distribution, sample prevalence, geographic breadth (unique TARA stations), and the fraction of cluster members also lacking DIAMOND nr hits (*>*90% nr-dark classifies a family as “nr-dark” versus “Pfam-only-dark”).

##### Environmental correlation (Step 8)

Pfam-dark-family abundances were CLR-transformed (pseudocount = 1.0) and tested for Spearman correlation against AEF embedding dimensions and GEE environmental variables, with Benjamini-Hochberg FDR correction (*q <* 0.05), matching the statistical framework applied to Pfam domains. At the primary family-profile threshold of *E <* 10^−9^, this yielded 557,892 family–environment association tests (30,994 prevalent families), of which 410,030 (73.5%) were significant at FDR *<* 0.05. Gradient boosting regressors (sklearn `GradientBoostingRegressor`; 100 trees, max depth 4, 5-fold CV) compared predictive performance using Pfam-only, Pfam-dark-family-only, and combined feature sets.

The full pipeline is orchestrated by a Slurm dependency-chained submission script (submit_pipeline.sh) that chains all steps via --dependency=afterok. All pipeline scripts, parameters, and provenance documentation are available in the code repository under 08_novel_domains/.

#### Methodological comparison with the Pfam-based analysis

**Methodological comparison with the Pfam-based analysis.**
| Aspect | Main Pfam pipeline | Pfam-dark family pipeline |
| --- | --- | --- |
| Annotation source | Pfam-A (curated) | *De novo* HMM profiles |
| Annotation E-value | $10^{-9}$ | $10^{-9}$ (primary); $10^{-5}$ (sensitivity) |
| CLR pseudocount | +1.0 | +1.0 |
| Correlation test | Spearman rank | Spearman rank |
| FDR method | Benjamini-Hochberg | Benjamini-Hochberg |
| FWER permutation | 1,000 iterations | Not implemented (exploratory) |
| Prevalence filter | $\geq 5\%$ of samples | $\geq 10$ samples |
| Environmental variables | AEF (64 dim.) for correlation screen; GEE + WOA23 (37 vars) for CCA and bidirectional modeling | AEF + GEE merged |
| Predictive model | Neural network | Gradient boosting |

#### InterProScan cluster cross-checks

For the 33,950-HMM global catalogue, representative sequences from the 100 domains ranked highest by prevalence, environmental coupling, and total abundance were searched against InterPro using the European Bioinformatics Institute (EBI) InterProScan representational state transfer (REST) API (Table S4). In a separate photosynthetic-anchor-neighbor analysis, 58,570 unannotated proteins adjacent to photosynthetic anchor genes were clustered independently with MMseqs2, yielding 35,667 representatives; this count is unrelated to the *≥*50-member filter that defined the 33,950 global-catalogue HMMs. All 35,667 representatives from the independent screen were searched against the full InterPro consortium member databases using InterProScan 5.74-105.0^86^ (36-chunk SLURM array job, 1,000 sequences per chunk). Analyses included Pfam, PANTHER, Gene3D, SUPERFAMILY, SMART, CDD, ProSiteProfiles, ProSitePatterns, PRINTS, NCBIfam, Hamap, PIRSF, SFLD, FunFam, MobiDBLite, Phobius, TMHMM, SignalP, Coils, and AntiFam. GO term mappings (`--goterms`) and pathway annotations (`--pathways`) were enabled. Input sequences were cleaned of stop-codon characters prior to submission. Chunk results were concatenated into a single merged output with a domain_id column linking annotations back to cluster representatives.

#### Structure prediction and homology searches

Structure prediction used a two-tier approach. First, 3,600 representative sequences spanning the full range of environmental coupling strengths (10 rank bins, 360 per bin, stratified by maximum *|ρ|*) were predicted using Boltz-2 v2.2.1^43^ (MIT-licensed) and AlphaFold 3 (below), providing the primary structural characterization.

Second, as an initial homology screen, the ten highest-ranked Pfam-dark families (by composite ranking of prevalence, *|ρ|*, and abundance; 943–1,468 assemblies each; 303–1,297 aa) were predicted using the ESMFold v1 API^8^ (api.esmatlas.com) and searched against PDB and AlphaFold Database via Foldseek. For sequences exceeding the 400-residue API limit (8 of 10), the first 400 residues were submitted; this truncation may omit C-terminal regions and limits the scope of negative homology results. NOVEL_33041 was not among the locally retained queries for this screen, so a PDB homology result is not available for that family.

For Boltz-2, multiple sequence alignments were constructed from MMseqs2 30% identity cluster members (median 66 per cluster) aligned with MAFFT, then projected onto the query sequence. Predictions completed in 25.5 h with 0 failures (Figure S7, Data S10).

As an independent structural assessment, the same 3,600 rank-binned sequences were predicted using AlphaFold 3 v3.0.1^44^ in template-free mode. Pre-built MMseqs2 30%-identity cluster MSAs (median 66 sequences per alignment) were supplied via the unpairedMsaPath field (AF3 v2 JSON dialect) with --run_data_pipeline=false to bypass jackhmmer/template search, ensuring that the MSA source matched the Boltz-2 predictions. Inference used 4 recycles, 1 model seed, and 5 diffusion samples per seed (AF3 defaults). Predictions completed in *∼*16 h wall time (median *∼*1 min per job; Figure S7, Data S10).

Predicted structures (PDB format) were searched against the AlphaFold Database (Swiss-Prot subset) and the Protein Data Bank using Foldseek^45^ (`foldseek search`, *E <* 10^−3^, --alignment-type 2), which encodes backbone geometry as a structural alphabet for rapid 3D-structure comparison. Alignment TM-scores^107^ were extracted to assess structural similarity: TM-score *>* 0.5 indicates the same fold, while *<* 0.3 suggests no structural relationship.

#### Cross-taxonomic selection analysis

Published per-gene selection metrics were used to compare genes without a Pfam or InterPro annotation (dark) with genes carrying at least one annotation (white) in four lineages (Data S9). For *Chlamydomonas reinhardtii*, per-gene *π*_N_ and *π*_S_ values from 20 strains^36^ were obtained from Dryad (doi:10.5061/dryad.1n0g6, SupplementalDS3) and mapped to NCBI assembly GCF_000002595.2. Genes with *π*_S_ = 0 or fewer than five single-nucleotide polymorphisms were excluded, leaving 4,085 dark and 7,038 white genes. Groups were compared by a two-sided Mann–Whitney *U* test; the median difference was bootstrapped 10,000 times (seed 42), and an SNP-count-matched analysis controlled for gene-length-related coverage. For *Seminavis robusta*, published *π*_N_*/π*_S_ values from 48 strains^37^ were partitioned by InterPro status. For *Synechococcus* CC9311 and CC9902, published per-gene *d*_N_*/d*_S_ values^38^ were partitioned by Pfam-A `hmmsearch` hits (*E <* 10^−9^). For *Thalassiosira pseudonana*, published Phylogenetic Analysis by Maximum Likelihood (PAML) site-model results from seven strains^39^ were partitioned by the same Pfam criterion, and proportions of positively selected genes were compared by Fisher’s exact test. Per-gene inputs, partitions, scripts, and full results are provided in Data S9 and Tables S11–S12.

#### Within-genome NOVEL_33041 analysis

The 66 NOVEL_33041 copies in the *Dunaliella* ROIL assembly were aligned at the protein level with MAFFT and back-translated to a 270-codon alignment. Pairwise *d*_N_*/d*_S_ was calculated for all 2,145 sequence pairs by the Nei–Gojobori method with Jukes–Cantor correction. A maximum-likelihood tree was inferred with IQ-TREE 2 using the Q.mammal+G4 model and 1,000 ultrafast bootstrap replicates. Copy locations were summarized by contig gene count, and expression evidence, metagenome detections, similarity searches, and localization heuristics were evaluated as detailed in the Supplemental Text. Summary statistics are in Table S13; source paths are listed in the Supplemental Text.

#### Dark-proteome physicochemical analysis

To compare the biochemical properties of Pfam-annotated and Pfam-dark proteins, 500,000 ORFs were randomly subsampled from each class (seed = 42) using `seqtk sample`. Thirteen per-ORF physicochemical metrics were computed. Isoelectric point (pI) and molecular weight were calculated with BioPython^90^ `ProteinAnalysis`. GRAVY (grand average of hydropathy) scores used the Kyte–Doolittle scale.^108^ Intrinsic disorder propensity was estimated as the fraction of disorder-promoting residues ({A, R, G, S, Q, P, E, K}) following Dunker et al.,^109^ with order propensity defined symmetrically ({W, C, F, I, Y, V, L, N}); the remaining four residues (D, H, M, T) are classified as neutral. Shannon entropy of the per-sequence amino acid frequency vector quantified compositional complexity (*H* = *−* ^∑^ *p_i_* log_2_ *p_i_* over the 20 standard residues). Low-complexity regions (LCRs) were identified by sliding a 12-residue window across each ORF and flagging positions where local Shannon entropy fell below 1.5 bits; LCR fraction is the proportion of residues covered by at least one such window (computed for the first 100,000 ORFs per class). Net charge per residue (NCPR; *f_K_* + *f_R_ − f_D_ − f_E_*) and fraction of charged residues (FCR; *f_K_* + *f_R_* + *f_D_* + *f_E_*) followed the Das–Pappu framework.^110^ A codon-usage proxy for genomic GC content was computed as the difference in fractional abundance of GC-rich codon amino acids ({A, G, P, R}) minus AT-rich codon amino acids ({F, Y, I, N, K}). Secondary structure propensity was estimated from mean Chou–Fasman^111^ *α*-helix and *β*-strand parameters per ORF. All sequences were filtered to the 20 standard amino acids; ORFs shorter than 5 residues were excluded. The bulk algal proteome was omitted from comparisons because *∼*87% of it consists of dark proteins, making it a near-duplicate of the dark class (Figure 6).

#### Protein language model learnability analysis

Five GPT-2 Small models^40^ were trained from scratch on five protein-sequence corpora under identical hyperparameters using nanoGPT.^41^ Training code and analysis results are available at Zenodo^63^ (Data S11); model checkpoints^62^ and source sequences^82^ are available on Hugging Face.

##### Datasets

Three biological corpora were constructed from the TARA-Oceans algal proteome: annotated photosynthetic proteins (“white”; 66.5M training tokens, 7.4M validation tokens), a 400,000-sequence subsample of the algal proteome (“algae”; 58.9M/6.5M tokens), and Pfam-dark proteins (“dark”; 63.1M/7.0M tokens). Two random controls were generated: composition-matched random sequences preserving the empirical algal unigram amino-acid frequency distribution (“random algae”; 55.4M/6.2M tokens), and uniform independent and identically distributed (i.i.d.) sequences drawn from 20 amino acids with equal probability 1*/*20 (“random full”; 55.4M/6.2M tokens). All corpora used a character-level tokeniser with no sub-word segmentation. Biological datasets use vocabulary size 23 (20 standard amino acids, ambiguity symbol X, separator >, newline); random corpora omit X (vocabulary 22). Sequences were stored in flat-text format (one per line), tokenised as uint16 arrays, and split 90/10 by character position for training/validation.

##### Architecture and training

All five runs used the GPT-2 Small architecture: 12 transformer layers, 12 attention heads, 768-dimensional embeddings, 1,024-token context window, 0.2 dropout, no bias terms (*∼*85M parameters). Optimiser: AdamW (*β*_1_ = 0.9, *β*_2_ = 0.99). Learning rate: cosine decay from 5 *×* 10^−4^ to 5 *×* 10^−5^ over 40,000 iterations with 200-iteration linear warm-up. Effective batch size: 12 *×* 2 gradient-accumulation steps *×* 1,024 tokens *≈* 24,576 tokens per update. Validation loss was evaluated every 1,000 iterations (40 evaluation points); the checkpoint with the lowest validation loss was saved. Training was logged via Weights & Biases.

##### Cross-entropy interpretation and bounds

The model minimises mean negative log-likelihood (cross-entropy) in nats; a cross-entropy of *L* nats corresponds to perplexity *e^L^*. The theoretical entropy floor for the uniform random corpus is *H* = ln(20) *≈* 2.996 nats; the trained model reached 2.993 nats, confirming correct implementation. The composition-matched random floor is lower than ln(20) due to non-uniform residue probabilities; the trained model at 2.855 nats is consistent with learning unigram frequencies only. Any loss reduction below the composition-matched floor reflects higher-order statistical regularities in biological sequences.

##### Receiver operating characteristic (ROC) analysis

For each of the five checkpoints, per-sequence mean negative log-likelihood was computed on 1,000 held-out validation sequences per corpus (balanced random sample, sequence lengths 30–512 tokens). Tokens were remapped through a shared character alphabet to correct for vocabulary-ID differences between biological (vocabulary 23) and random (vocabulary 22) tokenisers. The per-sequence score 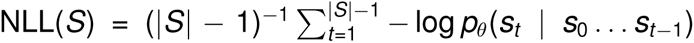 served as the discriminant (lower NLL = model prefers the sequence). Area under the ROC curve (AUROC) values and 95% bootstrap confidence intervals (5,000 resamples) were computed for all pairwise binary classification tasks.

##### Statistical comparisons

Pairwise validation-loss comparisons used the Mann-Whitney *U* test (one-tailed) on the last 10 evaluation steps (stable convergence window). Cohen’s *d* was computed on the same window. Final validation-loss differences are reported with 95% percentile bootstrap confidence intervals (9,999 resamples). Learnability scores are defined as *L*(*D*) = 100 *×* (*L*_floor_ *− L_D_*)*/*(*L*_floor_ *− L*_best_), where *L*_floor_ is the random full validation loss and *L*_best_ is the white validation loss.

#### Validation and sensitivity analyses

##### Temporal holdout

We linked 973 TARA and OSD assemblies to collection dates through the PANGAEA station registry (doi:10.1594/PANGAEA.842237) and OSD metadata. For the primary temporal test, XGBoost models were trained on 2009–2010 TARA samples (*n* = 354) and tested without overlap on 2011–2012 samples (*n* = 417); a secondary split trained on 2010–2011 and tested on 2009 plus 2012. Pfam abundances were CLR-transformed (pseudocount = 1) and reduced to 100 principal components. Models used the spatial-CV hyperparameters (max depth = 6, learning rate = 0.1, 200 estimators, subsample = 0.8, column sample by tree = 0.8). Because the TARA cruise trajectory links sampling year to ocean region, these holdouts test temporal persistence and geographic extrapolation jointly.

##### Geographic and dataset sensitivity

Spatial-block sensitivity used 2°, 5°, and 10° grid cells for SST prediction (*n* = 1,279). Leave-one-basin-out CV trained on six of seven named basins and tested on the omitted basin in turn, using CLR-transformed Pfam features (pseudocount = 0.5) and fixed XGBoost settings (200 estimators, max depth = 4, learning rate = 0.05); performance was summarized both by equally weighted basin scores and pooled held-out predictions. Data-source sensitivity repeated SST prediction and CCA using TARA metagenomes alone (*n* = 773 and 772, respectively). Size-fraction sensitivity restricted the analysis to 150 TARA 20–180 µm assemblies (90 spatial blocks); 8,378 domains meeting 5% prevalence were CLR-transformed (pseudocount = 0.5), and the same 10-fold spatial-block design was applied. Extended results are reported in Tables S6–S7 and Supplemental Text.

##### Lineage and prevalence controls

The lineage-controlled Pfam-dark-family-versus-Pfam-domain comparison used the 1,133-sample intersection with both count matrices, 18 environmental variables, and all 15 RuBisCO lineage counts. Counts were CLR-transformed (pseudo-count = 1), and features present in fewer than 10 samples were excluded. Partial Spearman correlations were computed by rank-transforming variables and using QR-decomposition projection to remove the 15 continuous lineage covariates. *|ρ|* distributions were compared by Mann–Whitney *U* test with a bootstrap 95% confidence interval for the enrichment ratio. A separate prevalence control matched Pfam-dark families and Pfam domains within 20 prevalence bins before comparison.

##### Variance partitioning

Distance-based redundancy analysis (db-RDA)^112^ used the 786 samples with complete environmental and geographic metadata. Euclidean distance on CLR-transformed Pfam abundances, equivalent to Aitchison distance, was projected onto 174 principal-coordinate axes retaining 80.0% of positive-eigenvalue variance. Predictors comprised 24 standardized GEE and WOA23 environmental variables and a geographic matrix of latitude, longitude, and 18 distance-based Moran eigenvector maps.^113^ Adjusted *R*^2^ was partitioned into pure environmental, shared spatial–environmental, pure geographic, and residual fractions. Conditional Freedman–Lane tests with 999 permutations assessed the pure environmental and pure geographic fractions; an unconditional row-permutation test assessed the full model.

### Statistical analysis

All statistical analyses were performed in Python 3.10.12 using scipy 1.9.3,^95^ statsmodels 0.13.5,^98^ pandas 1.5.3,^97^ numpy 1.24.3,^96^ xgboost 1.7.4,^22^ shap 0.41.0,^23^ scikit-learn 1.2.2,^94^ umap-learn 0.5.3,^28^ and PyTorch 2.1.^92^ GPT-2 protein language models were trained using nanoGPT^41^ with PyTorch.

Unless another unit is stated, *n* denotes independent biological records (metagenome assemblies, transcriptomes, or reference proteomes), not proteins, domains, tests, permutations, or cross-validation folds. The broad input collection included all 2,357 available records from the specified sources. The domain-analysis set required a completed algaGPT classification and a non-empty Pfam profile (*n* = 2,044). Geographic analyses further required valid coordinates and, where applicable, a named basin; AEF analyses used the 995 samples with AEF records, including 26 with partial zero filling during feature preparation. Target-specific models used complete cases for the relevant response and predictors, so their sample sizes are reported with each analysis. Spearman calculations omitted missing pairs using `nan_policy=’omit’`. No sample was excluded on the basis of an observed association or model outcome; the 68-point IQR filter in Figure 2A was used only for visualization and not for inferential analyses.

Spearman correlations tested 761,472 domain–dimension pairs (11,898 Pfams *×* 64 dimensions; *n* = 995 samples with AEF records), with Benjamini–Hochberg FDR controlled at 5%. Rank-based tests did not assume normality. We did not use the parametric Wilks’ Λ approximation for CCA inference because multivariate normality was not expected; significance was evaluated by row and spatial-block permutation. Predictive performance was evaluated out of sample by 5-fold CV for the embedding screen and 10-fold spatial-block CV for the primary bidirectional models. Model residual normality was not required for the tree-based regressors. Permutation tests assumed exchangeability at their stated unit; spatial-block permutations preserved within-block autocorrelation.

Centers and dispersion are identified with each result: skewed distributions are summarized by the median and interquartile range, approximately symmetric replicate or fold distributions by the mean and standard deviation, and resampling uncertainty by percentile 95% confidence intervals. Primary bidirectional-model uncertainty is the cross-fold standard deviation from 10-fold spatial-block CV; Figure 3E additionally reports 95% bootstrap confidence intervals from 1,000 resamples of held-out 80/20-split predictions. Exact sample sizes, test direction, correction family, and permutation or bootstrap counts are stated in the corresponding methods, legends, or tables.

This observational computational study had no treatment allocation or experimental blinding; randomization and investigator blinding were therefore not applicable. Random seeds were fixed where stochastic fitting or subsampling was used (seed = 42 unless otherwise stated), and permutation labels or spatial blocks were reassigned computationally as described above. We analyzed the full available dataset after prespecified technical and metadata-completeness filters. No a priori power calculation was performed because the study was exploratory and sample sizes were determined by public-data and metadata availability.

All computation was performed on the Jubail high-performance computing (HPC) cluster at New York University Abu Dhabi unless otherwise noted. GPU tasks used NVIDIA A100 GPUs (40 GB) managed by SLURM. At the reported throughput of *∼*280 sequences/s, LA^4^SR inference across 447.7 million sequences corresponds to *∼*444 serial processing hours; retained local records do not establish the concurrent GPU count or elapsed wall time. Neural network training (environment-to-Pfam projection, protein language models) required *∼*6 h on A100 GPUs. Boltz-2 structure prediction (3,600 sequences) required 25.5 h on an NVIDIA GB10 GPU. AlphaFold 3 structure prediction (3,600 sequences) required *∼*16 h on 2*×* NVIDIA RTX A6000 GPUs (48 GB each; A^2^S^2^ Lab, NYU Abu Dhabi). InterProScan annotation of the 35,667 representatives from the independent photosynthetic-anchor-neighbor screen was parallelised as a 36-chunk SLURM array job. Downstream statistical analyses (correlation, XGBoost, CCA, dimensionality reduction) required *∼*12 h of additional central-processing-unit (CPU) or GPU time.

### Additional resources

Analysis code (Code S1):^59^ https://doi.org/10.5281/zenodo.22910378

Trained algaGPT classifier:^60^ https://huggingface.co/GreenGenomicsLab/algaGPT

Bidirectional XGBoost models:^61^ https://huggingface.co/GreenGenomicsLab/TARA-XGBoost-Bidirectional

VICReg joint embedding model:^47^ https://huggingface.co/GreenGenomicsLab/TARA-WorldModel-VICReg

Dark-whiteGPLM checkpoints:^62^ https://huggingface.co/SarahDaakour/dark-whiteGPLM

Dark-whiteGPLM training data:^82^ https://huggingface.co/datasets/SarahDaakour/dark-whiteGPLM-data

Dark-whiteGPLM code:^114^ https://github.com/SarahD4/dark-whiteGPLM

TARA Oceans data portal:^3,57^ https://www.ebi.ac.uk/metagenomics/

MMETSP transcriptomes:^18,58^ https://www.ncbi.nlm.nih.gov/bioproject/PRJNA231566

Pfam database:^12,68^ https://ftp.ebi.ac.uk/pub/databases/Pfam/releases/Pfam37.2/

AlphaEarth embeddings:^13,69^ https://developers.google.com/earth-engine

## SUPPLEMENTAL INFORMATION

### SUPPLEMENTAL FIGURES

**Figure S1.**
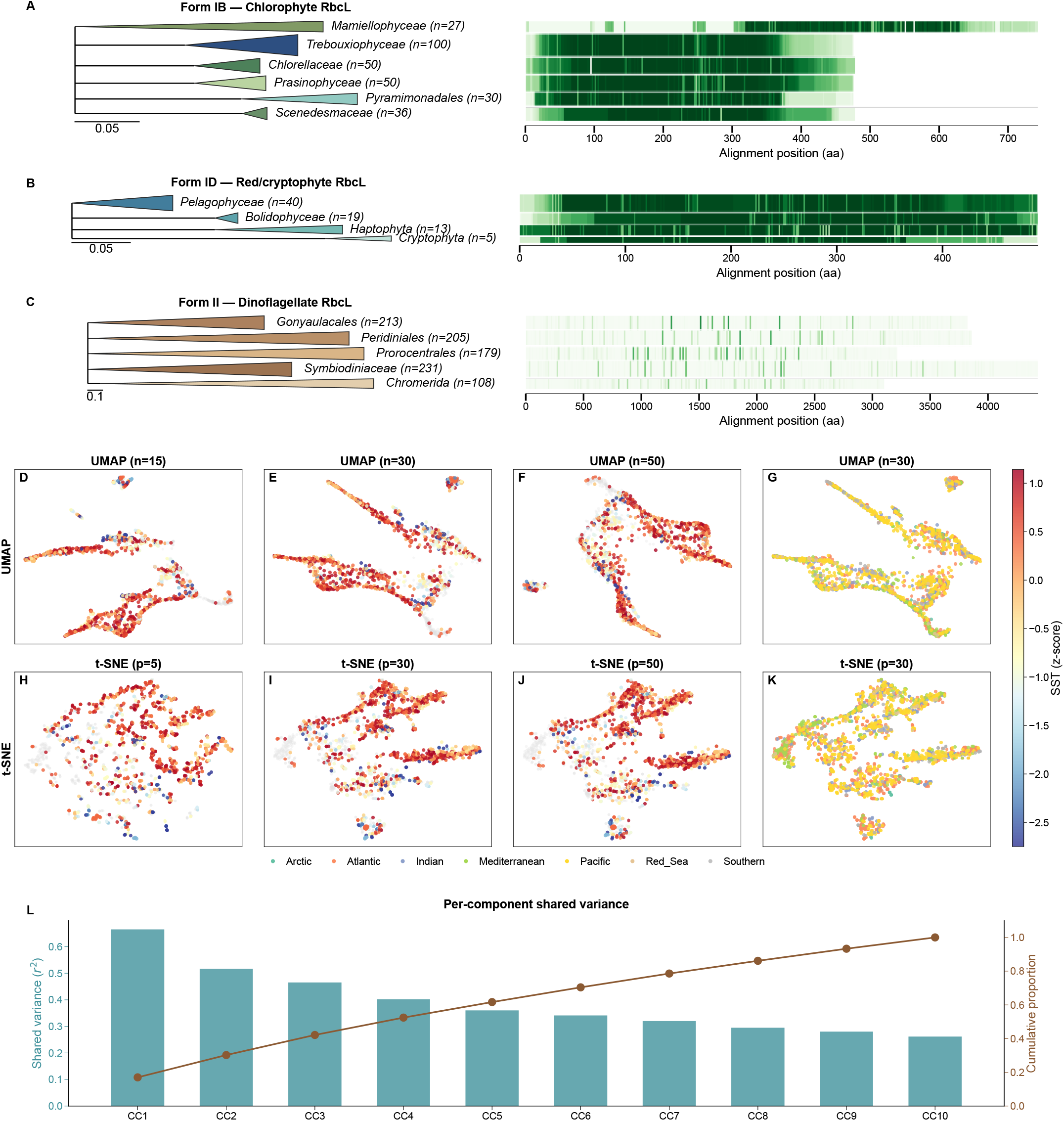
RuBisCO RbcL phylogeny (A–C), dimensionality-reduction parameter sensitivity (D–K), and canonical shared-variance decomposition (L), related to Figures 1 and 2. (A–C) Collapsed phylogenetic trees for 15 ribulose-1,5-bisphosphate carboxylase/oxygenase (RuBisCO) lineages, with conservation calculated within each lineage’s multiple-sequence alignment. Form IB contains six chlorophyte lineages (293 reference sequences; alignment lengths 475–743 amino acids [aa]), Form ID contains four red-lineage taxa (Pelagophyceae, Bolidophyceae, Haptophyta, and Cryptophyta; 77 sequences; 488–490 aa), and Form II contains five myzozoan lineages, four dinoflagellate lineages (Symbiodiniaceae, Peridiniales, Gonyaulacales, and Prorocentrales) plus Chromerida (936 sequences; 3,096–4,431 aa). Form IB and ID trees were inferred with IQ-TREE 2, ModelFinder, and 1,000 ultrafast bootstrap replicates; the Form II tree used FastTree with WAG+CAT and six *Rhodospirillum rubrum* Protein Data Bank structures as the omitted outgroup. Values beside lineage names give sequence counts. (D–F) Uniform Manifold Approximation and Projection (UMAP) of 1,810 samples represented by 20,318 raw Pfam count columns (unstandardized) from the permissive Pfam-A v37.2 screen (*E <* 10*^−^*^5^) at 15, 30, and 50 neighbors, colored by mean sea surface temperature (SST). (G) UMAP at 30 neighbors colored by ocean basin. (H–J) t-distributed Stochastic Neighbor Embedding (t-SNE) at perplexities 5, 30, and 50, colored by SST. (K) t-SNE at perplexity 30 colored by basin. (L) Per-component squared canonical correlation and cumulative fraction of shared variance across ten environment–Pfam canonical correlation analysis (CCA) components; CC1 accounts for 17% of the total.

**Figure S2.**
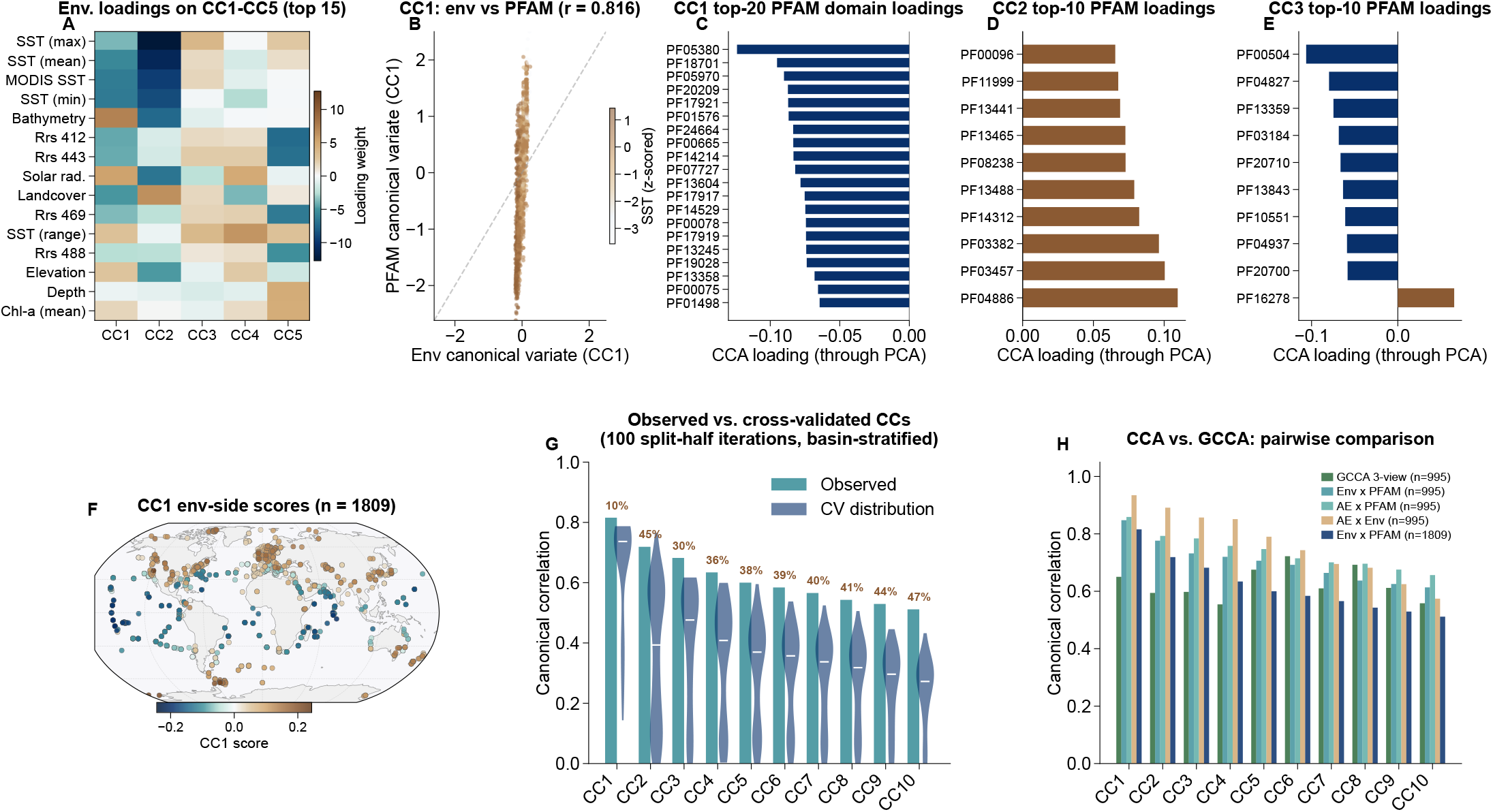
Comprehensive canonical correlation analysis (CCA), related to Figure 2. Panels A–G show the two-view raw-environment–Pfam CCA (*n* = 1,809). (A) Fifteen largest absolute environmental loadings on CC1–CC5; sea surface temperature (SST) dominates CC1–CC2. (B) Environment-versus Pfam-side CC1 scores (*r* = 0.816), colored by SST. (C– E) Highest-loading Pfam domains for CC1–CC3; domain loadings are back-projected through the 100-component principal component analysis (PCA). (F) Global distribution of environment-side CC1 scores. (G) Full-data canonical correlations versus held-out split-half correlations across 100 basin-stratified iterations; violins show the iteration distributions and markers show medians. CC1 shrinks by 9.7%; CC6–CC10 are exploratory. (H) Canonical correlations from three-view generalized CCA (GCCA; AlphaEarth Foundations [AEF] + raw environment + Pfam, *n* = 995) and four two-view CCAs, including the full environment–Pfam analysis (*n* = 1,809) and matched-sample analyses (*n* = 995).

**Figure S3.**
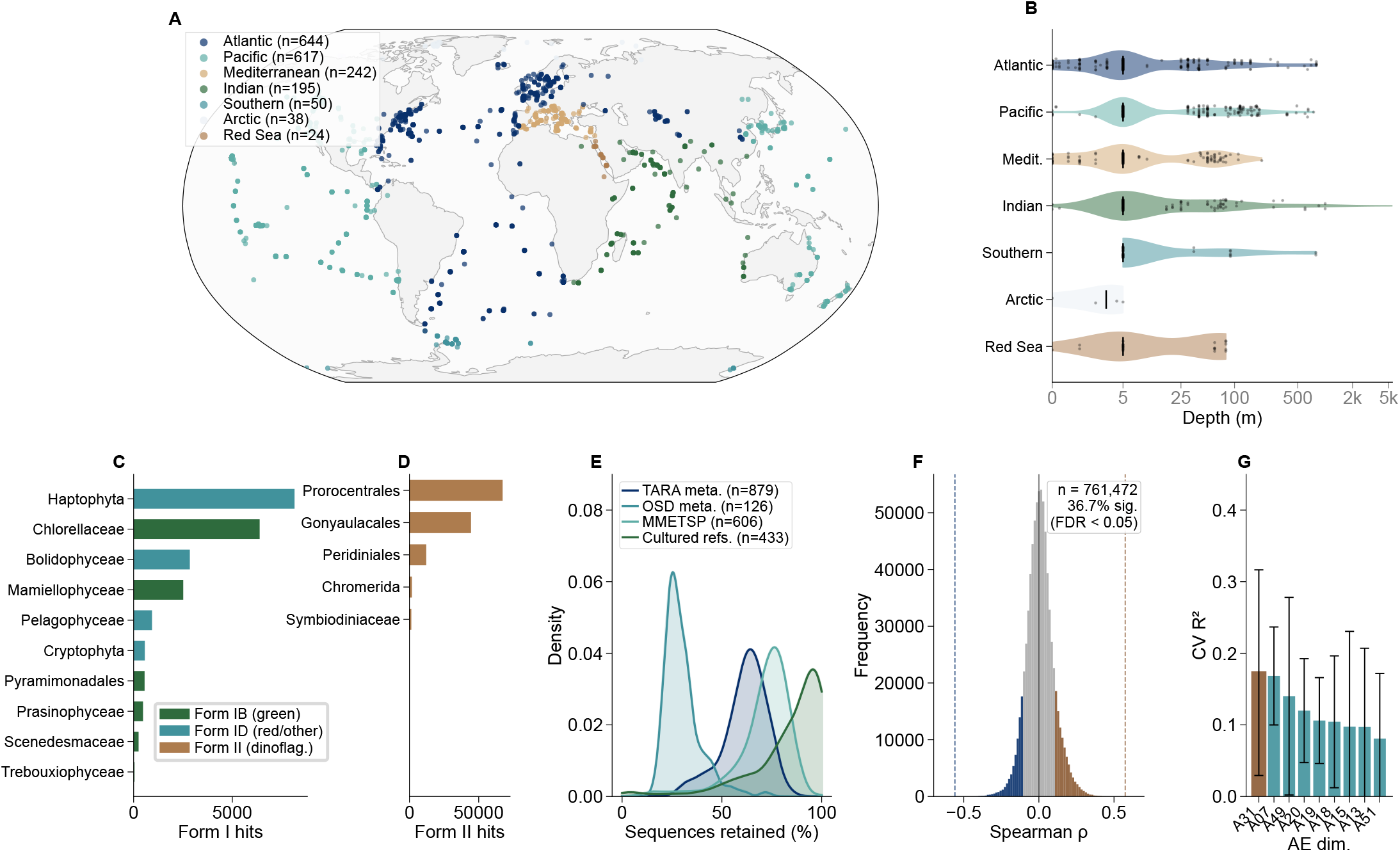
Basin coverage, RuBisCO lineage composition, and domain–environment correlation, related to Figures 1 and 2. (A) Global distribution of 1,810 geolocated samples across seven basins; labels give sample counts. (B) Log-scaled sampling-depth distributions by basin; violins show sample distributions and internal boxes show medians and interquartile ranges. (C–D) Form I (22,743 hits) and Form II (126,990 hits) RuBisCO lineage composition, totaling 149,733 hits across 15 lineages; panels use independent count axes. (E) algaGPT retention percentage by source type (TARA metagenomes, *n* = 879; Ocean Sampling Day [OSD], *n* = 126; Marine Microbial Eukaryote Transcriptome Sequencing Project [MMETSP], *n* = 606; cultured references, *n* = 433). (F) Distribution of 761,472 Spearman correlations between 11,898 Pfam domains and 64 AlphaEarth Foundations dimensions (*n* = 995); 36.7% pass Benjamini–Hochberg false discovery rate (FDR) *<* 0.05. The solid line marks zero and dashed lines mark the observed minimum and maximum *ρ*. (G) Mean *±* standard deviation five-fold CV *R*^2^ for the nine best-predicted AlphaEarth dimensions using domain composition (*n* = 995).

**Figure S4.**
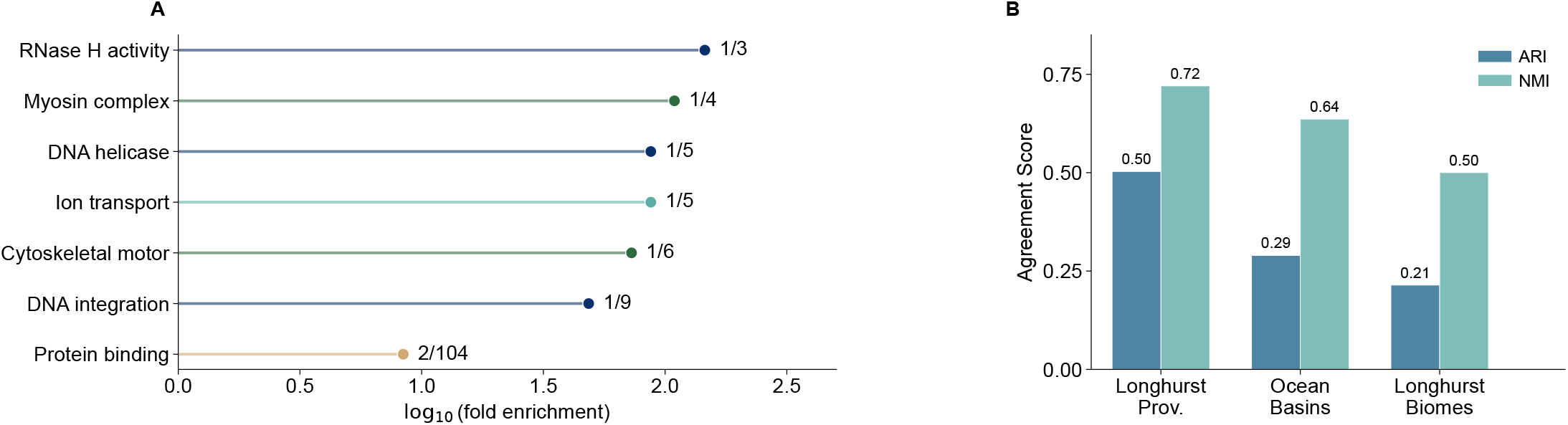
Gene Ontology (GO) enrichment and biogeographic agreement, related to Figures 3 and 4. (A) Hypergeometric GO enrichment among the 22 Pfam domains with held-out test *R*^2^ *>* 0.3 in the top-100 single-split (80/20) forward screen (background: 9,611 Pfam domains; only GO terms carried by at least three background domains were tested). Seven terms pass Benjamini–Hochberg false discovery rate (FDR) *q <* 0.05; the x-axis shows log_10_ fold enrichment. Labels give *k/N*, the number of test-set domains carrying the term over the number of background domains carrying it; six of the seven terms rest on a single domain (*k* = 1). (B) Adjusted Rand index (ARI) and normalized mutual information (NMI) between Hierarchical Density-Based Spatial Clustering of Applications with Noise (HDBSCAN) functional biomes and three geographic schemes: Longhurst provinces, ocean basins, and Longhurst biomes (*n* = 1,310 non-noise samples of the 1,810 GPS-mapped samples from all source types; 500 HDBSCAN noise samples excluded).

**Figure S5.**
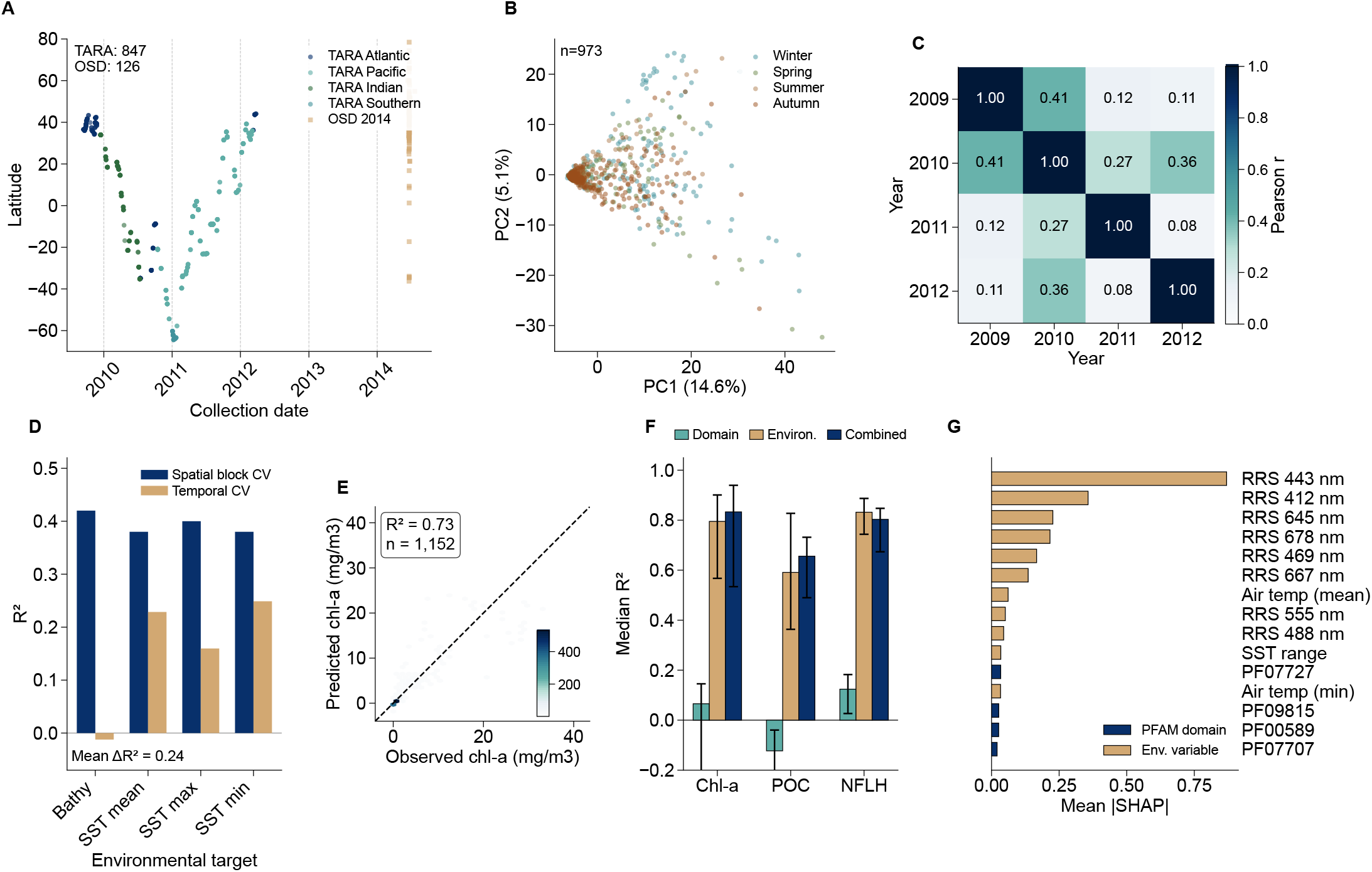
Temporal analysis of domain–environment coupling (A–D) and productivity proof-of-concept under spatial block cross-validation (CV; E–G), related to Figure 2 and Discussion. (A) Sampling timeline and geographic coverage for 847 TARA assemblies collected in 2009–2012 and 126 Ocean Sampling Day (OSD) assemblies collected in 2014; colors indicate basin. (B) Principal component analysis (PCA) of the 500 most variable Pfam domains across 973 dated samples, colored by season (permutational multivariate analysis of variance *R*^2^ = 0.025). (C) Pairwise Pearson correlations between annual vectors of per-domain environmental association coefficients; values are shown in each cell. (D) Reverse-model *R*^2^ under 10-fold spatial-block CV and temporal holdout, in which models are trained on earlier collection years and tested on a later year, for four environmental targets; the in-panel annotation gives the mean spatial-minus-temporal *R*^2^ difference across the four targets. (E) Observed versus predicted chlorophyll-a for the combined domain-plus-environment XGBoost model, pooling held-out predictions from 10 spatial folds (*n* = 1,152); hexagons encode sample density and the dashed line is identity. (F) Median fold *R*^2^ with interquartile range across the same 10 spatial folds for domain-only, environment-only, and combined models of chlorophyll-a and particulate organic carbon (POC; *n* = 1,152 each) and normalized fluorescence line height (NFLH; *n* = 1,151). (G) Top 15 mean absolute SHapley Additive exPlanations (SHAP) feature values for the combined chlorophyll-a model, colored as environmental or Pfam-domain features.

**Figure S6.**
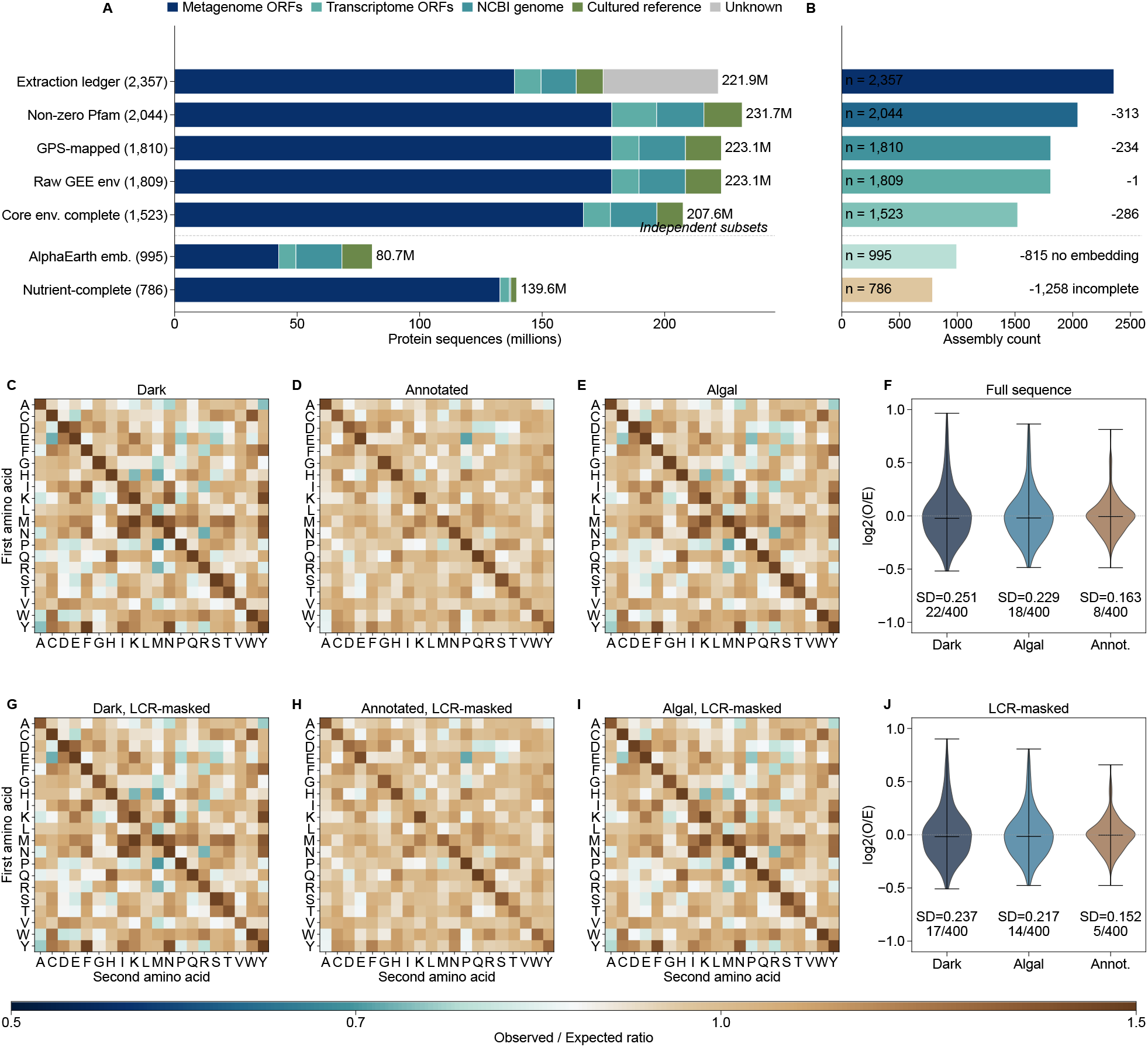
Sample accounting through the analysis pipeline (A–B) and dipeptide observed/expected (O/E) analysis (C–J), related to Methods, Table S1, and Figure 6. (A) Algal protein counts by source and analysis inventory. The 221.9-million early extraction ledger and 231.7-million source-resolved domain-analysis inventory are distinct inventories, not consecutive attrition stages; subsequent subsets derive from the 2,044-sample domain-analysis set. (B) Assembly counts, inclusion criteria, and exclusions at each stage. The linear stages are nested (2,357 *→* 2,044 *→* 1,810 *→* 1,809 *→* 1,523); the AlphaEarth (995) and nutrient-complete GEE + WOA23 (786) subsets branch independently, with exclusion counts computed relative to the 1,810 GPS-mapped samples (815 without AEF records) and the 2,044-sample domain-analysis set (1,258 without the complete 24-variable matrix), respectively. (C–E) Dipeptide O/E-ratio heatmaps for all 400 dipeptides in the dark, annotated, and algal proteomes (500,000 sequences per class; seed = 42). (F) Distributions of full-sequence log_2_(O*/*E) across the 400 dipeptides. Standard deviations are 0.251 (dark), 0.229 (algal), and 0.163 (annotated); respectively 22, 18, and 8 dipeptides have *|* log_2_(O*/*E)*| >* 0.5, and the dark-to-annotated SD ratio is 1.54. (G–I) O/E-ratio heatmaps after masking low-complexity regions (LCRs; 12-residue window, Shannon entropy *<* 1.5 bits). (J) LCR-masked log_2_(O*/*E) distributions. Standard deviations are 0.237 (dark), 0.217 (algal), and 0.152 (annotated); biased-dipeptide counts are 17, 14, and 5, and the dark-to-annotated SD ratio remains 1.56.

**Figure S7.**
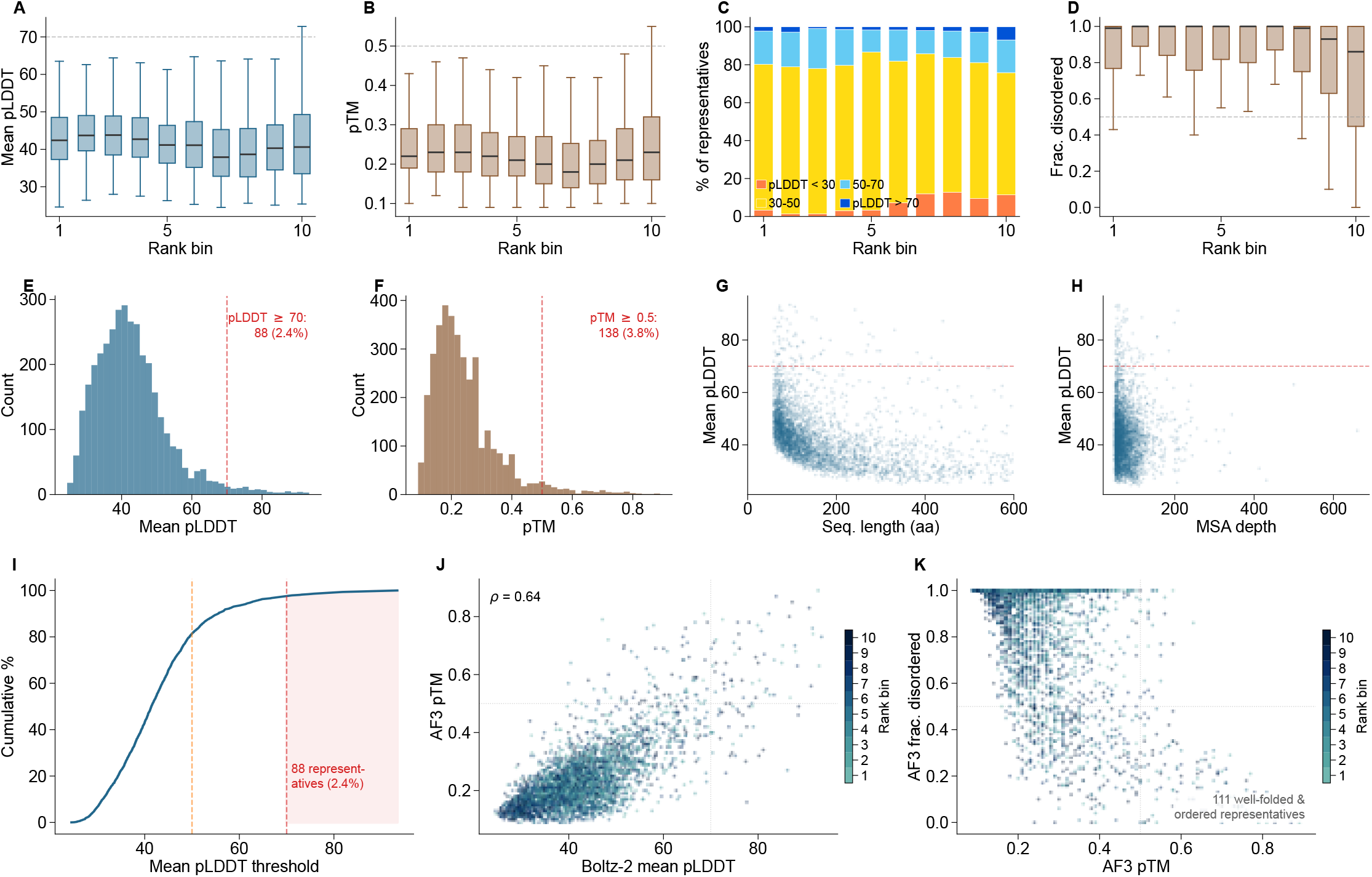
Structure-prediction confidence analysis of 3,600 rank-binned Pfam-dark protein-family representatives using Boltz-2 and AlphaFold 3, related to Figure 7. **(A)** Boltz-2 mean predicted local distance difference test (pLDDT) by environmental coupling rank bin (360 representatives per decile; bin 1 = strongest coupling). Dashed line: pLDDT 70 confident-fold threshold. **(B)** AlphaFold 3 (AF3) predicted template modeling score (pTM) by rank bin. Dashed line: pTM 0.5 threshold. In A, B, and D, boxes show medians and interquartile ranges, whiskers extend to 1.5 times the interquartile range, and outliers are omitted from display. **(C)** Boltz-2 confidence category breakdown per bin (pLDDT thresholds: 30, 50, 70). **(D)** AF3 fraction disordered by rank bin. Dashed line: 0.5 threshold. **(E)** Distribution of Boltz-2 mean pLDDT across all 3,600 representatives. Dashed line: pLDDT 70; 88 representatives (2.4%) had mean pLDDT *≥* 70. **(F)** Distribution of AF3 pTM; 138 representatives (3.8%) exceed pTM 0.5. **(G)** Boltz-2 mean pLDDT versus sequence length. Dashed line: pLDDT 70. **(H)** Boltz-2 mean pLDDT versus multiple-sequence-alignment (MSA) depth (MMseqs2 cluster size). Dashed line: pLDDT 70. **(I)** Cumulative distribution of Boltz-2 mean pLDDT. Dashed lines mark pLDDT 50 and 70; the shaded region marks the 88 representatives with mean pLDDT *≥* 70. **(J)** Boltz-2 mean pLDDT versus AF3 pTM, colored by rank bin (1–10; Spearman *ρ* = 0.64). **(K)** AF3 pTM versus AF3 fraction disordered, colored by rank bin (1–10). Lower-right quadrant: 111 well-folded and ordered representatives (pTM *≥* 0.5, fraction disordered *<* 0.5).

### SUPPLEMENTAL TABLES

**Table S1.**
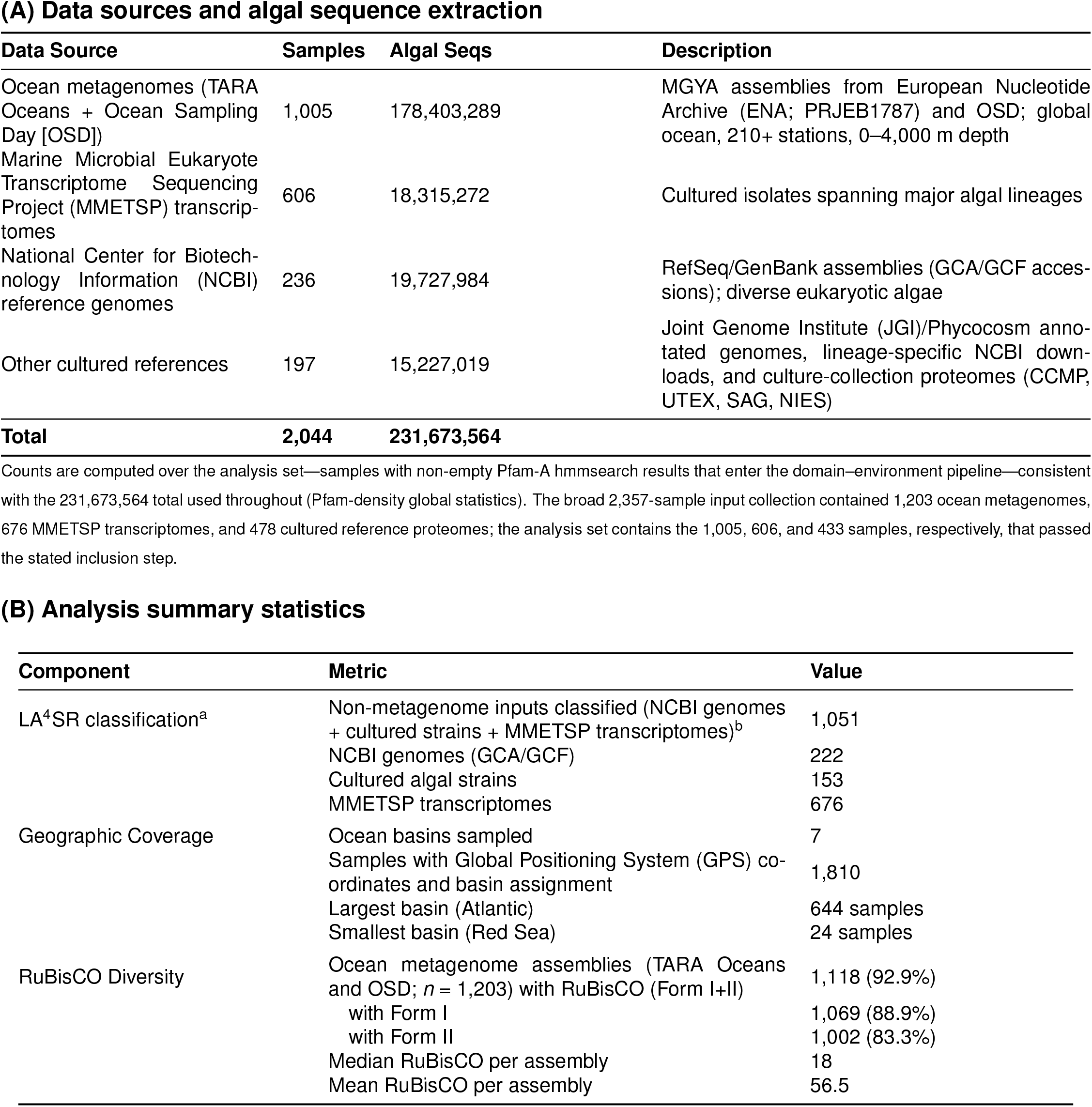
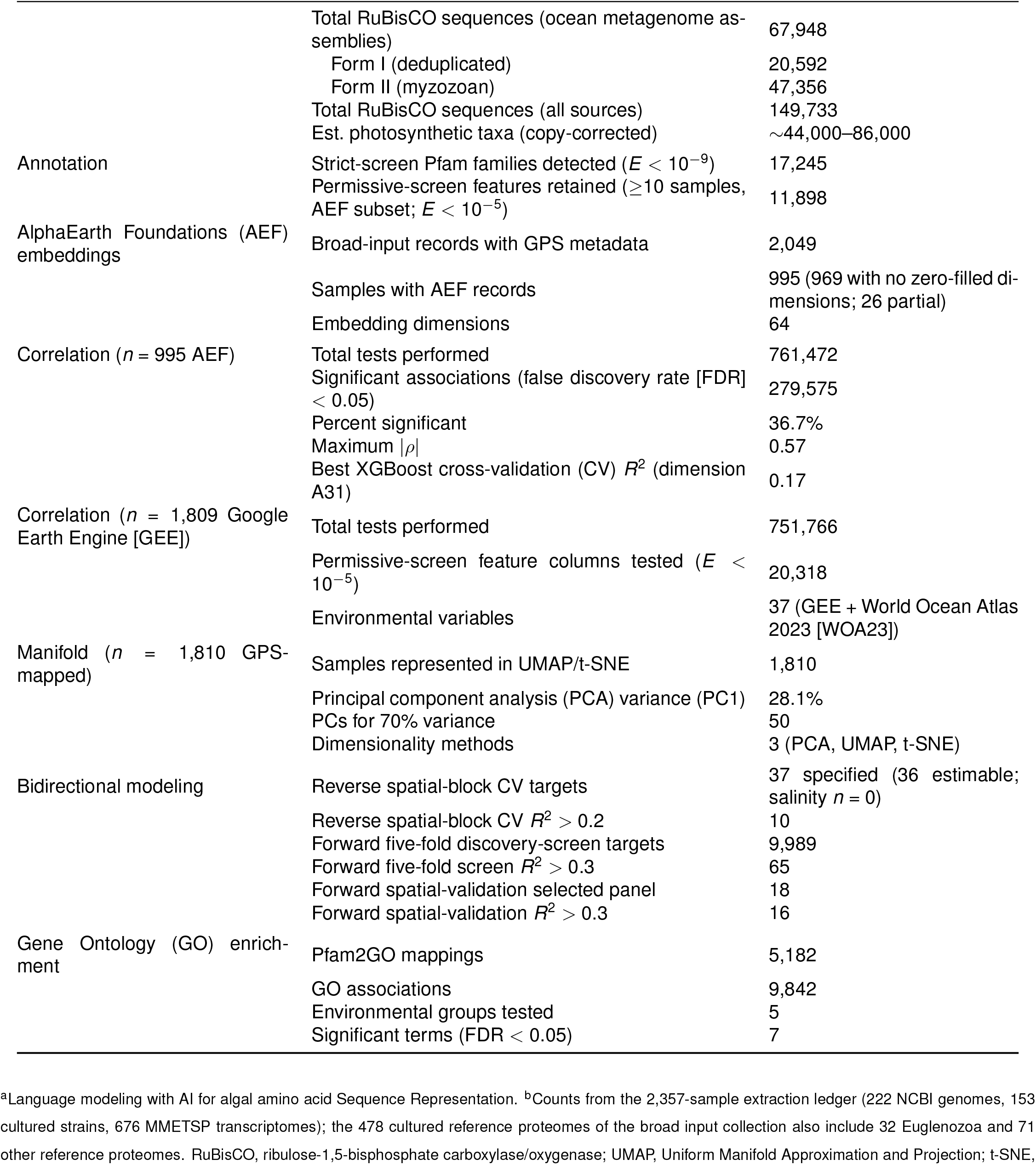
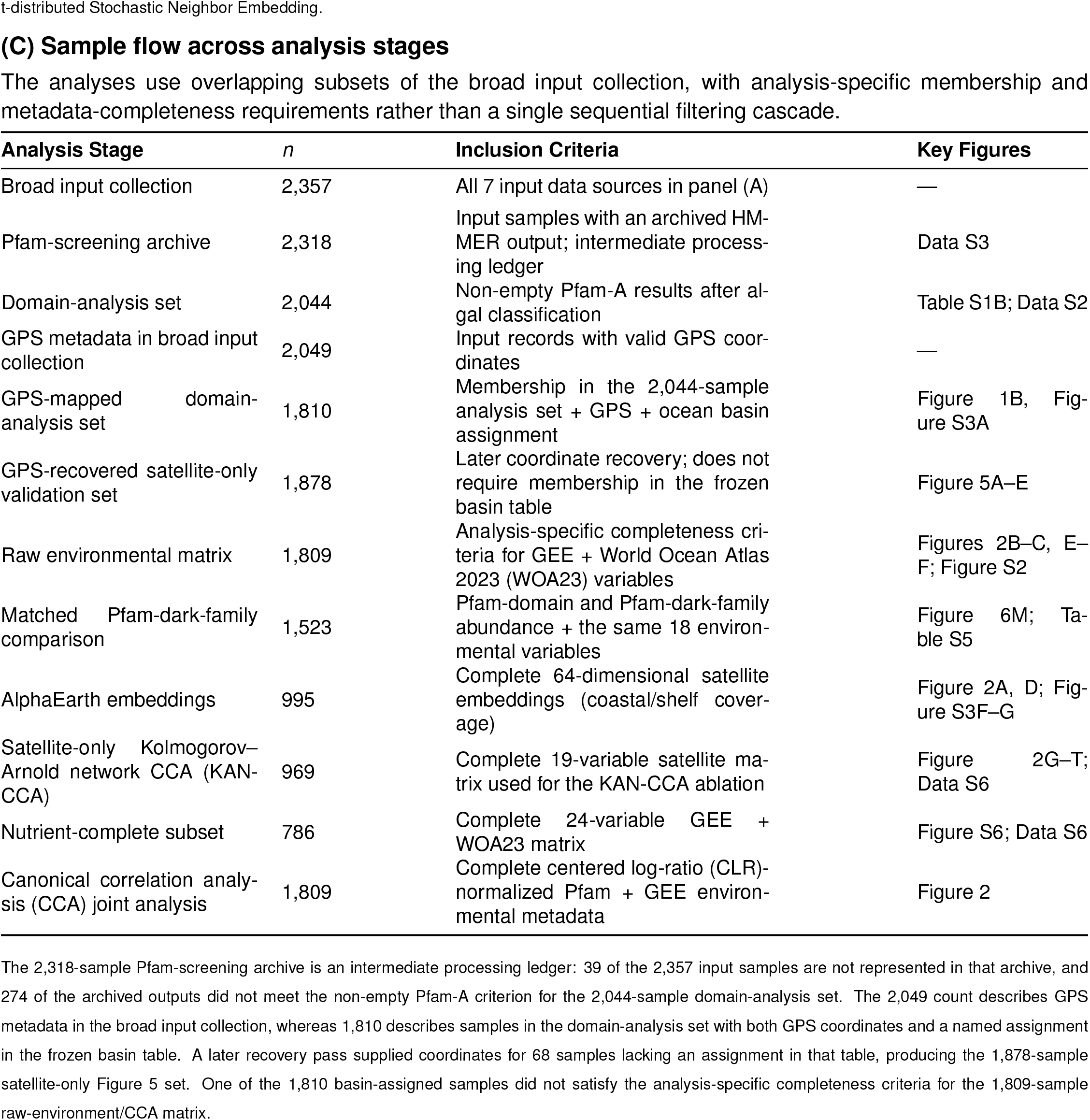
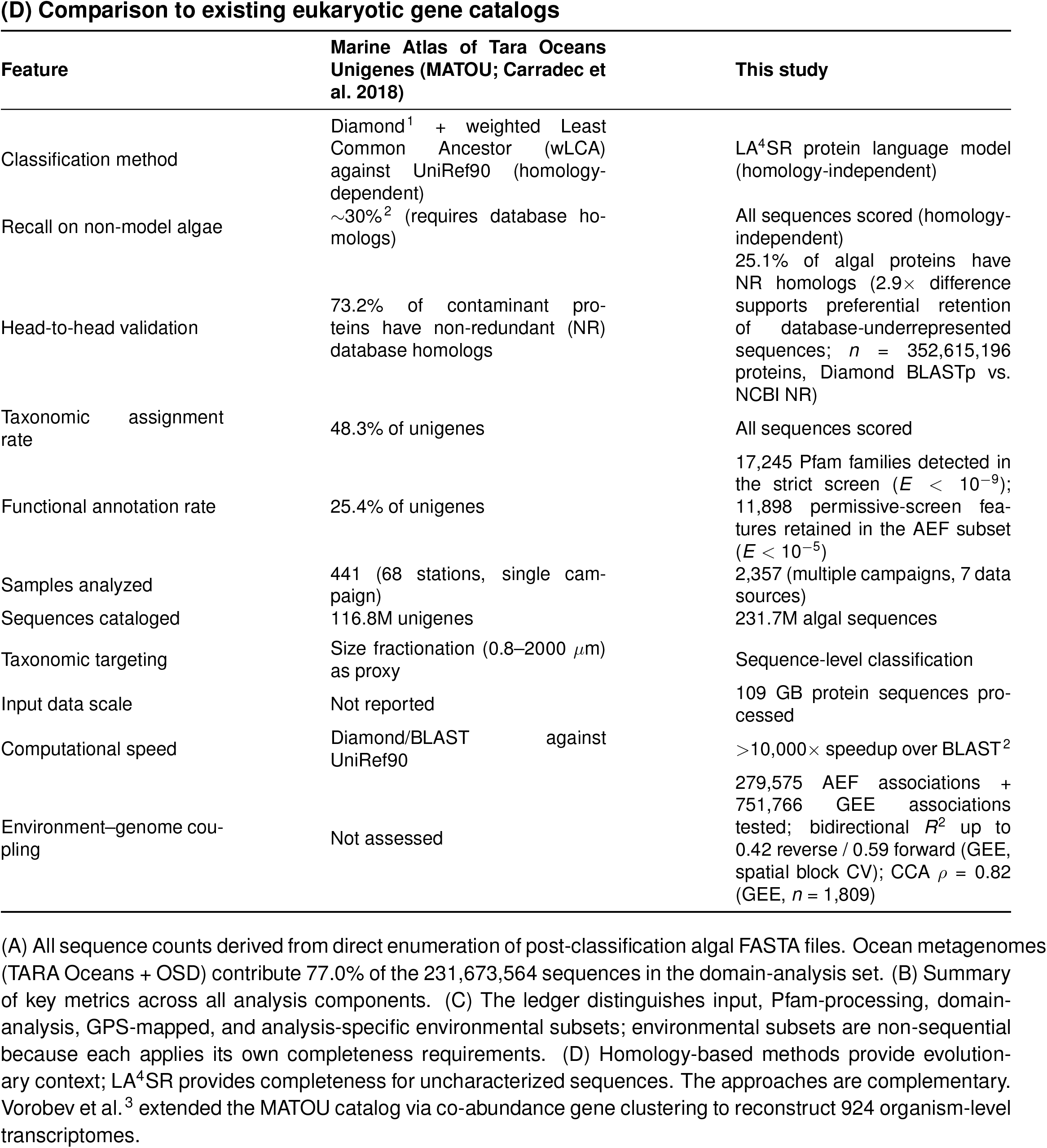
Full dataset and analysis summary, related to Figures 1–5 and Methods.

**Table S2.**
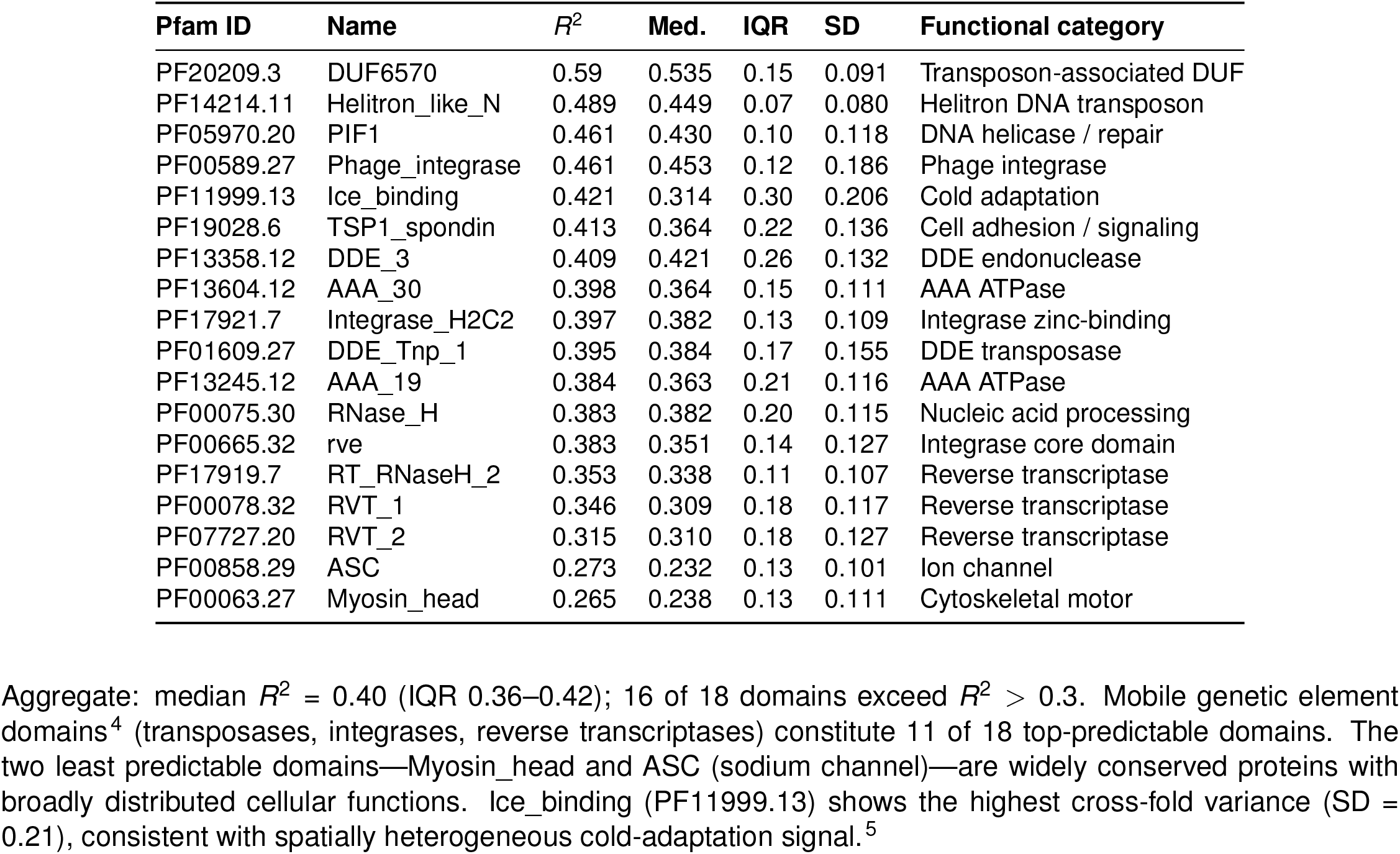
Forward model individual domain performance under spatial block CV, related to Figure 5. Per-domain prediction accuracy for 18 Pfam domains selected in an earlier standard five-fold screen and then evaluated under 10-fold spatial block cross-validation (2° grid cells; *n* = 1,810 samples). Because selection and spatial validation use overlapping samples, these values condition on screen selection and are not selection-independent estimates; the maximum may be optimistic. Domains are ranked by overall *R*^2^. Functional category was assigned from Pfam clan membership and InterPro annotations. Median *R*^2^, interquartile range (IQR), and standard deviation (SD) were computed across 10 spatial folds.

**Table S3. Combined Form I and Form II RuBisCO per-sample detection across a separately enumerated 2,372-sample full-protein marker-search inventory (15 lineages), related to Figure 1**

Per-sample RuBisCO sequence counts across 15 algal and myzozoan lineages for all 2,372 samples in the full-protein marker-search inventory (1,203 ocean metagenome assemblies [TARA Oceans and OSD], 677 MMETSP transcriptomes, 222 reference genomes, and 270 cultured strains). This inventory was enumerated independently for the RuBisCO analysis. The 15 records absent from the classifier input collection had sequences that were too short to align or came from non-target species. Columns report total predicted proteins, total RuBisCO sequences, Form I green-lineage (IB) and red-lineage (ID) subtotals, Form II (myzozoan) subtotal, and per-lineage counts for 10 Form I lineages (Mamiellophyceae, Prasinophyceae, Pyramimonadales, Chlorellaceae, Trebouxiophyceae, Scenedesmaceae, Pelagophyceae, Bolidophyceae, Haptophyta, Cryptophyta) and 5 Form II lineages (Symbiodiniaceae, Peridiniales, Gonyaulacales, Prorocentrales, Chromerida). Form I RuBisCO is chloroplast-encoded and typically single-copy per photosynthetic genome;^6^ Form II is nuclear-encoded in myzozoans and may be multi-copy in some dinoflagellate species.^7,8^

**Summary statistics:** 2,237 of 2,372 samples (94.3%) contained at least one RuBisCO sequence. Total: 149,733 unique sequences (Form I: 22,743; Form II: 126,990). Among the 1,203 ocean metagenome assemblies (TARA Oceans and OSD): 1,118 (92.9%) detected RuBisCO (Form I in 1,069 [88.9%], Form II in 1,002 [83.3%]), with a median of 18 distinct sequences per assembly (mean 56.5, range 0–721) spanning a median of 6 co-occurring photosynthetic lineages (mean 6.2).

*Full table (2,372 rows × 21 columns) provided as a machine-readable TSV in Data S4*.

**Table S4.** InterPro cross-check of top 100 Pfam-dark protein families against current InterPro member databases, related to Figure 6. Pfam-dark protein families were defined by whole-sequence MMseqs2 clustering of proteins with no Pfam-A v37.2 match at the strict threshold of *E <* 10^−9^. The clustering required 80% bidirectional coverage and did not delimit independent domains. To test whether these families would be reclassified by an integrated database, a different tool, and a different query sequence, representatives from the 100 most prevalent and environmentally coupled families were searched against the full InterPro database (which includes Pfam among 20+ member databases) using the InterProScan representational state transfer application programming interface (REST API) provided by the European Bioinformatics Institute (EBI). InterProScan differs from the original pipeline in two material ways: (i) it applies member-database-specific search parameterization and gathering thresholds rather than the uniform *E <* 10^−9^ cutoff, and (ii) it searches the cluster representative rather than the individual member sequences used in the original `hmmsearch`. Any Pfam matches reported below therefore reflect these methodological differences, not a contradiction with the original dark-proteome classification. Families were ranked by a composite score combining prevalence (number of assemblies), environmental coupling (max *|ρ|*), and total abundance. **Summary of InterPro cross-check results.**

**Summary of InterPro cross-check results.**
| Category | Count | % |
| --- | --- | --- |
| Families submitted | 100 | 100 |
| No match from any database | 41 | 41 |
| Soft predictions only (disorder, coiled-coil, transmembrane [TM], signal peptide) | 24 | 24 |
| Functional database match (Pfam, PANTHER, CDD, Gene3D, etc.) | 30 | 30 |
| API errors/timeouts | 5 | 5 |
| **With Pfam match via InterProScan** | **5** | **5** |
| With integrated InterPro entry (IPR accession) | 14 | 14 |
| No functional annotation (no match + soft-only + errors) | 70 | 70 |

The five families that returned Pfam hits via InterProScan—but not in the original hmmsearch at *E <* 10^−9^—correspond to mobile genetic elements (retrotransposon polyproteins, 3/5) and conserved house-keeping proteins (viral DNA polymerase, histone; 2/5). The dominant functional database providing coverage beyond Pfam was PANTHER (28/100 families), which classifies proteins into broad evolutionary families. Overall, 95% returned no Pfam hit from InterProScan and 86% lacked any integrated InterPro entry, supporting their classification as largely uncharacterized protein families rather than universally database-absent sequences.

#### Independent photosynthetic-anchor-neighbor InterProScan screen (35,667 representatives)

This second screen is distinct from the global catalogue of 33,950 hidden Markov models (HMMs) assessed in the top-100 cross-check above. It began with 58,570 unannotated proteins neighboring photosynthetic anchor genes, which were clustered independently with MMseqs2 to yield 35,667 representatives; the 35,667 count is therefore not a pre-filter total for the 33,950 HMMs. All representatives from this independent screen were scanned against InterPro using InterProScan 5.74-105.0 in a 36-chunk array job. InterProScan uses current Pfam releases and its own gathering thresholds, so a small fraction of sequences classified as unannotated in the source screen may return Pfam hits under these different conditions. Results:

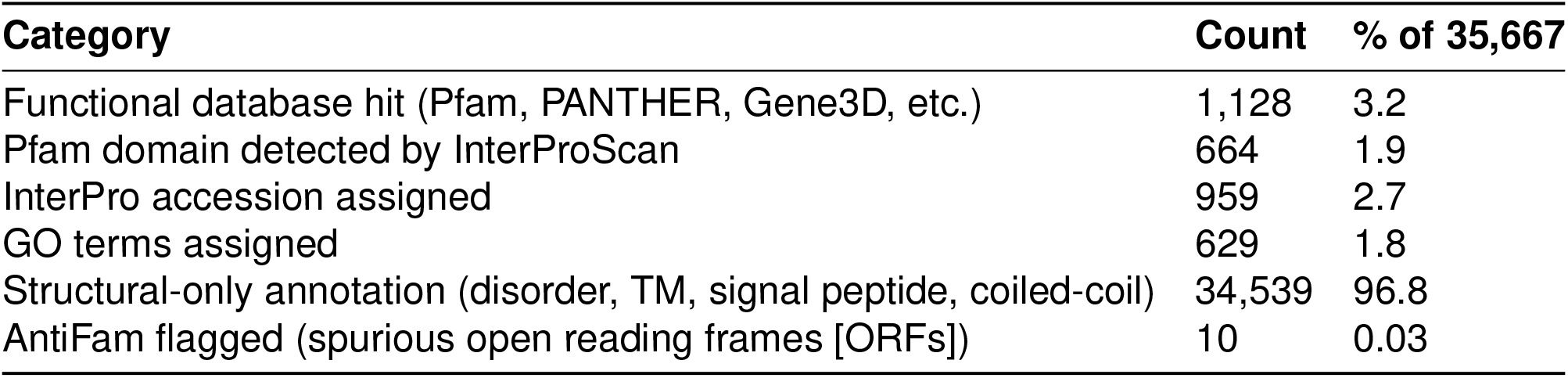

In this independent set, 98.1% of cluster representatives returned no Pfam hit under InterProScan’s updated database and search parameterization, 96.8% lacked any functional annotation from InterPro member databases, and AntiFam flagged only 10 of 35,667 sequences (0.03%) as potential spurious ORFs. The 664 sequences (1.9%) that did return Pfam hits are dominated by Chlorophyll A-B binding protein (PF00504; 64 representatives), Leucine Rich Repeats (PF00560/PF13516; 31), and WD repeats (PF00400; 14). These repeat and photosynthesis-associated domains lie near the boundary of detection sensitivity and were detected here using updated Pfam releases and InterProScan’s gathering thresholds. The low annotation rate independently supports extensive uncharacterized sequence space near photosynthetic genes, but it does not constitute a full-catalogue scan of the 33,950 HMMs. Structural annotations were common (MobiDBLite disorder: 12,047 sequences; Phobius signal/TM: 7,928; TMHMM: 1,296).

**Table S5.**
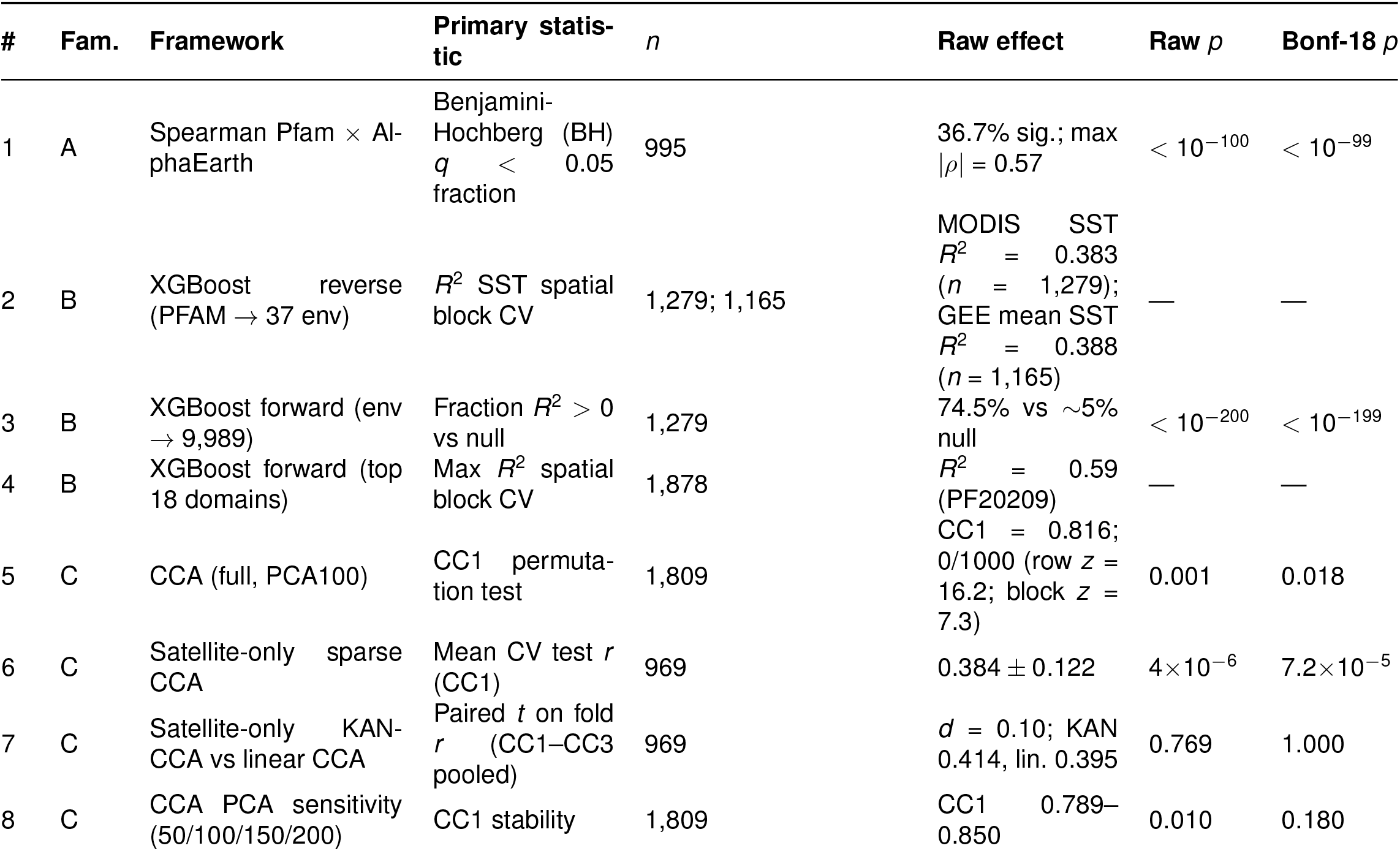
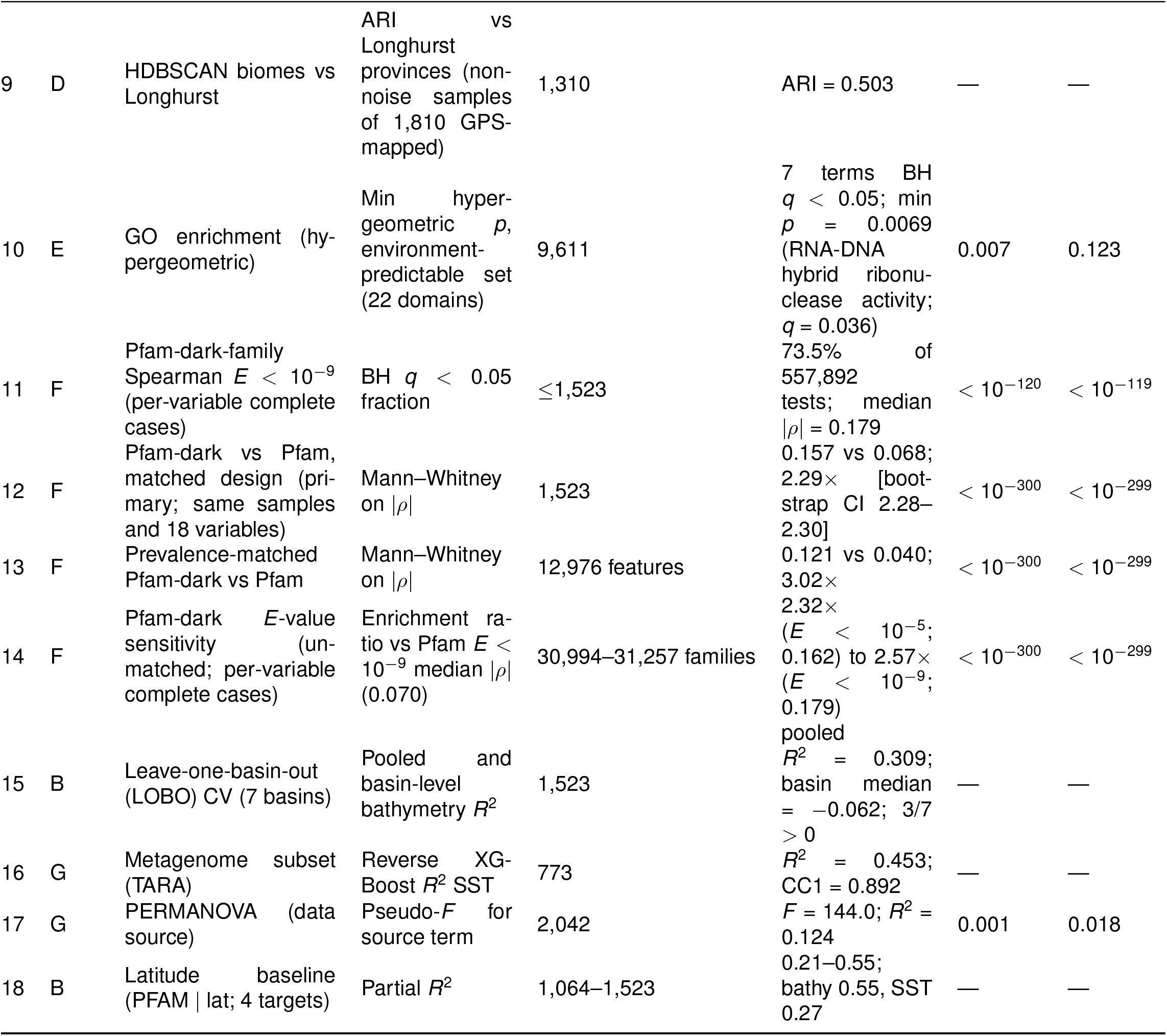
Framework-level multiple-testing table, related to Discussion. We enumerated the 18 analytical frameworks used in the manuscript and paired each with its primary effect estimate and sample count. For the 12 frameworks with a documented primary *p*- or *q*-value, we also report the raw value and a conservative Bonferroni-18 correction (raw value *×*18, capped at 1.0); 9 of these 12 retain corrected *p <* 0.05. The six cross-validation, agreement, or sensitivity frameworks without a separate null-hypothesis test (F2, F4, F9, F15, F16, and F18) are marked with dashes and excluded from the inferential count. All reported statistics are traceable to the source files listed in the machine-readable provenance table. Because several frameworks address variants of the same underlying claim, we group them into seven claim families (column “Family”): **A** Pfam–environment correlation discovery (F1); **B** bidirectional domain–environment prediction (F2–F4, F15, F18); **C** joint embedding structure (F5–F8); **D** biome partitioning (F9); **E** GO functional enrichment (F10); **F** Pfam-dark protein-family characterization (F11–F14); and **G** data-source sensitivity (F16– F17). These families organize convergent evidence and are not treated as additional hypothesis tests. HDB-SCAN denotes Hierarchical Density-Based Spatial Clustering of Applications with Noise; ARI, adjusted Rand index; PERMANOVA, permutational multivariate analysis of variance; SST, sea surface temperature; KAN-CCA, Kolmogorov–Arnold network canonical correlation analysis; CI, confidence interval.

Dashes indicate cross-validation, agreement, or sensitivity results for which no separate null-hypothesis test was performed. Raw *p*-values expressed as *<* 10^−^*^k^* correspond to framework-level effect sizes whose analytical minimum is far below conventional reporting thresholds (e.g., million-scale Mann– Whitney comparisons, 761,472 Spearman tests, or 557,892 BH-tested Pfam-dark-family correlations); these entries are capped at a reporting bound of 10^−100^ to 10^−300^ as documented in the machine-readable source table deposited with Data S1.

**Table S6.** Leave-one-basin-out cross-validation results, related to Results and Discussion. Leave-one-basin-out (LOBO) CV trains on 6 basins and tests on the held-out basin, cycling through all 7 basins (Arctic, Atlantic, Indian, Mediterranean, Pacific, Red Sea, Southern). LOBO XGBoost models used *n*_estimators_ = 200, max depth = 4, and learning rate = 0.05 (CLR-transformed Pfam features, pseudocount = 0.5; 20,318 domains); these settings differ from the 10-fold spatial block CV models (max depth = 6, learning rate = 0.1). The SST target is MODIS SST. Each basin is held out in turn; the model trains on the remaining 6 basins. Basins with *n*_test_ *<* 5 are excluded. We report both the median of the seven basin-level *R*^2^ values, which weights basins equally, and pooled *R*^2^ computed after concatenating all held-out predictions, which weights the represented samples and target variation. Prediction errors are summarized as mean absolute error (MAE).

| Target | Basin | $n_{\text{test}}$ | $R^2$ | MAE |
| --- | --- | --- | --- | --- |
| Bathymetry (m) | Arctic | 34 | -2.38 | 532 |
|  | Atlantic | 543 | 0.30 | 1,419 |
|  | Indian | 169 | 0.28 | 1,307 |
|  | Mediterranean | 131 | -1.88 | 1,442 |
|  | Pacific | 581 | 0.42 | 1,246 |
|  | Red Sea | 19 | -80.1 | 1,819 |
|  | Southern | 46 | -0.06 | 1,377 |
|  | *Median basin [IQR]* | 1,523 | -0.06 [-2.13-0.29] |  |
|  | *Pooled held-out predictions* | 1,523 | 0.31 | 1,326 |
| MODIS SST (°C) | Arctic | 30 | -47.6 | 10.4 |
|  | Atlantic | 428 | -0.0005 | 5.34 |
|  | Indian | 148 | -8.35 | 5.72 |
|  | Mediterranean | 127 | -8.13 | 2.95 |
|  | Pacific | 486 | 0.12 | 4.27 |
|  | Red Sea | 19 | -17.5 | 3.74 |
|  | Southern | 41 | -757 | 16.2 |
|  | *Median basin [IQR]* | 1,279 | -8.35 [-32.6--4.07] |  |
|  | *Pooled held-out predictions* | 1,279 | 0.24 | 5.18 |

**Table S7.**
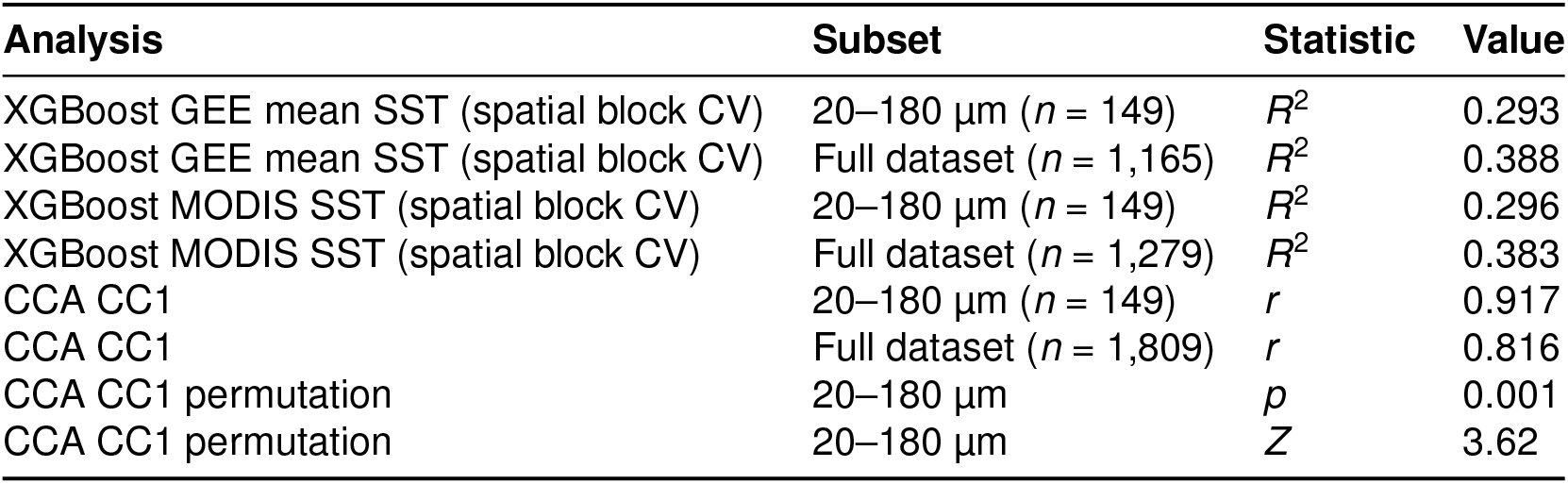
Size-fraction sensitivity: 20–180 µm TARA subset, related to Discussion. Size-fraction sensitivity analysis restricted to TARA metagenome assemblies from the 20–180 µm fraction (*n* = 150 assemblies; 90 spatial blocks on a 2° grid). Size-fraction assignments were derived from the MGnify assembly metadata mapping. XGBoost hyperparameters: *n*_estimators_ = 200, max depth = 6, learning rate = 0.1; CLR-transformed Pfam features (pseudocount = 0.5; 8,378 domains at 5% prevalence threshold). Spatial blocks: 2° grid, 10-fold CV. CCA: 20 PCA components from 100-component reduction (55.4% variance retained), 24 environmental variables. MODIS denotes the Moderate Resolution Imaging Spectroradiometer.

**Table S8.** Variance inflation factors for 23 environmental variables, related to Figure 2 and Methods. Variance inflation factors for 23 environmental variables (*n* = 1,101 complete-case samples). VIF = *∞* indicates exact linear dependence. Variables with VIF *>* 10 (collinearity concern) are marked with ^∗∗^.

| Variable | VIF | Max $\vert r\vert $ |
| --- | --- | --- |
| sst_max_c | $\infty^{**}$ | 0.940 |
| sst_min_c | $\infty^{**}$ | 0.947 |
| sst_range_c | $\infty^{**}$ | 0.539 |
| rrs_667 | 21,655.6** | 0.998 |
| rrs_678 | 21,650.8** | 0.998 |
| rrs_547 | 10,548.1** | 0.999 |
| rrs_555 | 5,654.6** | 0.999 |
| rrs_469 | 3,924.6** | 0.967 |
| rrs_531 | 2,267.6** | 0.990 |
| rrs_443 | 1,698.3** | 0.979 |
| rrs_488 | 1,349.6** | 0.948 |
| rrs_645 | 721.9** | 0.997 |
| sst_mean_c | 259.6** | 0.996 |
| modis_sst_mean_c | 211.3** | 0.996 |
| rrs_412 | 161.8** | 0.979 |
| nflh_mean | 121.4** | 0.680 |
| chl_mean_mg_m3 | 6.2 | 0.826 |
| poc_mean_mg_m3 | 4.3 | 0.826 |
| chl_max_mg_m3 | 3.1 | 0.609 |
| chl_min_mg_m3 | 2.2 | 0.527 |
| bathymetry_m | 2.0 | 0.414 |
| solar_rad_mj_m2 | 1.5 | 0.333 |
| distance_to_coast_km | 1.2 | 0.262 |

**Table S9.**
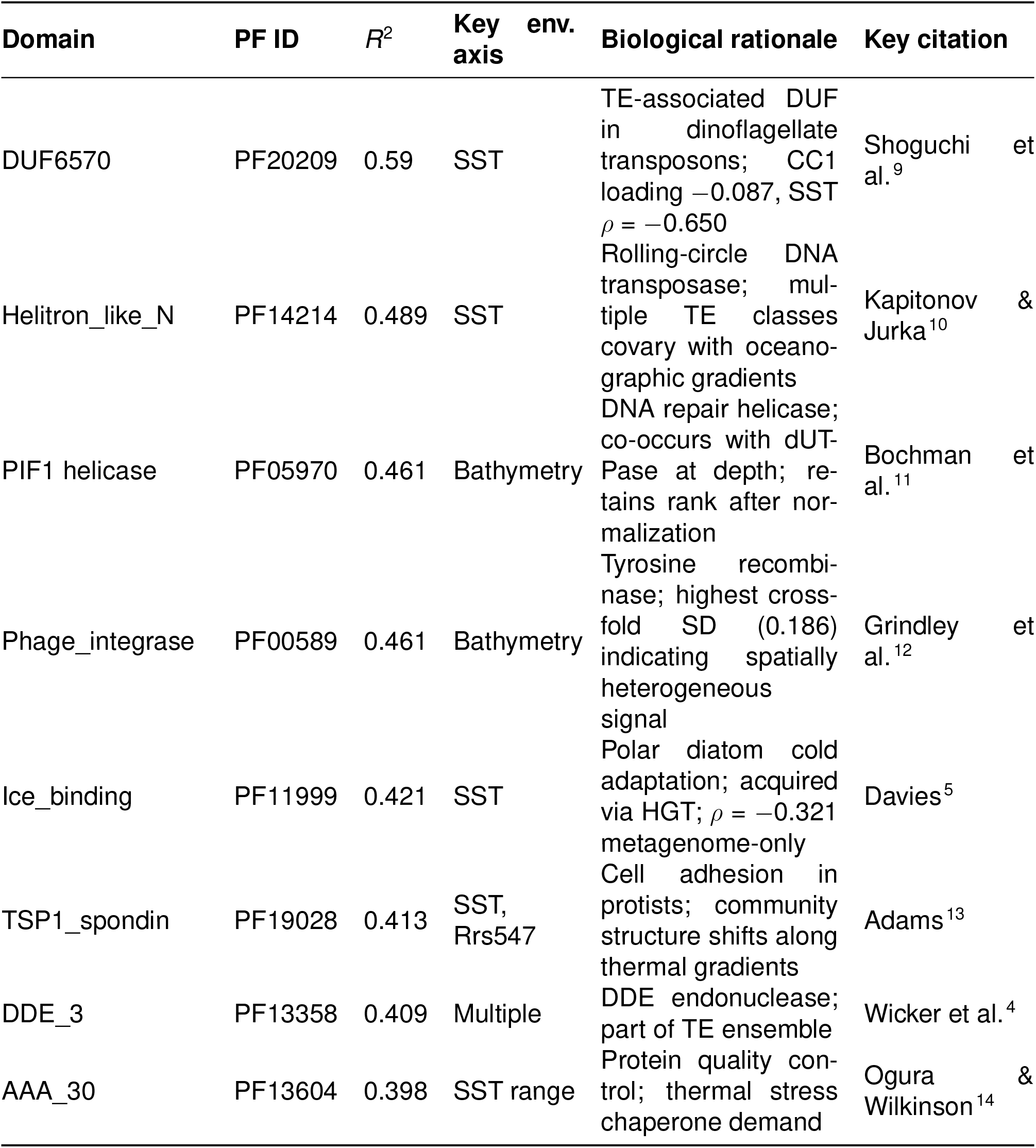
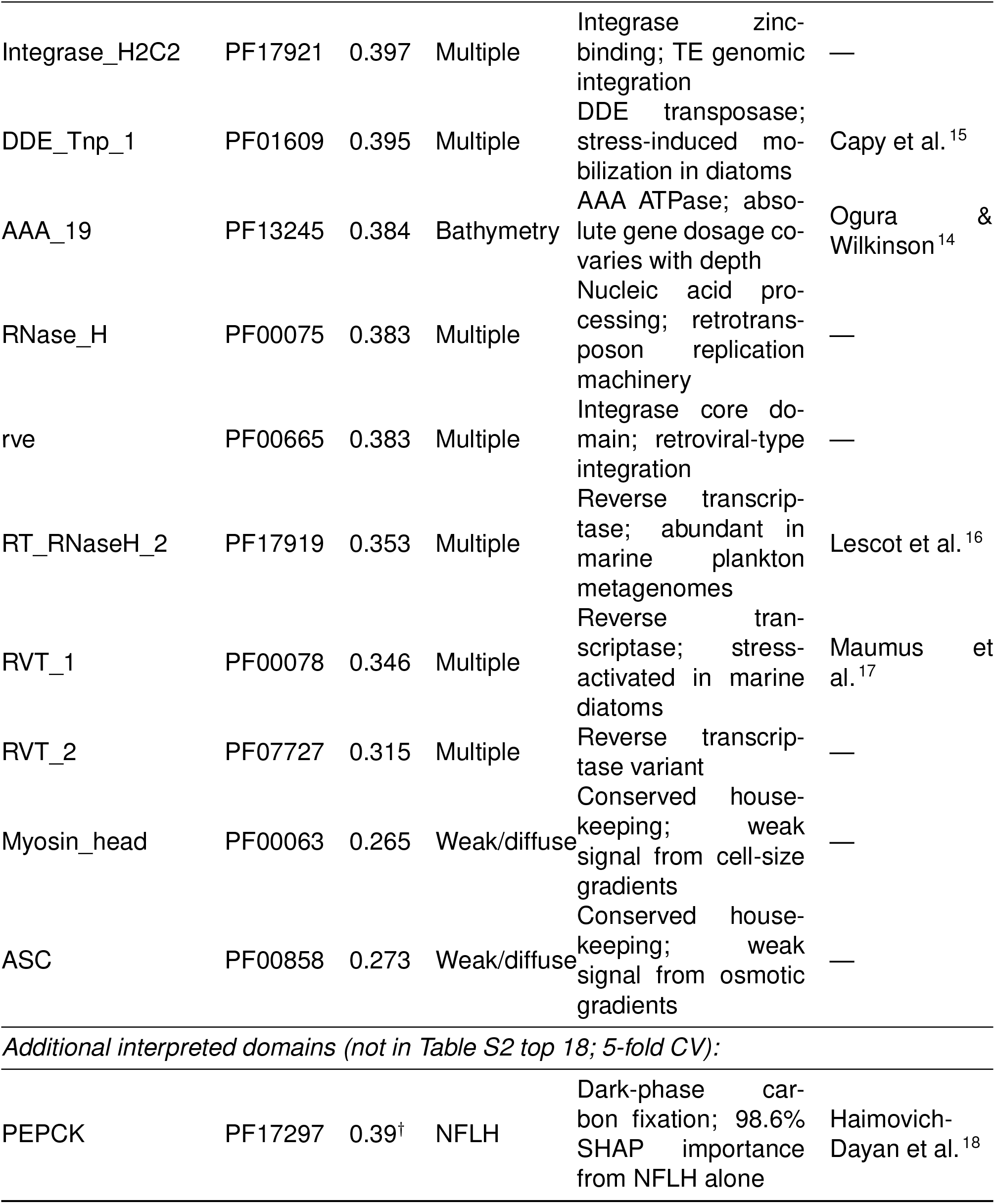
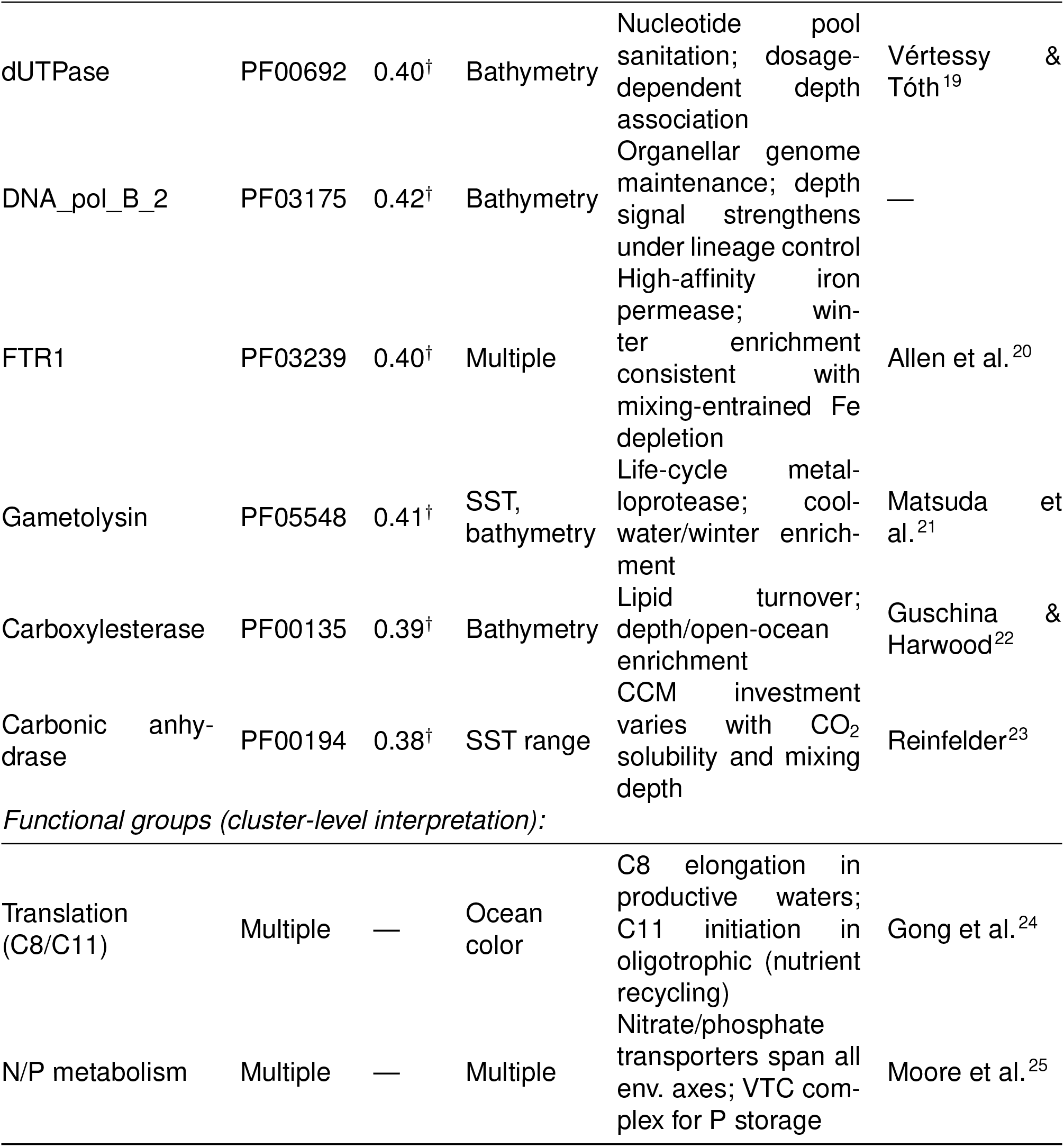
Biological interpretation of environment-predictive domains, related to Figures 3–5. *R*^2^ from 10-fold spatial block CV (Table S2) except where noted; 5-fold CV values marked with ^†^. SHAP denotes SHapley Additive exPlanations.

**Table S10.** Dipeptide O/E bias summary across three proteome fractions, related to Figure 6. Dipeptide O/E bias summary across three proteome fractions (500,000 sequences each).

| Proteome | Full sequence |  |  | LCR-masked |  |  |
| --- | --- | --- | --- | --- | --- | --- |
|  | SD | IQR | *n* biased | SD | IQR | *n* biased |
| Dark | 0.251 | 0.292 | 22/400 (5.5%) | 0.237 | 0.284 | 17/400 (4.3%) |
| Algal | 0.229 | 0.257 | 18/400 (4.5%) | 0.217 | 0.260 | 14/400 (3.5%) |
| Annotated | 0.163 | 0.171 | 8/400 (2.0%) | 0.152 | 0.163 | 5/400 (1.3%) |
| Dark/Annotated ratio | 1.54× | 1.71× | 2.75× | 1.56× | 1.74× | 3.40× |

SD and IQR of log<sub>2</sub>(O/E) across 400 dipeptides. “Biased” = |log<sub>2</sub>(O/E)| > 0.5 (>1.41-fold). LCR mask: 12-residue window, Shannon entropy < 1.5 bits.

**Table S11.** Dark vs. white proteome *π*_N_*/π*_S_ in *C. reinhardtii*, related to Figure 6. Dark vs. white proteome *π*_N_*/π*_S_ in *C. reinhardtii* (re-partition of Flowers et al. 2015 data by Pfam annotation status).

| Metric | Dark | White |
| --- | --- | --- |
| $n$ (after filter) | 4,085 | 7,038 |
| Median $\pi_N/\pi_S$ | 0.277 | 0.121 |
| IQR | [0.152, 0.429] | [0.051, 0.223] |
| Mean $\pi_N/\pi_S$ | 0.337 | 0.157 |
| Aggregate $\pi_N/\pi_S$ ($\sum \pi_N / \sum \pi_S$) | 0.277 | 0.151 |

**Table S12.** Cross-taxonomic summary of dark-vs-white selection across four species, related to Figure 6. Cross-taxonomic summary of dark-vs-white selection across four species. Fold denotes the within-study dark/white contrast for the metric shown; *π*_N_*/π*_S_, *d*_N_*/d*_S_, and positive-selection incidence operate at different evolutionary scales and are not pooled as a common rate. All partitions use Pfam-A domain annotation at *E <* 10^−9^ (or InterPro for *S. robusta*).

| Species | Lineage | Metric | $n_{\text{dark}}$ | $n_{\text{white}}$ | Fold | 95% CI | $p$ |
| --- | --- | --- | --- | --- | --- | --- | --- |
| *C. reinhardtii* | Chlorophyta | $\pi_N/\pi_S$ | 4,085 | 7,038 | 2.30 | — | $< 10^{-300}$ |
| *S. robusta* | Bacillariophyta | $\pi_N/\pi_S$ | 7,537 | 13,205 | 1.46 | [1.42, 1.51] | $5.7 \times 10^{-160}$ |
| *Synechococcus* | Cyanobacteria | $d_N/d_S$ | 1,255 | 3,363 | 1.84 | [1.75, 1.94] | $3.3 \times 10^{-117}$ |
| *T. pseudonana* | Bacillariophyta | pos. sel. incidence | 5,408 | 6,265 | 1.90<sup>†</sup> | — | $4.9 \times 10^{-21}$ |

<sup>†</sup>Incidence ratio: 9.3% dark vs. 4.9% white genes with site-model evidence for positive selection. The separately calculated dark-fraction enrichment was 1.34 (95% CI [1.27, 1.42]); that interval is not a CI for the 1.90 incidence ratio.

**Table S13.** NOVEL_33041 within-genome *d*_N_*/d*_S_ summary, related to Supplemental Text. NOVEL_33041 within-genome *d*_N_*/d*_S_ summary (66 copies, Nei-Gojobori method).

| Metric | Value |
| --- | --- |
| Global $\omega$ (mean $d_N$ / mean $d_S$) | 0.33 |
| Median per-pair $\omega$ | 0.32 |
| Pairs with $\omega < 1$ | 1,957 / 2,009 finite pairs (97.4%) |
| Mean $d_N$ | 0.022 |
| Mean $d_S$ | 0.066 |
| Pairs with $d_S < 0.01$ (gene conversion) | 352 / 2,145 (16.4%) |
| Pairs identical ($d_N = 0$, $d_S = 0$) | 26 / 2,145 (1.2%) |

**Table S14. Provenance of the 478 cultured reference genome assemblies used as reference proteomes, related to Methods**

Provided as a separate Excel file (TableS14_reference_genome_provenance.xlsx). One row per reference genome assembly (478 rows), giving the file name as used in the pipeline and in Data S2, organism and strain, NCBI accession, assembly name, submitter, BioProject, and release date for NCBI assemblies (284; GenBank and RefSeq), and the study that published each genome. Most of the remaining 194 assemblies were published in Nelson et al. (2021; https://doi.org/10.1016/j.chom.2020.12.005) or Nelson et al. (2019; https://doi.org/10.1016/j.isci.2018.12.035).

### SUPPLEMENTAL DATASETS

**Data S1. Domain–environment association and modeling results, related to Figures 2–5, Figure S3, and Methods**

Pfam domain count matrices (CLR-transformed and raw), Pfam–GEE Spearman correlation tables (20,318 raw feature columns from the permissive Pfam-A v37.2 screen at *E <* 10^−5^ *×* 37 environmental variables; *n* = 1,809), Pfam–AlphaEarth correlation tables (the same 20,318 raw feature columns *×* 64 embedding dimensions; *n* = 995), SHapley Additive exPlanations (SHAP) feature importance values from XGBoost reverse models (32 environmental targets), PCA/UMAP/t-SNE embeddings of the Pfam manifold, CCA joint embeddings and canonical loadings, XGBoost bidirectional cross-validated predictions (spatial block CV with 2° grid cells), per-sample summaries with GPS coordinates and ocean basin assignments, stratified correlation results by data source (metagenome, transcriptome, reference genome), and temporal hold-out cross-validation results (train 2009–2010, test 2011–2012). Deposited at Zenodo (concept DOI): https://doi.org/10.5281/zenodo.18538438.

**Data S2. LA^4^SR-classified algal protein sequences, related to Figure 1 and Methods**

The deposit contains 2,044 FASTA files from the mixed-source domain-analysis set: 1,005 ocean metagenomes, 606 MMETSP transcriptomes, 236 NCBI reference genomes, and 197 other cultured references. Each file contains proteins assigned to the algal class by LA^4^SR using the GPT-2-based algaGPT model with top-*k* sampling (*k* = 10; Methods). Total: 231.7 million algal sequences across the 2,044-sample analysis set (Table S1). Deposited at Zenodo (concept DOI): https://doi.org/10.5281/zenodo.18728836.

**Data S3. Pfam-A hmmsearch results archive, related to Figures 2–4, Figure S3, and Methods**

Per-sample `.aa.hmmsearch.tbl` tabular output files from HMMER3 `hmmsearch`^26^ against Pfam-A v37.2^27,28^ (strict threshold, E-value *<* 10^−9^; tabular output) for 2,318 processed samples from the mixed-source input collection, packaged as a `.tar.gz` archive. This intermediate processing ledger contains 2,318 of the 2,357 input samples; the 2,044 outputs with non-empty Pfam-A results define the domain-analysis set (Table S1C). Each file contains per-domain hit coordinates, E-values, and bit scores and enables independent re-thresholding, per-domain quantification, and domain architecture analysis without re-running `hmmsearch`. Deposited at Zenodo (concept DOI): https://doi.org/10.5281/zenodo.18786751.

**Data S4. RuBisCO lineage analysis package, related to Figure 1 and Methods**

A single consolidated Zenodo deposit (concept DOI: https://doi.org/10.5281/zenodo.18786775) containing the complete RuBisCO lineage analysis package, organized into three subdirectories with per-subdirectory READMEs and a top-level SHA-256 checksum manifest:

**form_I_chloroplast/per_sample_results_extended/**: 2,372 per-sample hmmsearch count tables from hmmsearch using 15 narrow lineage-specific rbcL HMMs, ten Form I lineages (Mamiellophyceae, Prasinophyceae, Pyramimonadales, Chlorellaceae, Trebouxiophyceae, Scenedesmaceae, Pelagophyceae, Bolidophyceae, Haptophyta, Cryptophyta) and five Form II myzozoan lineages (Symbiodiniaceae, Peridiniales, Gonyaulacales, Prorocentrales, Chromerida), plus broad green and red Form I rbcL profiles, run against LA^4^SR-extracted algal proteomes.

**form_II_nuclear/full_protein_results/**: 2,372 per-sample hmmsearch count tables from a curated Form II rbcL HMM (primarily dinoflagellate nuclear-encoded Form II) run against full predicted proteomes (not restricted to algal-classified sequences, since Form II rbcL is nuclear-encoded in dinoflagellates).

**hmm_profiles_and_alignments/**: The 18 .hmm profile files (with hmmpress-generated binary indexes), the 18 corresponding MAFFT reference alignments (.aln), and 49 input sequence files (raw, length-filtered, and CD-HIT 95% non-redundant sets with .clstr cluster membership files) from which the profiles were constructed, enabling full reproduction of the lineage-specific hmmsearch analysis.

This consolidated deposit supersedes three earlier separate deposits.

**Data S5. AlphaEarth satellite embedding matrix, related to Figures 3–5 and Methods**

A single TSV file containing 1,810 GPS-mapped analysis-set samples and 64 AlphaEarth embedding dimensions (A00–A63), extracted via the Google Earth Engine AlphaEarth foundation model at 10 m resolution. The table includes all 1,810 sample rows; 995 have AEF records, of which 969 contain no zero-filled dimensions and 26 were partially zero-filled during downstream feature preparation. Columns include sample identifiers, GPS coordinates (latitude, longitude), ocean basin assignment, and embedding dimensions. Despite terrestrial training, multiple dimensions correlate with oceanographic gradients (*|ρ|* up to 0.68 for SST and bathymetry; see main text). Deposited at Zenodo (concept DOI): https://doi.org/10.5281/zenodo.18786761.

**Data S6. KAN-CCA and sparse CCA cross-validation results, related to Figure 2 (panels G–T) and Methods**

The deposit contains two analysis variants. The nutrient-expanded sparse CCA files use 786 samples with complete metadata for 24 environmental variables. The KAN B-spline activations, fold-level KAN-CCA results, and domain-sparsity ablation use the earlier satellite-only complete-case matrix (969 samples; 19 environmental variables; ten 2° spatial-block folds) at *k* = 6, 13, 50, 100, and 500 domain features. Files are provided in TSV and JSON formats. Deposited at Zenodo (concept DOI): https://doi.org/10.5281/zenodo.18786766.

**Data S7. Pfam-dark protein-family discovery results, related to Figures 5–6, Table S4, and Methods**

Family-profile HMMs (`.hmm`; 33,950 whole-protein clusters from MMseqs2 clustering of 201 million Pfam-dark proteins at 30% sequence identity and 80% bidirectional coverage), a family count matrix (2,044 samples *×* 33,950 families), per-sample hmmsearch tblout files (`.tar.gz`), characterization table (prevalence across assemblies, geographic breadth, nr-dark fraction, cluster size), environmental correlation results (Spearman *ρ* and FDR-corrected *q*-values for 18 environmental variables), Pfam-dark-family-vs-Pfam-domain effect-size comparison (primary matched design, 1,523 samples and 18 environmental variables: median *|ρ|* = 0.157 vs. 0.068, 2.29-fold enrichment), ESMFold-predicted structures for the ten highest-ranked families (10 PDB files; sequences truncated to 400 aa per API limit), per-family pLDDT summary statistics and per-residue pLDDT confidence profiles, and Foldseek structural homology search results against PDB and AlphaFold Database (best-hit TM-scores, E-values, and alignment statistics). The package also includes InterProScan annotations for 35,667 representatives from the separate photosynthetic-anchor-neighbor screen; these representatives are not members of the 33,950-HMM count matrix. Deposited at Zenodo (concept DOI): https://doi.org/10.5281/zenodo.18786771.

**Data S8. DIAMOND BLASTp comparison results, related to Figure 1 and Methods**

Per-sample and summary statistics from DIAMOND BLASTp^1^ searches of LA^4^SR-classified algal and non-algal sequences against NCBI NR (E-value *<* 10^−9^, very-sensitive mode). The comparison covers 352,615,196 proteins from the 2,044-sample mixed-source domain-analysis set, a defined subset of the 447.7 million proteins submitted to LA^4^SR; proteins from the 313 input samples outside the domain-analysis set were not included. The deposit contains per-sample cross-tabulation, source-stratified summaries, an execution log, and the Python analysis script. Non-algal-classified sequences are 2.9-fold more likely to have NR homologs than algal-classified sequences (73.2% vs. 25.1%), supporting preferential retention of database-underrepresented sequences in the algal class. Deposited at Zenodo (concept DOI): https://doi.org/10.5281/zenodo.19441355.

**Data S9. Dark-proteome selection analysis and generative protein language model (GPLM) training sequences, related to the cross-taxonomic validation section and Table S12**

Per-gene selection data and dark/white proteome partitions cover four species spanning *∼*1.5 Gyr of photosynthetic evolution: *Chlamydomonas reinhardtii* (17,535 per-gene rows, *π*_N_*/π*_S_; Flowers et al. 2015^29^), *Seminavis robusta* (36,254 per-gene rows, *π*_N_*/π*_S_; Osuna-Cruz et al. 2020^30^), *Synechococcus* CC9311 and CC9902 (2,891 and 2,307 per-gene rows, *d*_N_*/d*_S_; Tai & Palenik 2011^31^), and *Thalassiosira pseudonana* (3,280 rows, the genes with nominal *p <* 0.05 in the published PAML positive-selection table; Koester et al. 2013^32^). These row counts are the complete per-gene sheets and include genes excluded from the tested subsets; the gene sets actually compared (e.g., 4,085 dark and 7,038 white *C. reinhardtii* genes after filtering; 20,891 *S. robusta* genes with *π*_N_*/π*_S_ data; 11,673 *T. pseudonana* proteins partitioned) are given in the Supplemental Text and Tables S11–S12. Each per-gene sheet records the original selection metric, our Pfam/InterPro-based dark/white partition, and domain annotations. Accompanying summary, provenance, and analysis scripts enable reproduction.

The same Data S9 release contains approximately 15 million Pfam-dark proteins and 15 million Pfam-annotated proteins from the 2,044-sample mixed-source ELF-NET domain-analysis set, each divided 90/10 into training and holdout subsets. Pfam-A hmmsearch tables accompany the annotated sequences. Proteins were assigned to the algal class by the 70-million-parameter GPT-2/nanoGPT-based algaGPT classifier. Deposited at Zenodo (concept DOI): https://doi.org/10.5281/zenodo.20486973.

**Data S10. Boltz-2 and AlphaFold 3 structure predictions for 3,600 rank-binned Pfam-dark protein-family representatives, related to Figure 7, Figure S7, and Methods**

Predicted three-dimensional structures for 3,600 representative Pfam-dark protein families were generated independently with Boltz-2 v2.2.1^33^ and AlphaFold 3. Sequences were rank-binned by environmental coupling strength (maximum *|ρ|* across environmental variables) into 10 deciles of 360 sequences each. The complementary outputs contain Boltz-2 PDB structures, confidence JSON files, mapping and rank-bin manifest tables, MSA cluster sizes, and a pLDDT summary, together with AlphaFold 3 structures, per-residue pLDDT, PAE matrices, pTM scores, and mapping information. Deposited at Zenodo (concept DOI): https://doi.org/10.5281/zenodo.20508680.

**Data S11. Protein language model learnability analysis, related to Figure 6 and Methods**

Training code, configuration files, analysis scripts, five trained GPT-2 Small checkpoints (*∼*5 GB), and source protein sequences (FASTA files, tokenized train/val splits; *∼*1.1 GB) for the five-corpus protein language model learnability comparison. The five corpora—annotated photosynthetic proteins (“white”; 66.5M training tokens), algal proteome (58.9M tokens), Pfam-dark proteins (63.1M tokens), composition-matched random sequences (55.4M tokens), and uniform random sequences (55.4M tokens)—were trained under identical GPT-2 Small hyperparameters (12 layers, 12 heads, 768-dimensional embeddings, 1,024-token context, 40,000 iterations; Methods). Code and results deposited at Zenodo (concept DOI): https://doi.org/10.5281/zenodo.20933332. Model checkpoints: https://huggingface.co/SarahDaakour/dark-whiteGPLM. Training data: https://huggingface.co/datasets/SarahDaakour/dark-whiteGPLM-data.

### SUPPLEMENTAL TEXT

#### Protein language models extract algal genomes from metagenomic dark matter

The pipeline presented here differs fundamentally from previous TARA Oceans eukaryotic analyses in its classification methodology, and this difference determines what fraction of the marine algal proteome is accessible (Table S1).

Previous large-scale analyses of TARA eukaryotic sequences—including the Marine Atlas of Tara Oceans Unigenes (MATOU)^34^ and subsequent functional analyses—relied on homology-based classification: Diamond^1^ alignment against UniRef90 with weighted Least Common Ancestor (wLCA) taxonomy assignment. These methods require high sequence similarity to database references. For well-characterized model organisms, homology-based recall is high; for non-model algal lineages with limited reference genomes, BLAST-type searches typically return hits for only *∼*30% of predicted proteins.^2^ Carradec et al. reported that 48.3% of MATOU unigenes received taxonomic assignment at any level, meaning more than half the eukaryotic metagenome was unclassifiable.^34^ The unclassified fraction—genomic “dark matter”—was excluded from downstream functional analysis.

A complementary approach applied co-abundance gene clustering to the same MATOU catalog, grouping unigenes with correlated abundance profiles across 365 metagenomes to reconstruct 924 metagenomics-based transcriptomes (MGTs) representing individual eukaryotic organisms.^3^ This organism-centric strategy partially overcomes the gene-centric limitation of the original catalog and enabled functional insights such as lineage-specific dimethylsulfoniopropionate (DMSP) cycling. However, the MGT approach inherits the taxonomic assignment limitations of the underlying MATOU catalog and remains dependent on reference-based homology for functional annotation.

LA^4^SR operates on a different principle. As a transformer-based protein language model fine-tuned on the full breadth of available algal genomics data, LA^4^SR classifies sequences based on learned amino acid composition patterns rather than sequence similarity to known references.^2^ Every input sequence receives a classification score regardless of whether it has database homologs, achieving near-complete recall on algal proteomes where BLAST-type methods recover less than a third of sequences.^2^ This distinction is not incremental—it determines whether the majority of the algal proteome is visible or invisible to analysis. The computational difference is proportionate: LA^4^SR achieves *>*10,000-fold speedup over BLAST-based classification,^2^ enabling processing of the full 364 GB input dataset—a scale at which homology search against comprehensive databases such as NR is computationally prohibitive.

The consequences propagate through the pipeline. Because LA^4^SR provides a classification score for every sequence regardless of homology, the downstream PFAM annotation, environmental association, and bidirectional modeling presented here operate on a more complete set of algal sequences than homology-based catalogs can access. The broad 2,357-sample input collection spans 5.3-fold more samples than MATOU’s 441 samples from 68 stations,^34^ and the 2,044-sample domain-analysis set contains 231.7 million algal sequences (Table S1). These sequences include both the homology-accessible fraction and sequences lacking database homologs that previous analyses discarded. However, this classification completeness does not guarantee functional annotation: sequences without Pfam-A hits remain functionally uncharacterized regardless of whether they are classified as algal by LA^4^SR or missed entirely by homology-based methods. The advantage is access—all sequences are included in the analysis—not interpretation of currently unannotated sequences. The 279,575 significant domain–AlphaEarth associations (FDR *<* 0.05; *n* = 995) and bidirectional predictability from direct GEE environmental variables (reverse-model *R*^2^ up to 0.42 under spatial block CV, for bathymetry with *n* = 1,523) reported here thus reflect functional coupling across the annotatable portion of the full algal proteome.

#### Algal proteome extraction details

The initial Form I-only RuBisCO search across 1,203 ocean metagenome assemblies (TARA Oceans and OSD) detected RuBisCO in 1,069 (88.9%), with a median of 13 distinct sequences per assembly (mean 19.3, range 1–113). This count excludes dinoflagellates, which encode Form II RuBisCO in the nuclear genome rather than the plastid genome; Form II is also the RuBisCO form of chromerids (the fifth Form II lineage in our panel).^7,8^ Because LA^4^SR targets chloroplast gene structure signatures, nuclear-encoded dinoflagellate rbcL may not be captured by the algal-classified protein subset. To address this, we extended the hidden Markov model (HMM) panel with five myzozoan Form II lineages (four dinoflagellate lineages, Symbiodiniaceae, Peridiniales, Gonyaulacales, and Prorocentrales, plus Chromerida; 1,183 reference sequences after polyprotein splitting) and rescanned *all* predicted proteins per assembly (Methods). The extended 15-lineage search detected RuBisCO in 1,118 of the 1,203 ocean metagenome assemblies (92.9%), up from 1,069 (88.9%) with Form I alone. Form II was detected in 1,002 assemblies (83.3%), yielding 47,356 myzozoan sequences—dominated by Prorocentrales (28,908 sequences in 917 assemblies) and Gonyaulacales (11,532 in 764). With both forms combined, the median rose from 13 to 18 distinct RuBisCO sequences per assembly (mean 56.5, range 0–721) spanning a median of 6 co-occurring photosynthetic lineages per assembly (mean 6.2). Combined with MMETSP and reference genomes, total unique RuBisCO detections reach 149,733 sequences (Form I: 22,743; Form II: 126,990; Table S3). The Form I count is the union of sequences detected by the broad green-lineage (Form IB) and red-lineage (Form ID) HMMs; counting green and red hits separately and summing would yield 42,757, but 20,014 sequences hit both broad profiles and are resolved to a single lineage assignment at the lineage-specific stage, so the deduplicated Form I total is 22,743. Because Form I rbcL is typically single-copy per plastid genome,^6^ each Form I detection approximates one photosynthetic taxon. Form II rbcL, however, is nuclear-encoded and multi-copy in dinoflagellates; raw Form II counts therefore overestimate taxon diversity. Zhang and Lin^8^ reported *∼*3 copies per genome in *Prorocentrum minimum*, but copy number varies substantially across dinoflagellate species (2–6 or more copies). Applying the central estimate of *∼*3 copies per genome yields *∼*65,000 photosynthetic algal taxa (*≈*22,743 + 126,990/3). Assuming 2–6 copies per genome brackets this estimate at *∼*44,000–86,000 taxa (*≈*22,743 + 126,990/6 to 22,743 + 126,990/2). This is an order-of-magnitude approximation and a lower bound, subject to additional caveats: (i) pseudogenes may inflate sequence counts;^6^ (ii) assembly fragmentation can split multi-exon rbcL loci into separate contigs, artificially increasing apparent copy number;^35^ and (iii) the correction assumes uniform copy number across all dinoflagellate lineages, whereas empirical data are available only for a few species.^8^

#### Satellite environmental context details

To characterize the relationship between AlphaEarth dimensions and direct oceanographic measurements, we correlated all 64 embedding dimensions against 29 GEE environmental variables for samples with AEF records and environmental metadata. Of 1,856 dimension–variable pairs tested, 819 (44.1%) achieved significance at FDR *<* 0.05, with maximum *|ρ|* = 0.68. Multiple dimensions correlated strongly (*|ρ| >* 0.5) with known oceanographic gradients: A08 with sea surface temperature maximum (*ρ* = *−*0.68), A49 with mean SST (*ρ* = 0.68), A21 with bathymetry (*|ρ|* = 0.67), A41 with bathymetry (*ρ* = 0.59), A23 with chlorophyll-a maximum (*ρ* = 0.57), and A03 with MODIS SST (*ρ* = 0.57). Remote sensing reflectance at ocean-color wavelengths (412–488 nm) correlated with multiple dimensions at *|ρ| >* 0.5. The hub dimensions most associated with PFAM domain composition—A31, A25, and A14— mapped to coastal/depth gradients (A31: bathymetry *ρ* = *−*0.53), thermal seasonality (A25: SST range *ρ* = 0.40), and temperature combined with coastal proximity (A14: air temperature *ρ* = 0.52, distance to coast *ρ* = 0.47), indicating that domain–embedding associations reflect identifiable environmental gradients rather than opaque latent features.

#### Temporal stability of domain–environment coupling

The main analyses draw on cross-sectional data spanning TARA Oceans (2009–2012) and OSD (2014) campaigns. To assess whether domain–environment associations persist across time, we linked 973 of the 1,003 TARA Oceans and OSD assemblies in the 1,810-sample GPS-mapped set (97.0%) to collection dates via the PANGAEA station registry (DOI: 10.1594/PANGAEA.842237)^36^ and analyzed temporal stability across four years of sampling.

##### Temporal analysis methods

For TARA assemblies, GPS coordinates were matched to the nearest of 211 PANGAEA stations by Euclidean distance in latitude-longitude space (threshold *<*0.5 degrees; median match distance 0.055 degrees; 847 of 877 TARA assemblies matched). For OSD assemblies, collection dates were obtained from OSD metadata via coordinate matching (threshold *<*0.2 degrees; 126 of 126 OSD assemblies matched). Hemisphere-aware seasonal assignment used meteorological seasons: Northern Hemisphere (latitude *≥* 0): DJF = winter, MAM = spring, JJA = summer, SON = autumn; Southern Hemisphere (latitude *<* 0): seasons reversed (DJF = summer, etc.).

Year-to-year correlation stability was computed by subsetting TARA assemblies by year (2009: *n* = 186; 2010: *n* = 244; 2011: *n* = 340; 2012: *n* = 77), computing Spearman correlations between 200 top-variable PFAM domains and 37 environmental variables within each year subset, and computing pairwise Pearson correlations of the resulting rho vectors across all six year pairs. Sign concordance rate was the fraction of domain–environment pairs with the same sign across all four years.

PERMANOVA tested seasonal effects on PFAM composition using Bray-Curtis dissimilarity on raw counts of the top 500 most-variable domains (*n* = 973 samples). The pseudo-*F* statistic was computed as the ratio of between-group to within-group sum of squared distances; *p*-values were obtained from 999 permutations of season labels. Analysis of similarity (ANOSIM)^37^ tested within-station versus between-station dissimilarity using 117 TARA stations as the grouping factor. Mantel tests^38^ correlated temporal distance (absolute days between collection) with Bray-Curtis distance, stratified by ocean basin; partial Mantel tests controlled for geographic distance (Euclidean in latitude-longitude) using 999 permutations.

Temporal cross-validation used strict year-based hold-out: training on 2009–2010 samples (*n* = 354 with non-missing target values of the 430 dated 2009–2010 samples) and testing on 2011–2012 samples (*n* = 417). PFAM abundances were CLR-transformed (pseudocount = 1) and reduced to 100 principal components; XGBoost models used identical hyperparameters to the spatial block CV (max_depth = 6, learning_rate = 0.1, n_estimators = 200, subsample = 0.8, colsample_bytree = 0.8). A secondary split (train 2010–2011, test 2009 + 2012) provided robustness assessment. Forward temporal hold-out (environment *→* individual domain abundance) used the same year-based splits and XGBoost hyperparameters applied to the top 200 domains by spatial *R*^2^ (*n* = 973 dateable samples; 25 environmental features matching the spatial CV analysis).

##### Temporal analysis results

Adjacent sampling years showed correlated domain–environment association profiles (2009–2010: *r* = 0.41; 2010–2011: *r* = 0.27), with correlation decaying over longer intervals (2009–2012: *r* = 0.11; mean pairwise *r* = 0.22, range 0.08–0.41). Sign concordance across all four years was 25.0%, indicating that while the strongest associations persist, the full association landscape shifts year to year (mean pairwise *r* = 0.22).

Seasonal effects on PFAM composition were statistically significant (PERMANOVA: pseudo-*F* = 8.38, *p* = 0.001, *n* = 973; effect persistent across ocean basins), with over half (103/200) of the most variable domains showing seasonal differential abundance at FDR *<* 0.05, predominantly peaking in winter. The PFAM manifold captured this temporal structure: UMAP2 correlated with month-of-year (*r* = 0.35 for cosine-encoded month; *p <* 10^−28^), and samples collected closer in time were closer in PFAM space (Mantel *ρ* = 0.10, *p* = 0.005). The seasonal *R*^2^ of 0.025 is consistent with spatial geography dominating over temporal seasonality in structuring domain composition—an expected pattern for cross-sectional expedition data.

Within-station variance analysis (117 unique stations, median 6 assemblies/station) revealed that within-station samples were significantly more similar than between-station samples (Bray-Curtis 0.86 vs. 0.92; ANOSIM *R* = 0.056, *p* = 0.001), but a partial Mantel test controlling for geographic distance eliminated this signal (*ρ* = *−*0.04, *p* = 1.0). The apparent temporal clustering thus reflects spatial autocorrelation along the expedition track rather than independent temporal dynamics.

Year-based cross-validation provided the most stringent test, though its interpretation requires care. Training XGBoost models on 2009–2010 samples (*n* = 354) and testing on 2011–2012 (*n* = 417) creates a strict split with no sample overlap. However, because the TARA expedition sailed continuously from 2009 to 2012, this year-based partition also separates samples geographically: different years correspond largely to different ocean basins along the cruise track. The split therefore tests geographic extrapolation to new regions as much as temporal persistence, and the environmental targets themselves (AlphaEarth annual satellite embeddings for 2017 onward, GEE climatologies) are not contemporaneous with sampling. With this caveat, the year-based hold-out reduced predictive performance compared to spatial block CV: SST *R*^2^ = 0.16–0.25 versus spatial *R*^2^ = 0.38–0.40 (mean Δ*R*^2^ = *−*0.17 across the three SST targets; mean spatial-minus-temporal difference across the four targets in Figure S5D = 0.24); bathymetry *R*^2^ = *−*0.01 versus spatial *R*^2^ = 0.42 (Δ*R*^2^ = *−*0.43). The bathymetry failure is expected given this design: bathymetry is time-invariant, so the year-based split tests whether depth–composition associations learned at one set of geographic locations transfer to locations visited in later years—a geographic extrapolation test, not a temporal one. The SST retention (*R*^2^ = 0.16–0.25) is more informative, indicating that domain–temperature coupling generalizes across ocean regions not represented in training, retaining 40–65% of predictive variance despite the geographic shift.

Extending the year-based hold-out to the forward direction (environment *→* individual domain abundance) revealed more severe attenuation: among the top 200 domains by spatial *R*^2^, none achieved positive *R*^2^ under either split (primary median *R*^2^ = *−*0.28, secondary median *R*^2^ = *−*0.34; *n* = 973 dateable samples; Data S1). This asymmetry between forward and reverse directions is expected: the reverse model aggregates thousands of domains to predict a single environmental variable, averaging out region-specific noise and capturing the shared environmental signal, while single-domain predictions from a handful of environmental features are dominated by stochastic variation among geographic regions. The asymmetry thus reflects the statistical distinction between multivariate-to-univariate projection (robust to geographic shift) and univariate-to-univariate prediction (fragile), not necessarily temporal instability.

These results have two implications. First, domain–environment coupling established from spatial co-occurrence generalizes to geographically independent ocean regions for aggregate predictions (PFAM *→* SST, *R*^2^ = 0.16–0.25 under year-based geographic hold-out), while individual domain predictions do not transfer across regions. Second, the TARA expedition’s continuous spatial trajectory makes year and geography inseparable in this dataset, precluding a clean test of temporal stability independent of geographic extrapolation. The 2014 OSD samples (*n* = 126) provide partial temporal and geographic independence (2–5 year gap from TARA, different sampling design) but concentrate in a single month (June–July), precluding seasonal decomposition. Future designs with repeated temporal sampling at fixed locations would enable stronger inferences about temporal versus geographic contributions to prediction attenuation.

#### Extended bidirectional modeling results

##### GO enrichment and per-environment-group domain interpretation

To characterize the functional roles of environment-associated domains, we performed Gene Ontology (GO)^39^ enrichment analysis using hypergeometric tests on PFAM sets defined by SHapley Additive exPlanations (SHAP) importance. Mapping 5,182 PFAMs to 9,842 GO associations via Pfam2GO,^40^ we tested for functional over-representation in temperature-, depth-, light-, and optical-associated domain sets.

In the temperature-associated PFAM set, the top-ranked domain (PF11999, Ice_binding) implicates cold adaptation mechanisms. Heat shock protein domains (HSP20/alpha-crystallin family, PF00011) also ranked among temperature-predictive features, consistent with thermal stress response pathways.^41^ Transposase domains (DDE_Tnp_IS66, DDE_Tnp_1) appeared prominently across multiple environmental groups, and TE-associated domains recur in Figure 4 clusters spanning distinct oceanographic axes: elevation (C1; rve), bathymetry (C15; RVT_2), thermal variability (C6; DDE_Tnp_4, Integrase_H2C2), stable temperature (C12; DDE_1), and warm SST (C13; RVT_1), indicating that mobile element abundance covaries with multiple oceanographic axes rather than a single driver.^15^ The prominence of transposase domains among environment-predictive features raises the question of whether their importance reflects biology or artifact (see Discussion in main text). Two sensitivity analyses address this. First, excluding all 23 transposase-family domains (DDE superfamily, integrases, reverse transcriptases) from five-fold CV of the XGBoost reverse model (*n* = 1,279 GEE subset; each fold 80/20) changed mean fold bathymetry *R*^2^ by only *−*1.4% (0.4909*→*0.4842) and MODIS SST *R*^2^ by +0.6% (0.6076*→*0.6114), with 99.7% of significant domain–environment correlations retained. Thus, non-TE features retain predictive signal when these domains are excluded; the ablation does not estimate the importance of the removed TE features. (These *R*^2^ values differ from the 10-fold spatial-block CV values reported in the main text because this sensitivity analysis uses standard five-fold CV.) Second, normalizing PFAM counts to relative abundance (dividing by total domains per sample) did not reduce transposase SHAP rankings: DDE_Tnp_IS66 (PF03050) moved from rank 6 to rank 2 for bathymetry prediction, and HTH_Tnp_IS1 (PF01527) remained at rank 4. This normalization argues against total-domain-count scaling as the sole explanation for the transposase rankings; the relative composition of mobile-element domains remains informative for environmental prediction.

For depth-associated PFAMs, dUTPase (PF00692) and PIF1 helicase (PF05970) ranked highest, both involved in DNA repair and genome maintenance—functions potentially important under pressure stress^42^ or low-light conditions at depth. Light-associated domains included PEPCK (phosphoenolpyruvate carboxykinase, PF17297), linking irradiance to gluconeogenesis and carbon metabolism.^18^

Environment-predictable PFAMs (the 22 domains with held-out *R*^2^ *>* 0.3 in a top-100 single-split forward screen; background 9,611 domains) showed significant GO enrichment (7 terms at FDR *<* 0.05; Figure S4A): RNA-DNA hybrid ribonuclease activity (GO:0004523), myosin complex (GO:0016459), DNA helicase activity (GO:0003678), monoatomic ion transport (GO:0006811), cytoskeletal motor activity (GO:0003774), DNA integration (GO:0015074), and protein binding (GO:0005515). This functional signature—encompassing nucleic acid processing, cytoskeletal dynamics, and ion homeostasis—indicates that environment-predictable domains encode housekeeping and structural functions with systematic oceanographic distributions. Light-associated domains separately showed enrichment for photosystem II oxygen-evolving complex (GO:0009654; raw *p* = 0.0016, FDR = 0.074) and photosynthesis-related terms, consistent with photosynthetic machinery varying with irradiance.^43^ However, many environment-associated domains lacked GO annotations, and algal-specific functions remain incompletely characterized in current databases.^44^

##### AEF bicluster detailed descriptions

The 16 row clusters (C1–C16) partition 716 PFAM domains into groups with coherent environmental associations, ranging in size from 21 (C4) to 91 (C12) domains. Temperature dominates as the primary environmental signature: seven clusters associate with SST or air temperature variants, including cold-water clusters C3 (53 domains, SST min*−*; top PFAMs: DNA_pol_B, Chloroa_b-bind) and C14 (30 domains, SST max*−*; Citrate_synt, HSP90), variable-temperature clusters C6 (81 domains, SST range+; Photo_RC, Arf), C9 (82 domains, SST range+; AAA, Carb_anhydrase), and C12 (91 domains, SST range*−*; GIDA, 2OG-FeII_Oxy), and warm-water clusters C4 (21 domains, air temp.+; Actin) and C13 (36 domains, MODIS SST+; ketoacyl-synt). Bathymetry and elevation partition into complementary clusters: C1 (29 domains, elevation+; ATP-synt_ab) and C2 (67 domains, bathymetry*−*; ABC_tran) capture depth gradients from opposing directions. Chlorophyll-associated cluster C5 (29 domains, Chl-a mean*−*; Histone, Histone_H2A_C) links chromatin remodeling to low-productivity waters. Ocean color clusters capture the bio-optical gradient: C8 (27 domains, Rrs443*−*; GTP_EFTU) contains translation machinery characteristic of phytoplankton-rich waters, while C11 (22 domains, Rrs412+; TFIIA) shows transcriptional regulators reflecting metabolic efficiency in oligotrophic conditions. This functional dichotomy—growth maximization versus resource efficiency—maps directly onto the bio-optical gradient detected by satellite remote sensing.

Biological interpretation reveals coherent functional signatures across environmental axes. Cold-water clusters are enriched for photosynthetic machinery (Photo_RC, Chloroa_b-bind in C3 and C6) and core metabolic enzymes, consistent with cold, nutrient-rich waters supporting phytoplankton productivity;^45,46^ the co-occurrence of glycosyl hydrolases in C3 may further indicate heterotrophic supplementation among cold-adapted species.^47^ In C6, thermal variability creates stress imbalance of the photosynthetic machinery, which can drive reactive oxygen species (ROS) production;^48^ the co-enrichment of mobile elements (DDE_Tnp_4, Integrase_H2C2) and DNA methylase in this cluster is consistent with stress-induced transposon mobilization in response to photosynthetic redox imbalance.^15,49^ Variable-temperature cluster C9 contains carbonic anhydrase, a core component of the carbon-concentrating mechanism (CCM): because CO_2_ solubility decreases with rising temperature, SST variability would drive fluctuating CO_2_ availability and select for CCM capacity.^23^ Stable-temperature cluster C12 includes caleosin, a lipid-droplet-associated protein found in oil bodies of *Chlorella* and other microalgae, suggesting a role in lipid storage under thermally stable conditions.^50^ Warm-water clusters C4 (Actin, eRF1_2) and C13 (ketoacyl-synt) link elevated temperature to cell division and fatty acid synthesis, respectively; temperature influences microalgal lipid content and composition, with species-specific responses that may partly reflect underlying taxonomic signal.^51^ Among depth-associated clusters, C2 contains HSP70, which responds not only to thermal stress but also to light, chemical, and life-stage cues;^52^ its enrichment at depth may additionally reflect hydrostatic pressure as a contributing stressor. Chlorophyll-associated cluster C5 links chromatin remodeling (Histone, Histone_H2A_C) to low-productivity waters; Yan et al.^53^ demonstrated that nitrogen deprivation induces correlated changes in chromatin reorganization and gene expression in the marine microalga *Nannochloropsis oceanica*, suggesting that the chromatin enrichment may reflect nutrient-stress-driven genome compaction rather than preparation for cell division. The co-occurrence of ubiquitin-proteasome components (JAB, zf-C3HC4) in this cluster is consistent with enhanced protein turnover under nutrient limitation.^54^ HSP90 in C14 is expressed as a heat-shock response in *Chlamydomonas*, linking this cool-water cluster to thermal stress response pathways.^55^ This environmental organization of protein domains—from photosynthetic cold-water clusters through ion homeostasis in nearshore C16 (Ion_trans, AAA_30)— demonstrates that the marine algal functional repertoire is structured along canonical oceanographic axes.^56^

##### Partial correlation analysis: controlling for taxonomic composition

To assess whether domain-environment associations persist beyond simple taxonomic sorting, we computed partial Spearman correlations controlling for dominant RuBisCO lineage identity (assigned per sample from the lineage with highest RuBisCO count among ten Form I algal lineages; *n* = 1,005 metagenome samples). The partial correlation analysis used Form I lineages. Of the top 20 associations by *|ρ|*, all 20 retained significance (*p <* 0.05, Bonferroni-corrected) after lineage correction, with a mean *|ρ|* reduction of *−*5.5%; that is, partial correlations were on average 5.5% stronger than the zero-order correlations. In a supplementary analysis sampling the strongest association per GEE environmental variable across 12 key oceanographic variables, 11 of 12 retained significance after lineage correction, with a mean *|ρ|* reduction of 20.1%. Physical oceanographic variables (SST, bathymetry, distance to coast) showed minimal attenuation under lineage correction (6–25% *|ρ|* reduction), while productivity-related variables (chlorophyll-a, particulate organic carbon [POC]) showed larger reductions (*∼*45%) but remained significant, suggesting these are more strongly influenced by taxonomic composition. The single association that lost significance (precipitation) also changed sign, indicating it was entirely driven by lineage sorting. Across both analyses, 31 of 32 tested associations retained significance and sign after controlling for lineage identity. This test used the dominant lineage at the RuBisCO level to capture phylum-level taxonomic turnover (e.g., haptophyte- vs. chlorophyte-dominated communities).

##### Lineage-controlled Pfam-dark-family versus Pfam-domain enrichment comparison

A finer-grained lineage control tests whether the 2.29-fold Pfam-dark-family-over-Pfam-domain enrichment in environmental coupling is an artifact of lineage composition. We computed partial Spearman correlations on the 1,133-sample intersection carrying both count matrices, 18 GEE+WOA environmental variables, and RuBisCO lineage data, using all 15 RuBisCO lineage counts (10 Form I lineages, six green and four red, plus 5 Form II myzozoan lineages) as continuous covariates via QR projection on rank-transformed variables. CLR normalization (pseudocount +1) was applied to counts; features present in fewer than 10 samples were excluded. Without lineage control on this sample subset, Pfam-dark families showed 2.08-fold enrichment over Pfam domains (13,330 Pfam domains, 31,143 Pfam-dark families; median *|ρ|*: 0.1202 vs. 0.0579). After controlling for 15 RuBisCO lineage covariates, the enrichment ratio decreased to 1.83-fold (95% CI [1.82, 1.85]; partial median *|ρ|*: 0.1303 vs. 0.0710; Mann-Whitney *p <* 10^−300^), a 22.5% attenuation of the excess above parity. The enrichment persists after lineage correction, arguing against lineage-composition covariates as the sole explanation.

##### Metagenome-only association sensitivity

To assess whether the combination of metagenomes, transcriptomes, and reference genomes biases the reported associations, we repeated key analyses on the ocean metagenome subset alone (TARA Oceans and OSD; *n* = 1,005). Domain–environment correlations for directly measured oceanographic variables persisted: 72% of top SST associations and 86% of top normalized fluorescence line height (NFLH) associations retained FDR significance (*n* = 773), with 96% and 100% sign concordance, respectively. The strongest metagenome-only association (PF11999.13 *×* SST, *ρ* = *−*0.321, *p* = 5.86 *×* 10^−20^) remained significant. Stratified AlphaEarth embedding analysis (Data S1) showed that domain–embedding coupling was strongest in the metagenome stratum (21.7% FDR-significant, *n* = 364), exceeding both transcriptome (2.3%, *n* = 240) and reference genome (3.3%, *n* = 391) strata. These sensitivity results indicate that cultured or reference samples do not solely account for the reported associations.

#### Pfam-to-AlphaEarth projection model

As an independent validation that domain composition encodes sufficient environmental information to reconstruct satellite-derived landscape descriptors, we developed a neural projection model mapping Pfam profiles to AlphaEarth embeddings. This model is not used for primary analyses in the Results; it serves as a methodological complement to the XGBoost and CCA frameworks, confirming that domain– environment coupling supports cross-modal prediction. The model architecture consists of a deep multilayer perceptron with bottleneck layers: input batch normalization, followed by fully-connected layers (9,611 *→* 2,048 *→* 512 *→* 128 *→* 64 units) with ReLU activations and dropout regularization (0.2–0.3). Total model capacity is approximately 20.8 million parameters.

Compositional data were transformed using the centered log-ratio (CLR):^57^

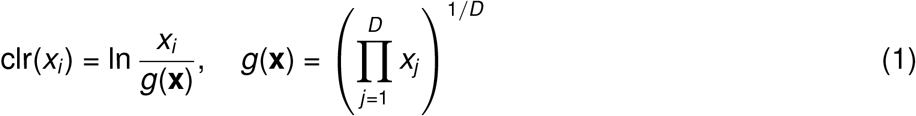

where *x_i_* is the abundance of domain *i* and *g*(**x**) is the geometric mean across all *D* domains. CLR was chosen over the isometric log-ratio (ILR) transform because CLR components maintain a one-to-one correspondence with individual domains, which is essential for per-domain SHAP decomposition, forward modeling of individual domain abundances, and biological interpretability; ILR components are log-ratios between domain groups, precluding direct domain-level attribution.^58^ Zero counts in PFAM abundance matrices were replaced with a pseudocount of 0.5 prior to CLR transformation to avoid undefined logarithms while minimizing perturbation to non-zero values. Training data comprised 995 samples with complete CLR-normalized Pfam profiles (9,611 domains) and AEF records (64 dimensions; 969 with no zero-filled dimensions and 26 partially zero-filled). Data were split using geographic holdout: Mediterranean Sea samples (n=242) were reserved for testing to evaluate generalization to unseen oceanographic regions. Remaining samples were partitioned 80/20 for training (n=603) and validation (n=150).

The model was trained using AdamW optimizer^59^ with initial learning rate 10^−3^, cosine annealing schedule, and weight decay 10^−4^. A combined loss function weighted mean squared error (MSE) reconstruction (80%) and cosine similarity (20%) to optimize both magnitude and direction in embedding space. Cosine similarity measures the angular alignment between predicted (**y***^*) and target (**y**) embedding vectors:

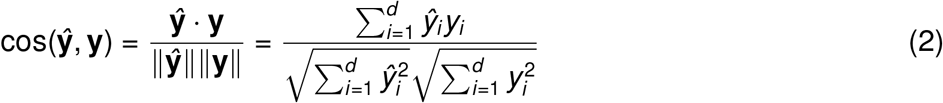

where *d* is the embedding dimensionality (64 for AlphaEarth). This metric is bounded [*−*1, 1], with 1 indicating identical direction regardless of magnitude. Early stopping with 30-epoch patience on validation loss prevented overfitting. Training used automatic mixed-precision (AMP) computation on NVIDIA GPUs.

Model performance was evaluated using per-dimension *R*^2^, mean *R*^2^, and cosine similarity between predicted and actual embeddings. Baselines included XGBoost regression (64 independent models) and ridge regression. Negative controls verified that shuffled features and random sample pairings produced near-zero *R*^2^, confirming the model learns meaningful Pfam-to-AlphaEarth mappings rather than spurious patterns.

#### Extended CCA: higher-order canonical variates and nonlinearity analysis

Individual interpretation of CC2–CC5 revealed that the canonical structure captures multiple independent environmental gradients beyond the dominant thermal axis (Figure S2A). CC2 (*r* = 0.72) captured a coastal–pelagic gradient, with distance to coast (*ρ* = +0.56), bathymetry (*ρ* = *−*0.54), and SST maximum (*ρ* = *−*0.53) as the three strongest correlates. CC3 (*r* = 0.68) captured a productivity–seasonality gradient dominated by normalized fluorescence line height (*ρ* = *−*0.43) and SST range (*ρ* = +0.42), reflecting phytoplankton biomass variation coupled with thermal seasonality. CC4 (*r* = 0.63) combined SST seasonality (*ρ* = +0.33) with solar radiation (*ρ* = +0.31), while CC5 (*r* = 0.60) loaded exclusively on short-wavelength ocean color bands (rrs_443: *ρ* = *−*0.63; rrs_412: *ρ* = *−*0.62; rrs_469: *ρ* = *−*0.60), capturing a water clarity gradient inversely related to productivity. Split-half cross-validation (100 iterations, stratified by ocean basin) confirmed that CC1 is the most stable axis, with only 9.7% out-of-sample shrinkage (observed *r* = 0.82, cross-validated median *r* = 0.74), whereas CC2–CC5 showed 30–45% shrinkage (Figure S2G), consistent with CCA’s known tendency to capitalize on noise in higher-order components.^60^ The first five components nevertheless retained meaningful out-of-sample signal (cross-validated median *r ≥* 0.37).

We evaluated sparsity and nonlinearity in two analysis variants (Methods). A nutrient-expanded sparse CCA used 786 samples with complete metadata for 24 environmental variables. The KAN-CCA nonlinearity analysis used a separate satellite-only complete-case matrix (969 samples; 19 environmental variables). In the primary 10-fold comparison on the 13-domain sparse subset, KAN-CCA and linear sparse CCA did not differ (fold-averaged test correlation across CC1–CC3: 0.414 vs. 0.395, paired *p* = 0.769; per-component *p >* 0.46; Table S5, F7). In the separate sparsity-ablation run at *k* = 13 domain features, KAN-CCA did not outperform linear sparse CCA on CC1 (+15.8%, *p* = 0.201) but improved CC2 (+57.4%, *q* = 0.012) and CC3 (+65.9%, *q* = 0.006; Benjamini–Hochberg FDR across three components). These secondary-axis findings are exploratory. Full feature weights, domain identities, and KAN spline analyses are detailed below and summarized in Figure 2G–T.

#### Sparse CCA and KAN-CCA details

The nutrient-expanded L1-penalized sparse CCA used 786 samples and 24 environmental variables, including dissolved nutrients. It imposed strong environmental-side sparsity (3–5 features per component at optimal *c_u_* = 50, *c_v_*= 1.5; mean test correlation CC1 = 0.373 *±* 0.137) while retaining broad domain-side coverage; mean test correlation averaged 0.20 across 12 components. On CC1, only four environmental features carried non-zero weight: SST (0.660), dissolved oxygen (*−*0.748), phosphate (*−*0.047), and particulate organic carbon (*−*0.045), indicating that the primary domain–environment axis is driven by a temperature–oxygen gradient with minor nutrient modulation. A nutrient-dominated component emerged on CC4 (nitrate *−*0.898, phosphate *−*0.416, silicate *−*0.136), and a mixing-dominated component on CC10 (mixed layer depth [MLD] 0.907, silicate 0.321).

Separately, KAN-CCA^61^ was evaluated on 969 samples with 19 satellite-derived environmental variables. In the sparsity-ablation run at *k* = 13, it did not outperform linear CCA on CC1 (+15.8%, *p* = 0.201) but improved CC2 (+57.4%, *q* = 0.012) and CC3 (+65.9%, *q* = 0.006; Benjamini–Hochberg FDR across three components; Figure 2G–H); the primary 10-fold comparison on the same 13-domain subset found no difference (0.414 vs. 0.395 fold-averaged across CC1–CC3; *p* = 0.769). Thus the dominant coupling was largely linear in this analysis, whereas secondary variates showed exploratory evidence of nonlinear structure.

The 13 highest-weighted domains on CC1 are functionally convergent: twelve of the 13 encode transposable element (TE) or TE-associated machinery spanning the complete retrotransposon replication cycle: reverse transcriptase (PF00078, RVT_1; CC1 weight 0.113), three RNase H domains (PF17917, RT_RNaseH; PF17919, RT_RNaseH_2; PF00075, RNase_H), integrase core and zinc-binding domains (PF00665, rve; PF17921, Integrase_H2C2), Pao retrotransposon peptidase (PF05380, Peptidase_A17; CC1 weight 0.110), DDE transposase (PF01609, DDE_Tnp_1), Helitron rolling-circle transposase (PF14214, Helitron_like_N), retrotransposon endonuclease (PF14529, Exo_endo_phos_2), and two domains of unknown function found in retrotransposon polyproteins and eukaryotic transposon proteins (PF18701, DUF5641; PF20209, DUF6570). The sole non-TE domain in the top 13 was PF24664 (CC1 weight 0.076). The six highest-weighted TE domains (PF00078, PF17917, PF17919, PF00665, PF17921, PF05380) appeared in all 10 cross-validation folds, confirming robust selection. This TE enrichment independently converges with the supervised forward model finding that 11 of 18 most environment-predictable domains are mobile genetic elements (below), indicating that two methodologically independent frameworks—unsupervised sparse CCA and supervised XGBoost—identify the same functional signal.

The dominance of TE domains in environment-coupled features has a plausible biological basis in dinoflagellate genome architecture. Dinoflagellates possess among the largest known eukaryotic genomes (1–250 Gb),^62^ with extensive LTR retrotransposon content documented in *Symbiodinium minutum* ^9^ and reviewed by Wisecaver and Hackett.^62^ Because metagenomic assemblies from dinoflagellate-rich waters inherit the TE-heavy composition of their source genomes, TE domain abundance may serve as a compositional proxy for dinoflagellate community fraction. This interpretation corresponds to hypothesis (i) of the main-text Discussion—taxonomic covariation—and suggests that the environment– TE coupling captured by CCA reflects, at least in part, the spatial distribution of dinoflagellate-dominated communities along oceanographic gradients. A direct test of this proxy hypothesis—lineage-resolved partial correlations controlling for the five myzozoan Form II lineage counts (four dinoflagellate lineages plus Chromerida)—is presented below and indicates that TE–environment coupling persists after taxonomic correction, implying a component of within-lineage TE variation beyond pure taxonomic sorting.

The environmental features showing strongest nonlinearity in KAN spline activations—remote sensing reflectance at 443 nm (*R*^2^ = 0.28), 488 nm (*R*^2^ = 0.32), and normalized fluorescence line height (nflh; *R*^2^ = 0.36)—overlap with satellite bands used in operational harmful algal bloom (HAB) monitoring.^63,64^ The nonlinear structure of these features in CC2–CC3 in the *k* = 13 ablation run (CC2 +57.4%, *q* = 0.012; CC3 +65.9%, *q* = 0.006) but not CC1 (+15.8%, *p* = 0.201) is consistent with a two-regime coupling architecture: the leading canonical variate captures a broad, linear temperature– bathymetry gradient, while secondary variates capture threshold-like community transitions. However, several caveats apply: (i) the dataset contains no confirmed HAB events, as TARA Oceans was not designed to capture bloom dynamics; (ii) these spectral bands respond to many non-bloom processes including chromophoric dissolved organic matter (CDOM) absorption, sediment resuspension, and routine phytoplankton succession; and (iii) whether dinoflagellate TE abundance tracks bloom propensity or merely community composition cannot be resolved from cross-sectional data. We therefore treat the HAB-band overlap as a hypothesis-generating observation rather than evidence of bloom-specific coupling. The disappearance of KAN-CCA’s advantage at high dimensionality (*k* = 500; CC2: *−*42.2%, *p* = 0.012) confirms that the nonlinearity is localized to the sparse TE-dominated feature set rather than being a general property of domain–environment associations. Learned B-spline activation functions for these features are shown in Figure 2I–T.

#### Extended biological interpretation of environment-predictive domains

The following domain-level interpretations illustrate how the analytical framework recovers biologically coherent signals from SHAP-ranked environment-predictive domains. Table S9 summarizes all interpreted domains; three cases with the most distinctive biological signatures are discussed in detail below.

##### Ice-binding protein (PF11999) and cold adaptation

PF11999 is the single strongest individual predictor of sea surface temperature (SHAP importance 0.207 for SST_max_). Ice-binding proteins containing the DUF3494 domain inhibit ice recrystallization in polar diatoms;^5^ in *Fragilariopsis cylindrus*, IBP genes form a multigene family acquired via horizontal gene transfer from bacteria.^65–67^ SHAP importance is concentrated on temperature variables with negligible importance for non-thermal variables. PF11999 falls in Figure 3 cluster C2 (30 domains; dominant association Rrs555+) and is not among the 716 domains clustered in Figure 4. The metagenome-only validation confirmed *ρ* = *−*0.321 (*p* = 5.86 *×* 10^−20^; *n* = 773), demonstrating that this association is not driven by cultured reference genome inclusion.

##### dUTPase (PF00692) and PIF1 helicase (PF05970): genome maintenance at depth

dUTPase and PIF1 co-occur among depth-predictive features, implicating genome maintenance as a depth-structured functional axis. dUTPase hydrolyzes dUTP to prevent uracil misincorporation,^19^ and its depth enrichment may reflect reduced photoenzymatic repair at depth^68^ and elevated replication error rates under hydrostatic pressure.^42^ PIF1 is enriched in DNA repair (GO:0006281; *p* = 0.004 among temperature-associated domains) and DNA helicase activity (GO:0003678; FDR = 0.036) among environment-predictable domains. The two domains diverge after copy-number normalization: dUTPase drops from rank 2 to outside the top 20, whereas PIF1 retains its ranking, indicating that absolute gene dosage drives dUTPase’s depth association while PIF1 reflects relative compositional enrichment. After controlling for dominant RuBisCO lineage identity (seven dominant-lineage groups with *≥* 5 samples among the ten Form I lineages), PIF1’s partial correlation retains 76.9% of the zero-order signal (*ρ*_partial_ = *−*0.388, *p* = 1.67 *×* 10^−30^, *n* = 810).

##### PEPCK (PF17297) and light-responsive carbon metabolism

PEPCK is near-exclusively associated with normalized fluorescence line height (98.6% of total SHAP importance from NFLH; 9 of 14 variables have exactly zero importance). In marine diatoms, PEPCK shows diel periodicity, peaking during the dark phase and contributing to light-independent carbon fixation via *β*-carboxylation.^18^ Koley et al.^69^ reported that PEPCK is not used under autotrophic conditions in *Chlamydomonas reinhardtii* but may become active during the dark phase. The near-exclusive association with fluorescence—rather than chlorophyll concentration or temperature—is consistent with PEPCK modulating carbon flux during transitions in photosynthetic activity rather than tracking phytoplankton biomass.

The predictability hierarchy—TE domains at the top (*R*^2^ = 0.31–0.59), adaptive-function domains (Ice_binding, PIF1, PEPCK) in the middle, housekeeping domains (Myosin_head, ASC; *R*^2^ = 0.27) at the bottom—recapitulates functional expectations and serves as an internal calibration for the forward modeling framework. The 2.2-fold *R*^2^ difference between the most predictable TE domain and the least predictable housekeeping domains (0.59 vs. 0.27) quantifies this separation.

#### Domain-level biological interpretation

##### Domain-by-domain forward model interpretation

In the forward direction—can environmental conditions predict which protein domains will be abundant?—the 18 most predictable domains achieved median *R*^2^ = 0.40 (IQR: 0.36–0.42; *n* = 1,810 samples with both PFAM profiles and environmental metadata, including metagenomes and transcriptomes; Table S2), with 16 of 18 exceeding *R*^2^ *>* 0.3. The top-ranked domain was DUF6570 (PF20209; *R*^2^ = 0.59), a domain of unknown function found in eukaryotic transposon proteins, followed by Helitron_like_N (PF14214; *R*^2^ = 0.49, rolling-circle DNA transposase), Phage_integrase (PF00589; *R*^2^ = 0.46), and PIF1 helicase (PF05970; *R*^2^ = 0.46, DNA repair). Mobile genetic element domains—transposases, integrases, and reverse transcriptases— constituted 11 of the 18 most predictable domains (Table S2), consistent with the SHAP-based transposase enrichment described above (GO enrichment and per-environment-group domain interpretation). Among the least predictable of the 18 were Myosin_head (PF00063; *R*^2^ = 0.27) and the amiloride-sensitive sodium channel (PF00858; *R*^2^ = 0.27)—housekeeping domains with broadly conserved cellular functions that may vary less systematically with oceanographic gradients. Ice_binding (PF11999; *R*^2^ = 0.42) showed the highest cross-fold variance (SD = 0.21 vs. mean SD = 0.12), indicating spatially heterogeneous cold-adaptation signal that is strong in some ocean regions and weak in others. The predictability hierarchy is biologically informative: TE domains (DUF6570, Helitron_like_N, Phage_integrase) occupy the top ranks, consistent with mobile element load tracking environmental gradients (see Discussion), while non-TE domains in the top 18 implicate specific adaptive axes— Ice_binding reflects polar thermal adaptation, PIF1 helicase implicates DNA repair under UV or oxidative stress, and Myosin_head may track cell-size variation along productivity gradients. The contrast between highly predictable mobile element domains and weakly predictable housekeeping domains (Myosin_head *R*^2^ = 0.27, sodium channel *R*^2^ = 0.27) suggests that environment-responsive and conserved genomic compartments are functionally distinct. Calibration plots for the five most predictable non-phage domains (Figure 5A–E) confirm that predictions track observed abundances along the identity line, with characteristic mean regression; Figure 5F shows FTR1 (PF03239) from the separate five-fold screen.

#### GO enrichment analysis

GO enrichment analysis^39^ (5,182 PFAMs mapped to 9,842 GO associations via Pfam2GO^40^) confirmed that environment-predictable domains are functionally coherent. Among the 22 domains with held-out *R*^2^ *>* 0.3 in the top-100 single-split forward screen, the seven FDR-significant terms (Figure S4A) (*q <* 0.05) spanned RNA-DNA hybrid ribonuclease activity (GO:0004523), cytoskeletal motor activity (GO:0003774, GO:0016459), DNA helicase activity (GO:0003678), ion transport (GO:0006811), DNA integration (GO:0015074), and protein binding (GO:0005515)—implicating nucleic acid processing, cytoskeletal dynamics, and ion homeostasis as the functional axes most responsive to environment. Per-environment-group analysis revealed distinct signatures: temperature-associated domains were enriched for DNA repair (GO:0006281, fold enrichment 9.3*×*; raw *p* = 0.004, FDR = 0.14) and chromatin structure, depth-associated domains for lyase activity and membrane processes, and light-associated domains for photosystem II oxygen evolution (GO:0009654, 32.0*×*; raw *p* = 0.0016, FDR = 0.074) and photosynthetic electron transport. These enrichments indicate that the XGBoost forward models capture biologically interpretable environment–function relationships. Excluding all 23 transposase-family domains from the reverse models left performance nearly unchanged (bathymetry *R*^2^ 0.491*→*0.484; SST 0.608*→*0.611; Figure 3D), so the non-transposase features alone carry the reverse-model signal (see GO enrichment and per-environment-group domain interpretation, above).

##### Lineage-resolved transposase analysis

The TE-as-taxonomic-proxy hypothesis—that TE domain abundance tracks dinoflagellate community fraction rather than independent environmental coupling— predicts that TE–environment correlations should collapse after controlling for lineage composition. Three analyses tested this prediction (Methods). No TE–lineage pair exceeded *|ρ| >* 0.5 (maximum: rve *×* Prorocentrales, *ρ* = 0.38); TE domains showed significantly elevated variance within Form II-positive samples (Mann–Whitney *p* = 3.4 *×* 10^−4^); and partial correlations controlling for the five myzozoan Form II lineage counts (four dinoflagellate lineages plus Chromerida; *n* = 1,438) retained all 44 TE–environment associations after Bonferroni correction (mean 111% of zero-order *ρ* for SST). These results indicate that TE–environment coupling is not explained by myzozoan community composition at the lineage level, consistent with a component of within-lineage TE variation alongside taxonomic covariation.

#### Robustness and sensitivity analyses

##### Metagenome-only XGBoost and CCA analysis (data source confounding)

To directly address whether the combination of metagenomes, transcriptomes, and reference genomes inflates the reported coupling signals, we restricted both XGBoost SST prediction and CCA to the TARA Oceans metagenome subset (*n* = 773 with valid SST for XGBoost; *n* = 772 for CCA). Under 10-fold spatial block CV (2° grid, 123 blocks), the metagenome-only XGBoost achieved*R*^2^ = 0.453 for sst_mean_c and *R*^2^ = 0.435 for modis_sst_mean_c—*higher* than the full-dataset values of *R*^2^ = 0.388 (*n* = 1,165) and 0.383 (*n* = 1,279), respectively. CCA on the metagenome-only subset yielded CC1 = 0.892, exceeding the full-dataset CC1 of 0.816 (*n* = 1,809). In these analyses, restricting the data source did not weaken the coupling signal, arguing against inflation driven solely by mixed source types.

##### PERMANOVA batch-effect analysis

PERMANOVA on Bray-Curtis distances (999 permutations; *n* = 2,042) confirmed that data source explains a significant fraction of PFAM compositional variation: coarse data source (metagenome/ transcriptome/reference) *R*^2^ = 0.124 (pseudo-*F* = 144.0, *p* = 0.001); fine-grained dataset (9 levels) *R*^2^ = 0.218 (pseudo-*F* = 71.0, *p* = 0.001). However, within the SST-available subset (*n* = 1,165), SST quartiles explained only *R*^2^ = 0.039 (pseudo-*F* = 15.7, *p* = 0.001) via PERMANOVA—consistent with XGBoost capturing nonlinear variance that group-means-based PERMANOVA cannot detect (*R*^2^ = 0.38–0.45). The Mantel test between PFAM Bray-Curtis and SST Euclidean distances was weak (*r* = 0.010, *p* = 0.466), indicating that pairwise sample-level distance structure is not organized by SST alone. The higher coupling *R*^2^ in the metagenome-only analysis is consistent with source heterogeneity adding noise rather than generating the signal.

##### Latitude baseline for all flagship targets

To quantify the unique predictive contribution of PFAM domain composition beyond geographic position, three XGBoost models were trained under 10-fold spatial block CV (2° grid) for each of four flagship targets: (1) latitude-only (absolute latitude as sole feature); (2) PFAM-only (CLR + PCA100); (3) PFAM + latitude.

| Target | *n* | $R^2(\text{lat})$ | $R^2(\text{PFAM})$ | $R^2(\text{combined})$ | partial $R^2(\text{PFAM}\vert \text{lat})$ | partial $R^2(\text{lat}\vert \text{PFAM})$ |
| --- | --- | --- | --- | --- | --- | --- |
| SST | 1,279 | 0.796 | 0.375 | 0.852 | 0.273 | 0.763 |
| Bathymetry | 1,523 | -0.321 | 0.400 | 0.411 | 0.554 | 0.020 |
| Nitrate | 1,064 | 0.670 | 0.302 | 0.741 | 0.214 | 0.629 |
| Dissolved O<sub>2</sub> | 1,064 | 0.838 | 0.393 | 0.873 | 0.215 | 0.790 |

The targets partition into two regimes. SST, nitrate, and dissolved oxygen are latitude-dependent (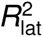 = 0.67–0.84), so latitude dominates these predictions (partial 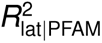 = 0.63–0.79), while PFAM adds 21–27% unique variance (partial 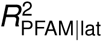 = 0.21–0.27). Bathymetry is latitude-independent (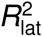 = *−*0.32): latitude is uninformative, and PFAM carries the predictive signal (partial 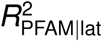 = 0.55; partial 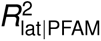 = 0.02). The distinction is physically interpretable: SST, surface nitrate, and dissolved oxygen all covary with latitude via solar heating and meridional overturning, whereas bathymetry depends on tectonic and depositional processes unrelated to latitude. PFAM contributes unique variance across all four flagship targets, and the contribution is largest for the one target (bathymetry) where latitude provides no baseline.

##### Spatial block size sensitivity

To assess robustness to spatial autocorrelation at longer correlation lengths, we re-ran XGBoost SST prediction (modis_sst_mean_c) under three spatial block sizes (*n* = 1,279): 2° (*R*^2^ = 0.375, 301 blocks; this rerun used a different, equal-size fold assignment from the primary spatial block CV, which gave *R*^2^ = 0.383), 5° (*R*^2^ = 0.365, 210 blocks), and 10° (*R*^2^ = 0.344, 136 blocks). The decline from 2° to 10° is 8.3% relative, with substantial signal retained at all scales. Fold-level variability increased with larger blocks (IQR: [0.29, 0.44] at 2° vs. [0.17, 0.45] at 10°), reflecting more uneven fold sizes, but the overall *R*^2^ remained well above zero at all scales.

##### CCA PCA component sensitivity

CCA was recomputed with 50, 100 (baseline), 150, and 200 PCA components to assess sensitivity to the dimensionality reduction threshold (*n* = 1,809):

| PCA components | Variance retained | CC1 | CC2 | CC3 |
| --- | --- | --- | --- | --- |
| 50 | 72.3% | 0.789 | 0.641 | 0.614 |
| 100 (baseline) | 76.0% | 0.816 | 0.719 | 0.682 |
| 150 | 78.5% | 0.842 | 0.766 | 0.741 |
| 200 | 80.4% | 0.850 | 0.789 | 0.776 |

CC1 is stable across all tested thresholds (0.789–0.850), monotonically increasing with additional components. The 100-component baseline retains 76% of variance and represents a conservative choice: adding components strengthens rather than weakens coupling. All canonical correlations increase monotonically, consistent with additional PCA dimensions capturing genuine signal rather than noise.

##### CCA component retention criterion

We retained 10 CCA components for representation after comparison with the standard row-permutation null (1,000 shuffles; Figure 2E): all 10 observed canonical correlations exceed the null distribution’s 99th percentile (0.496), and CC1–CC6 exceed the null maximum (0.582). Components CC2–CC10 are secondary or exploratory because the prespecified spatial-block permutation test was applied only to CC1; CC4–CC10 also show 36–47% cross-validation shrinkage (Figure S2G).

##### Spatial-block CCA permutation test

The standard row-shuffle permutation test destroys sample–environment pairings but preserves geographic neighbor redundancy within the PFAM matrix. Because oceanographic data carry spatial autocorrelation, this row-shuffle null is permissive—nearby samples share similar domain profiles regardless of the specific environmental pairing. To provide a conservative estimate that matches the spatial block CV design used for XGBoost validation, we recomputed the CCA CC1 permutation null under block shuffling within 2° grid cells (*n* = 1,809 samples across 434 blocks; 1,000 permutations). Block shuffling reassigns entire grid cells of PFAM rows to different environmental contexts, preserving within-block spatial autocorrelation under the null hypothesis.

Under spatial-block shuffling, the null CC1 distribution has mean = 0.639, SD = 0.024, and maximum = 0.773. The observed CC1 of 0.816 exceeds all 1,000 block-permuted values (*z* = 7.3, permutation *p <* 0.001; 0/1,000 permutations *≥* observed). Under the standard row shuffle run alongside it, the null has mean = 0.364, SD = 0.028, maximum = 0.572 (*z* = 16.2; Table S5, row 5); the separately seeded 1,000-permutation row-shuffle null plotted in Figure 2E has maximum 0.582. The block-shuffle null mean is 75% higher than the row-shuffle null mean (0.639 vs. 0.364), confirming that spatial autocorrelation inflates the row-shuffle null. The effect size under the conservative block-shuffle test (*z* = 7.3) is approximately half that of the row shuffle (*z* = 16.2), but the observed CC1 remains well above the block-shuffle null maximum, and no permutation exceeds the observed value.

##### Environmental variable collinearity

Of the 37 environmental variables, several groups—temperature variants and remote-sensing reflectance (Rrs) bands—introduce collinearity. Variance inflation factor (VIF) analysis on the 23 GEE-derived variables with *≤*50% missing data (*n* = 1,101 complete-case samples) identified 16 variables with VIF *>* 10 (Table S8). Three SST variables (sst_max, sst_min, sst_range) exhibit VIF = *∞*, indicating exact linear dependence within the SST group; the maximum pairwise *|r |* among the five temperature variables is 0.996 (sst_mean vs. MODIS sst_mean). Rrs bands show similarly extreme collinearity: rrs_667 and rrs_678 reach VIF *>* 21,000 with pairwise *|r |* = 0.998, and 15 variable pairs exceed *|r | >* 0.9 overall. Seven variables retain VIF *<* 10: chlorophyll (mean, max, min), POC mean, bathymetry, solar radiation, and distance to coast.

This collinearity does not affect XGBoost predictions (tree-based models are invariant to monotonic transformations and collinear features) but can destabilize CCA loadings for individual variables. We mitigate this through PCA dimensionality reduction prior to CCA (retaining 76% of variance in 100 components), which implicitly decorrelates the input space. SHAP importance is reported per environmental category (temperature, bathymetry, productivity, ocean color) rather than per individual variable where collinearity is a concern.

The VIF analysis covers 23 of the 32 GEE-derived variables after excluding columns with *>*50% missing values. The five WOA23 variables (nitrate, phosphate, silicate, oxygen, and mixed layer depth) were excluded from this VIF calculation; they are included where stated in the nutrient-augmented analyses.

##### CLR versus raw-count top-domain comparison

To quantify the impact of compositional normalization on domain-level rankings, we compared the top 50 domains by maximum *|ρ|* between raw-count and CLR-transformed Spearman correlations (*n* = 13,335 domains tested). This comparison used the AlphaEarth subset (*n* = 995 samples; 13,335 domains *×* 64 AEF dimensions; 853,423 associations tested) and is distinct from the primary 11,898-domain screen (279,575 significant associations). At the association level, 279,834 raw and 268,909 CLR associations were FDR-significant (*<* 0.05), with 127,664 significant under both; overall, 65.6% of the 853,423 associations had the same significance status under both normalizations (both significant or both non-significant). At the domain level, the top-50 domains ranked by max *|ρ|* showed Jaccard overlap of 0.205; ranked by number of significant associations, Jaccard was 0.235. Domain-level rank correlation between raw and CLR max *|ρ|* was *ρ* = 0.017 (*p* = 0.045). These results confirm that while the overall number of significant associations is comparable between normalizations, the *identity* of top-ranked domains differs substantially. Both perspectives are presented in Data S1; the manuscript uses CLR-transformed inputs for all quantitative modeling.

##### Prevalence-matched Pfam-dark-family versus Pfam-domain coupling

To control for the possibility that Pfam-dark-family–environment coupling enrichment reflects prevalence differences rather than functional divergence, we performed a bin-matched comparison (20 prevalence bins) between Pfam-dark families (*n* = 31,257; correlations over the 2,044-sample family matrix with per-variable complete cases) and Pfam domains (*n* = 9,611; 1,809-sample Pfam matrix), using 18 environmental variables. This comparison does not use the matched sample set of the primary analysis, and its Pfam-domain baseline differs accordingly. Unmatched median *|ρ|*: Pfam-dark families = 0.179, Pfam domains = 0.041 (4.3-fold). After prevalence matching (*n* = 6,488 per group): Pfam-dark families = 0.121, Pfam domains = 0.040 (3.0-fold; Mann–Whitney *p <* 10^−300^). The 2.29-fold enrichment reported in the manuscript (primary matched design: same 1,523 samples and 18 variables; median *|ρ|* = 0.157 vs. 0.068; Table S5, F12) is therefore conservative with respect to prevalence; after further prevalence matching, the enrichment increases to 3.0-fold.

##### Leave-one-basin-out cross-validation

Leave-one-basin-out (LOBO) CV trains on 6 basins and tests on the held-out basin, cycling through all 7 basins (Arctic, Atlantic, Indian, Mediterranean, Pacific, Red Sea, Southern) to produce a distribution of out-of-basin *R*^2^ values (Table S6). LOBO XGBoost models used *n*_estimators_ = 200, max depth = 4, and learning rate = 0.05 (CLR-transformed PFAM features, pseudocount = 0.5); these settings differ from the 10-fold spatial block CV models (max depth = 6, learning rate = 0.1). The SST target is MODIS SST. We summarize performance both across equally weighted basin-level scores and across all pooled held-out predictions.

**Bathymetry** (*n* = 1,523): Pooled predictions across all held-out samples yield *R*^2^ = 0.309 (MAE = 1,326 m). Three of 7 basin-level scores are positive: Pacific (*R*^2^ = 0.42, MAE = 1,246 m, *n* = 581), Atlantic (*R*^2^ = 0.30, MAE = 1,419 m, *n* = 543), and Indian (*R*^2^ = 0.28, MAE = 1,307 m, *n* = 169). These basins contain 1,293 of 1,523 evaluated samples (84.9%). The equal-basin median is *R*^2^ = *−*0.06 [IQR: *−*2.13–0.29], reflecting poor transfer in the Mediterranean (*R*^2^ = *−*1.88, *n* = 131), Arctic (*R*^2^ = *−*2.38, *n* = 34), Red Sea (*R*^2^ = *−*80.1, *n* = 19), and Southern Ocean (*R*^2^ = *−*0.06, *n* = 46). Thus, the bathymetry–domain relationship transfers across the bulk of the represented, data-rich open-ocean sample domain but is not uniform across basins.

**SST** (*n* = 1,279): Pooled held-out predictions yield *R*^2^ = 0.239 (MAE = 5.18°C), but only the Pacific holdout has positive basin-level performance (*R*^2^ = 0.12, MAE = 4.27°C, *n* = 486). All other basins—including the large Atlantic (*R*^2^ = *−*0.0005, *n* = 428) and Indian (*R*^2^ = *−*8.35, *n* = 148) test sets—produce *R*^2^ *<* 0; the equal-basin median is *−*8.35 [IQR: *−*32.6–*−*4.07]. The positive pooled score captures performance across the full global SST range, whereas the basin-level scores test improvement over each held-out basin’s own mean. SST–domain mappings therefore retain global-scale signal but do not transfer consistently within individual basins. Spatial block CV (MODIS SST *R*^2^ = 0.38, *n* = 1,279) quantifies coupling within geographically proximate samples; LOBO defines the broader applicability domain.

##### Within-TARA 20–180 µm size-fraction sensitivity analysis

To test whether the headline domain–environment coupling holds within a single algal-dominated size fraction, we restricted to TARA metagenome assemblies from the 20–180 µm fraction (*n* = 150 assemblies; 90 spatial blocks on a 2° grid). Size-fraction assignments were derived from the MGnify assembly metadata mapping (Table S7). CLR-transformed PFAM abundances (pseudocount = 0.5; 8,378 domains passing 5% prevalence) were reduced by PCA, and 20 components (55.4% variance retained) entered the CCA; XGBoost used CLR-transformed features directly.

**XGBoost reverse SST prediction** (10-fold spatial block CV, 2° blocks): *R*^2^ = 0.29 for GEE mean SST (*n* = 149 with SST data), compared to *R*^2^ = 0.39 for the same target in the full dataset (*n* = 1,165). The 24% reduction in *R*^2^ (0.293 vs. 0.388) accompanies a 7.8-fold reduction in sample size (MODIS SST: 0.296 vs. 0.383, a 23% reduction, with an 8.6-fold reduction from *n* = 1,279); with 15 samples per fold on average, fold-level *R*^2^ ranged from *−*0.07 to 0.56 (median = 0.33), indicating that the variance is driven by small fold sizes rather than absent signal.

**CCA CC1** (row-shuffle permutation, 1,000 iterations): observed CC1 = 0.917 exceeds the full-dataset CC1 (= 0.816). Permutation null: mean = 0.703, SD = 0.059, max = 0.910; *p* = 0.001, *Z* = 3.62. The increase in CC1 within a taxonomically more homogeneous subset is consistent with reduced inter-fraction compositional noise.

#### VICReg joint embedding analysis

As an alternative to the XGBoost bidirectional framework, we evaluated a self-supervised joint embedding approach using VICReg (Variance-Invariance-Covariance Regularization)^70^ to align environmental and PFAM representations in a shared latent space without requiring labeled targets during the alignment phase.

##### Architecture and training

The VICReg dual-encoder consists of two independent multilayer perceptron (MLP) encoders—one for 24 environmental variables (satellite-derived SST, chlorophyll-*a*, ocean color bands, and AlphaEarth embedding dimensions) and one for 20 PFAM module scores (PCA-reduced from CLR-transformed domain profiles)—that project into a shared 32-dimensional latent space. Each encoder is a two-layer MLP (*∼*53K–64K parameters; dropout = 0.3). The VICReg loss comprises three terms: invariance (mean squared error between paired environmental and PFAM embeddings; weight = 25.0), variance (hinge loss ensuring each latent dimension maintains standard deviation *≥* 1; weight = 25.0), and covariance (penalizing off-diagonal covariance to decorrelate latent dimensions; weight = 1.0). Training used 1,810 stations with GPS-matched environmental and PFAM data; the productivity prediction subset comprised 1,151 samples with complete chlorophyll-*a*, POC, and NFLH measurements. A linear prediction head was trained on concatenated environmental and PFAM embeddings to predict each productivity target.

##### Cross-modal alignment succeeds

The trained embeddings show meaningful cross-modal alignment. Mean cosine similarity between paired environmental and PFAM embeddings reaches 0.61 (random pairing baseline: *∼*0.0), indicating that the two modalities are mapped to overlapping regions of the latent space. HDBSCAN clustering of the joint latent space recovers Longhurst biogeochemical provinces at ARI = 0.52 (NMI = 0.61), comparable to the ARI = 0.503 obtained from HDBSCAN functional biomes of domain abundances (*n* = 1,310 non-noise samples; Figure S4B). The embedding therefore captures genuine biogeographic structure rather than noise.

##### Predictive transfer fails under spatial block CV

Despite successful alignment, downstream productivity prediction fails to generalize across ocean basins. Results under two CV designs:

*6-fold leave-one-basin-out (LOBO).* The joint model (environment + PFAM embeddings) achieves POC *R*^2^ = 0.532 versus an environment-only XGBoost baseline of *R*^2^ = 0.422. The improvement (+0.110 *R*^2^) is not statistically significant (Cohen’s *d* = 0.026, *p* = 0.38 by paired permutation test), and performance is unstable across folds: two basins (Indian Ocean, South Pacific) show improvement while the remaining four show degradation or marginal change.

*9-fold basin-and-latitude spatial CV.* This design split the Atlantic and Pacific each into three latitude bands and used the Mediterranean, Indian Ocean plus Red Sea, and Arctic plus Southern Ocean as the remaining folds; it differs from the 10-fold 2° grid-cell spatial block CV used for the primary XGBoost models. The MLP prediction head produces negative *R*^2^ on held-out blocks across all three PFAM dimensionalities tested:

| PFAM dims | Mean POC $R^2$ | Worst fold $R^2$ | Worst fold basin |
| --- | --- | --- | --- |
| 20 | −2.05 | −7.83 | Mediterranean |
| 32 | −3.14 | −12.61 | Pacific-Mid |
| 64 | −4.22 | −18.44 | Mediterranean |

Negative *R*^2^ indicates predictions worse than predicting the training-set mean. The failures are concentrated on basins with distinctive environmental regimes: the Mediterranean (semi-enclosed, high salinity, oligotrophic) and mid-Pacific (vast open-ocean gyre with minimal continental influence). A sign test across all 14 target–dimensionality configurations (7 targets *×* 2 PFAM dim settings with complete results) finds 1 of 14 positive (*p* = 0.9999), confirming systematic rather than stochastic failure.

##### Diagnosis: architecture confound, not signal absence

Three lines of evidence indicate that the failure is architectural rather than reflecting absence of a domain–environment alignment signal:

1. The alignment metrics (cosine similarity 0.61, ARI 0.52) confirm that VICReg successfully learns a shared representation; the failure occurs only when this representation is used for out-of-basin prediction.
2. XGBoost achieves positive *R*^2^ under its 10-fold 2° spatial block CV design (MODIS SST*R*^2^ = 0.38, bathymetry *R*^2^ = 0.42; main text), demonstrating that the signal exists and is extractable by tree-based methods.
3. The catastrophic failures are basin-specific (Mediterranean, Pacific-Mid), consistent with distribution shift rather than global signal absence. These basins have environmental distributions that differ from the training distribution in ways that the MLP extrapolates poorly.

The dominant factor is that XGBoost’s axis-aligned tree partitioning handles distribution shift in small tabular datasets (*n ≈* 1,100) more effectively than shallow neural networks, which must learn smooth decision boundaries that can fail catastrophically when test distributions fall outside the training manifold.

##### Avenues for improvement

Possible improvements include basin-aware normalization, heavier augmentation during pre-training, and replacing the MLP prediction head with a tree-based decoder; architecture and checkpoints are deposited at Hugging Face^71^ for reproduction.

#### Dipeptide composition bias in the dark proteome

The first-order physicochemical profiling (Figure 6) establishes that dark proteins are shorter, more basic, more disordered, and of lower sequence complexity than annotated proteins. We extended this characterization to second-order composition—dipeptide observed/expected (O/E) ratios—to test whether the dark proteome also differs in nearest-neighbor amino acid usage patterns.

##### Dipeptide O/E analysis

For each of the three proteome fractions (dark, annotated, algal; 500,000 randomly subsampled sequences per class, seed = 42), we computed the 20 *×* 20 dipeptide O/E matrix, where O*/*E(*i*, *j*) = count(*i*, *j*)*/*[ *f* (*i*) *× f* (*j*) *× N*_dp_ ], with *f* (*i*) the single-amino-acid frequency derived from dipeptide marginals and *N*_dp_ the total dipeptide count. We quantified bias spread as the standard deviation (SD) of log_2_(O/E) across all 400 dipeptides and counted dipeptides with *|*log_2_(O*/*E)*| >* 0.5 (*>*1.41-fold deviation from expectation) as exhibiting bias.

The dark proteome shows wider dipeptide bias than the annotated proteome across every measure (Figure S6C–F). The SD of log_2_(O/E) is 0.251 for the dark proteome versus 0.163 for the annotated proteome—a 1.54-fold difference—with the algal proteome intermediate at 0.229 (Table S10). The dark proteome has 22 dipeptides exceeding the bias threshold (5.5% of 400) compared with 8 for the annotated proteome (2.0%) and 18 for the algal proteome (4.5%). The most over-represented dipeptides in the dark proteome are KK (O/E = 1.952, log_2_ = +0.965), YY (1.887, +0.916), and QQ (1.809, +0.855); the most under-represented are PM (0.698, *−*0.518), HM (0.723, *−*0.468), and EC (0.723, *−*0.467). The annotated proteome spans a narrower O/E range (0.713–1.757; maximum QQ, followed by PP at 1.682) than the dark proteome (0.698–1.952). The mean absolute log_2_(O/E) is 0.189 (dark) versus 0.116 (annotated), confirming that the dark proteome deviates from amino acid independence 1.63-fold more than annotated proteins on average.

##### LCR-masked control

Because the dark proteome has lower sequence complexity (Shannon entropy 3.87 vs. 4.05 bits; Figure 6), the wider dipeptide bias could in principle reflect repetitive low-complexity regions (LCRs) rather than genuine compositional skew. To control for this, we repeated the analysis after masking LCRs using a 12-residue sliding window with Shannon entropy *<* 1.5 bits—matching the manuscript’s LCR definition—replacing masked residues with X and excluding them from all frequency and dipeptide counts.

LCR masking removed 1.7% of dark residues, 1.0% of annotated residues, and 1.6% of algal residues (Figure S6G–J). The dark-to-annotated SD ratio after masking is 0.237*/*0.152 = 1.56-fold, essentially unchanged from the unmasked ratio of 1.54-fold. The number of biased dipeptides decreased modestly (dark: 22 *→* 17; annotated: 8 *→* 5; algal: 18 *→* 14), but the rank ordering and relative differences persisted. The wider dipeptide bias in the dark proteome is therefore not an artifact of low-complexity sequence.

##### Interpretation

The dark proteome’s wider dipeptide O/E spread—maintained after LCR masking—indicates compositional skew at the nearest-neighbor level that exceeds what is observed in functionally annotated proteins. This pattern is consistent with rapid lineage-specific divergence under weak purifying selection: proteins under relaxed constraint accumulate context-dependent amino acid biases (e.g., codon usage bias, mutational biases in AT-rich genomes) that structured proteins under selection would not tolerate. The hierarchy—dark *>* algal *>* annotated—parallels the first-order physicochemical hierarchy (Figure 6) and the GC-proxy gradient (GC-proxy +0.213 vs. +0.064), reinforcing the interpretation that the dark proteome occupies a distinct region of sequence space shaped by evolutionary rate rather than by functional constraint on specific dipeptide motifs.

#### Population-genomic evidence for relaxed purifying selection on the dark proteome

The environmental coupling enrichment and dipeptide bias of the dark proteome are both consistent with relaxed purifying selection—an expectation that can be tested directly with population-genomic data. If Pfam-dark genes experience weaker functional constraint, they should accumulate nonsynonymous polymorphisms at a higher rate relative to synonymous polymorphisms (*π*_N_*/π*_S_) than Pfam-annotated genes.

##### Data source

Flowers et al.^29^ sequenced 20 strains of *Chlamydomonas reinhardtii* (12 field-collected, 8 laboratory) and reported per-gene *π*_N_ and *π*_S_ for 17,290 nuclear gene models (JGI Phytozome v5.6). Data were obtained from Dryad (doi:10.5061/dryad.1n0g6, SupplementalDS3).

##### Dark/white partition

Each gene was classified as **dark** (no Pfam-A domain annotation) or **white** (*≥*1 Pfam-A domain) using UniProt InterPro/Pfam annotations for *C. reinhardtii*, mapped via NCBI GFF3 locus tags (GCF_000002595.2). Of the 17,290 DS3 genes, 11,631 (67.3%) mapped successfully: 4,395 dark and 7,236 white. Genes with *π*_S_ = 0 or total SNPs *<* 5 were excluded, yielding 4,085 dark and 7,038 white genes for analysis.

##### Results

Median *π*_N_*/π*_S_ is 2.30-fold higher in dark-proteome genes than in Pfam-annotated genes (0.277 vs. 0.121; Mann-Whitney *U* = 21,368,216, *p <* 10^−300^; two-sided; Table S11). The aggregate ratio 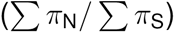, which avoids per-gene ratio noise, shows a consistent 1.84-fold difference.

Bootstrap resampling (10,000 replicates, seed = 42) yields a 95% CI on the median difference of [0.147, 0.164].

##### Controls

###### Gene-length control

Dark genes are modestly longer (median 69 vs. 60 total SNPs; Mann-Whitney *p* = 1.91 *×* 10^−5^). Resampling dark genes to match the white gene SNP-count distribution (IQR 30–119 SNPs) retains a 2.04-fold difference (*n*_matched_ = 1,739; *p* = 2.09 *×* 10^−190^), ruling out a length artifact.

###### Comparison with published values

Flowers et al. report *π*_N_*/π*_S_ = 0.41 for unannotated genes versus 0.16 for annotated genes (2.56-fold) using their own annotation categories. Our Pfam-based partition yields a consistent but slightly lower ratio (2.30-fold), as expected because Pfam annotation is more conservative than the original study’s combined annotation pipeline, shifting some weakly annotated genes from white to dark.

##### Interpretation

The elevated *π*_N_*/π*_S_ in Pfam-dark genes confirms that these genes experience weaker purifying selection than annotated genes. This is consistent with the dark proteome’s faster sequence divergence beyond homology detection thresholds and provides a population-genomic mechanism for the observational bias identified in the main text: genes under relaxed constraint diverge faster, fall below Pfam’s homology detection threshold, and accumulate in the unannotated fraction. The 2.30-fold *π*_N_*/π*_S_ enrichment in a single well-studied alga provides complementary within-species evolutionary evidence for the faster-diverging proteome whose cross-community environmental coupling was quantified in the 1,523-sample matched analysis.

Input data are from Dryad (doi:10.5061/dryad.1n0g6; Flowers et al. 2015); per-gene values and analysis scripts are deposited in Data S9.

##### Cross-taxonomic validation

To test whether relaxed purifying selection on Pfam-dark genes is a general pattern rather than a *Chlamydomonas*-specific finding, we applied the same dark/white partition to three additional species spanning *∼*1.5 Gyr of divergence (Table S12).

**Seminavis robusta** (Bacillariophyta, pennate diatom). Osuna-Cruz et al.^30^ sequenced 48 strains and reported per-gene *π*_N_ and *π*_S_ for 20,891 genes. Genes were partitioned by InterPro annotation status (integrating Pfam, PANTHER, CDD, and other member databases): 7,537 dark (no InterPro hit) and 13,205 white (*≥*1 hit). Dark genes show 1.46-fold higher median *π*_N_*/π*_S_ (0.171 vs. 0.117; Mann-Whitney *p* = 5.7 *×* 10^−160^; bootstrap 95% CI on fold: 1.42–1.51).

**Synechococcus** (Cyanobacteria). Tai and Palenik^31^ estimated per-gene *d*_N_*/d*_S_ for two reference genomes (CC9311 clade I, CC9902 clade IV) using environmental metagenomic reads from coastal California. We ran hmmsearch (Pfam-A, domain *E <* 10^−9^) on both NCBI proteomes to assign dark/white status. Dark genes show 1.88-fold (CC9311; median 0.179 vs. 0.095; *p* = 1.2 *×* 10^−68^; *n*_dark_ = 706, *n*_white_ = 1,698) and 1.80-fold (CC9902; 0.182 vs. 0.101; *p* = 1.1 *×* 10^−50^; *n*_dark_ = 549, *n*_white_ = 1,665) higher *d*_N_*/d*_S_. Combined: 1.84-fold (*p* = 3.3 *×* 10^−117^). Dark genes are also 66-fold enriched among positively selected genes (*d*_N_*/d*_S_ *>* 1) in CC9311 (Fisher OR = 66.0, *p* = 1.4 *×* 10^−25^). Note: *d*_N_*/d*_S_ measures between-lineage divergence, not within-species polymorphism; both metrics reflect selection but at different evolutionary timescales.

**Thalassiosira pseudonana** (Bacillariophyta, centric diatom). Koester et al.^32^ tested gene models (11,390 reported) across 7 strains for positive selection using PAML site models. We ran hmmsearch (Pfam-A, domain *E <* 10^−9^) on the 11,673 proteins of the NCBI GCF_000149405.2 proteome: 5,408 dark, 6,265 white. Among 809 Bonferroni-significant positively selected genes, the positive-selection incidence is 1.90-fold higher in dark than in white genes (9.3% vs. 4.9%; Fisher OR = 2.0, *p* = 4.9 *×* 10^−21^). Separately, the dark fraction among positively selected genes is 1.34-fold its genome-wide value (62.2% vs. 46.3%; bootstrap 95% CI on this enrichment fold: 1.27–1.42); this interval does not apply to the 1.90-fold incidence ratio.

Across green algae (*∼*1 Gyr divergence from diatoms), pennate and centric diatoms, and cyanobacteria (*∼*1.5–2 Gyr divergence from eukaryotes), the within-study contrasts consistently indicate altered selective constraint in Pfam-dark genes. Because *π*_N_*/π*_S_, *d*_N_*/d*_S_, and positive-selection incidence operate at different evolutionary scales, their fold contrasts were not pooled as a common evolutionary rate.

Per-gene data, provenance, and analysis scripts for all four species are deposited in Data S9.

#### Boltz-2 structure prediction of Pfam-dark protein-family representatives

To assess whether structurally coherent folds exist among the Pfam-dark protein families, we predicted three-dimensional structures for 3,600 representative sequences using Boltz-2 v2.2.1^33^ (MIT-licensed; run on NVIDIA GB10 GPU). Sequences were drawn from 10 rank bins (360 per bin, stratified by maximum *|ρ|* across environmental variables) spanning the full range of environmental coupling strengths.

Multiple sequence alignments were constructed from existing MMseqs2 30% identity cluster members (median 66 members per cluster, max 660) aligned with MAFFT, then projected onto the query sequence to meet Boltz-2’s fixed-length MSA requirement.

#### NOVEL_33041: case study of an uncharacterized Pfam-dark protein family

The tightest single Pfam-dark-family–environment coupling in the dataset (*ρ* = 0.566, bathymetry; *n* = 1,523) belongs to NOVEL_33041, a cluster representative derived from the *Dunaliella* ROIL 10x Genomics assembly. This family has no Pfam or integrated InterPro functional annotation. DIAMOND returned five weak or partial NCBI nr matches—four hypothetical proteins and one alignment to the non-reverse-transcriptase region of a reverse-transcriptase-labelled multidomain protein—but none assigned a credible function. NOVEL_33041 was not included in the locally retained Foldseek query set, so a PDB homology result is not available. Below we present the evidence assembled to evaluate whether it encodes a genuine, functional protein family.

##### Within-genome distribution

The *Dunaliella* ROIL assembly contains 66 copies of NOVEL_33041 on 66 distinct contigs, spanning all contig size classes: 20 copies on chromosome-scale scaffolds (*≥*100 predicted genes), 21 on sub-chromosomal scaffolds (10–99 genes), and 25 on short contigs (*<*10 genes). The largest scaffold carrying a copy contains 1,110 predicted genes. This dispersed distribution across all contig size classes argues against assembly fragmentation as the primary explanation for copy number.

##### Within-genome selection

Pairwise *d*_N_*/d*_S_ across all 66 copies (Nei-Gojobori method with Jukes-Cantor correction; 2,145 pairs from a 270-codon alignment back-translated onto the MAFFT protein alignment) yields the values summarized in Table S13.

This *ω* = 0.33 is comparable to conserved eukaryotic gene families (*ω* = 0.1–0.5) and is incompatible with pseudogenization. The high proportion of very-low-*d*_S_ pairs across different contigs (16.4%) is consistent with concerted evolution via gene conversion homogenizing the family. Three copies show elevated mean *ω* (0.81–0.96), suggesting possible subfunctionalization, but even the highest (0.96) remains consistent with relaxed purifying selection rather than confirmed positive selection. Protein-level diversity across all 66 copies is *π* = 0.038 per site (47 unique sequences; mean pairwise identity 96.0%, median 98.5%). IQ-TREE2 inference (Q.mammal+G4 model, 1,000 UFBoot replicates) recovers a structured phylogeny with a core clade of near-identical copies (including 17 protein-identical copies on 17 different contigs) and a divergent subclade at 74–85% identity, indicating the family is not uniformly young.

##### Expression evidence

NOVEL_33041 is detected in four independent MMETSP transcriptomes from cultured Chlamydomonadales: *Chlamydomonas leiostraca* (MMETSP1391, *E* = 1.3 *×* 10^−29^, bitscore 103.8), *Chlamydomonas* sp. (MMETSP1180, *E* = 2.4 *×* 10^−15^), *C. euryale* (MMETSP0063, *E* = 8.6 *×* 10^−14^), and *C. chlamydogama* (MMETSP1392, *E* = 2.5 *×* 10^−12^). All represent poly(A)-selected mRNA, providing direct evidence of transcription. Additionally, environmental read recruitment detects the family in 118 ocean metagenome samples with 2,348 cumulative hits.

##### Cross-phylum conservation

The family is detected across algal phyla spanning *∼*1.5 Gyr of divergence: Chlorophyta (Volvocales including *Dunaliella*, *Chlamydomonas*, *Volvox* ; Chlorellales; Sphaeropleales), Stramenopiles (*Pelagomonas calceolata*, *E* = 3.8 *×* 10^−31^), Haptophyta (*Tisochrysis lutea*, *E* = 4.0 *×* 10^−16^), Euglenozoa (*E* = 1.1 *×* 10^−21^), and Charophyta (*Klebsormidium nitens*, *E* = 5 *×* 10^−6^). A total of 118 assemblies across 118 unique samples carry the family.

##### DIAMOND BLASTp against NCBI nr

Only five hits at *E <* 10^−6^ were recovered from the full NCBI nr database (*∼*700 million sequences, June 2026). Four are annotated as hypothetical proteins: two from Volvocales green algae (*Chlamydomonas* sp. UWO 241, a psychrophilic Antarctic isolate; *Astrephomene gubernaculifera*), and two from freshwater metagenome-assembled genomes (99.6% identical to each other; likely the same gene binned into two MAGs). The fifth hit (MCA3444670.1, *Rhodobacter* MAG) is annotated as “reverse transcriptase family protein,” but this annotation derives from the N-terminal domain of a 1,091-aa multi-domain protein; the DIAMOND alignment maps exclusively to the C-terminal domain (*>*500 residues from the RT catalytic motif YADD at position 292), and InterProScan detects no RT or any other domain in NOVEL_33041. The RT annotation is spurious.

##### Compositional signature

The 198-aa representative has an unusual amino acid composition: 14 tryptophans (7.1%, 6.5*×* the Swiss-Prot average), 9 cysteines (4.5%, 3.3*×* enriched), an estimated net charge of +9.7 at pH 7 from BioPython `ProteinAnalysis`, and near-complete depletion of isoleucine (1 residue) and tyrosine (1 residue). Two cysteine motifs—CVC (positions 86–88) and CPC (positions 113–115)—are separated by a universally conserved HVFW tetrapeptide. The HVFW motif is 100% conserved across all 19 taxonomic representatives spanning five phyla; the CPC motif is conserved in 84%. ESMFold predicts a largely disordered structure (mean pLDDT 0.48), with a semi-structured core at residues 84–123 (mean pLDDT 0.64) anchored by the CVC/CPC disulfide pairs and Trp89 (pLDDT 0.76, the highest in the protein). ESMFold-predicted S*_γ_*–S*_γ_* distances between the CVC and CPC motifs (3.2–5.5 Å) are consistent with disulfide bonding in the folded core.

##### Subcellular localization

Heuristic analysis identifies a possible but atypical signal peptide (hydrophobic core at positions 15–22, displaced from the N-terminus by charged residues R5 and R8); no transmembrane domain is detected. The protein’s cysteine-rich motifs, tryptophan enrichment, cationic net charge, and genomic co-localization with an antifreeze type I protein (IPR000104) are consistent with a possible extracellular or surface-associated environmental-interface role, but localization remains untested. No known domain matches the combined CVC… HVFW… CPC + Trp-richness signature.

#### Size-fraction rationale and operational scope

TARA Oceans collected samples across multiple size fractions (0.8–5, 5–20, 20–180, 180–2000 µm), but analyses were not stratified by size fraction for the following reasons. First, microalgal cells undergo large size changes across life stages—vegetative cells, gametes, resting cysts, and colonial forms of a single species can span two or more filter fractions—so size fraction is an unreliable proxy for taxonomy at the species level, let alone at the protein domain level. Second, the dataset integrates ocean metagenomes (TARA plus Ocean Sampling Day), MMETSP transcriptomes, and cultured reference proteomes from NCBI GenBank/RefSeq, Phytozome, and algal culture collections (Table S1); only the TARA Oceans subset carries size-fraction metadata, and stratification would discard the OSD, transcriptome, and reference samples (1,167 of the 2,044-sample domain-analysis set are not TARA Oceans assemblies) while biasing results toward a single campaign. Third, this study operates at the protein domain level rather than the organism level: the same domain family appears across size fractions via shared gene families, making fraction-based partitioning less informative for domain–environment coupling than for organism–environment coupling. Fourth, LA^4^SR classifies sequences individually, removing the assumption that all proteins from a given size fraction belong to the same taxonomic group. Finally, PERMANOVA testing confirmed that data source identity explains only *R*^2^ = 0.12 of domain composition variance, and restricting to TARA Oceans metagenomes alone *increased* rather than decreased coupling strength (see Metagenome-only XGBoost and CCA analysis, above), indicating that multi-source heterogeneity attenuates rather than inflates the reported associations.

##### Operational scope: “algal” as a sequence-level, not taxonomic, designation

The “algal” label is an LLM-mediated sequence-level classification, not a taxonomic one. LA^4^SR assigns individual protein sequences to the algal fraction based on learned compositional features; it does not assign organisms, size fractions, or taxonomic identities. Non-photosynthetic marine protists that share evolutionary ancestry with algae—apicomplexans, plastid-loss dinoflagellates, choanoflagellates, kinetoplastids, kleptoplastic ciliates—retain genomic signatures of photosynthetic heritage and are correctly included in the algal-derived functional repertoire (Methods). The downstream domain–environment associations therefore reflect coupling between a sequence-defined functional space and the physical environment, independent of organismal taxonomy. The empirical evidence supports this framing: LA^4^SR recovers RuBisCO from 1,069 of 1,203 ocean metagenome assemblies (TARA Oceans and OSD; 88.9%), with TARA Oceans assemblies spanning all four collected size fractions, including the bacterially dominated 0.8–5 µm fraction (Results; Figure 1C–Q). Recovery of a chloroplast-encoded photosynthetic marker across size fractions that the textbook mapping assigns to bacteria (0.8–5 µm), small eukaryotes (5– 20 µm), and large eukaryotes (20–180 µm) is direct evidence that filter-fraction boundaries do not bound algal biology: marine algal cell sizes span more than three orders of magnitude across taxa and life stages,^72^ and much of this life-cycle size variation—gametes, resting cysts, colonial disaggregation— remains uncharacterized for the majority of marine algal species.^73^ The 20–180 µm fraction is therefore a TARA Oceans sampling choice, not a biological boundary.

#### Prospective validation opportunities

All validation in this study—spatial block CV and cross-basin holdout—operates within data collected by the same expeditions (TARA Oceans and OSD). Cross-campaign validation—applying the trained PFAM*→*SST model to metagenomes from an independent expedition—is the outstanding test of generalizability. Several public datasets are suited to this purpose: the Tara Pacific expedition^74^ provides coral reef metagenomes across Indo-Pacific gradients, the Malaspina 2010 circumnavigation^75^ sampled the deep ocean globally, and time-series stations (Hawaii Ocean Time-series,^76^ Bermuda Atlantic Time-series Study^77^) offer repeated sampling at fixed locations that would decouple spatial and temporal variation entirely. Each design tests a different facet of the environmental filtering hypothesis: Tara Pacific would assess whether domain–temperature coupling transfers across biomes and size fractions; Malaspina would test the depth–composition signal independently of the TARA track; and time-series stations would isolate seasonal environmental forcing from biogeographic structure. Retained predictive skill on independent campaigns (*R*^2^ *>* 0 for SST) would strengthen the case that domain–environment coupling reflects a general ecological pattern rather than expedition-specific confounding; complete failure would constrain the framework’s scope to within-campaign description rather than cross-campaign prediction.

#### Variance partitioning via distance-based redundancy analysis

Distance-based redundancy analysis (db-RDA) following Borcard et al.^78^ was performed on the 786-sample subset with complete environmental and geographic metadata. Euclidean distance on centered log-ratio (CLR)-transformed Pfam abundances—equivalent to Aitchison distance—was projected onto 174 principal coordinate axes capturing 80.0% of positive eigenvalue variance. Two predictor matrices were constructed: (1) an environmental matrix of 24 standardized Google Earth Engine and World Ocean Atlas 2023 (GEE+WOA23) variables; (2) a geographic matrix comprising latitude, longitude, and 18 distance-based Moran eigenvector maps (dbMEMs)^79^ capturing spatial autocorrelation at multiple scales. Variance was partitioned into four fractions: pure environmental [a], shared (spatially structured environmental) [b], pure geographic [c], and residual [d]. Conditional Freedman–Lane permutation tests (999 permutations) assessed the pure environmental and pure geographic fractions; an unconditional row-permutation test assessed the full model.

| Fraction | Adj. $R^2$ | % of explained | Permutation $p$ |
| --- | --- | --- | --- |
| Pure environmental [a] | 0.029 | 36.7% | 0.001 |
| Shared env\|geo [b] | 0.017 | 20.9% | — |
| Pure geographic [c] | 0.034 | 42.4% | 0.001 |
| Residual [d] | 0.920 | — | — |
| Environmental total [a+b] | 0.046 | 57.6% | — |
| Geographic total [b+c] | 0.050 | 63.3% | — |
| **Full model [a+b+c]** | **0.080** | **100%** | 0.001 |

The pure environmental and pure geographic fractions and the full model are significant at *p* = 0.001 (999 permutations). The total explained variance (adj. *R*^2^ = 0.080) is low in absolute terms given the high dimensionality of the response matrix (9,466 Pfam domains) and the ecological complexity of global ocean metagenomes. Pure geographic variance (adj. *R*^2^ = 0.034) exceeds pure environmental variance (adj. *R*^2^ = 0.029) by 1.15-fold, supporting an independent contribution from dispersal limitation or historical biogeographic barriers. The shared fraction (adj. *R*^2^ = 0.017, 20.9% of explained variance) captures spatially structured environmental variation. The combined environmental signal (pure + shared) accounts for 57.6% of explained variance, whereas pure geography accounts for 42.4%; this partition is consistent with contributions from both environmental and geographic structure.

Output deposited in Data S1.

#### Meta-analytical multiplicity

The study employs approximately 18 analytical frameworks (Spearman screening, CLR sensitivity, XGBoost forward/inverse modeling, SHAP decomposition, CCA/GCCA/sparse CCA/KAN-CCA, spatial block cross-validation, temporal hold-out, permutation null, TE enrichment analysis, dark proteome discovery, and structural validation). Although each addresses a distinct scientific question—association discovery, predictive validation, feature attribution, dimensionality reduction, robustness testing, and functional characterization—the meta-analytical multiplicity means that some individual findings could arise by chance across so many frameworks. The central conclusions do not rest on any single test: SST predictability emerges independently in XGBoost, CCA, and spatial block designs; TE enrichment appears in both SHAP attribution and forward prediction rankings; and the temporal hold-out attenuation relative to spatial block CV is a negative result consistent across all evaluated targets. This convergence across supervised, unsupervised, and sensitivity frameworks mitigates—though does not eliminate—the risk of framework-level false discovery.

#### Transposable element experimental evidence

Hypothesis (ii) of the main-text Discussion—stress-induced TE activation—is supported by direct experimental evidence in marine microalgae. In the diatom *Phaeodactylum tricornutum*, the LTR retrotransposon Blackbeard is transcriptionally activated under nitrate starvation via promoter hypomethylation,^17^ while a second retrotransposon, Surcouf, is specifically induced by thermal stress.^80^ Cold stress activates LTR retrotransposons in the diatom *Leptocylindrus aporus*, where TE-related transcripts are the single most enriched category under temperature downshift.^81^ Long-term ocean warming drives epigenetic release of TEs via loss of H3K27me3 in *Phaeodactylum*.^82^ In dinoflagellates, retrotransposon-mediated retroposition has reshaped *∼*23% of *Symbiodinium* genes, with bursts coinciding with periods of environmental transition.^83^ Critically, reverse transcriptase genes—the enzymatic core of retrotransposon replication—reach up to 13.5% of total gene abundance in large-size-fraction TARA Oceans metagenomes, with active transcription confirmed in matched metatranscriptomes.^16^ The stress–TE activation link is a eukaryote-wide phenomenon,^15,49^ but these marine-specific studies establish that the mechanism operates in the taxa and environments represented in our dataset.

The analytical framework recovers biologically coherent signals without supervision: cold-adaptation machinery predicts temperature, genome maintenance predicts depth, carbon-flux regulation predicts photosynthetic activity, and macronutrient cycling domains carry importance across multiple oceanographic axes. These associations are correlative, consistent with the cross-sectional design.

#### Harmful algal bloom spectral overlap

The satellite bands where KAN-CCA identifies significant nonlinear structure—rrs at 443 nm, 488 nm, and normalized fluorescence line height—overlap with bands used in operational harmful algal bloom (HAB) detection algorithms.^63,64^ However, several caveats preclude causal interpretation. First, the dataset contains no confirmed HAB events; the TARA Oceans sampling campaign was not designed to capture bloom dynamics. Second, these spectral bands respond to many non-HAB processes— including CDOM absorption, sediment resuspension, and routine phytoplankton community turnover— so nonlinear structure in these bands does not imply bloom-specific transitions. Third, the nonlinearity is concentrated in CC2–CC3, whose consensus domain features include TE domains (Helitron_like_N, DUF6570) alongside non-TE domains, but whether dinoflagellate TE abundance tracks bloom propensity or merely community composition is unresolved. The implied two-regime architecture—linear temperature–depth gradients on CC1, threshold-like community transitions on CC2–CC3—*could* reflect spectral signatures associated with shifts between dinoflagellate-dominated and other phytoplankton assemblages. The available cross-sectional data cannot resolve this question.

#### Limitations of the study

All domain–environment associations reflect co-occurrence in cross-sectional data; the bidirectional modeling framework establishes shared statistical structure but does not establish causal direction. AEF records were available for 995 of 1,810 GPS-mapped domain-analysis samples (55.0%): 969 contained no zero-filled dimensions and 26 were partially zero-filled during feature preparation. Missing records were concentrated in open-ocean locations, limiting generalizability to pelagic environments. Table S5 inventories 18 analytical frameworks and groups related analyses into seven claim families. Twelve frameworks have a documented primary *p*- or *q*-value; 9 of these 12 retain *p <* 0.05 after a conservative Bonferroni-18 correction. The remaining six frameworks report cross-validation, agreement, or sensitivity effects without assigning an undocumented null-hypothesis test.

The biological sampling campaigns (TARA Oceans, 2009–2012; OSD, 2014) predate the AlphaEarth annual satellite embeddings, which are available for 2017 onward (Earth Engine collection GOOGLE/SATELLIT all available annual layers combined with ImageCollection.mosaic() before extraction), so those environmental embeddings are not contemporaneous with biological sampling; temporal analysis of domain– environment coupling stability is reported in Supplemental Text and Figure S5.

Geographic bias toward the Atlantic and Pacific, with limited sampling in polar basins (Southern Ocean *n* = 50, Arctic *n* = 38), precludes robust inferences about domain–environment coupling under polar oceanographic conditions. Dissolved nutrient concentrations (nitrate, phosphate, silicate) and dissolved oxygen are derived from WOA23 climatological annual means at 1° resolution^84^ rather than *in situ* measurements concurrent with sample collection; this introduces spatial smoothing (mesoscale features *<*100 km are unresolved) and temporal mismatch (climatology vs. actual conditions at time of sampling). Mixed layer depth values are similarly climatological.^85^ These limitations are partially offset by the large sample size and global coverage, but nutrient-domain associations should be interpreted as reflecting large-scale biogeochemical gradients rather than local transient conditions. We tested whether incorporating these WOA23 nutrient fields into the CCA environmental matrix altered the joint embedding; because the climatological annual means lack the spatiotemporal specificity of the satellite-derived variables matched to actual sampling coordinates and dates, we retained the 1,809-sample CCA without WOA23 nutrient fields as the primary analysis and report the nutrient-augmented analyses (786 samples; 24 variables) separately for reference. Top-down controls (viral lysis, grazing, parasitism)^86^ are absent from the environmental characterization. Classification recall varies by lineage depending on representation in the training data (e.g., underrepresented or phylogenetically distant lineages show lower retention). PFAM annotation depends on database completeness. The Pfam-dark family HMMs were built from whole-protein 30% identity clusters with 80% bidirectional coverage; no independent domain boundaries were inferred. This threshold lies in the protein-family twilight zone^87^ where sequence similarity does not guarantee homology. The Pfam-dark-family enrichment is stable across a 10,000-fold range of hmmsearch E-value thresholds (10^−9^ to 10^−5^; 2.57- to 2.32-fold in the unmatched design, Table S5, F14; the primary matched 2.29-fold comparison is shown in Figure 6M), confirming detection-threshold robustness, but identity-threshold sensitivity at 20% and 40% has not been evaluated. The initial correlation discovery screen uses raw PFAM counts; the top-ranked domains differ between raw and CLR normalizations (Jaccard = 0.21 at the top-50 level; Data S1), with 34% of tested associations changing FDR-significance status under compositional correction. The top-18 interpreted domains are validated under spatial-block CV on CLR-transformed inputs, but the targets were selected in an earlier standard five-fold screen on an overlapping cohort; the reported spatial scores are therefore conditional on selection and the maximum may be optimistic. All modeling analyses use CLR-transformed inputs. Size-fraction metadata was not used to stratify analyses (see Size-fraction rationale and operational scope, above). The dataset contains no confirmed harmful algal bloom events, so the overlap between nonlinear KAN-CCA features and HAB-monitoring spectral bands remains observational.

## References

1. Field, C.B., Behrenfeld, M.J., Randerson, J.T., and Falkowski, P. (1998). Primary production of the biosphere: integrating terrestrial and oceanic components. Science 281, 237–240. doi: 10.1126/science.281.5374.237.

2. Gorelick, N., Hancher, M., Dixon, M., Ilyushchenko, S., Thau, D., and Moore, R. (2017). Google Earth Engine: Planetary-scale geospatial analysis for everyone. Remote Sensing of Environment 202, 18–27. doi: 10.1016/j.rse.2017.06.031.

3. Pesant, S., Not, F., Picheral, M., Kandels-Lewis, S., Le Bescot, N., Gorsky, G., Iudicone, D., Karsenti, E., Speich, S., Troublé, R., et al. (2015). Open science resources for the discovery and analysis of Tara Oceans data. Scientific Data 2, 150023. doi: 10.1038/sdata.2015.23.

4. Planes, S., Allemand, D., Agostini, S., Banaigs, B., Boissin, E., Boss, E., Bourdin, G., Bowler, C., Douville, E., Flores, J.M. et al. (2019). The Tara Pacific expedition—a panecosystemic approach of the “-omics” complexity of coral reef holobionts across the Pacific Ocean. PLOS Biology 17, e3000483. doi: 10.1371/journal.pbio.3000483.

5. Carradec, Q., Pelletier, E., Da Silva, C., Alberti, A., Seeleuthner, Y., Blanc-Mathieu, R., Lima-Mendez, G., Rocha, F., Tirichine, L., Labadie, K., et al. (2018). A global ocean atlas of eukaryotic genes. Nature Communications 9, 373. doi: 10.1038/s41467-017-02342-1.

6. de Vargas, C., Audic, S., Henry, N., Decelle, J., Mahé, F., Logares, R., Lara, E., Berney, C., Le Bescot, N., Probert, I., et al. (2015). Ocean plankton. Eukaryotic plankton diversity in the sunlit ocean. Science 348, 1261605. doi: 10.1126/science.1261605.

7. Rives, A., Meier, J., Sercu, T., Goyal, S., Lin, Z., Liu, J., Guo, D., Ott, M., Zitnick, C.L., Ma, J. et al. (2021). Biological structure and function emerge from scaling unsupervised learning to 250 million protein sequences. Proceedings of the National Academy of Sciences 118, e2016239118. doi: 10.1073/pnas.2016239118.

8. Lin, Z., Akin, H., Rao, R., Hie, B., Zhu, Z., Lu, W., Smetanin, N., Verkuil, R., Kabeli, O., Shmueli, Y. et al. (2023). Evolutionary-scale prediction of atomic-level protein structure with a language model. Science 379, 1123–1130. doi: 10.1126/science.ade2574.

9. Régimbeau, A., Aumont, O., Bowler, C., Guidi, L., Jackson, G.A., Karsenti, E., Memery, L., Tagliabue, A., and Eveillard, D. (2025). Unveiling the link between phytoplankton molecular physiology and biogeochemical cycling via genome-scale modeling. Science Advances 11, eadq3593. doi: 10.1126/sciadv.adq3593.

10. Vorobev, A., Dupouy, M., Carradec, Q., Delmont, T.O., Annamalé, A., Wincker, P., and Pelletier, E. (2020). Transcriptome reconstruction and functional analysis of eukaryotic marine plankton communities via high-throughput metagenomics and metatranscriptomics. Genome Research 30, 647–659. doi: 10.1101/gr.253070.119.

11. Violle, C., Navas, M.L., Vile, D., Kazakou, E., Fortunel, C., Hummel, I., and Garnier, E. (2007). Let the concept of trait be functional! Oikos 116, 882–892. doi: 10.1111/j.0030-1299.2007.15559.x.

12. Mistry, J., Chuguransky, S., Williams, L., Qureshi, M., Salazar, G.A., Sonnhammer, E.L., Tosatto, S.C., Paladin, L., Raj, S., Richardson, L.J. et al. (2021). Pfam: The protein families database in 2021. Nucleic Acids Research 49, D412–D419. doi: 10.1093/nar/gkaa913.

13. Brown, C.F., Kazmierski, M.R., Pasquarella, V.J., Rucklidge, W.J., Samsikova, M., Zhang, C., Shelhamer, E., Lahera, E., Wiles, O., Ilyushchenko, S., et al. (2025). AlphaEarth foundations: An embedding field model for accurate and efficient global mapping from sparse label data. doi: 10.48550/arXiv.2507.22291. arXiv:2507.22291.

14. Salazar, G., Paoli, L., Alberti, A., Huerta-Cepas, J., Ruscheweyh, H.J., Cuenca, M., Field, C.M., Coelho, L.P., Cruaud, C., Engelen, S. et al. (2019). Gene expression changes and community turnover differentially shape the global ocean metatranscriptome. Cell 179, 1068–1083. doi: 10.1016/j.cell.2019.10.014.

15. Sunagawa, S., Coelho, L.P., Chaffron, S., Kultima, J.R., Labadie, K., Salazar, G., Djahanschiri, B., Zeller, G., Mende, D.R., Alberti, A. et al. (2015). Structure and function of the global ocean microbiome. Science 348, 1261359. doi: 10.1126/science.1261359.

16. Nelson, D.R., Jaiswal, A.K., Ismail, N.S., Mystikou, A., and Salehi-Ashtiani, K. (2025). Panmicroalgal dark proteome mapping via interpretable deep learning and synthetic chimeras. Patterns 6, 101373. doi: 10.1016/j.patter.2025.101373.

17. Kopf, A., Bicak, M., Kottmann, R., Schnetzer, J., Kostadinov, I., Lehmann, K., Fernandez-Guerra, A., Jeanthon, C., Rahav, E., Ullrich, M. et al. (2015). The ocean sampling day consortium. GigaScience 4, 27. doi: 10.1186/s13742-015-0066-5.

18. Keeling, P.J., Burki, F., Wilcox, H.M., Allam, B., Allen, E.E., Amaral-Zettler, L.A., Armbrust, E.V., Archibald, J.M., Bharti, A.K., Bell, C.J. et al. (2014). The Marine Microbial Eukaryote Transcriptome Sequencing Project (MMETSP): Illuminating the functional diversity of eukaryotic life in the oceans through transcriptome sequencing. PLOS Biology 12, e1001889. doi: 10.1371/journal.pbio.1001889.

19. Eddy, S.R. (2011). Accelerated profile HMM searches. PLoS Computational Biology 7, e1002195. doi: 10.1371/journal.pcbi.1002195.

20. Reagan, J.R., Boyer, T.P., Garcia, H.E., Locarnini, R.A., Baranova, O.K., Bouchard, C., Cross, S.L., Mishonov, A.V., Paver, C.R., Seidov, D. et al. (2024). World Ocean Atlas 2023. NOAA National Centers for Environmental Information. URL: https://www.ncei.noaa.gov/access/world-ocean-atlas-2023/. Accessed 2026-03-20.

21. de Boyer Montégut, C., Madec, G., Fischer, A.S., Lazar, A., and Iudicone, D. (2004). Mixed layer depth over the global ocean: An examination of profile data and a profile-based climatology. Journal of Geophysical Research: Oceans 109, C12003. doi: 10.1029/2004JC002378.

22. Chen, T., and Guestrin, C. (2016). XGBoost: A scalable tree boosting system. In Proceedings of the 22nd ACM SIGKDD international conference on knowledge discovery and data mining. pp. 785–794. doi: 10.1145/2939672.2939785.

23. Lundberg, S.M., and Lee, S.I. (2017). A unified approach to interpreting model predictions. Advances in Neural Information Processing Systems 30. URL: https://papers.nips.cc/paper/2017/hash/8a20a8621978632d76c43dfd28b67767-Abstract.html. arXiv preprint arXiv:1705.07874.

24. Morse, D., Salois, P., Markovic, P., and Hastings, J.W. (1995). A nuclear-encoded form II RuBisCO in dinoflagellates. Science 268, 1622–1624. doi: 10.1126/science.7777861.

25. Tabita, F.R., Satagopan, S., Hanson, T.E., Kreel, N.E., and Scott, S.S. (2008). Distinct form I, II, III, and IV Rubisco proteins from the three kingdoms of life provide clues about Rubisco evolution and structure/function relationships. Journal of Experimental Botany 59, 1515–1524. doi: 10.1093/jxb/erm361.

26. Zhang, H., and Lin, S. (2003). Complex gene structure of the form II Rubisco in Prorocentrum minimum (Dinophyceae). Journal of Phycology 39, 1160–1171. doi: 10.1111/j.0022-3646.2003.03-055.x.

27. Benjamini, Y., and Hochberg, Y. (1995). Controlling the false discovery rate: A practical and powerful approach to multiple testing. Journal of the Royal Statistical Society: Series B (Methodological) 57, 289–300. doi: 10.1111/j.2517-6161.1995.tb02031.x.

28. McInnes, L., Healy, J., and Melville, J. (2018). UMAP: Uniform manifold approximation and projection for dimension reduction. arXiv preprint arXiv:1802.03426. URL: https://arxiv.org/abs/1802.03426.

29. Hotelling, H. (1936). Relations between two sets of variates. Biometrika 28, 321–377. doi: 10.1093/biomet/28.3-4.321.

30. Liu, Z., Wang, Y., Vaidya, S., Ruehle, F., Halverson, J., Soljačić, M., Hou, T.Y., and Tegmark, M. (2024). KAN: Kolmogorov-Arnold networks. arXiv preprint arXiv:2404.19756. doi: 10.48550/arXiv.2404.19756.

31. Shoguchi, E., Shinzato, C., Kawashima, T., Gyoja, F., Mungpakdee, S., Koyanagi, R., Takeuchi, T., Hisata, K., Tanaka, M., Fujiwara, M. et al. (2013). Draft assembly of the *Symbiodinium minutum* nuclear genome reveals dinoflagellate gene structure. Current Biology 23, 1399–1408. doi: 10.1016/j.cub.2013.05.062.

32. Allen, A.E., LaRoche, J., Maheswari, U., Lommer, M., Schauer, N., Lopez, P.J., Finazzi, G., Fernie, A.R., and Bowler, C. (2008). Whole-cell response of the pennate diatom *Phaeodactylum tricornutum* to iron starvation. Proceedings of the National Academy of Sciences 105, 10438–10443. doi: 10.1073/pnas.0711370105.

33. Campello, R.J., Moulavi, D., and Sander, J. (2013). Density-based clustering based on hierarchical density estimates. In Pacific-Asia Conference on Knowledge Discovery and Data Mining. Springer pp. 160–172. doi: 10.1007/978-3-642-37456-2_14.

34. Hanson, C.A., Fuhrman, J.A., Horner-Devine, M.C., and Martiny, J.B.H. (2012). Beyond biogeographic patterns: processes shaping the microbial landscape. Nature Reviews Microbiology 10, 497–506. doi: 10.1038/nrmicro2795.

35. Wilson, B.A., Foy, S.G., Neme, R., and Masel, J. (2017). Young genes are highly disordered as predicted by the preadaptation hypothesis of de novo gene birth. Nature Ecology & Evolution 1, 0146. doi: 10.1038/s41559-017-0146.

36. Flowers, J.M., Hazzouri, K.M., Pham, G.M., Rosas, U., Bahmani, T., Khraiwesh, B., Nelson, D.R., Jijakli, K., Abdrabu, R., Harris, E.H. et al. (2015). Whole-genome resequencing reveals extensive natural variation in the model green alga *Chlamydomonas reinhardtii*. Plant Cell 27, 2353–2369. doi: 10.1105/tpc.15.00492.

37. Osuna-Cruz, C.M., Bilcke, G., Vancaester, E., De Decker, S., Bones, A.M., Winge, P., Poulsen, N., Bulankova, P., Verhelst, B., Audoor, S., et al. (2020). The *Seminavis robusta* genome provides insights into the evolutionary adaptations of benthic diatoms. Nature Communications 11, 3320. doi: 10.1038/s41467-020-17191-8.

38. Tai, V., Poon, A.F.Y., Paulsen, I.T., and Palenik, B. (2011). Selection in coastal *Synechococcus* (Cyanobacteria) populations evaluated from environmental metagenomes. PLoS ONE 6, e24249. doi: 10.1371/journal.pone.0024249.

39. Koester, J.A., Swanson, W.J., and Armbrust, E.V. (2013). Positive selection within a diatom species acts on putative protein interactions and transcriptional regulation. Molecular Biology and Evolution 30, 422–434. doi: 10.1093/molbev/mss242.

40. Radford, A., Wu, J., Child, R., Luan, D., Amodei, D., and Sutskever, I. (2019). Language models are unsupervised multitask learners. OpenAI blog 1, 9.

41. Karpathy, A. (2023). nanoGPT. https://github.com/karpathy/nanoGPT. Minimal GPT training framework.

42. Steinegger, M., and Söding, J. (2017). MMseqs2 enables sensitive protein sequence searching for the analysis of massive data sets. Nature Biotechnology 35, 1026–1028. doi: 10.1038/nbt.3988.

43. Passaro, S., Corso, G., Wohlwend, J., Reveiz, M., Thaler, S., Somnath, V.R., Getz, N., Portnoi, T., Roy, J., Stark, H. et al. (2025). Boltz-2: Towards accurate and efficient binding affinity prediction. bioRxiv. doi: 10.1101/2025.06.14.659707.

44. Abramson, J., Adler, J., Dunger, J., Evans, R., Green, T., Pritzel, A., Ronneberger, O., Willmore, L., Ballard, A.J., Bambrick, J. et al. (2024). Accurate structure prediction of biomolecular interactions with AlphaFold 3. Nature 630, 493–500. doi: 10.1038/s41586-024-07487-w.

45. van Kempen, M., Kim, S.S., Tumescheit, C., Mirdita, M., Lee, J., Gilchrist, C.L., Söding, J., and Steinegger, M. (2024). Fast and accurate protein structure search with Foldseek. Nature Biotechnology 42, 243–246. doi: 10.1038/s41587-023-01773-0.

46. Bardes, A., Ponce, J., and LeCun, Y. (2022). VICReg: Variance-invariance-covariance regularization for self-supervised learning. In International Conference on Learning Representations. URL: https://openreview.net/forum?id=xm6YD62D1Ub.

47. Nelson, D.R., and Salehi-Ashtiani, K. (2026). TARA-WorldModel-VICReg: Joint environment–proteome embedding model via self-supervised learning. Hugging Face. URL: https://huggingface.co/GreenGenomicsLab/TARA-WorldModel-VICReg VICReg-aligned joint embedding checkpoints and hyperparameter sweep results.

48. Vértessy, B.G., and Tóth, J. (2009). Keeping uracil out of DNA: Physiological role, structure and catalytic mechanism of dUTPases. Accounts of Chemical Research 42, 97–106. doi: 10.1021/ar800114w.

49. Haimovich-Dayan, M., Garfinkel, N., Ewe, D., Marcus, Y., Gruber, A., Wagner, H., Kroth, P.G., and Kaplan, A. (2013). The role of C4 metabolism in the marine diatom *Phaeodactylum tricornutum*. New Phytologist 197, 177–185. doi: 10.1111/j.1469-8137.2012.04375.x.

50. Maumus, F., Allen, A.E., Mhiri, C., Hu, H., Jabbari, K., Vardi, A., Grandbastien, M.A., and Bowler, C. (2009). Potential impact of stress activated retrotransposons on genome evolution in a marine diatom. BMC Genomics 10, 624. doi: 10.1186/1471-2164-10-624.

51. Egue, F., Chenais, B., Tastard, E., Marchand, J., Hiard, S., Gateau, H., Hermann, D., Morant-Manceau, A., Casse, N., and Caruso, A. (2015). Expression of the retrotransposons Surcouf and Blackbeard in the marine diatom *Phaeodactylum tricornutum* under thermal stress. Phycologia 54, 617–627. doi: 10.2216/15-52.1.

52. Pargana, A., Musacchia, F., Sanges, R., Russo, M.T., Ferrante, M.I., Bowler, C., and Zingone, A. (2020). Intraspecific diversity in the cold stress response of transposable elements in the diatom *Leptocylindrus aporus*. Genes 11, 9. doi: 10.3390/genes11010009.

53. Song, B., Morse, D., Song, Y., Fu, Y., Lin, X., Wang, W., Cheng, S., Chen, W., Liu, X., and Lin, S. (2017). Comparative genomics reveals two major bouts of gene retroposition coinciding with crucial periods of *Symbiodinium* evolution. Genome Biology and Evolution 9, 2037–2047. doi: 10.1093/gbe/evx144.

54. Lescot, M., Hingamp, P., Kojima, K.K., Villar, E., Romac, S., Veluchamy, A., Boccara, M., Jaillon, O., Iudicone, D., Bowler, C. et al. (2016). Reverse transcriptase genes are highly abundant and transcriptionally active in marine plankton assemblages. The ISME Journal 10, 1134–1146. doi: 10.1038/ismej.2015.192.

55. Litchman, E., and Klausmeier, C.A. (2007). The role of functional traits and trade-offs in structuring phytoplankton communities: scaling from cellular to ecosystem level. Ecology Letters 10, 1170–1181. doi: 10.1111/j.1461-0248.2007.01117.x.

56. Richardson, L., Allen, B., Baldi, G., Beracochea, M., Bileschi, M.L., Burdett, T., Burgin, J., Caballero-Pérez, J., Cochrane, G., Colwell, L.J. et al. (2023). MGnify: the microbiome sequence data analysis resource in 2023. Nucleic Acids Research 51, D753–D759. doi: 10.1093/nar/gkac1080.

57. EBI MGnify (2023). TARA Oceans metagenome assemblies. European Bioinformatics Institute. URL: https://www.ebi.ac.uk/metagenomics/ MGYA accessions; see also Richardson et al. (2023).

58. National Center for Genome Resources (2014). Marine Microbial Eukaryote Transcriptome Sequencing Project (MMETSP). NCBI BioProject. URL: https://www.ncbi.nlm.nih.gov/bioproject/PRJNA231566 BioProject PRJNA231566; see also Keeling et al. (2014).

59. Nelson, D.R., Plouviez, M., Daakour, S., Jaiswal, A., Fu, W., Amin, S.A., and Salehi-Ashtiani, K. (2026). ELF-NET Code S1: analysis code. Zenodo. URL: 10.5281/zenodo.22910378. doi: 10.5281/zenodo.22910378.

60. Nelson, D.R., Jaiswal, A.K., Ismail, N.S., Mystikou, A., and Salehi-Ashtiani, K. (2025). algaGPT: Causal language model for binary classification of microalgal protein sequences. Hugging Face. URL: https://huggingface.co/GreenGenomicsLab/algaGPT nanoGPT-based classifier; model weights and tokenizer.

61. Nelson, D.R., and Salehi-Ashtiani, K. (2026). TARA-XGBoost-Bidirectional: Bidirectional XGBoost models linking marine environment and protein domain composition. Hugging Face. URL: https://huggingface.co/GreenGenomicsLab/TARA-XGBoost-Bidirectional 100 forward (environment to PFAM) and 31 reverse (PFAM to environment) XGBoost regressors with SHAP importance.

62. Daakour, S., Nelson, D.R., and Salehi-Ashtiani, K. (2026). Dark-whiteGPLM model check-points. Hugging Face. URL: https://huggingface.co/SarahDaakour/dark-whiteGPLM.

63. Daakour, S., Nelson, D.R., and Salehi-Ashtiani, K. (2026). dark-whiteGPLM: Character-level protein language models for marine microalgae proteomes. Zenodo. URL: 10.5281/zenodo.20933332. doi: 10.5281/zenodo.20933332.

64. Buchfink, B., Xie, C., and Huson, D.H. (2015). Fast and sensitive protein alignment using DIAMOND. Nature Methods 12, 59–60. doi: 10.1038/nmeth.3176.

65. Sayers, E.W., Cavanaugh, M., Clark, K., Pruitt, K.D., Sherry, S.T., Yankie, L., and Karsch-Mizrachi, I. (2023). GenBank 2023 update. Nucleic Acids Research 51, D141–D144. doi: 10.1093/nar/gkac1012.

66. Nelson, D.R., Chaiboonchoe, A., Fu, W., Hazzouri, K.M., Huang, Z., Jaiswal, A., Daakour, S., Mystikou, A., Arnoux, M., Sultana, M. et al. (2019). Potential for heightened sulfurmetabolic capacity in coastal subtropical microalgae. iScience 11, 450–465. doi: 10.1016/j.isci.2018.12.035.

67. Nelson, D.R., Hazzouri, K.M., Lauersen, K.J., Jaiswal, A., Chaiboonchoe, A., Mystikou, A., Fu, W., Daakour, S., Dohai, B., Alzahmi, A. et al. (2021). Large-scale genome sequencing reveals the driving forces of viruses in microalgal evolution. Cell Host & Microbe 29, 250–266.e8. doi: 10.1016/j.chom.2020.12.005.

68. EMBL-EBI (2024). Pfam protein families database, version 37.2. European Bioinformatics Institute. URL: https://ftp.ebi.ac.uk/pub/databases/Pfam/releases/Pfam37.2/ 24,076 protein family HMMs; November 2024 release; see also Mistry et al. (2021).

69. Google (2025). Satellite Embedding V1 (AlphaEarth Foundations) annual image collection. Google Earth Engine. URL: https://developers.google.com/earth-engine/datasets/catalog/GOOGLE_SATELLITE_EMBEDDING_V1_ANNUAL Collection ID: GOOGLE/SATELLITE_EMBEDDING/V1/ANNUAL; 64-dimensional satellite embeddings at 10 m resolution.

70. The Gene Ontology Consortium, Aleksander, S.A., Balhoff, J., Carbon, S., Cherry, J.M., Drabkin, H.J., Ebert, D., Feuermann, M., Gaudet, P., Harris, N.L., et al. (2023). The Gene Ontology knowledgebase in 2023. Genetics 224, iyad031. doi: 10.1093/genetics/iyad031.

71. Gene Ontology Consortium (2023). Pfam2GO mapping file. Gene Ontology. URL: http://current.geneontology.org/ontology/external2go/pfam2go Pfam-to-GO term mappings; see also Gene Ontology Consortium (2023).

72. Nelson, D.R. (2026). ELF-NET Data S1: Domain–environment association and modeling results. Zenodo. URL: 10.5281/zenodo.18538438. doi: 10.5281/zenodo.18538438.

73. Nelson, D.R. (2026). ELF-NET Data S2: algaGPT-classified algal protein sequences from the mixed-source domain-analysis set. Zenodo. URL: 10.5281/zenodo.18728836. doi: 10.5281/zenodo.18728836.

74. Nelson, D.R. (2026). ELF-NET Data S3: Pfam-A hmmsearch results for the mixed-source input collection. Zenodo. URL: 10.5281/zenodo.18786751. doi: 10.5281/zenodo.18786751.

75. Nelson, D.R., Plouviez, M., and Salehi-Ashtiani, K. (2026). ELF-NET Data S4: RuBisCO lineage analysis package. Zenodo. URL: 10.5281/zenodo.18786775. doi: 10.5281/zenodo.18786775.

76. Nelson, D.R. (2026). ELF-NET Data S5: AlphaEarth satellite embedding matrix for GPS-mapped domain-analysis samples. Zenodo. URL: 10.5281/zenodo.18786761. doi: 10.5281/zenodo.18786761 Publicly available under CC BY 4.0.

77. Nelson, D.R. (2026). ELF-NET Data S6: KAN-CCA and sparse CCA cross-validation results. Zenodo. URL: 10.5281/zenodo.18786766. doi: 10.5281/zenodo.18786766.

78. Nelson, D.R. (2026). ELF-NET Data S7: Novel domain discovery results from the Pfamdark proteome. Zenodo. URL: 10.5281/zenodo.18786771. doi: 10.5281/zenodo.18786771.

79. Nelson, D.R. (2026). ELF-NET Data S8: DIAMOND BLASTp comparison of algaGPT-classified proteins. Zenodo. URL: 10.5281/zenodo.19441355. doi: 10.5281/zenodo.19441355 Publicly available under CC BY 4.0.

80. Nelson, D.R. (2026). ELF-NET Data S9: Dark-proteome selection analysis and GPLM training sequences. Zenodo. URL: 10.5281/zenodo.20486973. doi: 10.5281/zenodo.20486973.

81. Nelson, D.R., Jaiswal, A., and Salehi-Ashtiani, K. (2026). ELF-NET Data S10: Boltz-2 and AlphaFold 3 structure predictions for 3,600 novel dark proteome domains. Zenodo. URL: 10.5281/zenodo.20508680. doi: 10.5281/zenodo.20508680.

82. Daakour, S., Nelson, D.R., and Salehi-Ashtiani, K. (2026). Dark-whiteGPLM training data. Hugging Face. URL: https://huggingface.co/datasets/SarahDaakour/dark-whiteGPLM-data.

83. Eddy, S.R. (2020). HMMER: biosequence analysis using profile hidden Markov models, version 3.3.2. Eddy Lab, Harvard University. URL: http://hmmer.org/ See also Eddy (2011).

84. Korf, I. (2004). Gene finding in novel genomes. BMC Bioinformatics 5, 59. doi: 10.1186/1471-2105-5-59.

85. Katoh, K., and Standley, D.M. (2013). MAFFT multiple sequence alignment software version 7: improvements in performance and usability. Molecular Biology and Evolution 30, 772–780. doi: 10.1093/molbev/mst010.

86. Blum, M., Chang, H.Y., Chuguransky, S., Grego, T., Kandasaamy, S., Mitchell, A., Nuka, G., Paysan-Lafosse, T., Qureshi, M., Raj, S. et al. (2021). The InterPro protein families and domains database: 20 years on. Nucleic Acids Research 49, D344–D354. doi: 10.1093/nar/gkaa977.

87. Fu, L., Niu, B., Zhu, Z., Wu, S., and Li, W. (2012). CD-HIT: accelerated for clustering the next-generation sequencing data. Bioinformatics 28, 3150–3152. doi: 10.1093/bioinformatics/bts565.

88. Minh, B.Q., Schmidt, H.A., Chernomor, O., Schrempf, D., Woodhams, M.D., von Haeseler, A., and Lanfear, R. (2020). IQ-TREE 2: new models and efficient methods for phylogenetic inference in the genomic era. Molecular Biology and Evolution 37, 1530–1534. doi: 10.1093/molbev/msaa015.

89. Price, M.N., Dehal, P.S., and Arkin, A.P. (2010). FastTree 2 – approximately maximum-likelihood trees for large alignments. PLoS ONE 5, e9490. doi: 10.1371/journal.pone.0009490.

90. Cock, P.J.A., Antao, T., Chang, J.T., Chapman, B.A., Cox, C.J., Dalke, A., Friedberg, I., Hamelryck, T., Kauff, F., Wilczynski, B. et al. (2009). Biopython: freely available Python tools for computational molecular biology and bioinformatics. Bioinformatics 25, 1422–1423. doi: 10.1093/bioinformatics/btp163.

91. Python Software Foundation (2023). Python language reference, version 3.10.12. URL: https://www.python.org/ RRID:SCR_008394.

92. Paszke, A., Gross, S., Massa, F., Lerer, A., Bradbury, J., Chanan, G., Killeen, T., Lin, Z., Gimelshein, N., Antiga, L. et al. (2019). PyTorch: An imperative style, high-performance deep learning library. In Advances in Neural Information Processing Systems vol. 32. pp. 8024–8035. URL: https://proceedings.neurips.cc/paper/2019/hash/bdbca288fee7f92f2bfa9f7012727740-Abstract.html.

93. Wolf, T., Debut, L., Sanh, V., Chaumond, J., Delangue, C., Moi, A., Cistac, P., Rault, T., Louf, R., Funtowicz, M. et al. (2020). Transformers: State-of-the-art natural language processing. In Proceedings of the 2020 Conference on Empirical Methods in Natural Language Processing: System Demonstrations. Association for Computational Linguistics pp. 38–45. URL: https://aclanthology.org/2020.emnlp-demos.6/. doi: 10.18653/v1/2020.emnlp-demos.6.

94. Pedregosa, F., Varoquaux, G., Gramfort, A., Michel, V., Thirion, B., Grisel, O., Blondel, M., Prettenhofer, P., Weiss, R., Dubourg, V. et al. (2011). Scikit-learn: Machine learning in Python. Journal of Machine Learning Research 12, 2825–2830. URL: https://www.jmlr.org/papers/v12/pedregosa11a.html.

95. Virtanen, P., Gommers, R., Oliphant, T.E., Haberland, M., Reddy, T., Cournapeau, D., Burovski, E., Peterson, P., Weckesser, W., Bright, J. et al. (2020). SciPy 1.0: fundamental algorithms for scientific computing in Python. Nature Methods 17, 261–272. doi: 10.1038/s41592-019-0686-2.

96. Harris, C.R., Millman, K.J., van der Walt, S.J., Gommers, R., Virtanen, P., Cournapeau, D., Wieser, E., Taylor, J., Berg, S., Smith, N.J., et al. (2020). Array programming with NumPy. Nature 585, 357–362. doi: 10.1038/s41586-020-2649-2.

97. McKinney, W. (2010). Data structures for statistical computing in Python. In Proceedings of the 9th Python in Science Conference. pp. 56–61. doi: 10.25080/Majora-92bf1922-00a.

98. Seabold, S., and Perktold, J. (2010). statsmodels: Econometric and statistical modeling with Python. In Proceedings of the 9th Python in Science Conference. pp. 92–96. doi: 10.25080/Majora-92bf1922-011.

99. Hoyer, S., and Hamman, J. (2017). xarray: N-D labeled arrays and datasets in Python. Journal of Open Research Software 5, 10. doi: 10.5334/jors.148.

100. van der Maaten, L., and Hinton, G. (2008). Visualizing data using t-SNE. Journal of Machine Learning Research 9, 2579–2605. URL: https://jmlr.org/papers/v9/vandermaaten08a.html.

101. Levy Karin, E., Mirdita, M., and Söding, J. (2020). MetaEuk—sensitive, high-throughput gene discovery, and annotation for large-scale eukaryotic metagenomics. Microbiome 8, 48. doi: 10.1186/s40168-020-00808-x.

102. Aitchison, J. (1982). The statistical analysis of compositional data. Journal of the Royal Statistical Society: Series B (Methodological) 44, 139–160. doi: 10.1111/j.2517-6161.1982.tb01195.x.

103. Nyholt, D.R. (2004). A simple correction for multiple testing for single-nucleotide polymorphisms in linkage disequilibrium with each other. The American Journal of Human Genetics 74, 765–769. doi: 10.1086/383251.

104. EMBL-EBI (2021). InterPro REST API for protein family annotations. European Bioinformatics Institute. URL: https://www.ebi.ac.uk/interpro/api See also Blum et al. (2021).

105. He, K., Zhang, X., Ren, S., and Sun, J. (2015). Delving deep into rectifiers: Surpassing human-level performance on ImageNet classification. In 2015 IEEE International Conference on Computer Vision (ICCV). IEEE pp. 1026–1034. doi: 10.1109/iccv.2015.123.

106. Loshchilov, I., and Hutter, F. (2019). Decoupled weight decay regularization. In International Conference on Learning Representations. URL: https://openreview.net/forum?id=Bkg6RiCqY7.

107. Zhang, Y., and Skolnick, J. (2005). TM-align: a protein structure alignment algorithm based on the TM-score. Nucleic Acids Research 33, 2302–2309. doi: 10.1093/nar/gki524.

108. Kyte, J., and Doolittle, R.F. (1982). A simple method for displaying the hydropathic character of a protein. Journal of Molecular Biology 157, 105–132. doi: 10.1016/0022-2836(82)90515-0.

109. Dunker, A.K., Lawson, J.D., Brown, C.J., Williams, R.M., Romero, P., Oh, J.S., Oldfield, C.J., Campen, A.M., Ratliff, C.M., Hipps, K.W. et al. (2001). Intrinsically disordered protein. Journal of Molecular Graphics and Modelling 19, 26–59. doi: 10.1016/S1093-3263(00)00138-8.

110. Das, R.K., and Pappu, R.V. (2013). Conformations of intrinsically disordered proteins are influenced by linear sequence distributions of oppositely charged residues. Proceedings of the National Academy of Sciences 110, 13392–13397. doi: 10.1073/pnas.1304749110.

111. Chou, P.Y., and Fasman, G.D. (1979). Prediction of the secondary structure of proteins from their amino acid sequence. Advances in Enzymology and Related Areas of Molecular Biology 47, 45–148. doi: 10.1002/9780470122921.ch2.

112. Borcard, D., Legendre, P., and Drapeau, P. (1992). Partialling out the spatial component of ecological variation. Ecology 73, 1045–1055. doi: 10.2307/1940179.

113. Borcard, D., and Legendre, P. (2002). All-scale spatial analysis of ecological data by means of principal coordinates of neighbour matrices. Ecological Modelling 153, 51–68. doi: 10.1016/s0304-3800(01)00501-4.

114. Daakour, S. (2026). Dark-whiteGPLM: Code repository. GitHub. URL: https://github.com/SarahD4/dark-whiteGPLM.

## References

1. Buchfink, B., Xie, C., and Huson, D.H. (2015). Fast and sensitive protein alignment using DIAMOND. Nature Methods 12, 59–60. doi: 10.1038/nmeth.3176.

2. Nelson, D.R., Jaiswal, A.K., Ismail, N.S., Mystikou, A., and Salehi-Ashtiani, K. (2025). Panmicroalgal dark proteome mapping via interpretable deep learning and synthetic chimeras. Patterns 6, 101373. doi: 10.1016/j.patter.2025.101373.

3. Vorobev, A., Dupouy, M., Carradec, Q., Delmont, T.O., Annamalé, A., Wincker, P., and Pelletier, E. (2020). Transcriptome reconstruction and functional analysis of eukaryotic marine plankton communities via high-throughput metagenomics and metatranscriptomics. Genome Research 30, 647–659. doi: 10.1101/gr.253070.119.

4. Wicker, T., Sabot, F., Hua-Van, A., Bennetzen, J.L., Capy, P., Chalhoub, B., Flavell, A., Leroy, P., Morgante, M., Panaud, O. et al. (2007). A unified classification system for eukaryotic transposable elements. Nature Reviews Genetics 8, 973–982. doi: 10.1038/nrg2165.

5. Davies, P.L. (2014). Ice-binding proteins: a remarkable diversity of structures for stopping and starting ice growth. Trends in Biochemical Sciences 39, 548–555. doi: 10.1016/j.tibs.2014.09.005.

6. Tabita, F.R., Satagopan, S., Hanson, T.E., Kreel, N.E., and Scott, S.S. (2008). Distinct form I, II, III, and IV Rubisco proteins from the three kingdoms of life provide clues about Rubisco evolution and structure/function relationships. Journal of Experimental Botany 59, 1515–1524. doi: 10.1093/jxb/erm361.

7. Morse, D., Salois, P., Markovic, P., and Hastings, J.W. (1995). A nuclear-encoded form II RuBisCO in dinoflagellates. Science 268, 1622–1624. doi: 10.1126/science.7777861.

8. Zhang, H., and Lin, S. (2003). Complex gene structure of the form II Rubisco in Prorocentrum minimum (Dinophyceae). Journal of Phycology 39, 1160–1171. doi: 10.1111/j.0022-3646.2003.03-055.x.

9. Shoguchi, E., Shinzato, C., Kawashima, T., Gyoja, F., Mungpakdee, S., Koyanagi, R., Takeuchi, T., Hisata, K., Tanaka, M., Fujiwara, M. et al. (2013). Draft assembly of the *Symbiodinium minutum* nuclear genome reveals dinoflagellate gene structure. Current Biology 23, 1399–1408. doi: 10.1016/j.cub.2013.05.062.

10. Kapitonov, V.V., and Jurka, J. (2007). Helitrons on a roll: eukaryotic rolling-circle transposons. Trends in Genetics 23, 521–529. doi: 10.1016/j.tig.2007.08.004.

11. Bochman, M.L., Judge, C.P., and Zakian, V.A. (2011). The Pif1 family in prokaryotes: what are our helicases doing in your bacteria? Molecular Biology of the Cell 22, 1955–1959. doi: 10.1091/mbc.e11-01-0045.

12. Grindley, N.D.F., Whiteson, K.L., and Rice, P.A. (2006). Mechanisms of site-specific recombination. Annual Review of Biochemistry 75, 567–605. doi: 10.1146/annurev.biochem.73.011303.073908.

13. Adams, J.C. (2004). The thrombospondins. The International Journal of Biochemistry & Cell Biology 36, 961–968. doi: 10.1016/j.biocel.2004.01.004.

14. Ogura, T., and Wilkinson, A.J. (2001). AAA^+^ superfamily ATPases: common structure—diverse function. Genes to Cells 6, 575–597. doi: 10.1046/j.1365-2443.2001.00447.x.

15. Capy, P., Gasperi, G., Biémont, C., and Bazin, C. (2000). Stress and transposable elements: coevolution or useful parasites? Heredity 85, 101–106. doi: 10.1046/j.1365-2540.2000.00751.x.

16. Lescot, M., Hingamp, P., Kojima, K.K., Villar, E., Romac, S., Veluchamy, A., Boccara, M., Jaillon, O., Iudicone, D., Bowler, C. et al. (2016). Reverse transcriptase genes are highly abundant and transcriptionally active in marine plankton assemblages. The ISME Journal 10, 1134–1146. doi: 10.1038/ismej.2015.192.

17. Maumus, F., Allen, A.E., Mhiri, C., Hu, H., Jabbari, K., Vardi, A., Grandbastien, M.A., and Bowler, C. (2009). Potential impact of stress activated retrotransposons on genome evolution in a marine diatom. BMC Genomics 10, 624. doi: 10.1186/1471-2164-10-624.

18. Haimovich-Dayan, M., Garfinkel, N., Ewe, D., Marcus, Y., Gruber, A., Wagner, H., Kroth, P.G., and Kaplan, A. (2013). The role of C4 metabolism in the marine diatom *Phaeodactylum tricornutum*. New Phytologist 197, 177–185. doi: 10.1111/j.1469-8137.2012.04375.x.

19. Vértessy, B.G., and Tóth, J. (2009). Keeping uracil out of DNA: Physiological role, structure and catalytic mechanism of dUTPases. Accounts of Chemical Research 42, 97–106. doi: 10.1021/ar800114w.

20. Allen, A.E., LaRoche, J., Maheswari, U., Lommer, M., Schauer, N., Lopez, P.J., Finazzi, G., Fernie, A.R., and Bowler, C. (2008). Whole-cell response of the pennate diatom *Phaeodactylum tricornutum* to iron starvation. Proceedings of the National Academy of Sciences 105, 10438–10443. doi: 10.1073/pnas.0711370105.

21. Matsuda, Y., Koseki, M., Shimada, T., and Saito, T. (1995). Purification and characterization of a vegetative lytic enzyme responsible for liberation of daughter cells during the proliferation of *Chlamydomonas reinhardtii*. Plant and Cell Physiology 36, 681–689. doi: 10.1093/oxfordjournals.pcp.a078809.

22. Guschina, I.A., and Harwood, J.L. (2006). Lipids and lipid metabolism in eukaryotic algae. Progress in Lipid Research 45, 160–186. doi: 10.1016/j.plipres.2006.01.001.

23. Reinfelder, J.R. (2011). Carbon concentrating mechanisms in eukaryotic marine phytoplankton. Annual Review of Marine Science 3, 291–315. doi: 10.1146/annurev-marine-120709-142720.

24. Gong, W., Paerl, H., and Marchetti, A. (2018). Eukaryotic phytoplankton community spatiotem-poral dynamics as identified through gene expression within a eutrophic estuary. Environmental Microbiology 20, 1095–1111. doi: 10.1111/1462-2920.14049.

25. Moore, C.M., Mills, M.M., Arrigo, K.R., Berman-Frank, I., Bopp, L., Boyd, P.W., Galbraith, E.D., Geider, R.J., Guieu, C., Jaccard, S.L. et al. (2013). Processes and patterns of oceanic nutrient limitation. Nature Geoscience 6, 701–710. doi: 10.1038/ngeo1765.

26. Eddy, S.R. (2020). HMMER: biosequence analysis using profile hidden Markov models, version 3.3.2. Eddy Lab, Harvard University. URL: http://hmmer.org/ See also Eddy (2011).

27. Mistry, J., Chuguransky, S., Williams, L., Qureshi, M., Salazar, G.A., Sonnhammer, E.L., Tosatto, S.C., Paladin, L., Raj, S., Richardson, L.J. et al. (2021). Pfam: The protein families database in 2021. Nucleic Acids Research 49, D412–D419. doi: 10.1093/nar/gkaa913.

28. EMBL-EBI (2024). Pfam protein families database, version 37.2. European Bioinformatics Institute. URL: https://ftp.ebi.ac.uk/pub/databases/Pfam/releases/Pfam37.2/ 24,076 protein family HMMs; November 2024 release; see also Mistry et al. (2021).

29. Flowers, J.M., Hazzouri, K.M., Pham, G.M., Rosas, U., Bahmani, T., Khraiwesh, B., Nelson, D.R., Jijakli, K., Abdrabu, R., Harris, E.H. et al. (2015). Whole-genome resequencing reveals extensive natural variation in the model green alga *Chlamydomonas reinhardtii*. Plant Cell 27, 2353–2369. doi: 10.1105/tpc.15.00492.

30. Osuna-Cruz, C.M., Bilcke, G., Vancaester, E., De Decker, S., Bones, A.M., Winge, P., Poulsen, N., Bulankova, P., Verhelst, B., Audoor, S., et al. (2020). The *Seminavis robusta* genome provides insights into the evolutionary adaptations of benthic diatoms. Nature Communications 11, 3320. doi: 10.1038/s41467-020-17191-8.

31. Tai, V., Poon, A.F.Y., Paulsen, I.T., and Palenik, B. (2011). Selection in coastal *Synechococcus* (Cyanobacteria) populations evaluated from environmental metagenomes. PLoS ONE 6, e24249. doi: 10.1371/journal.pone.0024249.

32. Koester, J.A., Swanson, W.J., and Armbrust, E.V. (2013). Positive selection within a diatom species acts on putative protein interactions and transcriptional regulation. Molecular Biology and Evolution 30, 422–434. doi: 10.1093/molbev/mss242.

33. Passaro, S., Corso, G., Wohlwend, J., Reveiz, M., Thaler, S., Somnath, V.R., Getz, N., Portnoi, T., Roy, J., Stark, H. et al. (2025). Boltz-2: Towards accurate and efficient binding affinity prediction. bioRxiv. doi: 10.1101/2025.06.14.659707.

34. Carradec, Q., Pelletier, E., Da Silva, C., Alberti, A., Seeleuthner, Y., Blanc-Mathieu, R., Lima-Mendez, G., Rocha, F., Tirichine, L., Labadie, K., et al. (2018). A global ocean atlas of eukaryotic genes. Nature Communications 9, 373. doi: 10.1038/s41467-017-02342-1.

35. Ayling, M., Clark, M.D., and Leggett, R.M. (2020). New approaches for metagenome assembly with short reads. Briefings in Bioinformatics 21, 584–594. doi: 10.1093/bib/bbz020.

36. Pesant, S., Not, F., Picheral, M., Kandels-Lewis, S., Le Bescot, N., Gorsky, G., Iudicone, D., Karsenti, E., Speich, S., Troublé, R., et al. (2015). Open science resources for the discovery and analysis of Tara Oceans data. Scientific Data 2, 150023. doi: 10.1038/sdata.2015.23.

37. Clarke, K.R. (1993). Non-parametric multivariate analyses of changes in community structure. Australian Journal of Ecology 18, 117–143. doi: 10.1111/j.1442-9993.1993.tb00438.x.

38. Mantel, N. (1967). The detection of disease clustering and a generalized regression approach. Cancer Research 27, 209–220.

39. The Gene Ontology Consortium, Aleksander, S.A., Balhoff, J., Carbon, S., Cherry, J.M., Drabkin, H.J., Ebert, D., Feuermann, M., Gaudet, P., Harris, N.L., et al. (2023). The Gene Ontology knowledgebase in 2023. Genetics 224, iyad031. doi: 10.1093/genetics/iyad031.

40. Gene Ontology Consortium (2023). Pfam2GO mapping file. Gene Ontology. URL: http://current.geneontology.org/ontology/external2go/pfam2go Pfam-to-GO term mappings; see also Gene Ontology Consortium (2023).

41. Lindquist, S., and Craig, E.A. (1988). The heat-shock proteins. Annual Review of Genetics 22, 631–677. doi: 10.1146/annurev.ge.22.120188.003215.

42. Bartlett, D.H. (2002). Pressure effects on in vivo microbial processes. Biochimica et Biophysica Acta – Protein Structure and Molecular Enzymology 1595, 367–381. doi: 10.1016/S0167-4838(01)00357-0.

43. Falkowski, P.G., and Raven, J.A. (2007). Aquatic Photosynthesis. 2nd ed.. Princeton University Press. ISBN 978-1-4008-4972-7. doi: 10.1515/9781400849727.

44. Keeling, P.J., Burki, F., Wilcox, H.M., Allam, B., Allen, E.E., Amaral-Zettler, L.A., Armbrust, E.V., Archibald, J.M., Bharti, A.K., Bell, C.J. et al. (2014). The Marine Microbial Eukaryote Transcriptome Sequencing Project (MMETSP): Illuminating the functional diversity of eukaryotic life in the oceans through transcriptome sequencing. PLOS Biology 12, e1001889. doi: 10.1371/journal.pbio.1001889.

45. Sarmiento, J.L., and Gruber, N. (2006). Ocean Biogeochemical Dynamics. Princeton, NJ: Princeton University Press. ISBN 978-0-691-01707-5. doi: 10.1515/9781400849079.

46. Behrenfeld, M.J., and Falkowski, P.G. (1997). Photosynthetic rates derived from satellite-based chlorophyll concentration. Limnology and Oceanography 42, 1–20. doi: 10.4319/lo.1997.42.1.0001.

47. Stoecker, D.K., Hansen, P.J., Caron, D.A., and Mitra, A. (2017). Mixotrophy in the marine plankton. Annual Review of Marine Science 9, 311–335. doi: 10.1146/annurev-marine-010816-060617.

48. Asada, K. (2006). Production and scavenging of reactive oxygen species in chloroplasts and their functions. Plant Physiology 141, 391–396. doi: 10.1104/pp.106.082040.

49. Casacuberta, E., and González, J. (2013). The impact of transposable elements in environmental adaptation. Molecular Ecology 22, 1503–1517. doi: 10.1111/mec.12170.

50. Lin, I.P., Jiang, P.L., Chen, C.S., and Tzen, J.T. (2012). A unique caleosin serving as the major integral protein in oil bodies isolated from *Chlorella* sp. cells cultured with limited nitrogen. Plant Physiology and Biochemistry 61, 80–87. doi: 10.1016/j.plaphy.2012.09.008.

51. Converti, A., Casazza, A.A., Ortiz, E.Y., Perego, P., and Del Borghi, M. (2009). Effect of temperature and nitrogen concentration on the growth and lipid content of *Nannochloropsis oculata* and *Chlorella vulgaris* for biodiesel production. Chemical Engineering and Processing: Process Intensification 48, 1146–1151. doi: 10.1016/j.cep.2009.03.006.

52. Feder, M.E., and Hofmann, G.E. (1999). Heat-shock proteins, molecular chaperones, and the stress response: evolutionary and ecological physiology. Annual Review of Physiology 61, 243–282. doi: 10.1146/annurev.physiol.61.1.243.

53. Yan, T., Wang, K., Feng, K., Gao, X., Jin, Y., Wu, H., Zhang, W., and Wei, L. (2023). Remodeling of the 3D chromatin architecture in the marine microalga *Nannochloropsis oceanica* during lipid accumulation. Biotechnology for Biofuels and Bioproducts 16, 129. doi: 10.1186/s13068-023-02378-0.

54. Bassham, D.C. (2007). Plant autophagy—more than a starvation response. Current Opinion in Plant Biology 10, 587–593. doi: 10.1016/j.pbi.2007.06.006.

55. Schroda, M., Hemme, D., and Mühlhaus, T. (2015). The Chlamydomonas heat stress response. The Plant Journal 82, 466–480. doi: 10.1111/tpj.12816.

56. Longhurst, A.R. (2006). Ecological Geography of the Sea. 2nd ed.. Elsevier/Academic Press. ISBN 978-0-12-455521-1. URL: https://www.educate.elsevier.com/book/details/9780124555211.

57. Aitchison, J. (1982). The statistical analysis of compositional data. Journal of the Royal Statistical Society: Series B (Methodological) 44, 139–160. doi: 10.1111/j.2517-6161.1982.tb01195.x.

58. Gloor, G.B., Macklaim, J.M., Pawlowsky-Glahn, V., and Egozcue, J.J. (2017). Microbiome datasets are compositional: and this is not optional. Frontiers in Microbiology 8, 2224. doi: 10.3389/fmicb.2017.02224.

59. Loshchilov, I., and Hutter, F. (2019). Decoupled weight decay regularization. In International Conference on Learning Representations. URL: https://openreview.net/forum?id=Bkg6RiCqY7.

60. González, I., Déjean, S., Martin, P.G., and Baccini, A. (2008). CCA: An R package to extend canonical correlation analysis. Journal of Statistical Software 23, 1–14. doi: 10.18637/jss.v023.i12.

61. Liu, Z., Wang, Y., Vaidya, S., Ruehle, F., Halverson, J., Soljačić, M., Hou, T.Y., and Tegmark, M. (2024). KAN: Kolmogorov-Arnold networks. arXiv preprint arXiv:2404.19756. doi: 10.48550/arXiv.2404.19756.

62. Wisecaver, J.H., and Hackett, J.D. (2011). Dinoflagellate genome evolution. Annual Review of Microbiology 65, 369–387. doi: 10.1146/annurev-micro-090110-102841.

63. Stumpf, R.P., Culver, M.E., Tester, P.A., Tomlinson, M., Kirkpatrick, G.J., Pederson, B.A., Truby, E., Ransibrahmanakul, V., and Soracco, M. (2003). Monitoring *Karenia brevis* blooms in the Gulf of Mexico using satellite ocean color imagery and other data. Harmful Algae 2, 147–160. doi: 10.1016/S1568-9883(02)00083-5.

64. Hu, C., Muller-Karger, F.E., Taylor, C., Carder, K.L., Kelble, C., Johns, E., and Heil, C.A. (2005). Red tide detection and tracing using MODIS fluorescence data: A regional example in SW Florida coastal waters. Remote Sensing of Environment 97, 311–321. doi: 10.1016/j.rse.2005.05.013.

65. Mock, T., Otillar, R.P., Strauss, J., McMullan, M., van Staal, P., et al. (2017). Evolutionary genomics of the cold-adapted diatom *Fragilariopsis cylindrus*. Nature 541, 536–540. doi: 10.1038/nature20803.

66. Bayer-Giraldi, M., Uhlig, C., John, U., Mock, T., and Valentin, K. (2010). Antifreeze proteins in polar sea ice diatoms: diversity and gene expression in the genus *Fragilariopsis*. Environmental Microbiology 12, 1041–1052. doi: 10.1111/j.1462-2920.2009.02149.x.

67. Raymond, J.A., and Kim, H.J. (2012). Possible role of horizontal gene transfer in the colonization of sea ice by algae. PLoS ONE 7, e35968. doi: 10.1371/journal.pone.0035968.

68. Buma, A.G., Helbling, E.W., de Boer, M.K., and Villafañe, V.E. (2001). Patterns of DNA damage and photoinhibition in temperate South-Atlantic picophytoplankton exposed to solar ultraviolet radiation. Journal of Photochemistry and Photobiology B: Biology 62, 9–18. doi: 10.1016/S1011-1344(01)00156-7.

69. Koley, S., Foley, K., Perrine, Z., Morley, S.A., Kambhampati, S., Gomez, O., Chu, K.L., Chou, Y.H., Wei, M., Tzeng, S.C. et al. (2026). Metabolic rewiring and biomass redistribution enable optimized mixotrophic growth in *Chlamydomonas*. Proceedings of the National Academy of Sciences 123. doi: 10.1073/pnas.2522572123.

70. Bardes, A., Ponce, J., and LeCun, Y. (2022). VICReg: Variance-invariance-covariance regularization for self-supervised learning. In International Conference on Learning Representations. URL: https://openreview.net/forum?id=xm6YD62D1Ub.

71. Nelson, D.R., and Salehi-Ashtiani, K. (2026). TARA-WorldModel-VICReg: Joint environment– proteome embedding model via self-supervised learning. Hugging Face. URL: https://huggingface.co/GreenGenomicsLab/TARA-WorldModel-VICReg VICReg-aligned joint embedding checkpoints and hyperparameter sweep results.

72. Finkel, Z.V., Beardall, J., Flynn, K.J., Quigg, A., Rees, T.A.V., and Raven, J.A. (2010). Phytoplankton in a changing world: cell size and elemental stoichiometry. Journal of Plankton Research 32, 119–137. doi: 10.1093/plankt/fbp098.

73. Ellegaard, M., and Ribeiro, S. (2018). The long-term persistence of phytoplankton resting stages in aquatic ‘seed banks’. Biological Reviews 93, 166–183. doi: 10.1111/brv.12338.

74. Planes, S., Allemand, D., Agostini, S., Banaigs, B., Boissin, E., Boss, E., Bourdin, G., Bowler, C., Douville, E., Flores, J.M. et al. (2019). The Tara Pacific expedition—a pan-ecosystemic approach of the “-omics” complexity of coral reef holobionts across the Pacific Ocean. PLOS Biology 17, e3000483. doi: 10.1371/journal.pbio.3000483.

75. Duarte, C.M. (2015). Seafaring in the 21st century: The Malaspina 2010 circumnavigation expedition. Limnology and Oceanography Bulletin 24, 11–14. doi: 10.1002/lob.10008.

76. Karl, D.M., and Lukas, R. (1996). The Hawaii Ocean Time-series (HOT) program: Background, rationale and field implementation. Deep-Sea Research Part II 43, 129–156. doi: 10.1016/0967-0645(96)00005-7.

77. Steinberg, D.K., Carlson, C.A., Bates, N.R., Johnson, R.J., Michaels, A.F., and Knap, A.H. (2001). Overview of the US JGOFS Bermuda Atlantic Time-series Study (BATS): a decade-scale look at ocean biology and biogeochemistry. Deep-Sea Research Part II 48, 1405–1447. doi: 10.1016/S0967-0645(00)00148-X.

78. Borcard, D., Legendre, P., and Drapeau, P. (1992). Partialling out the spatial component of ecological variation. Ecology 73, 1045–1055. doi: 10.2307/1940179.

79. Borcard, D., and Legendre, P. (2002). All-scale spatial analysis of ecological data by means of principal coordinates of neighbour matrices. Ecological Modelling 153, 51–68. doi: 10.1016/s0304-3800(01)00501-4.

80. Egue, F., Chenais, B., Tastard, E., Marchand, J., Hiard, S., Gateau, H., Hermann, D., Morant-Manceau, A., Casse, N., and Caruso, A. (2015). Expression of the retrotransposons Surcouf and Blackbeard in the marine diatom *Phaeodactylum tricornutum* under thermal stress. Phycologia 54, 617–627. doi: 10.2216/15-52.1.

81. Pargana, A., Musacchia, F., Sanges, R., Russo, M.T., Ferrante, M.I., Bowler, C., and Zingone, A. (2020). Intraspecific diversity in the cold stress response of transposable elements in the diatom *Leptocylindrus aporus*. Genes 11, 9. doi: 10.3390/genes11010009.

82. Hong, T., Mo, J., Li, T., Huang, N., Liu, W., Liang, H., Pei, P., Li, P., Chen, J., and Du, H. (2025). Multi-omics analysis reveals adaptation strategies of marine diatom to long-term ocean warming: resource allocation trade-offs and epigenetic regulation. Plant, Cell & Environment 48, 5089–5103. doi: 10.1111/pce.15482.

83. Song, B., Morse, D., Song, Y., Fu, Y., Lin, X., Wang, W., Cheng, S., Chen, W., Liu, X., and Lin, S. (2017). Comparative genomics reveals two major bouts of gene retroposition coinciding with crucial periods of *Symbiodinium* evolution. Genome Biology and Evolution 9, 2037–2047. doi: 10.1093/gbe/evx144.

84. Reagan, J.R., Boyer, T.P., Garcia, H.E., Locarnini, R.A., Baranova, O.K., Bouchard, C., Cross, S.L., Mishonov, A.V., Paver, C.R., Seidov, D. et al. (2024). World Ocean Atlas 2023. NOAA National Centers for Environmental Information. URL: https://www.ncei.noaa.gov/access/world-ocean-atlas-2023/. Accessed 2026-03-20.

85. de Boyer Montégut, C., Madec, G., Fischer, A.S., Lazar, A., and Iudicone, D. (2004). Mixed layer depth over the global ocean: An examination of profile data and a profile-based climatology. Journal of Geophysical Research: Oceans 109, C12003. doi: 10.1029/2004JC002378.

86. Worden, A.Z., Follows, M.J., Giovannoni, S.J., Wilken, S., Zimmerman, A.E., and Keeling, P.J. (2015). Rethinking the marine carbon cycle: Factoring in the multifarious lifestyles of microbes. Science 347, 1257594. doi: 10.1126/science.1257594.

87. Rost, B. (1999). Twilight zone of protein sequence alignments. Protein Engineering, Design and Selection 12, 85–94. doi: 10.1093/protein/12.2.85.

